# Computing pathogenicity of mutations in human cytochrome P450 superfamily

**DOI:** 10.1101/2025.02.07.637054

**Authors:** Somnath Mondal, Pranchal Shrivastava, Rukmankesh Mehra

## Abstract

Cytochrome P450 (CYP) are heme-containing enzymes involved in the metabolism of drugs and endogenous compounds. Humans have 57 known CYPs, with >200 mutations linked to severe disorders, highlighting their role in disease etiology. To investigate their pathogenicity, we performed an in-depth computational study, comparing pathogenic mutations with non-pathogenic ones. First, we evaluated the effects of all possible mutations across 26 known CYP’s structures using five structure-and five sequence-based methods, aiming to reproduce pathogenesis. We computed 23,46,410 mutations, categorized into: all possible, non-pathogenic, and pathogenic. Comparisons consistently revealed a meaningful stability pattern: non-pathogenic > all > pathogenic. Second, we found that a significantly higher number of pathogenic mutations were buried in CYP structures, relating to their higher pathogenesis potential. Third, we analyzed mutation allele frequency relative to solvent accessibility. Fourth, we computed change in 48 amino acid properties upon mutation and identified three distinguishing features: Gibbs free energy, isoelectric point and volume. Fifth, the positive residue content was reduced significantly in diseased mutations, with arginine mutations being the main culprit, directly linked to isoelectric point change. Sixth, we found a higher propensity for pathogenic mutations in conserved sites, suggesting disruption of CYP function. Finally, analysis of heme versus substrate sites showed a higher frequency of pathogenic mutations in heme site, with arginine being the main mutating residue, possibly disrupting arginine-heme interactions. We provide the first comprehensive analysis of mutation effects across multiple CYPs. We model the chemical basis of CYP-related pathogenicity, paving way for a semi-quantitative model to predict diseases.

## 1. Introduction

The human cytochrome P450 (CYP450 or CYP) superfamily is a broad and diverse group of 57 enzymes, predominantly located in the liver but also present in other tissues [1–4]. These enzymes are crucial for metabolizing a wide range of substrates, including endogenous compounds like steroids, fatty acids, and vitamins, as well as exogenous substances such as drugs and toxins [5–7].

CYP450 enzymes are systematically classified into families and subfamilies according to the amino acid sequence identity [8]. Enzymes with ≥40% sequence identity are placed in a family (labeled with a number), while those with ≥ 55% identity are grouped into a subfamily (denoted by a letter). For instance, the enzymes sterol 27-hydroxylase and vitamin D3 24-hydroxylase share ≥40% sequence identity and therefore, placed in single family CYP27. However, since their identity is <55%, they are classified into separate subfamilies: CYP27A and CYP27B, respectively.

Mutations in CYP genes can greatly impact drug metabolism, influence disease risk, and affect other physiological functions [9,10]. This is due to their critical role in metabolism. CYP450 enzymes oxidize organic molecules, typically by adding an oxygen atom to the substrate. This function is essential for metabolizing a wide range of substances, including drugs, toxins (such as pollutants and carcinogens), and endogenous compounds (like fatty acids, cholesterol and steroid hormones) [11,12]. For instance, the enzymes from CYP1, CYP2, and CYP3 families metabolize about 80% of clinically prescribed drugs. CYP2D6 processes around 20% of these drugs, while CYP3A4 and CYP3A5 together metabolize 30%. The CYP4 family is involved in fatty acid metabolism, while the CYP8 and CYP27 families are implicated in bile acid synthesis. The CYP11 and CYP21 families contribute to steroid biosynthesis, and the CYP24 family is responsible for the inactivation of vitamin D [13]. These functions underscore the critical role of CYP450 enzymes in the human body.

CYP450 mutations cause several functional consequences including: (a) loss of function mutations, which reduce or eliminate enzyme activity, impairing the processing of endogenous substances; (b) gain of function mutations, which result in overactive enzymes, and accelerate drug metabolism, leading to an increased production of certain metabolites; and (c) polymorphisms, genetic variations that result in differing enzyme activity between individuals, often affecting their response to drugs [14–16]. Examples of CYP1A2, CYP2D6 and CYP2C19 illustrate how polymorphisms can result in either slow or rapid metabolism of substrates [17–19]. For example, CYP1A2 processes substances such as caffeine, carcinogens and theophylline. Mutations in this enzyme may cause slow metabolism, causing elevated caffeine levels that increase the risk of side effects like anxiety or heart palpitations [20]. On the other hand, certain CYP1A2 mutations may speed up caffeine metabolism, reducing its effectiveness [21,22]. Similarly, CYP2D6 is responsible for metabolizing a variety of drugs, including opioids, antipsychotics, and antidepressants [20]. Mutations in this enzyme can lead to poor metabolism of certain drugs, resulting in toxic drug levels [23]. While some CYP2D6 mutations may cause rapid metabolism, leading to subtherapeutic drug levels [24,25]. These variations highlight the importance of understanding the structure-function relationships of CYP450 enzymes.

Therefore, we asked a question of how a single amino acid change in CYP450 enzymes can be sufficient to trigger pathogenesis. Additionally, we aimed to determine whether the pathogenic effects of these mutations could be predicted or quantified based on the simple chemical properties of the amino acid alterations. To explore this, we reviewed state-of-the-art computational tools designed for rapid and efficient analysis of the potential functional impacts of missense mutations [26–28]. We then applied these tools to known missense mutations in CYP450 enzymes that are associated with pathogenesis, which are central to current research on CYP450-related etiologies. Mutations that alter protein folding stability can reduce the amount of functional protein (loss of function) or induce malicious misfolding (gain of function) [29,30]. Pathogenic missense mutations often disrupt protein interactions and stability, and can also impact other key factors such as metabolic activity, substrate binding, protein charge, solubility, and protein signaling or processing [31–38]. These effects are typically assessed through either structure-based methods, which rely on 3D protein structures or the amino acid sequences as inputs [39–41].

We performed the first most comprehensive analysis to date of mutation datasets across all CYP450 enzymes. Structures of twenty-six CYP450 enzymes were known in Protein Data Bank **(Table S1)** [42], and we used these structural data as input for our analysis. We investigated whether ten distinct computational methods could quantitatively assess the pathogenicity of CYP450 mutations, employing an analysis of variance (ANOVA) test across all possible mutations in these 26 different CYP isoforms. We also tackled the more complex task of distinguishing pathogenic from non-pathogenic mutations by using the area under the curve (AUC) of receiver operating characteristic (ROC) curves for all 26 CYPs across ten methods. These analyses were performed in parallel, using combined datasets from multiple CYPs to address dataset imbalance. We show that some methods effectively capture key features that differentiate pathogenic from non-pathogenic mutations, but our combined analysis of mutation datasets from multiple CYPs highlighted clear patterns of mutation effects across nine of the methods. Through detailed computational analysis across CYP enzymes, we find that arginine is an essential residue that is more prevalent in the heme binding cavity than substrate site and mutation of this residue causes serious consequences by significantly affecting the enzyme charge and disrupting the arginine interaction with heme group. The pathogenic mutations are predominant in the conserved protein regions compared to non-pathogenic. Our findings suggest a path toward creating unified, semi-quantitative chemical models for all human CYP450 enzymes, with potential predictive and clinical applications.

## 2. Methods

### 2.1. Hypothesis

CYP450 enzymes play a critical role in drug and xenobiotic metabolism. Numerous mutations have been reported that either disrupt their activity or are benign. Single nucleotide polymorphisms (SNPs) accumulate in the population, leading to protein mutations, most of which have no adverse effects, but some may impair protein function. Knowing the impact of these mutations is central to grasping their mechanism, broad substrate specificity, drug metabolism and drug-drug interactions. The CYP3A4 isoform, which metabolizes approximately 50% of marketed drugs [43], is the primary drug-metabolizing enzyme, although other CYP enzymes also contribute. Literature studies have focused on the impact of point mutations in CYP3A4 [44]. Surprisingly, only a single pathogenic mutation has been identified in CYP3A4, which might be due to the heavy dependency of metabolizing diverse substrates on this enzyme, suggesting that even a minor alteration to this enzyme could be toxic. Other CYP isoforms exhibit multiple natural mutations, including both pathogenic and non-pathogenic variants, as shown in our compiled data (**Table S2** and **Figure S1**). The effects of these mutations are often closely linked to protein folding stability. A disruption in protein fold stability can compromise its function, making it vital to understand these mechanisms. Therefore, scanning CYP isoforms for the impact of every possible mutation on protein folding stability is crucial for gaining insights into their evolutionary dynamics.

Computational estimates of the stability changes due to mutations are frequently used in protein design and evaluating disease-causing missense mutation, particularly in misfolding-prone proteins [45–48]. Such methods exhibit a decent level of accuracy [49–57], but their performance can vary depending on the mutation dataset. A key reason for this variability is that these methods are often trained on datasets that are imbalanced, with a bias toward predicting destabilizing effects. This bias results in a tendency to predict more destabilizing than stabilizing effects. A potential solution to address this imbalance is to divide the dataset into distinct mutation groups and compare the average effects across these groups. In this study, we conducted a systematic analysis of all possible mutation effects across 26 CYP isoforms (for which structures were available in the Protein Data Bank), using ten diverse sequence-and structure-based methods, and assessed pathogenicity through multiple independent computational approaches.

### 2.2. Datasets of CYP450 genetic variants and protein structures

We collected detailed data of missense genetic variants of all the human CYP450 enzymes from UniProt database [58] and classified the mutations into two categories: pathogenic (disease-causing) and non-pathogenic (non-disease-causing) as shown in **Table S2 and Figure S1**. There exist 316 pathogenic and 25,319 non-pathogenic natural mutations across the 57 human CYP isoforms. CYP2D6 has the highest number of non-pathogenic mutations (711 numbers), while CYP27C1 has only one non-pathogenic mutation. CYP21A2 has the most pathogenic variants, with 68 mutations. CYP3A4, the major drug-metabolizing enzyme, contains one pathogenic and 414 non-pathogenic mutations. These datasets provided a valuable foundation for systematically investigating the effects of mutations on CYP structure and function.

We compiled data on all available CYP450 structures from Protein Data Bank (PDB) [42] including their resolutions and the type of ligands (apo, substrate or inhibitor) bound to them, as presented in **Table S1** and **Figure S1b**. Out of the 57 human CYP isoforms, 26 have known 3D structures, with a total of 255 available structures. These structures were determined using X-ray crystallography, with good resolutions ranging from 1.4 Å to 3.7 Å (**Table S1**). We selected the representative structure for each CYP enzyme based on the highest available resolution. We focused on studying the 26 CYPs with known experimental structures and did not model other CYPs to avoid introducing structural errors into the mutation estimates. Notably, 16 of these 26 CYPs contained only non-pathogenic mutations, while 10 contained both pathogenic and non-pathogenic mutations.

### 2.3. Structure-based mutagenesis methods

Protein function is closely linked to its structure, and mutations can disrupt this function by altering protein folding stability (ΔΔG). To assess the impact of mutations on CYP450 enzymes, we computed ΔΔG using five structure-based methods, each based on different principles: I-MUTANT3.0 [59], mCSM [55], SIMBA [56], POPMUSIC [57], and SNPMUSIC [54] (**Table S3**). I-MUTANT3.0 is a machine-learning method that utilizes a support vector machine, while mCSM employs graph-based signatures. SIMBA uses a multiple linear regression model based on three simple properties, and POPMUSIC applies a mechanistic approach to calculate thermodynamic stability changes due to single-site mutations. SNPMUSIC uses solvent accessibility-dependent combinations of statistical potentials and artificial neural networks for prediction. Default parameters were used to compute ΔΔG (in kcal/mol) for all five methods.

We computed a total of 11,40,135 mutations across the 26 CYPs using the five structure-based methods. Since different methods use varying sign conventions to label ΔΔG values as stabilizing or destabilizing, we standardized the convention for this study. Specifically, we defined ΔΔG < 0 as indicating a destabilizing effect and ΔΔG > 0 as indicating a stabilizing effect.

### 2.4. Sequence-based mutagenesis methods

To investigate the mutation effects based on the CYP sequences (UniProt IDs in **Table S1**), we performed saturated mutagenesis using five sequence-based methods: ENVISION [52], META-SNP [49], SNAP2 [50], SHIFT [53], FATHMM [51] with default parameters (**Table S3**). These methods leverage amino acid conservation derived from multiple sequence alignments to predict the impact of missense mutations [60]. By considering the physicochemical properties of amino acids and potential structural alterations, these methods estimate mutation effects based on their types. We computed 12,06,275 mutations using the sequences of the 26 CYPs, and the resulting scores were used to interpret the mutation effects.

### 2.5. Residue classification based on solvent exposure

To identify the locations of natural mutations on the CYP structures, we calculated the relative solvent-accessible area (RSA) of the residues for ten enzyme structures with known pathogenic mutations using I-MUTANT3.0 [59]. Residues were categorized as buried or solvent-exposed based on an RSA threshold of <20% for buried residues, as previously defined [61].

### 2.6. Computing amino acid property change, and allele frequency

We collected allele frequency data for the missense mutations in CYPs from gnomAD (detailed in **supplementary section 1 of supplementary data**) [62]. Additionally, we retrieved data on 48 properties of the twenty standard amino acids [63] (**Table S4**) and computed the change in property (ΔP) upon mutation using the formula *ΔP = Pmutant – Pwild*. The allele frequencies and ΔP values were then correlated with the computed mutation effects.

**Figure 1.**
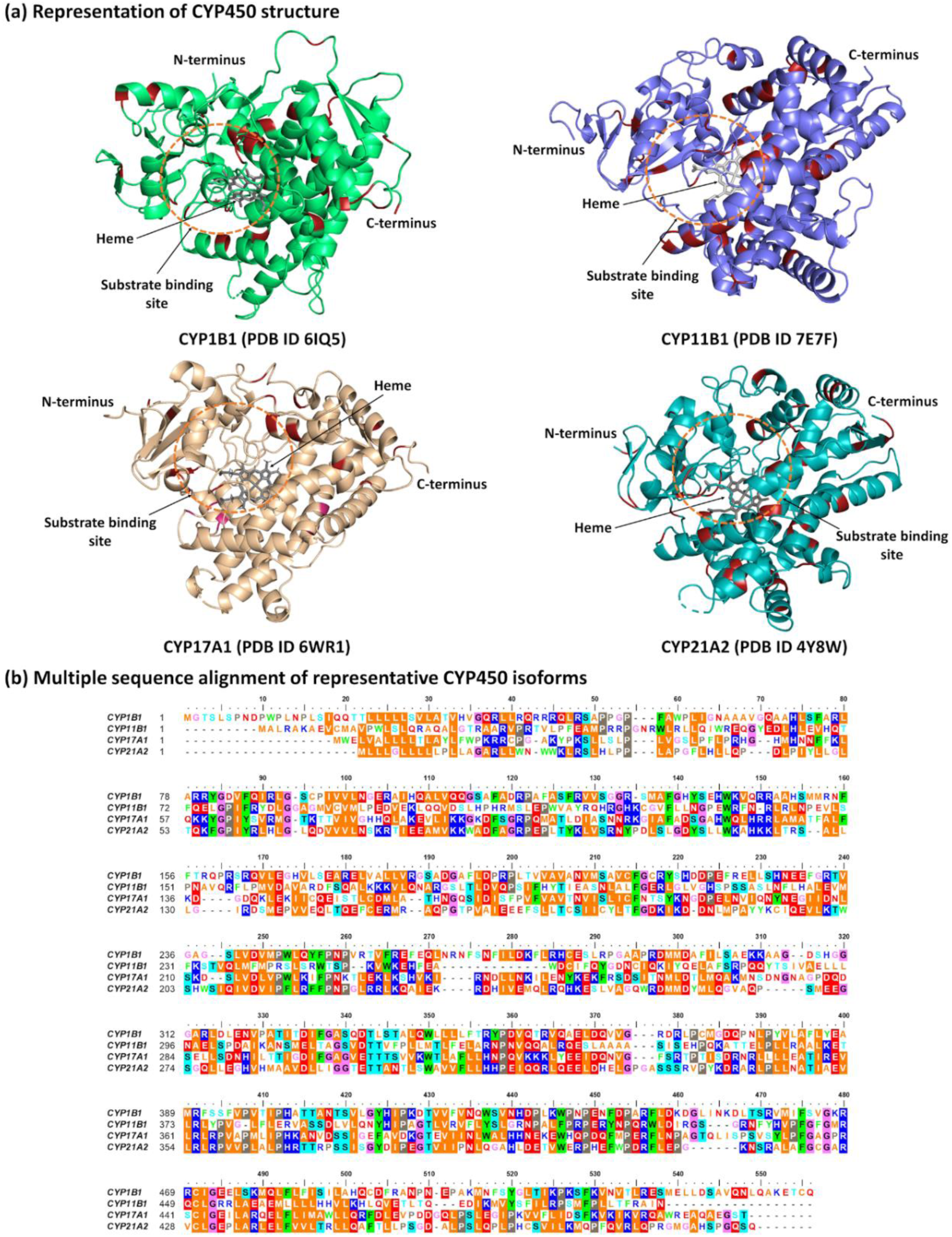
Representation of four CYP450 enzymes having dominant pathogenic mutation datasets. **(a)** CYP structures showing positions of pathogenic mutation in red color. **(b)** Sequence alignment of four CYPs using Clustal Omega [64]. Rsidues are colored based on their physiochemical properties. Aliphatic, aromatic, phosphorylatable, proline, glycine, positively and negatively charged residues are colored as orange, green, cyan, gray, pink, blue, and red respectively. Similar group of residues are represented in similar background color.

### 2.7. Statistical analysis and group comparisons

We employed ten diverse methods to create a comprehensive landscape of saturation mutagenesis, computing all possible mutations at each site of the enzyme. For analysis, the computed mutations were categorized into three groups: ‘all mutation’, ‘non-pathogenic’, and ‘pathogenic.’ ANOVA tests were conducted to examine significant differences between these mutation groups.

The area under the curve (AUC-ROC) values were plotted to evaluate the performance of the ten methods in classifying non-pathogenic and pathogenic mutations. The AUC-ROC curve was generated by plotting the true positive rate (TPR) against the false positive rate (FPR) using the Scikit-learn module in Python 3.7.8 [65], as described in **Supplementary Section 2**.

### 2.8. Calculation of conservation score and binding site analysis

The conservation of amino acid residues in the 26 enzyme sequences was analyzed using a script available at https://github.com/esbgkannan/kibby [66]. This approach estimates the conservation score based on sequence embeddings generated by protein language models like ESM2. These models, trained on extensive protein sequence datasets, learn representations (embeddings) that capture the underlying structure and function of the sequences. The conservation score is derived by comparing the sequence embeddings to those of other known sequences, bypassing the need for traditional sequence alignment [66]. Additionally, the Conserved Domain Database (CDD) was used for Batch CD-search with default parameters to identify conserved heme-and substrate-binding residues in CYPs [67].

## 3. Result and discussion

### 3.1. Estimation of pathogenicity in CYP450 variants

Missense mutations in CYP450 enzymes can impair their monooxygenation function, which may cause dysfunction in the metabolism of xenobiotics, steroids, fatty acids, and other compounds [3]. While random mutations are usually prone to reduce protein thermodynamic stability, mutations in the enzyme’s active site can specifically compromise its activity. Altering wild-type residues can shift the equilibrium between the folded and unfolded protein states toward the unfolded configuration [68,69]. We investigated whether sequence-based disease predictors, which incorporate evolutionary conservation data, provide a more accurate representation of pathogenicity compared to the thermodynamic instability of the folded state. Considering chemical alterations, these techniques measure the impact of mutation on protein function [70,71]. Relying on just one or two computational approaches may not be recommended due to their inherent error-prone nature, as results may be influenced by the choice of method used [72–78].

Figure S1a illustrates the number of mutations, and **Table S2** lists all mutations (non-pathogenic and pathogenic) found in each of the 26 CYPs. Pathogenic mutations were reported in only ten CYPs. Detailed mutation analysis using five structure-based methods is presented in Figures 2 and **S2-S6**. We compared the average effects of three mutation groups, namely, all possible, non-pathogenic and pathogenic to draw our conclusions. The analysis based on the single values of point mutation may be irrelevant because the major challenge with these methods is the unequal distribution of the training dataset. Although conclusions can be made based on the average values of each group, the stabilizing effects remain unclear. It is only through a comparative analysis of these groups that meaningful insights can be drawn.

**Figure 2.**
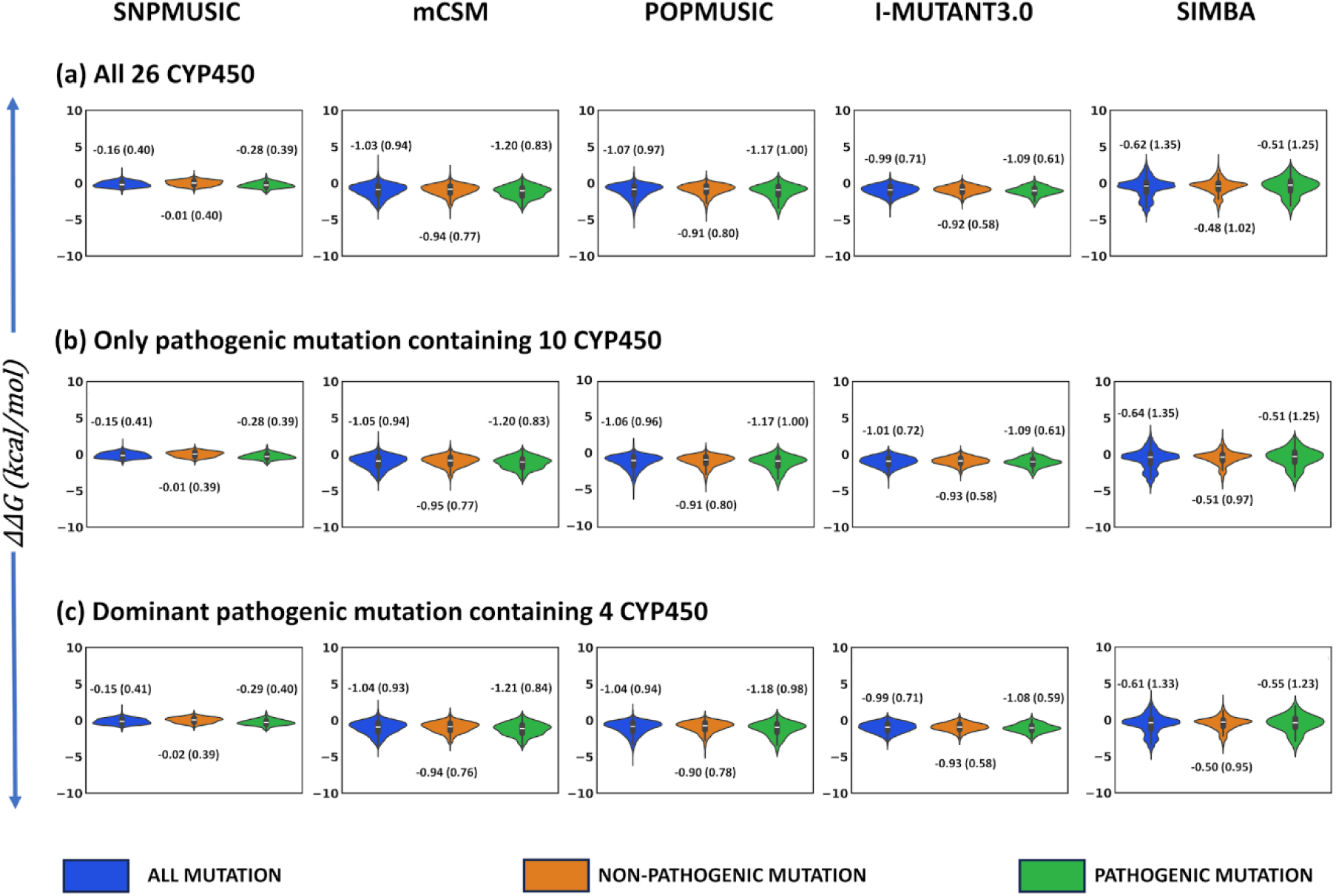
**Violin plots of the mutation groups comparisons using combined datasets of enzymes for the five structure-based methods**. The average and standard deviation (in brackets) for each group is shown near each violin. **(a)** Combined mutation dataset of 26 CYP450 for which natural mutations were known. **(b)** Combined mutation dataset of 10 CYP450 for which pathogenic mutations were known. **(c)** Combined dataset of 4 CYP450 for which the prominent pathogenic sets were available.

The comparisons between the three mutation groups revealed significant differences in the average effects using SNPMUSIC (**Figure S2**), as confirmed by ANOVA tests (**Table S5**). In SNPMUSIC, the overall pattern showed that pathogenic mutations tend to reduce protein stability more than non-pathogenic or all mutations (**Figure S2 and Table S6**), indicating that the pathogenic mutations may cause deleterious effect due to decreased stability. The average ΔΔG values followed the trend: non-pathogenic > all > pathogenic. In all 26 comparisons, the ΔΔG values for the non-pathogenic group were higher than those for the other groups, suggesting that non-disease-causing mutations are less likely to destabilize the protein. This means that these non-pathogenic mutations, which are favored by natural selection, do not significantly disrupt CYP stability or function, allowing them to accumulate in large numbers over evolutionary time. In contrast, pathogenic mutations, which are associated with the highest destabilization, are less favorable and therefore occur in fewer numbers.

The effect was particularly pronounced in the four CYPs with the highest number of pathogenic mutations, i.e., CYP1B1, CYP11B1, CYP17A1 and CYP21A2, each having more than 25 pathogenic mutations. However, the remaining six CYPs had fewer than 10 pathogenic mutations, and CYP3A4 and CYP8A1 each had only a single pathogenic mutation. As a result, comparing the average pathogenic effect for these CYPs with the larger mutation groups (including all possible and non-pathogenic) may not provide meaningful insights. Therefore, we analyzed data for each of five methods in three ways by combining mutation datasets of: (a) all 26 CYPs for which natural mutations were known, (b) 10 CYPs for which pathogenic mutations were known, and (c) 4 CYPs with the largest pathogenic mutation sets (Figure 2). Thus, to address the imbalance in mutation group sizes across the 26 CYPs, we also analyzed the combined datasets for the 10 and 4 CYPs separately.

Consistently, all three comparisons revealed significant stabilizing effects in the order of non-pathogenic > all > pathogenic (ANOVA test in **Table S5**), completely in agreement with the SNPMUSIC results for individual enzymes. These findings were clearly supported by the other three methods mCSM, POPMUSIC, and I-MUTANT3.0, as shown in Figure 2 (ANOVA tests in **Tables S5**). The differential effects were more dominant in the combined datasets compared to the analysis based on individual enzymes (**Figure S3-S5** and **Tables S7-S9**). Although SIMBA exhibited a slightly different stabilization pattern of non-pathogenic > pathogenic > all (Figure 2 and **Table S5**), it still indicated that non-pathogenic mutations are the most stabilizing (individual enzyme analysis shown in **Figure S6** and **Tables S10**).

All the above observations clearly indicate that the methodology we adopted, which involves analyzing combined datasets from multiple enzymes within the same superfamily, can reveal significant trends that are consistent across different methods.

In our case, four methods demonstrated a clear and meaningful stabilization pattern when mutation datasets from multiple enzymes were combined. This finding supports the conclusion that pathogenic mutations in CYP450 enzymes may lead to destabilization by shifting the equilibrium toward the unfolded state. While any mutation can potentially destabilize the enzyme by promoting an unfolded state [68,69], this effect is more pronounced in the pathogenic group compared to the non-pathogenic and all mutations groups, where the destabilizing effect is relatively milder.

Surprisingly, all five sequence-based methods consistently showed a significant differential trend in the group comparisons, strongly correlating with the results from the structure-based methods (**Figure S7-S11**). For individual CYP analysis, all methods displayed a general order of mutation impact from neutral to deleterious as non-pathogenic < all < pathogenic groups. The significant differences between the averages were checked using ANOVA tests (p-value < 0.05; **Tables S11-S15**). Clear differential effects were observed in the four CYPs with prominent pathogenic datasets: CYP1B1, CYP11B1, CYP17A1 and CYP21A2.

As with the structure-based methods, we also analyzed the sequence-based methods by combining mutation data from multiple enzymes in three different ways (Figure 3). This approach revealed consistent patterns, with mutations showing a clear order from neutral to deleterious as non-pathogenic < all < pathogenic groups across all methods (ANOVA tests in **Tables S16**). These results fully align with the findings from the structure-based methods. Notably, the effects became more prominent when we compared the combined mutation datasets from multiple enzymes. Even with the unbalanced dataset of 26 CYPs, clear trends emerged, which were further validated by narrowing down the analysis to ten and then four enzymes. This methodology strengthened our confidence in the results, as it consistently led to meaningful conclusions across multiple methods, CYP enzymes, and both structure-and sequence-based analyses.

**Figure 3.**
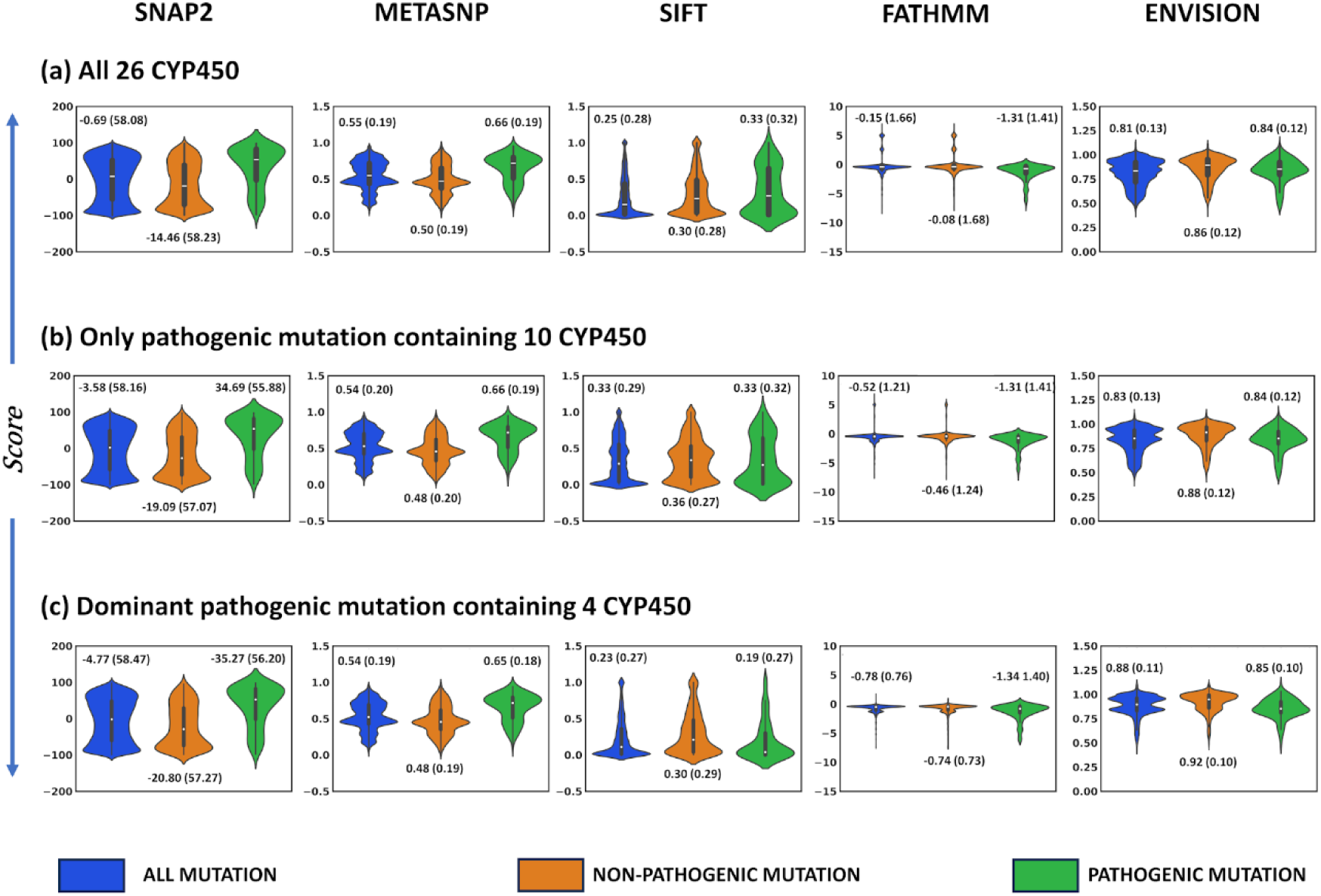
**Violin plots of the mutation groups computed using SNAP2, METASNP, SIFT, FATHMM, and ENVISION**. The average and standard deviation (in bracket) for each group is shown near each violin. **(a)** Combined mutation dataset of 26 CYP450 for which natural mutations were known. **(b)** Combined mutation dataset of 10 CYP450 for which pathogenic mutations were known. **(c)** Combined dataset of 4 CYP450 for which the prominent pathogenic sets were available.

These analyses suggest that the non-disease-causing natural mutations appear like wild-type, while pathogenic mutations exhibit deleterious effects. These effects can be effectively derived from the amino acid sequence information of the CYPs. Sequence-based methods outperformed structure-based methods for individual enzymes, indicating that the CYP sequences contain sufficient information to identify pathogenic variants. Since the amino acid sequence is linear, extracting patterns using sequence conservation, known variants, and machine learning models is relatively simpler and can provide valuable insights for predicting pathogenesis.

Protein structural methods rely upon the experimental dataset for training and 3D protein structures, which could be limiting factors [79,80]. Incorporating the effect of the residue conformations distant from the variant under scanning might also be the limiting factor and may introduce errors. The quality, domain integrity, and completeness of the protein structures are also critical factors that could impact the accuracy of these methods.

However, when a significant size of the datasets for the three groups were compared, the effects became predominant for both sequence-and structure-based methods, with both consistently revealing similar major trends. Our approach of combining mutation datasets from multiple CYPs (in Figure 2) successfully identified the meaningful patterns in the average behaviors.

Therefore, the significant findings from the group comparisons indicate that the pathogenic mutations showed a deleterious effect, which could be linked to changes in protein stability. Specifically, decreased stability in pathogenic mutations may lead to pathogenesis, whereas the non-pathogenic mutations exhibited relatively stable behavior, resembling wild-type residues. This could explain why numerous non-pathogenic mutations occurr naturally in CYPs without disrupting enzyme function, leading to their accumulation over time. In contrast, disease-causing mutations are less frequent, occurring only in certain CYPs, indicating that these residues play a critical role in maintaining enzyme stability and function. We propose that pathogenic mutations exert their effects by destabilizing protein folding.

### 3.2. Distinct behaviors of buried vs. exposed residues in pathogenic and non-pathogenic datasets

Previous studies have suggested the tendency of the pathogenic mutations to be present in the buried region, and surface site of the protein may involve in the molecular interactions that regulate the protein stability and function [60,72,81]. To investigate this, we computed the buriedness of the mutated residues in the pathogenic and non-pathogenic datasets for the ten enzymes with known diseased mutations. We examined the residue positions present in the selected protein structures. Interestingly, we found that the pathogenic set comprised a significantly higher number of buried residues than exposed, showing their tendency to occur in the buried regions of CYPs (Figure 4a). Specifically, 136 buried and 76 exposed residues were found in the pathogenic datasets of these ten CYPs, indicating a clear preference for pathogenic mutations to occur in buried regions. In contrast, natural mutations that enhance the fitness or stability of the protein are typically found on the protein surface, as previously observed [72,76]. While the non-pathogenic mutations showed no specific pattern in terms of buriedness (Figure 4b), meaning that a mutation can occur at any location on CYP enzymes, but those in buried regions are more likely to have a deleterious effect. This may be due to their possible role in substrate interaction, domain stability and enzyme function. In the non-pathogenic datasets of the ten CYPs, 1980 buried and 2021 exposed residues were identified.

**Figure 4.**
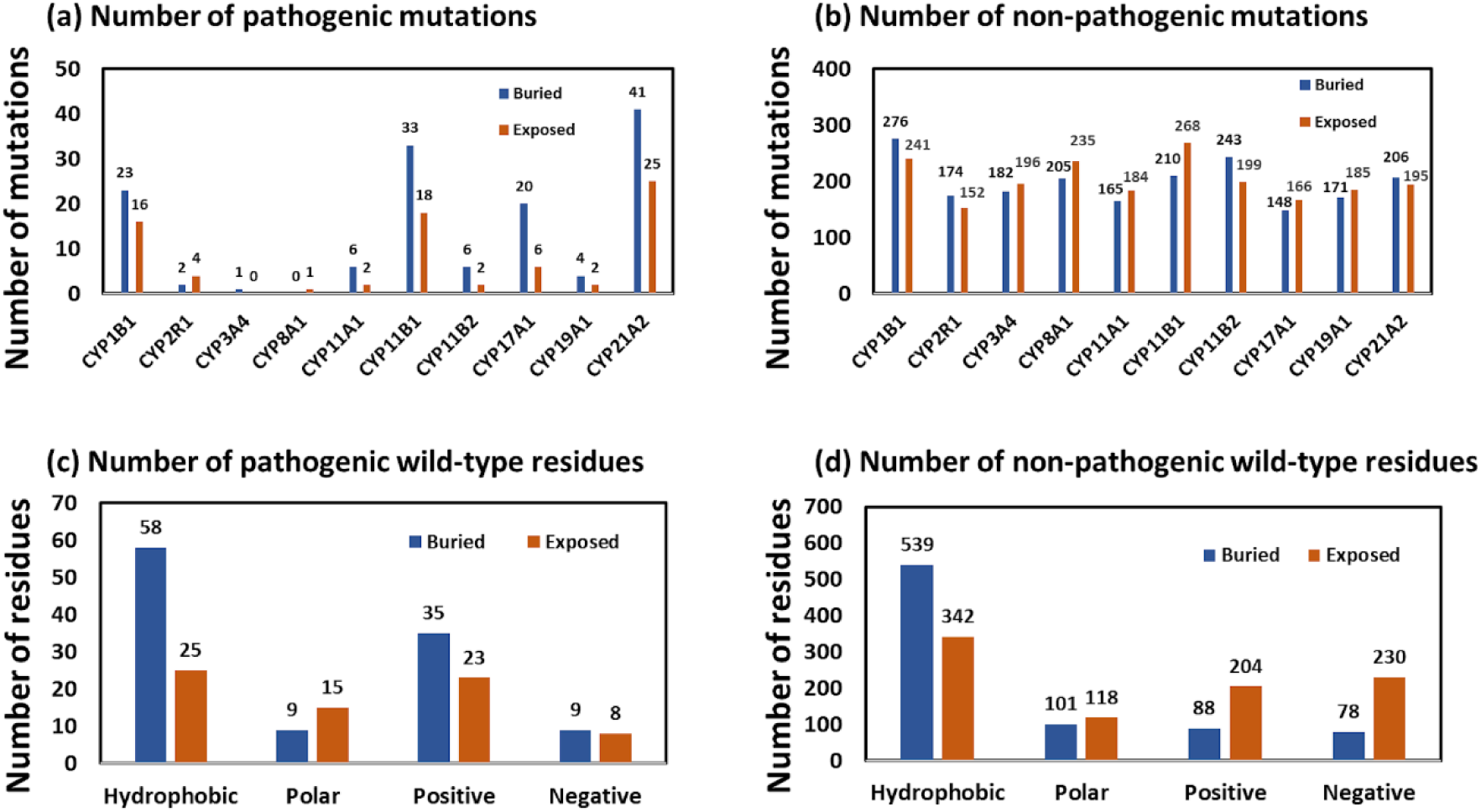
Distribution of pathogenic and non-pathogenic amino acids based on their distinct properties. Number of mutations of ten CYPs into buried and exposed regions forming: **(a)** pathogenic and **(b)** non-pathogenic datasets. Categorization of hydrophobic, polar, positive, and negative wild-type residues for the combined dataset of four CYPs (CYP1B1, CYP11B1, CYP17A1 and CYP21A2) into buried and solvent exposed undergoing: **(c)** pathogenic and **(d)** non-pathogenic mutations.

Understanding the properties of amino acids that are prone to mutation is crucial for assessing their broader effects. For this in-depth analysis, data from four CYPs with prominent mutations (CYP1B1, CYP11B1, CYP17A1 and CYP21A2) were combined and categorized based on the nature of the residues: hydrophobic, polar, positively and negatively charged. The protein core primarily consists of hydrophobic residues, which are necessary for packing, shaping, and maintaining the globular structure of the protein. In contrast, the surface region tends to be enriched in polar residues, as they interact with water molecules. Our analysis confirmed this pattern for CYP enzymes, with hydrophobic residues predominantly found in the buried regions and polar residues in the solvent-exposed areas (Figure 4c and d). Interestingly, the non-pathogenic mutations showed a higher number of charged residues in the solvent-accessible regions, which is expected [82]. However, the pathogenic mutations displayed a higher concentration of charged residues in the buried regions, again highlighting the tendency for disease-causing mutations to occur in these areas of the CYP enzymes. Therefore, mutations of charged residues in exposed regions do not appear to disrupt CYP enzyme function, but such mutations in buried regions may compromise the enzyme’s activity, as also previously reported [82]. Buried charges are expected to play key roles in ligand binding, catalysis or stabilizing structures [82]. The exact impact of such mutations may depend on additional factors, such as the nature of the amino acid substitution, whether the variant maintains or disrupts the charge, and if the mutated residue participates in intermolecular interactions.

Disease-causing mutations are much rarer than benign mutations. While they can disrupt interactions with other molecules when located in buried cavities, they can also occur on the protein surface. Nevertheless, pathogenesis may be governed by multiple factors such as changes in globularity, size, steric effect, intra and intermolecular electrostatic and hydrogen bonding interactions, etc., all contributing to the protein folding stability changes.

### 3.3. Statistical analysis of method accuracy using AUC-ROC

We applied ten methods to assess the pathogenicity of the mutations in 26 CYP450 enzymes and observed that four structural methods (SNPMUSIC, mCSM, POPMUSIC, I-MUTANT3.0) and all the sequence-based methods consistently reproduced similar effects across the three mutation groups (all, non-pathogenic and pathogenic). This consistency suggests that these methods are reliable for computing pathogenicity. These results were well supported by ANOVA test for differences in averages of mutation groups at p-value < 0.05.

To further test the potential of the methods in distinguishing the mutation groups, we performed an area under the receiver operating characteristic curve (AUC-ROC) analysis to complement our ANOVA results. By this point, we had confirmed that the four structural methods and five sequence-based methods consistently captured mutation effects related to folding stability and pathogenesis. We also recognized the imbalance between non-pathogenic and pathogenic datasets, which could potentially skew the trends. To address this, we performed the AUC-ROC analysis separately for individual CYPs (Figure 5a), as well as for combined datasets from ten CYPs (with at least one pathogenic mutation) and four CYPs (with a prominent number of pathogenic mutations) (Figure 5b).

**Figure 5.**
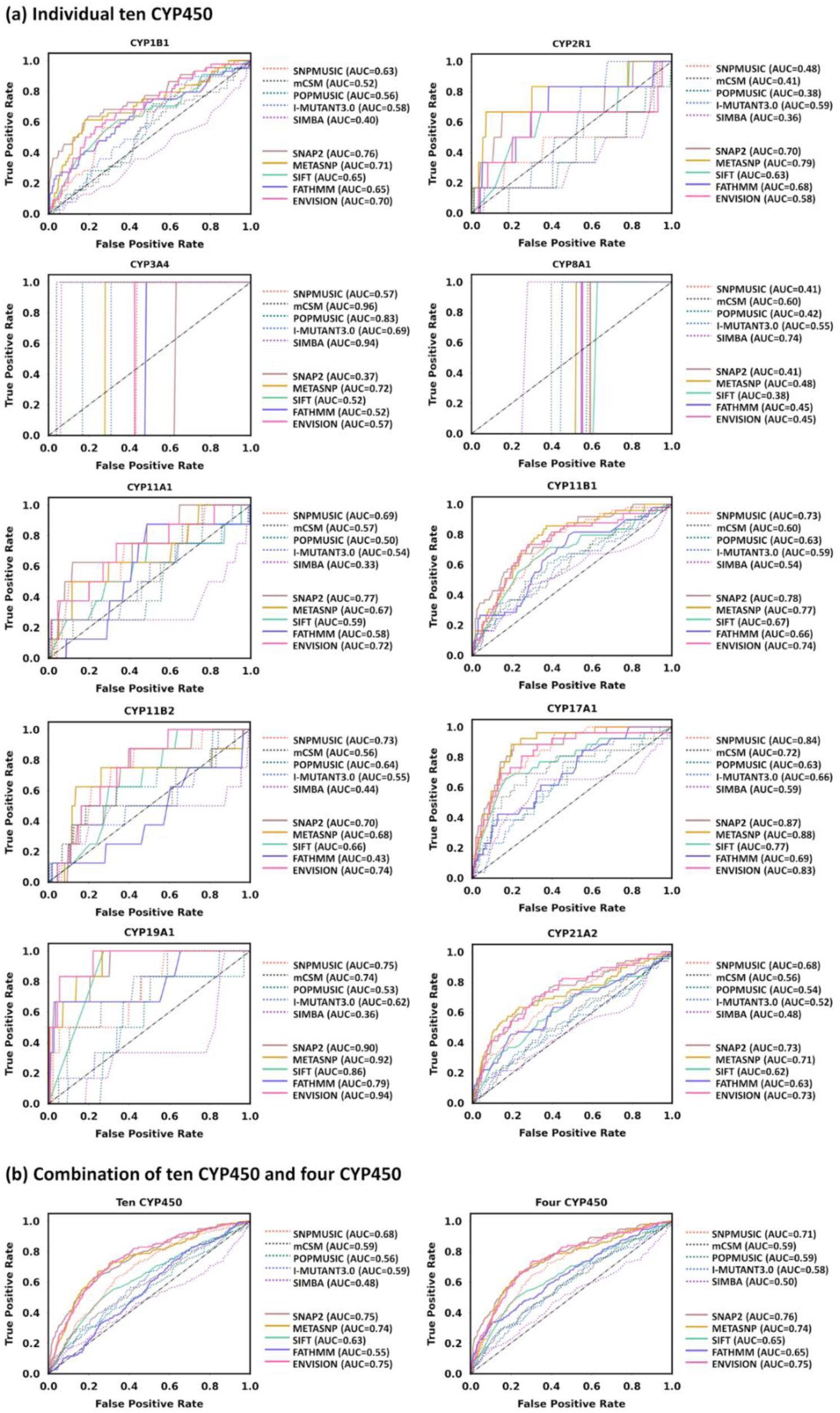
AUC-ROC plots of CYPs for ten structure-and sequence-based methods. **(a)** Individual dataset of ten CYPs. **(b)** Combined dataset of ten CYPs and four CYPs (CYP1B1, CYP11B1, CYP17A1, and CYP21A2). A higher value of area under the curve represents better efficiency of the method in distinguishing pathogenic from non-pathogenic mutations.

We observed decent performance of the nine methods (except SIMBA) (Figure 5b) in distinguishing pathogenic mutations from non-pathogenic ones, consistent with the results from the group comparison analysis. Overall, sequence-based methods outperformed structural methods. Specifically, one structural method, SNPMUSIC, along with three sequence-based methods – SNAP2, METASNP, and ENVISION – achieved AUC-ROC values of ≥0.7, indicating their optimal ability to segregate pathogenic from non-pathogenic mutations. These methods can therefore be confidently used for mutation scanning in CYP450 enzymes.

These results indicate that comparing the averages among mutation groups is an effective approach to differentiate pathogenic mutations from the reference datasets of non-pathogenic and random mutations. This was achieved consistently by mitigating training set biases in computational methods, whether structure-based or sequence-based, which were developed using different approaches. Statistical techniques, such as ANOVA, can validate the significant differences between groups, followed by AUC-ROC analysis to provide a rigorous estimate of the ability to distinguish pathogenic from non-pathogenic mutations. Methods that perform well in both analyses can be used with greater confidence. Although some methods may not perform well in rank predictions (e.g., AUC-ROC), if group comparisons reveal consistent average effects across different methods, it enhances confidence in the findings.

### 3.4. Analysis of allele frequency and change in amino acid properties

With the recent surge in population sequencing data, computational screening approaches have become crucial for interpreting variant phenotypes [83]. This data allows for the evaluation of how natural selection influences missense mutations across the entire allele frequency spectrum. According to clinical diagnostic filtrating criteria, high-frequency mutations are less likely to cause disease [84]. Therefore, we analyzed the allele frequencies of natural mutations in ten CYP450 enzymes with known pathogenic dataset (**Figure S12**).

Both non-pathogenic and pathogenic mutations were randomly distributed, with a few in both datasets exhibiting higher allele frequencies, suggesting that this association may not be well defined in CYPs. For most natural mutations, the same allele frequency was reported (indicated by dots appeared as straight lines in **Figure S12**), resulting in the random distribution of the data. We also analyzed the allele frequency in relation to relative solvent accessibility, SNAP2 score (the top-performing method in ROC analysis), and residue buriedness, as detailed in **Supplementary Figures S13-S15**, which also revealed random behavior. Nonetheless, allele frequency could offer valuable insights when sufficient data of variants and allele frequency are available and should be analyzed in pathogenicity prediction.

We explored the link between pathogenicity and changes in the 48 amino acid properties. This investigation aims to enhance our understanding on how CYP450 mutations contribute to pathogenicity and establish a benchmark for the semi-quantitative approach. We computed the mutation-induced change in property using the equation, *ΔP = Pmut - Pwild* and compared the average effects between the non-pathogenic and pathogenic variants, as shown in box and whisker plots in **Figures S16-S25** and analyzed using ANOVA tests in **Tables S17-S26**. We also correlated ΔP with ΔΔG/score as detailed in **Figure S26-S65** and **Table S27-30**.

Our analysis revealed several significantly different property changes between the two datasets for four CYPs with substantial numbers of pathogenic variants (CYP1B1, CYP11B1, CYP17A1 and CYP21A2) (**Table S31**). Notably, three properties: ΔG, pHi and v emerged as the most important differential descriptors between non-pathogenic and pathogenic mutations across the CYPs in general (found in five CYPs). ΔG is the unfolding Gibbs free energy and related to the protein folding stability, which could be linked with our previous observations on pathogenesis. The average change in ΔG was significantly lower for non-pathogenic than the pathogenic set (**Figure S66a, b**). This indicates that non-pathogenic mutations exhibit greater stability, whereas the pathogenic mutations influence larger changes in ΔG, possibly contributing to their deleterious effects.

pHi describes the isoelectric point of an amino acid, where it has no net charge, plays a critical role in protein folding by influencing solubility and stability at different pH levels. We observed that the average change in isoelectric point was significantly higher for the non-pathogenic dataset compared to the pathogenic one (**Figure S66a, b**). Generally, proteins with a higher isoelectric point tend to have a greater proportion of basic amino acids, such as arginine, lysine, or histidine. As a result, a higher isoelectric point typically indicated that the dataset had a prevalence of basic residues, leading to a net positive charge at pH levels below the isoelectric point. In our case, the non-pathogenic set might exhibit similar behavior more compared to the pathogenic set. It is interesting to observe here that CYPs contain a very high number of arginine residues (as analyzed in next section and Figure 6), which mutate to either pathogenic or non-pathogenic forms. Pathogenesis may occur when the original equilibrium (of high basic residues) is disturbed, altering the charge balance, while this charge is majorly maintained in non-pathogenic mutations.

**Figure 6.**
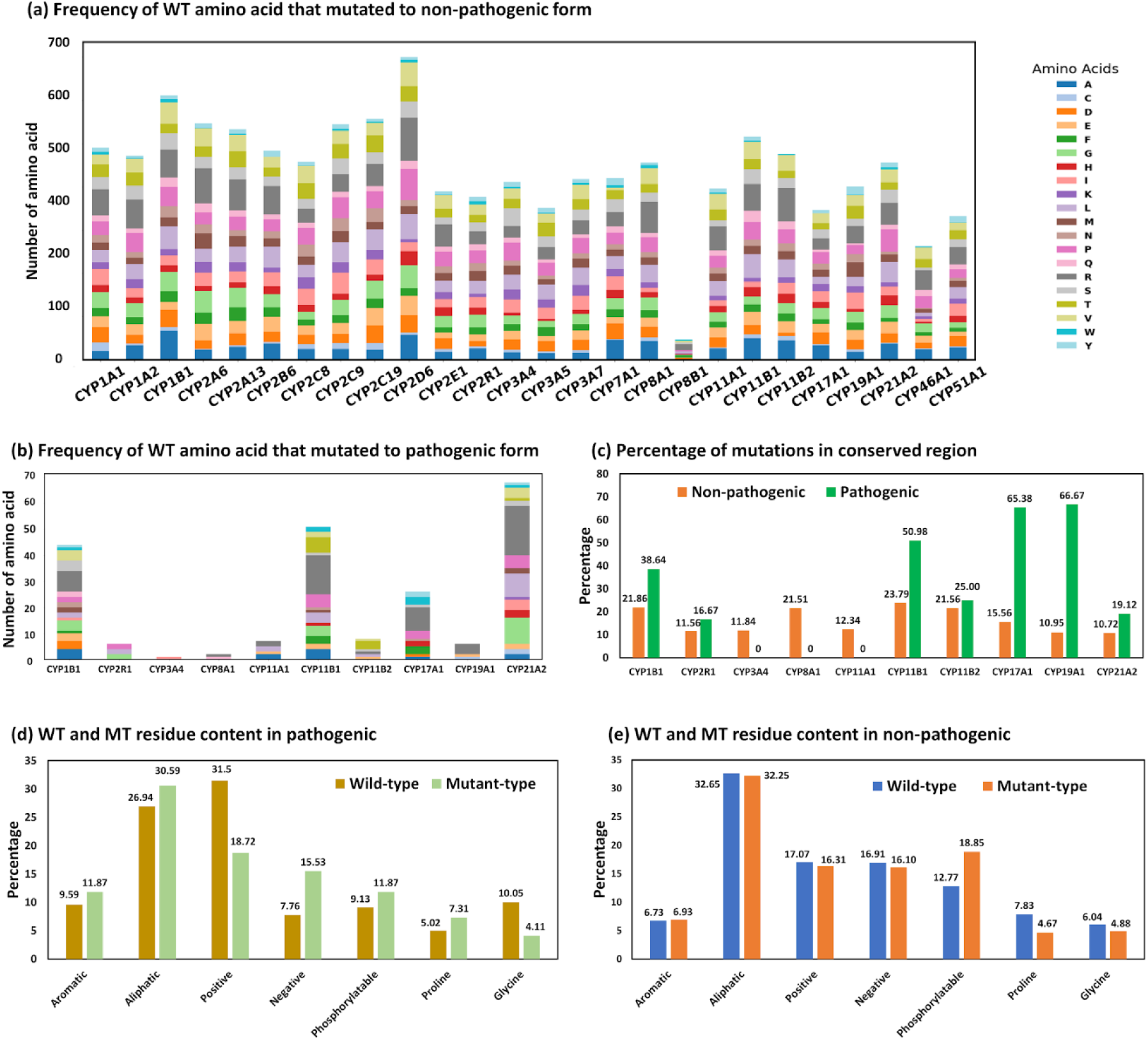
Conserveness and property analysis of the natural mutations in cytochrome P450 enzymes. **(a)** Frequency of the amino acids in the wild-type (WT) enzyme that mutated to pathogenic form. **(b)** Frequency of the amino acid in the wild-type (WT) enzyme that mutated to non-pathogenic form. **(c)** Percentage of mutations (pathogenic and non-pathogenic) in the conserved region of enzymes. The percentage of wild-type (WT) and mutant-type (MT) residues based on their properties for ten CYPs in **(d)** pathogenic and **(e)** non-pathogenic dataset.

The volume (v) of an amino acid, determined by the number of non-hydrogen sidechain atoms, influences protein folding by affecting the packing density and spatial arrangement of the protein structure. The change in amino acid volume showed distinct behavior across the four CYPs (**Figure S66a**), though, overall volume change appears comparatively smaller for non-pathogenic dataset (**Figure S66b**). Substitution by a larger amino acid will cause steric clashes and by a smaller residue will create a hole in the structure, both these events might disrupt the overall stability. Thus, the favorable mutant residues are expected to be of similar size to the wild type. The non-pathogenic mutations caused overall less change in volume than the pathogenic, defining a possible role of volume change in pathogenesis. These observations highlight three potential descriptors that could be crucial to define the pathogenicity across the broader CYP450 family.

### 3.5. Distribution of pathogenic mutation in conserved regions across CYPs

To identify the residues most frequently mutated in cytochromes, a comprehensive analysis of the wild-type amino acids that undergo mutation was conducted (Figure 6). Upon reviewing the occurrences of mutations at each wild-type position, it became evident that arginine was the most frequently observed target for pathogenesis, with a higher propensity for mutation across various cytochromes. This trend was particularly evident in four CYPs: CYP1B1, CYP11B1, CYP17A1 and CYP21A2 (Figure 6b). Arginine is a positively charged residue at physiological pH with a protonated amino group in its side chain and plays a critical role in enzyme function. Mutation at this site can reduce the charge of the protein and potentially disrupt its enzymatic mechanism, especially if the mutation occurs within the substrate or heme binding site of CYP enzymes. Several studies have linked mutations in arginine residues to pathogenic effects in CYP enzymes [85–91]. For instance, mutations in CYP11B1 gene, including p.R448H, are associated with 11β-hydroxylase deficiency (11β-OHD). This condition is a form of congenital adrenal hyperplasia (CAH) and results in impaired cortisol synthesis. Similarly, R181W and V386A mutations in CYP11B2 are associated with the development of 11β-hydroxylase deficiency (11β-OHD). R96Q mutation in CYP17A1 leads to a complete loss of 17α-hydroxylase/17,20-lyase enzyme activity. The R192H variant in CYP19A1 (aromatase) causes virilization in a 46, XX newborn and under-virilization in a 46, XY sibling, without affecting maternal virilization during pregnancy. Additionally, the R124H variant in CYP21A2 is involved in Japanese congenital adrenal hyperplasia (CAH) patients [85–91]. These findings underscore the critical role of arginine substitutions in the pathogenesis of various CYP450-related disorders.

Glycine and proline were also very commonly mutated residues (Figure 6b). Both these residues are typically found in the loop regions of the protein due to their unique geometry. Mutations in these residues may impair the enzyme activity by disrupting the loop conformation, particularly those near the binding site. These observations highlight the potential importance of arginine and additionally glycine and proline in the context of pathogenicity within CYP450 enzymes.

Arginine was also found to be the major mutated residue in the non-pathogenic dataset (Figure 6a). However, in this case the overall isoelectric point or charge of the protein remained relatively unaffected (analyzed in Figure 6d and e), likely due to substitutions with similar residues. This was supported by the smaller change in isoelectric point (ΔpHi) observed in the non-pathogenic mutations compared to the pathogenic ones. These findings suggest that while arginine sites are highly susceptible to mutations in CYP enzymes, pathogenicity is unlikely if the overall charge or isoelectric point is preserved. However, disruptions to these properties may lead to pathogenic outcomes.

Additionally, the non-pathogenic dataset included six general mutating residues: alanine, aspartic acid, glutamic acid, glycine, proline and valine (Figure 6a). Thus, arginine, glycine, and proline are particularly important for analyzing mutations in CYPs to better understand their role in pathogenesis, as further explored below.

We calculated the conservation score of residues involved in both deleterious and non-deleterious mutations using protein sequence embeddings generated from a protein language model [66]. This score represents how conserved a residue is across different protein sequences, indicating its structural or functional importance. A cutoff of ≥0.5 was used to define the highly conserved residues (**Table S32**). Notably, a higher percentage of pathogenic mutations were observed in the conserved regions of the enzymes compared to non-pathogenic mutations (Figure 6c). Across the ten CYPs, 36.53% of pathogenic mutations and 16.77% of non-pathogenic mutations were concentrated in the conserved regions. This suggests that a considerable portion of pathogenic variants (∼36%) may affect the conserved features essential for domain function of CYP enzymes. Conserved regions preserve the essential function of protein during evolution and may be necessary to maintain protein fold stability. Disturbance in these regions is likely to interrupt protein function or stability, which may explain the higher occurrence of pathogenic variants in the conserved regions of CYP enzymes.

The 20 amino acids were classified into seven groups for both pathogenic and non-pathogenic datasets: aliphatic (AVILMC), aromatic (FWY), negatively charged (DENQ), positively charged (HKR), phosphorylatable (STY), proline (P), and glycine (G). In the combined analysis of ten CYPs (with known pathogenic mutations), a thorough comparison of wild-type and mutant residues revealed clear trends (Figure 6d and e). Notably, the fraction of positively charged residues substantially decreased, and the negatively charged residues increased upon mutations in the pathogenic set. This trend is likely related to the crucial role of arginine mutations, which were highly frequent in both pathogenic and non-pathogenic datasets. However, in diseased mutations, the content of positively charged residues was significantly reduced, while it was maintained in the non-pathogenic set.

Similarly, the percentages of aromatic, aliphatic and negatively-charged residues were maintained in the non-pathogenic set, while they increased considerably in the pathogenic set. It means that the deleterious mutations in CYP enzymes may be driven by an increased presence of aromatic, aliphatic or negatively-charged residues, with the most prominent effect appears to be the reduction in positively charged residues.

The proline content increased upon mutations in pathogenic whereas it decreased in non-pathogenic sets. The trends for phosphorylatable and glycine residues were similar across both datasets. These observations suggest that an elevated proline content may be associated with disease-causing mutations, whereas its reduction might not induce pathogenicity. These distinctive features of pathogenic versus non-pathogenic mutations revealed significant trends that contribute to understanding the mechanisms of pathogenesis.

CYP450 enzymes contain a heme group and a substrate binding site, both of which are crucial for the enzyme’s function. The conserved residues within these sites play specific roles in the enzyme’s mechanism. We analyzed the residue composition and conservation of these sites using the CDD database [67] search (**Table S33**) in relation to pathogenic mutations across ten CYP enzymes. Our analysis revealed that a greater number of residues were mutated in the heme-binding site (total 23 mutations) compared to the substrate site (11 mutations) that cause pathogenesis. Arginine was particularly prominent, with seven mutations in the heme site compared to only one in the substrate pocket. This suggests that many pathogenic mutations are likely to affect the heme binding interactions with CYP enzymes rather than the substrate interactions. Given that heme contains two COO^-^ groups that likely interact with the positively charged amine group of arginine [92], disturbances in such electrostatic or salt-bridge equilibrium could disrupt the enzyme’s function.

This could be attributed to the well-defined nature of the heme-binding site versus the broader substrate-binding site, which has the potential to metabolize a wide variety of substrates. This indicates that the heme binding residues have specific roles, and mutations at this site are more likely to disrupt enzyme metabolism, thereby causing pathogenesis. On the contrary, any random mutation in the substrate site is comparatively less likely to restrict CYP enzyme activity due to the vast, flexible binding pocket that can accommodate diverse substrates. This flexibility allows the CYP enzymes to also process multiple substrates simultaneously and in various conformations, often resulting in atypical Michaelis-Menten kinetics.

We observed that typically three to four arginine were in the heme binding site of individual CYPs, which is distinct to the zero to one arginine in substrate site (except CYP3A4 that comprises four arginine). This distinction may be due to the crucial role of arginine in interacting with the heme group. Arginine residues are known to be involved in the structure and function of CYP enzymes [92,93]. The positively charged guanidinium group of arginine potentially interacts with the porphyrin ring of the heme in several CYP enzymes, contributing to the stability of the enzyme’s active site and optimizing its monooxygenation function [92,93].

Thus, the detailed analysis of mutations in CYP450 enzymes led to six key findings associated with their pathogenicity, as summarized in Figure 7. Computational methods proved effective in linking changes in protein fold stability to mutation effects through group comparisons. A series of computational approaches identified several major factors influencing pathogenicity in cytochrome P450 enzymes, including residue buriedness, amino acid property change, mutations at conserved sites, disruption of charge balance, arginine mutations, and distinct mutation contents in heme versus substrate sites. These insights are valuable for predicting the evolution of CYP450 enzymes and understanding their role in disease mechanisms.

**Figure 7.**
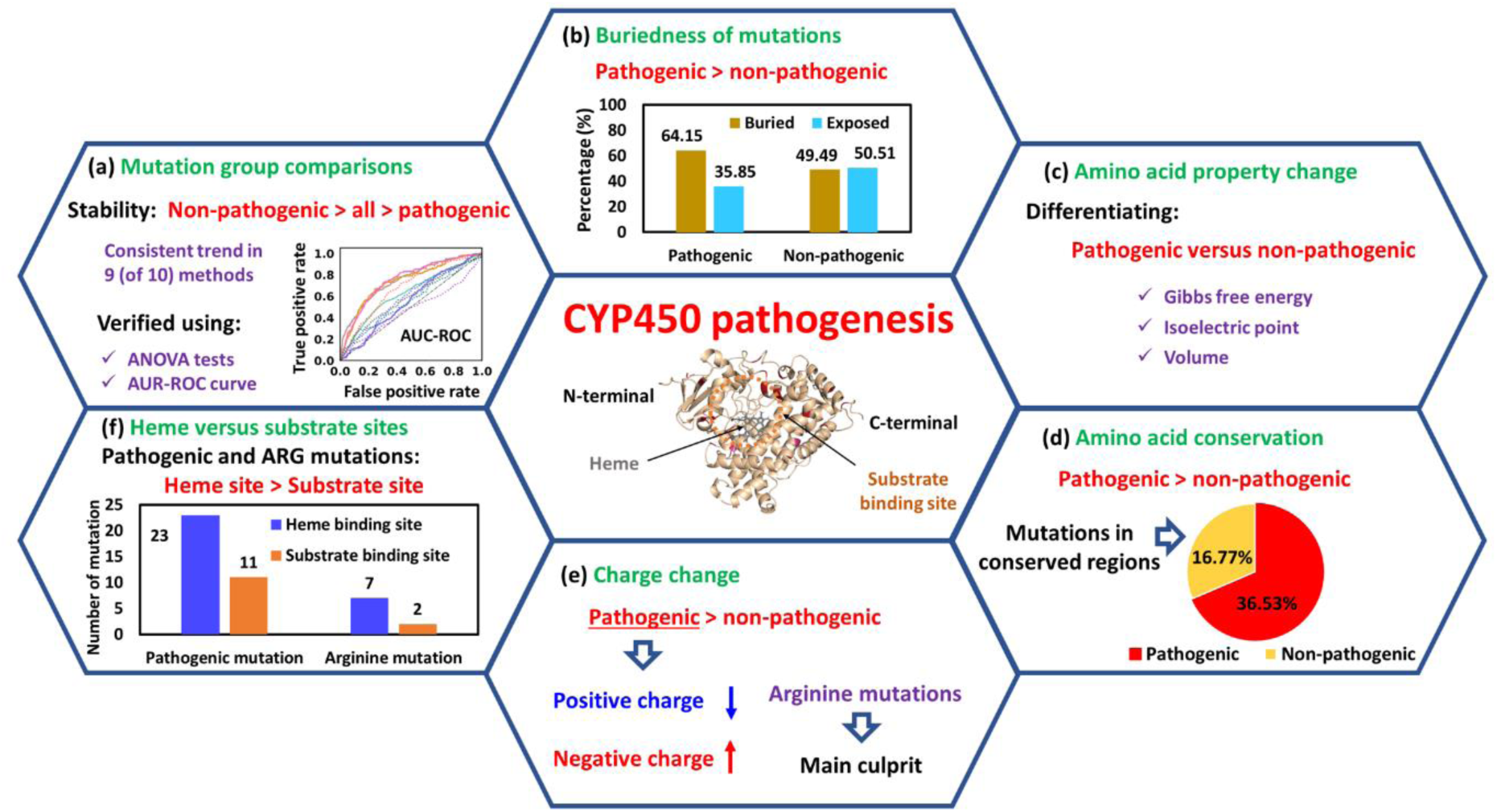
Major outcomes of the current study. **(a)** Comparison of mutation groups. **(b)** Buried mutations were prone to show pathogenic effects. **(c)** Three amino acid properties changes showed significant differences between pathogenic and non-pathogenic datasets. **(d)** Pathogenic mutations were relatively conserved. **(e)** Pathogenic mutations showed relatively greater charge change and arginine is the predominant mutating residue. **(f)** Pathogenic and arginine mutations were more populated in heme-binding site.

## 4. Conclusion

Cytochrome P450 are important class of enzymes primarily located in the liver and involved in the metabolism of wide array of substances including drugs, fatty acids, cholesterol, hormones, toxins and other endogenous compounds. These enzymes constitute a heme binding site and broad substrate cavity. Heme binds to the oxygen species and facilitates electron transfer for substrate oxidation. The substrate or active site catalyze reactions like dealkylation, hydroxylation and demethylation. Mutations in CYP enzymes are linked to pathogenesis by disrupting their metabolic functions. However, several mutations are known to show non-pathogenic or benign effects.

In this study, we thoroughly evaluated the pathogenicity of mutations in 26 cytochrome P450 enzymes (with known PDB structures) using multiple independent computational approaches. Initially, the analysis using four (of five) structure-guided methods consistently revealed a stabilizing pattern upon mutations as: non-pathogenic > all > pathogenic. Similarly, the five sequence-based methods showed consistent order from neutral to deleterious mutations as: non-pathogenic < all < pathogenic groups. This means that the pathogenic mutations might show the deleterious effects by reducing the folding stability more than the reference datasets of all possible mutations and non-pathogenic mutations. The non-pathogenic mutations exhibited the least destabilization (or comparatively increased stabilization) effects, even lower than all possible mutations. This indicates the comparatively higher stability of non-pathogenic mutations that makes them favorable. Such mutations are larger in number than pathogenic, due to their likely benign or increased stabilizing nature. We proposed a method of combining the mutation effects of multiple enzymes belonging to the same superfamily of enzymes to account for the imbalanced datasets. By merging the pathogenic, non-pathogenic, and all possible mutations for the twenty-six, ten, and four CYPs, we identified predominant and meaningful trends, and these results were further validated using ANOVA and AUC-ROC analyses.

Second, we found a significantly higher number of buried residues compared to the solvent exposed in the pathogenic mutations, while there was no such distinction in the non-pathogenic group. This concludes that while mutations can occur at any region of the CYP enzymes, the likelihood of deleterious effects is greater when they occur in the buried regions. Additionally, the content of charged residues was higher in the buried regions for the pathogenic mutations, whereas it was more prevalent in the solvent exposed region for non-pathogenic mutations. This indicates that the charged residue mutations in the buried sites are more likely to disrupt the enzyme’s function, while such mutations in the exposed regions seem to have little impact on CYP enzyme function.

Third, both pathogenic and non-pathogenic mutations displayed random distribution of the reported allele frequencies, with a few in both datasets exhibiting a higher allele frequency, indicating that this association may not be well defined in CYPs.

Fourth, three amino acid property changes, including Gibbs free energy (ΔG), isoelectric point (pHi) and volume (v) were identified as important differential descriptors between non-pathogenic and pathogenic mutations for CYP enzymes in general. ΔG change indicates a comparatively higher stability for non-pathogenic mutations. The isoelectric point change was notably higher for the non-pathogenic dataset than pathogenic dataset, implying a greater prevalence of basic amino acids such as arginine, lysine, or histidine in non-pathogenic mutations. The volume change was comparatively smaller for non-pathogenic mutations, suggesting that favorable mutations are expected to involve residues of a similar size to the wild-type.

Fifth, arginine was found to be the most frequently targeted residue for pathogenesis. CYPs contain a very high number of arginine residues, which are prone to mutation. In pathogenic mutations, the content of positively charged residues significantly decreased. Thus, pathogenesis may result from the disruption of the original equilibrium (of high basic residues) or the positive charge on the enzyme, while benign mutations maintain this charge. Additionally, an increased proportion of aromatic, aliphatic and negatively-charged residues were observed for the pathogenic set, while their content was retained for the non-diseased mutations.

Sixth, we found that pathogenic mutations have a higher propensity of being found in the conserved regions of CYP enzymes than non-pathogenic. This concludes that pathogenicity may result from the disruption of conserved regions that are critical for the enzyme’s essential functions or structural stability.

Seventh, we observed a considerably higher number of pathogenic mutations in the heme-binding site of CYPs compared to the substrate or active site. Here, arginine was a major mutating residue in the heme site, where its mutation could disrupt the interaction between arginine and heme. Notably, three to four arginine were located in the heme across multiple CYPs, distinct from the active site that contained mostly zero to one arginine. This concludes that arginine plays a critical role in interacting with the porphyrin ring of heme, potentially through its positively charged guanidinium group.

In brief, this study showcased the role of computational modeling in elucidating essential structure-function relations of proteins to better understand disease pathogenicity. CYP450 being an important superfamily of enzymes, the current investigations into the pathogenicity of mutations in these enzymes may be of substantial interest in clinical applications.

## Acknowledgements

RM and SM acknowledge SERB-SRG, Govt of India, for funding via grant SRG/2022/000304. RM acknowledges IIT Bhilai for supporting this work via the Research Initiation Grant (RIG), number 2005900. SM acknowledges the Department of Science and Technology, Govt of India, for providing the funds to carry out the research under file no. DST/INSPIRE Fellowship/2023/IF230110.

## CRediT authorship contribution statement

**Somnath Mondal:** Data curation, Formal analysis, Investigation, Software, Validation, Visualization, Writing – original draft. **Pranchal Shrivastava:** Data curation, Software. **Rukmankesh Mehra:** Conceptualization, Formal analysis, Funding acquisition, Investigation, Methodology, Project administration, Resources, Supervision, Validation, Visualization, Writing – original draft, Writing – review and editing.

## Supplementary material

Detailed analyses are provided in the supplementary information file “supplementary.pdf.” The file contains information on allele frequency, statistical analysis, mutation data, enzyme structures, enzyme sequences, protein fold stability methods, amino acid properties, conservation scores, heme and substrate binding sites.

## Data statement

The detailed data is provided in the supplementary file “supplementary.pdf.”

## References

[1] A.C. Drenth-van Maanen, I. Wilting, P.A.F. Jansen, Prescribing medicines to older people— How to consider the impact of ageing on human organ and body functions, Br. J. Clin. Pharmacol. 86 (2020) 1921–1930. 10.1111/bcp.14094.

[2] O.A. Almazroo, M.K. Miah, R. Venkataramanan, Drug Metabolism in the Liver, Clin. Liver Dis. 21 (2017) 1–20. 10.1016/j.cld.2016.08.001.

[3] F. Esteves, J. Rueff, M. Kranendonk, The central role of cytochrome p450 in xenobiotic metabolism—a brief review on a fascinating enzyme family, J. Xenobiotics 11 (2021) 94–114. 10.3390/jox11030007.

[4] S. Deb, S. Arrighi, Potential Effects of COVID-19 on Cytochrome P450-Mediated Drug Metabolism and Disposition in Infected Patients, Eur. J. Drug Metab. Pharmacokinet. 46 (2021) 185–203. 10.1007/s13318-020-00668-8.

[5] P. Manikandan, S. Nagini, Cytochrome P450 Structure, Function and Clinical Significance: A Review, Curr. Drug Targets 19 (2017) 38–54. 10.2174/1389450118666170125144557.

[6] L.L. Furge, F.P. Guengerich, Cytochrome P450 enzymes in drug metabolism and chemical toxicology: An introduction, Biochem. Mol. Biol. Educ. 34 (2006) 66–74. 10.1002/bmb.2006.49403402066.

[7] R. Kumar, S. Kapur, Cytochrome P450 Biocatalysts: A Route to Bioremediation, Available Online Www.Ijpras.Com Int. J. Pharm. Res. Allied Sci. 5 (2016) 113–123. www.ijpras.com.

[8] P. Shahabi, G. Siest, U.A. Meyer, S. Visvikis-Siest, Human cytochrome P450 epoxygenases: Variability in expression and role in inflammation-related disorders, Pharmacol. Ther. 144 (2014) 134–161. 10.1016/j.pharmthera.2014.05.011.

[9] I.M. Langmia, K.S. Just, S. Yamoune, J. Brockmöller, C. Masimirembwa, J.C. Stingl, CYP2B6 Functional Variability in Drug Metabolism and Exposure Across Populations— Implication for Drug Safety, Dosing, and Individualized Therapy, Front. Genet. 12 (2021) 1–21. 10.3389/fgene.2021.692234.

[10] S.C. Preissner, M.F. Hoffmann, R. Preissner, M. Dunkel, A. Gewiess, S. Preissner, Polymorphic cytochrome P450 enzymes (CYPs) and their role in personalized therapy, PLoS One 8 (2013) 1–12. 10.1371/journal.pone.0082562.

[11] D.W. Nebert, K. Wikvall, W.L. Miller, Human cytochromes P450 in health and disease, Philos. Trans. R. Soc. B Biol. Sci. 368 (2013). 10.1098/rstb.2012.0431.

[12] G.W.M. Chang, P.C.A. Kam, The physiological and pharmacological roles of cytochrome P450 isoenzymes, Anaesthesia 54 (1999) 42–50. 10.1046/j.1365-2044.1999.00602.x.

[13] N. Koppel, V.M. Rekdal, E.P. Balskus, Chemical transformation of xenobiotics by the human gut microbiota, Science (80-.). 356 (2017) 1246–1257. 10.1126/science.aag2770.

[14] P. Publisherlocation, B. Central, L. Publisherimprintname, B. Central, A.A. Articlecitationid, A.A. Articlelastpage, R. Onlinedate, A. Articlegrants, Key results, (2002) 12–15.

[15] C. Taylor, I. Crosby, V. Yip, P. Maguire, M. Pirmohamed, R.M. Turner, A review of the important role of cyp2d6 in pharmacogenomics, Genes (Basel). 11 (2020) 1–23. 10.3390/genes11111295.

[16] G.M. Shenfield, Genetic polymorphisms, drug metabolism and drug concentrations., Clin. Biochem. Rev. 25 (2004) 203–6. http://www.ncbi.nlm.nih.gov/pubmed/18458715%0Ahttp://www.pubmedcentral.nih.gov/articlerender.fcgi?artid=PMC1934960.

[17] D. Lesche, S. Mostafa, I. Everall, C. Pantelis, C.A. Bousman, Impact of CYP1A2, CYP2C19, and CYP2D6 genotype-and phenoconversion-predicted enzyme activity on clozapine exposure and symptom severity, Pharmacogenomics J. 20 (2020) 192–201. 10.1038/s41397-019-0108-y.

[18] J.M.J.L. Brouwer, K.J. Wardenaar, I.M. Nolte, E.J. Liemburg, P.M. Bet, H. Snieder, H. Mulder, D.C. Cath, B.W.J.H. Penninx, Association of CYP2D6 and CYP2C19 metabolizer status with switching and discontinuing antidepressant drugs: an exploratory study, BMC Psychiatry 24 (2024) 1–14. 10.1186/s12888-024-05764-6.

[19] J.M. Prather, C.M. Florez, A. Vargas, B. Soto, A. Ross, A. Harrison, A.H. Secrest, D.S. Willoughby, S. Kutter, L.W. Taylor, Impact of CYP1A2 Genotypes on the Ergogenic Effects and Subjective Mood States of Caffeine Ingestion in Resistance-Trained Women, Nutr. 16 (2024) 1–11. 10.3390/nu16162767.

[20] X. Men, Z.L. Taylor, V.S. Marshe, D.M. Blumberger, J.F. Karp, J.L. Kennedy, E.J. Lenze, C.F. Reynolds, C. Stefan, B.H. Mulsant, L.B. Ramsey, D.J. Müller, CYP2D6 Phenotype Influences Pharmacokinetic Parameters of Venlafaxine: Results from a Population Pharmacokinetic Model in Older Adults with Depression, Clin. Pharmacol. Ther. 115 (2024) 1065–1074. 10.1002/cpt.3162.

[21] Y. Tsubono, I. Tsuji, F. Medicine, A. Alevizos, G. Larios, A. Mariolis, Psoriasis and Risk of Myocardial Infarction, 297 (2014) 361–363.

[22] P. Chiarella, R. Sisto, Variability of caffeine metabolism by CYP1A2 polymorphism in different populations, (2023) 1–13.

[23] B. L., D. M.-L., D. P., A.-S. A., Molecular genetics of CYP2D6: Clinical relevance with focus on psychotropic drugs, Br. J. Clin. Pharmacol. 53 (2002) 111–122. http://www.embase.com/search/results?subaction=viewrecord&from=export&id=L34264265%0A10.1046/j.0306-5251.2001.01548.x.

[24] H.Ç. Lenk, R.L. Smith, K.S. O’Connell, O.A. Andreassen, E. Molden, Rapid Metabolism Underlying Subtherapeutic Serum Levels of Atypical Antipsychotics Preceding Clozapine Treatment: A Retrospective Analysis of Real-World Data, CNS Drugs 38 (2024) 473–480. 10.1007/s40263-024-01079-y.

[25] U.M. Stamer, F. Stuber, Pharmacogenetics of anesthetic and analgesic agents: CYP2D6 genetic variations [1], Anesthesiology 103 (2005) 1099. 10.1097/00000542-200511000-00027.

[26] V.E. Gray, K.R. Kukurba, S. Kumar, Performance of computational tools in evaluating the functional impact of laboratory-induced amino acid mutations, Bioinformatics 28 (2012) 2093–2096. 10.1093/bioinformatics/bts336.

[27] M. Petrosino, L. Novak, A. Pasquo, R. Chiaraluce, P. Turina, E. Capriotti, V. Consalvi, Analysis and interpretation of the impact of missense variants in cancer, Int. J. Mol. Sci. 22 (2021). 10.3390/ijms22115416.

[28] S. Castellana, T. Mazza, Congruency in the prediction of pathogenic missense mutations: State-of-the-art web-based tools, Brief. Bioinform. 14 (2013) 448–459. 10.1093/bib/bbt013.

[29] C.L. Worth, R. Preissner, T.L. Blundell, SDM - A server for predicting effects of mutations on protein stability and malfunction, Nucleic Acids Res. 39 (2011) 215–222. 10.1093/nar/gkr363.

[30] F. Chiti, C.M. Dobson, Protein misfolding, amyloid formation, and human disease: A summary of progress over the last decade, Annu. Rev. Biochem. 86 (2017) 27–68. 10.1146/annurev-biochem-061516-045115.

[31] S. Teng, T. Madej, A. Panchenko, E. Alexov, Modeling effects of human single nucleotide polymorphisms on protein-protein interactions, Biophys. J. 96 (2009) 2178–2188. 10.1016/j.bpj.2008.12.3904.

[32] Z. Wang, J. Moult, SNPs, protein structure, and disease, Hum. Mutat. 17 (2001) 263–270. 10.1002/humu.22.

[33] C. Ferrer-Costa, M. Orozco, X. De La Cruz, Characterization of disease-associated single amino acid polymorphisms in terms of sequence and structure properties, J. Mol. Biol. 315 (2002) 771–786. 10.1006/jmbi.2001.5255.

[34] T.G. Kucukkal, M. Petukh, L. Li, E. Alexov, Structural and physico-chemical effects of disease and non-disease nsSNPs on proteins, Curr. Opin. Struct. Biol. 32 (2015) 18–24. 10.1016/j.sbi.2015.01.003.

[35] S. Sunyaev, V. Ramensky, P. Bork, Towards a structural basis of human non-synonymous single nucleotide polymorphisms, Trends Genet. 16 (2000) 198–200. 10.1016/S0168-9525(00)01988-0.

[36] M. Petukh, T.G. Kucukkal, E. Alexov, On human disease-causing amino acid variants: Statistical study of sequence and structural patterns, Hum. Mutat. 36 (2015) 524–534. 10.1002/humu.22770.

[37] S. Stefl, H. Nishi, M. Petukh, A.R. Panchenko, E. Alexov, Molecular mechanisms of disease-causing missense mutations, J. Mol. Biol. 425 (2013) 3919–3936. 10.1016/j.jmb.2013.07.014.

[38] H. Nishi, M. Tyagi, S. Teng, B.A. Shoemaker, K. Hashimoto, E. Alexov, S. Wuchty, A.R. Panchenko, Cancer Missense Mutations Alter Binding Properties of Proteins and Their Interaction Networks, PLoS One 8 (2013). 10.1371/journal.pone.0066273.

[39] C. Geourjon, C. Combet, C. Blanchet, G. Deléage, Identification of related proteins with weak sequence identity using secondary structure information, Protein Sci. 10 (2001) 788–797. 10.1110/ps.30001.

[40] A. Wallqvist, Y. Fukunishi, L.R. Murphy, A. Fadel, R.M. Levy, Iterative sequence/secondary structure search for protein homologs: Comparison with amino acid sequence alignments and application to fold recognition in genome databases, Bioinformatics 16 (2000) 988–1002. 10.1093/bioinformatics/16.11.988.

[41] R.A. Laskowski, N. Tyagi, D. Johnson, S. Joss, E. Kinning, C. McWilliam, M. Splitt, J.M. Thornton, H. V. Firth, C.F. Wright, DDD Study, Integrating population variation and protein structural analysis to improve clinical interpretation of missense variation: Application to the WD40 domain, Hum. Mol. Genet. 25 (2016) 927–935. 10.1093/hmg/ddv625.

[42] C. Zardecki, S. Dutta, D.S. Goodsell, M. Voigt, S.K. Burley, RCSB Protein Data Bank: A Resource for Chemical, Biochemical, and Structural Explorations of Large and Small Biomolecules, J. Chem. Educ. 93 (2016) 569–575. 10.1021/acs.jchemed.5b00404.

[43] T.A.M. Mulder, R.A.G. van Eerden, M. de With, L. Elens, D.A. Hesselink, M. Matic, S. Bins, R.H.J. Mathijssen, R.H.N. van Schaik, CYP3A4∗22 Genotyping in Clinical Practice: Ready for Implementation?, Front. Genet. 12 (2021). 10.3389/fgene.2021.711943.

[44] M. Zhao, J. Ma, M. Li, Y. Zhang, B. Jiang, X. Zhao, C. Huai, L. Shen, N. Zhang, L. He, S. Qin, Cytochrome p450 enzymes and drug metabolism in humans, Int. J. Mol. Sci. 22 (2021) 1–16. 10.3390/ijms222312808.

[45] S. V. Nielsen, A. Stein, A.B. Dinitzen, E. Papaleo, M.H. Tatham, E.G. Poulsen, M.M. Kassem, L.J. Rasmussen, K. Lindorff-Larsen, R. Hartmann-Petersen, Predicting the impact of Lynch syndrome-causing missense mutations from structural calculations, PLOS Genet. 13 (2017) e1006739. 10.1371/journal.pgen.1006739.

[46] Y. Peng, J. Norris, C. Schwartz, E. Alexov, Revealing the Effects of Missense Mutations Causing Snyder-Robinson Syndrome on the Stability and Dimerization of Spermine Synthase, Int. J. Mol. Sci. 17 (2016) 77. 10.3390/ijms17010077.

[47] A. Strokach, C. Corbi-Verge, P.M. Kim, Predicting changes in protein stability caused by mutation using sequence-and structure-based methods in a CAGI5 blind challenge, Hum. Mutat. 40 (2019) 1414–1423. 10.1002/humu.23852.

[48] Y. Chen, H. Lu, N. Zhang, Z. Zhu, S. Wang, M. Li, PremPS: Predicting the impact of missense mutations on protein stability, PLoS Comput. Biol. 16 (2020) 1–22. 10.1371/journal.pcbi.1008543.

[49] T. Yasmin, In silico comprehensive analysis of coding and non-coding SNPs in human mTOR protein, PLoS One 17 (2022) 1–23. 10.1371/journal.pone.0270919.

[50] Y. Bromberg, G. Yachdav, B. Rost, SNAP predicts effect of mutations on protein function, Bioinformatics 24 (2008) 2397–2398. 10.1093/bioinformatics/btn435.

[51] H.A. Shihab, J. Gough, M. Mort, D.N. Cooper, I.N.M. Day, T.R. Gaunt, Ranking non-synonymous single nucleotide polymorphisms based on disease concepts, Hum. Genomics 8 (2014) 4–9. 10.1186/1479-7364-8-11.

[52] V.E. Gray, R.J. Hause, J. Luebeck, J. Shendure, M. Douglas, Mutagenesis Data, 6 (2019) 116–124. 10.1016/j.cels.2017.11.003.Quantitative.

[53] P.C. Ng, S. Henikoff, SIFT: Predicting amino acid changes that affect protein function, Nucleic Acids Res. 31 (2003) 3812–3814. 10.1093/nar/gkg509.

[54] F. Ancien, F. Pucci, M. Godfroid, M. Rooman, Prediction and interpretation of deleterious coding variants in terms of protein structural stability, Sci. Rep. 8 (2018) 1–11. 10.1038/s41598-018-22531-2.

[55] D.E.V. Pires, D.B. Ascher, T.L. Blundell, MCSM: Predicting the effects of mutations in proteins using graph-based signatures, Bioinformatics 30 (2014) 335–342. 10.1093/bioinformatics/btt691.

[56] O. Caldararu, T.L. Blundell, K.P. Kepp, Three Simple Properties Explain Protein Stability Change upon Mutation, J. Chem. Inf. Model. 61 (2021) 1981–1988. 10.1021/acs.jcim.1c00201.

[57] D. Gilis, M. Rooman, PoPMuSiC, an algorithm for predicting protein mutant stability changes: application to prion proteins., Protein Eng. 13 (2000) 849–856.

[58] A. Bateman, M.J. Martin, C. O’Donovan, M. Magrane, E. Alpi, R. Antunes, B. Bely, M. Bingley, C. Bonilla, R. Britto, B. Bursteinas, H. Bye-AJee, A. Cowley, A. Da Silva, M. De Giorgi, T. Dogan, F. Fazzini, L.G. Castro, L. Figueira, P. Garmiri, G. Georghiou, D. Gonzalez, E. Hatton-Ellis, W. Li, W. Liu, R. Lopez, J. Luo, Y. Lussi, A. MacDougall, A. Nightingale, B. Palka, K. Pichler, D. Poggioli, S. Pundir, L. Pureza, G. Qi, S. Rosanoff, R. Saidi, T. Sawford, A. Shypitsyna, E. Speretta, E. Turner, N. Tyagi, V. Volynkin, T. Wardell, K. Warner, X. Watkins, R. Zaru, H. Zellner, I. Xenarios, L. Bougueleret, A. Bridge, S. Poux, N. Redaschi, L. Aimo, G. ArgoudPuy, A. Auchincloss, K. Axelsen, P. Bansal, D. Baratin, M.C. Blatter, B. Boeckmann, J. Bolleman, E. Boutet, L. Breuza, C. Casal-Casas, E. De Castro, E. Coudert, B. Cuche, M. Doche, D. Dornevil, S. Duvaud, A. Estreicher, L. Famiglietti, M. Feuermann, E. Gasteiger, S. Gehant, V. Gerritsen, A. Gos, N. Gruaz-Gumowski, U. Hinz, C. Hulo, F. Jungo, G. Keller, V. Lara, P. Lemercier, D. Lieberherr, T. Lombardot, X. Martin, P. Masson, A. Morgat, T. Neto, N. Nouspikel, S. Paesano, I. Pedruzzi, S. Pilbout, M. Pozzato, M. Pruess, C. Rivoire, B. Roechert, M. Schneider, C. Sigrist, K. Sonesson, S. Staehli, A. Stutz, S. Sundaram, M. Tognolli, L. Verbregue, A.L. Veuthey, C.H. Wu, C.N. Arighi, L. Arminski, C. Chen, Y. Chen, J.S. Garavelli, H. Huang, K. Laiho, P. McGarvey, D.A. Natale, K. Ross, C.R. Vinayaka, Q. Wang, Y. Wang, L.S. Yeh, J. Zhang, UniProt: The universal protein knowledgebase, Nucleic Acids Res. 45 (2017) D158–D169. 10.1093/nar/gkw1099.

[59] E. Capriotti, P. Fariselli, R. Casadio, I-Mutant2.0: Predicting stability changes upon mutation from the protein sequence or structure, Nucleic Acids Res. 33 (2005) 306–310. 10.1093/nar/gki375.

[60] N. Tang, B. Dehury, K.P. Kepp, Computing the Pathogenicity of Alzheimer’s Disease Presenilin 1 Mutations, J. Chem. Inf. Model. 59 (2019) 858–870. 10.1021/acs.jcim.8b00896.

[61] C. Savojardo, M. Manfredi, P.L. Martelli, R. Casadio, Solvent Accessibility of Residues Undergoing Pathogenic Variations in Humans: From Protein Structures to Protein Sequences, Front. Mol. Biosci. 7 (2021) 1–9. 10.3389/fmolb.2020.626363.

[62] S. Gudmundsson, M. Singer-Berk, N.A. Watts, W. Phu, J.K. Goodrich, M. Solomonson, H.L. Rehm, D.G. MacArthur, A. O’Donnell-Luria, Variant interpretation using population databases: Lessons from gnomAD, Hum. Mutat. 43 (2022) 1012–1030. 10.1002/humu.24309.

[63] M.M. Gromiha, M. Oobatake, H. Kono, H. Uedaira, A. Sarai, Relationship between amino acid properties and protein stability: Buried mutations, J. Protein Chem. 18 (1999) 565–578. 10.1023/A:1020603401001.

[64] F. Sievers, A. Wilm, D. Dineen, T.J. Gibson, K. Karplus, W. Li, R. Lopez, H. McWilliam, M. Remmert, J. Söding, Fast, scalable generation of high-quality protein multiple sequence alignments using Clustal Omega, Mol. Syst. Biol. 7 (2011) 539.

[65] D.K. Barupal, O. Fiehn, Generating the blood exposome database using a comprehensive text mining and database fusion approach, Environ. Health Perspect. 127 (2019) 2825–2830. 10.1289/EHP4713.

[66] W. Yeung, Z. Zhou, L. Mathew, N. Gravel, R. Taujale, B. O’Boyle, M. Salcedo, A. Venkat, W. Lanzilotta, S. Li, N. Kannan, Tree visualizations of protein sequence embedding space enable improved functional clustering of diverse protein superfamilies, Brief. Bioinform. 24 (2023) 1–10. 10.1093/bib/bbac619.

[67] S. Lu, J. Wang, F. Chitsaz, M.K. Derbyshire, R.C. Geer, N.R. Gonzales, M. Gwadz, D.I. Hurwitz, G.H. Marchler, J.S. Song, N. Thanki, R.A. Yamashita, M. Yang, D. Zhang, C. Zheng, C.J. Lanczycki, A. Marchler-Bauer, CDD/SPARCLE: The conserved domain database in 2020, Nucleic Acids Res. 48 (2020) D265–D268. 10.1093/nar/gkz991.

[68] D. Daudé, C.M. Topham, M. Remaud-Siméon, I. André, Probing impact of active site residue mutations on stability and activity of Neisseria polysaccharea amylosucrase, Protein Sci. 22 (2013) 1754–1765. 10.1002/pro.2375.

[69] M. Lorch, J.M. Mason, R.B. Sessions, A.R. Clarke, Effects of mutations on the thermodynamics of a protein folding reaction: Implications for the mechanism of formation of the intermediate and transition states, Biochemistry 39 (2000) 3480–3485. 10.1021/bi9923510.

[70] H.A. Shihab, J. Gough, D.N. Cooper, P.D. Stenson, G.L.A. Barker, K.J. Edwards, I.N.M. Day, T.R. Gaunt, Predicting the Functional, Molecular, and Phenotypic Consequences of Amino Acid Substitutions using Hidden Markov Models, Hum. Mutat. 34 (2013) 57–65. 10.1002/humu.22225.

[71] S.E. Flanagan, A.M. Patch, S. Ellard, Using SIFT and PolyPhen to predict loss-of-function and gain-of-function mutations, Genet. Test. Mol. Biomarkers 14 (2010) 533–537. 10.1089/gtmb.2010.0036.

[72] R. Mehra, K.P. Kepp, Structural heterogeneity and precision of implications drawn from cryo-electron microscopy structures: SARS-CoV-2 spike-protein mutations as a test case, Eur. Biophys. J. 51 (2022) 555–568. 10.1007/s00249-022-01619-8.

[73] S. Thakur, R.K. Verma, K.P. Kepp, R. Mehra, Modelling SARS-CoV-2 spike-protein mutation effects on ACE2 binding, J. Mol. Graph. Model. 119 (2023) 108379. 10.1016/j.jmgm.2022.108379.

[74] K.T. Bæk, R. Mehra, K.P. Kepp, Stability and expression of SARS-CoV-2 spike-protein mutations, Mol. Cell. Biochem. 478 (2023) 1269–1280. 10.1007/s11010-022-04588-w.

[75] R. Mehra, K.P. Kepp, Computational analysis of Alzheimer-causing mutations in amyloid precursor protein and presenilin 1, Arch. Biochem. Biophys. 678 (2019). 10.1016/j.abb.2019.108168.

[76] R. Mehra, K.P. Kepp, Structure and Mutations of SARS-CoV-2 Spike Protein: A Focused Overview, ACS Infect. Dis. 8 (2022) 29–58. 10.1021/acsinfecdis.1c00433.

[77] O. Caldararu, R. Mehra, T.L. Blundell, K.P. Kepp, Systematic Investigation of the Data Set Dependency of Protein Stability Predictors, J. Chem. Inf. Model. 60 (2020). 10.1021/acs.jcim.0c00591.

[78] S. Thakur, K. Planeta Kepp, R. Mehra, Predicting virus Fitness: Towards a structure-based computational model, J. Struct. Biol. 215 (2023). 10.1016/j.jsb.2023.108042.

[79] Q. Wuyun, Y. Chen, Y. Shen, Y. Cao, G. Hu, W. Cui, J. Gao, W. Zheng, Recent Progress of Protein Tertiary Structure Prediction, Molecules 29 (2024). 10.3390/molecules29040832.

[80] M. Šikić, S. Tomić, K. Vlahoviček, Prediction of protein-protein interaction sites in sequences and 3D structures by random forests, PLoS Comput. Biol. 5 (2009). 10.1371/journal.pcbi.1000278.

[81] S.M. Cascarina, E.D. Ross, Natural and pathogenic protein sequence variation affecting prion-like domains within and across human proteomes, BMC Genomics 21 (2020) 1–18. 10.1186/s12864-019-6425-3.

[82] T. Kajander, P.C. Kahn, S.H. Passila, D.C. Cohen, L. Lehtiö, W. Adolfsen, J. Warwicker, U. Schell, A. Goldman, Buried Charged Surface in Proteins, Structure 8 (2000) 1203–1214. 10.1016/S0969-2126(00)00520-7.

[83] K. Djinovic-Carugo, O. Carugo, Missing strings of residues in protein crystal structures, Intrinsically Disord. Proteins 3 (2015) e1095697. 10.1080/21690707.2015.1095697.

[84] K. Fechter, A. Porollo, MutaCYP: Classification of missense mutations in human cytochromes P450, BMC Med. Genomics 7 (2014) 1–9. 10.1186/1755-8794-7-47.

[85] N. Bouchoucha, D. Samara-Boustani, A. V. Pandey, H. Bony-Trifunovic, G. Hofer, Y. Aigrain, M. Polak, C.E. Flück, Characterization of a novel CYP19A1 (aromatase) R192H mutation causing virilization of a 46,XX newborn, undervirilization of the 46,XY brother, but no virilization of the mother during pregnancies, Mol. Cell. Endocrinol. 390 (2014) 8–17. 10.1016/j.mce.2014.03.008.

[86] S.W.A. Mula-Abed, F.B. Pambinezhuth, M.K. Al-Kindi, N.B. Al-Busaidi, H.N. Al-Muslahi, M.A. Al-Lamki, Congenital adrenal hyperplasia due to 17-alpha-hydoxylase/17,20-lyase deficiency presenting with hypertension and pseudohermaphroditism: First case report from Oman, Oman Med. J. 29 (2014) 55–59. 10.5001/omj.2014.12.

[87] Y. Mitsuuchi, T. Kawamoto, A. Rösler, Y. Naiki, K. Miyahara, K. Toda, I. Kuribayashi, T. Orii, K. Yasuda, K. Miura, K. Nakao, H. Imura, S. Ulick, Y. Shizuta, Congenitally defective aldosterone biosynthesis in humans: The involvement of point mutations of the P-450C18 gene (CYP11B2) in CMO II deficient patients, Biochem. Biophys. Res. Commun. 182 (1992) 974– 979. 10.1016/0006-291X(92)91827-D.

[88] K. Dumic, T. Yuen, Z. Grubic, V. Kusec, I. Barisic, M.I. New, Two Novel CYP11B1 Gene Mutations in Patients from Two Croatian Families with 11 β-Hydroxylase Deficiency, Int. J. Endocrinol. 2014 (2014) 1–6. 10.1155/2014/185974.

[89] N. Katsumata, M. Ohtake, T. Hojo, E. Ogawa, T. Hara, N. Sato, T. Tanaka, Compound heterozygous mutations in the cholesterol side-chain cleavage enzyme gene (CYP11A) cause congenital adrenal insufficiency in humans, J. Clin. Endocrinol. Metab. 87 (2002) 3808–3813. 10.1210/jcem.87.8.8763.

[90] A. Ambalavanan, S.L. Girard, K. Ahn, S. Zhou, A. Dionne-Laporte, D. Spiegelman, C. V. Bourassa, J. Gauthier, F.F. Hamdan, L. Xiong, P.A. Dion, R. Joober, J. Rapoport, G.A. Rouleau, De novo variants in sporadic cases of childhood onset schizophrenia, Eur. J. Hum. Genet. 24 (2016) 944–948. 10.1038/ejhg.2015.218.

[91] B.A. Bejjani, D.W. Stockton, R.A. Lewis, K.F. Tomey, D.K. Dueker, M. Jabak, W.F. Astle, J.R. Lupski, Multiple CYP1B1 mutations and incomplete penetrance in an inbred population segregating primary congenital glaucoma suggest frequent de novo events and a dominant modifier locus, Hum. Mol. Genet. 9 (2000) 367–374. 10.1093/hmg/9.3.367.

[92] A.W. Munro, K.J. McLean, J.L. Grant, T.M. Makris, Structure and function of the cytochrome P450 peroxygenase enzymes, Biochem. Soc. Trans. 46 (2018) 183–196. 10.1042/BST20170218.

[93] P. Lafite, F. André, J.P. Graves, D.C. Zeldin, P.M. Dansette, D. Mansuy, Role of arginine 117 in substrate recognition by human cytochrome P450 2J2, Int. J. Mol. Sci. 19 (2018). 10.3390/ijms19072066.

