## Supplementary file for "Computing pathogenicity of mutations in human cytochrome P450 superfamily"

### Supplementary section 1. Details of allele frequency used for our analysis

The allele frequencies (AF) for the 10 pathogenic mutations containing CYPs were collected from the gnomAD database on 29-09-2023. Basically, allele frequency refers to how often a particular variant (allele) occurs at a specific location in the genome within a given population. In the context of the gnomAD database, allele count (AC) and allele number (AN) are important metrics used to describe the distribution of genetic variants in a population. The allele count refers to the total number of times a specific allele (variant) is observed in a population. The allele number is the total number of alleles observed at a specific location in the genome within the studied population.

$$Allele\ frequency\ (AF) = \frac{Allele\ count\ (AC)}{Allele\ number\ (AN)}$$

### Supplementary section 2. Details of statistical analysis

The AUC-ROC were plotted between two parameters: the true positive rate (TPR) and the false positive rate (FPR).

$$TPR\ or\ Sensitivity = \frac{TP}{TP + FN}$$

$$FPR = \frac{FP}{TN + FP}$$

$$Specificity = 1 - FPR$$

Where TP represents the true positive, FP is the false positive, TN is the true negative, and FN is the false negative instance.

**Table S1. The datasets of CYP450 enzymes along with their UniProt ID, structures and the known non-pathogenic and pathogenic mutations studied.** The values for the mutations in brackets represent the mutation positions present in the PDB structure.

| CYP450 enzyme | UniProt ID | PDB code | Apo | Substrate bound | Inhibitor bound | Resolution (in Å) | Non-pathogenic mutations | Pathogenic mutations |
|---|---|---|---|---|---|---|---|---|
| CYP1A1 | P04798 | 4I8V | 0 | 3 | 3 | 2.6 | 501 (476) | - |
| CYP1A2 | P05177 | 2HI4 | 0 | 0 | 1 | 1.95 | 473 (434) | - |
| CYP1B1 | Q16678 | 6IQ5 | 0 | 0 | 2 | 3.7 | 613 (517) | 44 (39) |
| CYP2A6 | P11509 | 2FDV | 3 | 1 | 10 | 1.65 | 548 (518) | - |
| CYP2A13 | Q16696 | 4EJI | 0 | 0 | 5 | 2.1 | 534 (508) | - |
| CYP2B6 | P20813 | 3IBD | 0 | 0 | 13 | 2.00 | 485 (452) | - |
| CYP2C8 | P10632 | 2NNJ | 0 | 0 | 5 | 2.28 | 459 (436) | - |
| CYP2C9 | P11712 | 5A5I | 1 | 7 | 5 | 2.00 | 546 (511) | - |
| CYP2C19 | P33261 | 4GQS | 0 | 0 | 1 | 2.87 | 558 (517) | - |
| CYP2D6 | P10635 | 3TBG | 1 | 3 | 10 | 2.10 | 711 (677) | - |
| CYP2E1 | P05181 | 3E6I | 0 | 3 | 3 | 2.20 | 390 (366) | - |
| CYP2R1 | QVVX0 | 3CZH | 0 | 3 | 0 | 2.30 | 372 (326) | 6 |
| CYP3A4 | P08684 | 1W0G | 3 | 11 | 83 | 1.80 | 414 (378) | 1 |
| CYP3A5 | P20815 | 7SV2 | 1 | 0 | 3 | 2.46 | 352 (333) | - |
| CYP3A7 | P24462 | 7MK8 | 0 | 0 | 1 | 2.15 | 419 (378) | - |
| CYP7A1 | P22680 | 3V8D | 1 | 2 | 0 | 1.90 | 413 (396) | - |
| CYP8A1 | Q16647 | 3B6H | 1 | 0 | 1 | 1.62 | 465 (440) | 1 |
| CYP8B1 | Q9UNU6 | 7LYX | 0 | 0 | 1 | 2.60 | 48 | - |
| CYP11A1 | P05108 | 3N9Y | 0 | 4 | 0 | 2.10 | 397 (349) | 8 |
| CYP11B1 | P15538 | 7E7F | 0 | 0 | 2 | 1.40 | 517 (478) | 51 |
| CYP11B2 | P19099 | 4FDH | 0 | 0 | 7 | 2.71 | 487 (442) | 8 |
| CYP17A1 | P05093 | 6WR1 | 0 | 4 | 12 | 1.85 | 347 (314) | 26 |
| CYP19A1 | P11511 | 3S7S | 0 | 2 | 9 | 3.21 | 402 (356) | 6 |
| CYP21A2 | P08686 | 4Y8W | 0 | 1 | 1 | 2.64 | 457 (401) | 68 (66) |
| CYP46A1 | Q9Y6A2 | 7N3L | 1 | 1 | 13 | 1.63 | 264 (228) | - |
| CYP51A1 | Q16850 | 4UHI | 1 | 1 | 5 | 2.04 | 333 (282) | - |

**Table S2.** All possible mutations of twenty-six CYP450 enzymes.

| | |
|---|---|
| **CYP1A1** | |
| Non-pathogenic | L2R,I5N,I5V,M7T,M7V,S8L,A9V,T10M,T10R,L13I,L13P,L14P,F19L,C20F,C20R,V22L,W24C,W24R,V25A,R27K,S29P,P31R,P31S,V33I,K35E,G36D,L37P,K38N,P41L,G42R,P43T,W44L,W44R,G45A,G45D,G45R,W46C,P47H,P47S,I49N,I49T,G50E,G50R,M52L,T54A,L55P,K57N,K57R,P59L,P59Q,H60P,L61V,A62P,A62T,L63Q,S64P,R65K,M66I,M66V,S67N,S67R,Q69H,Y70C,G71R,D72E,D72N,V73E,V73M,Q75P,Q75R,R77L,R77Q,I78T,G79D,S80F,T81I,P82S,V83M,V83W,V84M,L86V,G88D,G88S,G88V,D90G,D90N,T91I,T91N,T91P,R93L,R93P,R93Q,R93W,A95V,R98Q,R98W,Q99H,G100D,G100V,D101E,D101N,F103V,G105D,R106L,R106Q,R106W,P107T,D108A,D108N,D108Y,L109R,Y110L,T111A,T113S,L114I,I115V,G118D,S124G,P125T,D126A,D126E,D126G,D126N,P129L,P129S,V130G,V130L,A132P,A133P,R134C,R134H,R134S,R135L,R135Q,R135W,R136C,R136H,A138T,N140S,S144R,F145L,I147T,I147V,A148V,S149F,S149P,S153F,T155A,C157Y,L159P,E160K,E161A,E161G,E161K,E161Q,H162R,S164N,K165N,A167G,E168G,E168K,V169A,I171K,T173M,T173R,L177P,A179T,G180R,P181S,N185H,N185K,P186T,Y189C,V190M,V191L,V192L,V194M,N196D,N196S,A200P,A200S,A200V,C202G,G204A,G204C,G204D,G204V,R205Q,R205W,R206C,R206H,R206S,Y207D,D208V,H209N,H209Q,H209R,H211R,H211Y,E213K,L214V,S216C,S216G,N219K,N222S,F224S,G225E,G225R,G225W,E226D,V227M,V228A,V228F,G229S,P233Q,P233R,D235A,D235H,I237N,P238S,I239V,L240I,L240P,R241C,R241H,R241L,L243P,L243Q,L243R,P244A,N245D,N245K,P246A,P246S,L248P,N249I,N249K,N249S,F251V,K252Q,N255K,E256K,Y259C,Y259H,F261I,F261L,M262I,M262L,M262V,K264N,M265R,M265V,E268K,H269N,F273L,K275E,R279G,R279Q,R279W,D280N,I281V,D283N,D283Y,S284G,S284N,L285P,I286T,K292T,Q293E,Q293H,E296D,E296K,N299S,V300I,S303L,D304R,E305G,E305K,K306N,I308M,I308T,I308V,V311I,D313G,L314F,L314P,F315C,A317V,G318E,G318R,D320G,T323K,T324A,T324S,A325S,I326V,S327F,S329N,S329R,M331I,M331K,V334A,V334E,V334M,N336S,P337S,Q340H,R341K,K342N,I343L,Q344H,E346G,E346Q,L347P,D348E,T349I,T349K,V350A,I351T,G352S,R353G,S354A,R355Q,R355W,R356Q,R356W,P357L,R358Q,R358W,L359F,D361E,D361H,R362I,R362S,S363F,S363Y,H364Q,Y367C,M368L,M368V,E369D,E369Q,A370V,F371Y,I372S,L373P,E374G,T375I,T375N,T375P,F376Y,R377Q,S380A,S380F,F381L,V382A,V382I,P383H,P383L,P383S,I386S,I386V,P387L,H388D,H388P,H388R,S389G,S389R,T391A,R392G,D393A,D393N,D393V,T394A,T394R,L396V,Y400H,I401L,I401N,G404E,G404R,R405C,R405H,R405S,C406Y,V407D,V407I,V409I,Q411H,Q411P,W412R,I414T,I414V,N415I,N415T,H416N,H416Q,D417N,K419E,W421C,V422A,V422I,P424L,P424T,E426G,R431Q,R431W,D436V,I439V,D440N,D440V,D440Y,K441R,V442G,E445D,E445V,V447A,V447L,V447M,I448N,I448T,I449F,I449N,F450L,F450S,G451V,M452I,M452T,G453D,G453R,G453V,K454N,R455Q,R455W,C457F,C457Y,G459S,T461I,T461N,I462F,I462L,I462N,I462T,I462V,A463G,A463S,A463T,A463V,R464C,R464H,R464S,E466K,V467G,F468S,F470V,F470Y,I473N,L474P,L475P,Q476E,R477G,R477Q,R477W,V478G,V478M,E479V,S481G,S481T,V482M,P483S,P483T,G485R,V486E,V486L,V486M,V488M,D489A,D489Y,T491A,T491N,P492R,Y494C,G495W,T497N,M498T,K499N,H500D,H500L,H500Y,A501V,C503F,C503R,C503S,F506C,F506L,Q507E,Q509H,Q509R,R511C,R511H,R511L. |
| | **CYP1A2** |
| Non-pathogenic | L3F,L3S,S6P,S6T,S6Y,V7A,P8H,P8S,S10L,A11S,E13G,L14R,L15F,S18C,S18F,S18Y,A19P,A19T,A19V,I20M,F21L,F21Y,C22Y,F25C,F25S,V27M,L28I,G30C,R32K,R32M,P33H,P33R,R34Q,R34W,V35G,P36L,P36S,P36T,K37I,K37R,K40R,P42R,P42S,E44K,G47D,G47S,W48G,P49T,L50F,L51F,G52R,H53L,H53N,T56A,K59R,P61L,P61S,A64T,L65P,R67G,S69G,Q70E,R71C,R71H,Y72H,G73E,G73R,G73W,D74E,D74N,V75F,V75I,Q77E,R79C,R79H,R79L,I80T,G81D,S82F,T83M,P84T,V85L,V85M,L86P,S89G,R90C,R90G,R90H,R90 |

| | |
|---|---|
| | L,R90S,D92G,T93I,I94S,R95L,R95Q,R95W,A97D,A97T,A97V,L98Q,V99M,R100Q,R100W,D103N,D104N,K106Q,G107D,R108Q,R108W,D110N,L111F,Y112C,S114F,S114P,L116F,L116V,I117N,G120V,T124I,F125I,F125L,D128N,S129C,P131L,P131T,R136C,R136H,R136P,R137Q,R137W,R138C,R138H,L139M,N142S,A143D,A143T,T146N,S148C,I149V,A150D,A150T,S151F,D152N,P153A,P153L,A154T,C159S,E162K,H164R,V165G,E168K,E168Q,K170E,A171D,A171V,L172M,R175S,L176M,Q177K,E178K,M180V,G182E,G182R,P183A,P183L,H185Q,F186L,D187E,D187N,P188S,Q191P,V193E,V194A,A197T,A197V,N198S,V199I,I200T,G201R,G201S,G201V,M203T,M203V,F205V,G206R,G206V,F209L,P210L,E211A,E211G,S212C,D214N,E215K,M216I,S218N,S218R,V220M,T223A,T223P,H224N,H224Y,E225D,V227A,V227M,T229A,T229N,A230P,S231F,S232Y,G233R,N234K,P235A,P235L,P235S,L236W,F238S,F239L,R243C,R243H,L245Q,P246S,N247K,P248L,P248T,A249V,L250R,R252K,K254R,N257S,N257Y,Q258K,Q258L,Q265K,K266Q,K266R,T267A,T267I,Q269R,H271R,Y272H,D276H,D276N,K277N,K277T,V280I,R281Q,R281W,D282N,T284M,G285V,A286G,A286S,A286T,L287P,K289N,K289Q,K289R,H290R,S291R,K292Q,G294A,P295S,A297D,S298R,G299S,G299V,N300S,L301F,L301V,I302N,I302T,P303S,K306R,I307L,I307T,N309K,N309Y,L310H,L310P,V311G,I314M,I314V,G316A,A317T,A317V,G318E,G318R,F319C,T321A,T324I,T324R,S327A,S329I,S329R,M331I,M331R,M331T,Y332N,L333P,V334M,T335A,P337L,P337S,E338Q,I339T,Q340H,R341M,K342N,K345R,E346K,L347V,D348N,T349I,V350G,G352C,G352S,E354K,R355Q,R355W,R356Q,R356W,P357A,R358Q,R358W,L359I,D361E,R362S,P363L,P363S,L365R,P366S,L368V,A370T,E374V,F376L,F376S,R377P,R377Q,H378N,S379Y,L382W,T385A,T385N,I386F,I386T,I386V,P387T,H388N,H388Y,S389G,T390A,R392G,R392M,R392S,T394I,T395A,T395M,L396P,N397S,G398S,P402S,C406Y,V407A,V407D,V407I,V409E,V409I,N410K,V414G,V414I,V414L,N415S,H416Y,P418A,P418L,E419G,S425T,E426D,R428L,R428Q,R428W,E430A,E430D,E430Q,R431L,R431P,R431Q,R431W,F432S,T434P,A435S,D436G,D436H,D436N,T438A,T438I,T438N,T438P,P443L,L444F,S445N,E446G,K447M,M448I,M448L,M448T,M449I,M449L,L450P,M453T,G454S,R456C,R456H,R457P,R457Q,R457W,C458R,I459V,G460R,E461K,V462L,A464D,K465T,W466R,E467D,E467G,I468L,I468V,L470V,A473D,I474N,Q478H,L479P,E480Q,S482G,S482R,V483M,P484L,P484S,P485L,P485Q,G486D,G486S,V487M,K488E,V489I,D490N,T492P,Y495C,G496R,L497P,T498N,M499T,M499V,A502T,R503C,R503H,E505K,H506L,V507L,A509E,A509P,A509V,R510L,R510Q,R510W,R512C,R512H,F513C,S514P,S514Y,I515V,N516S. |
| | **CYP1B1** |
| Non-pathogenic | G2A,G2D,G2R,T3A,T3N,T3P,S4T,L5H,L5V,S6G,P7L,P7S,D9A,P10H,P10L,W11R,P12L,P12Q,P15T,S17C,S17F,S17P,I18L,I18S,Q19H,Q19R,Q20R,T22R,L23F,L23P,S28W,V29L,T32A,T32P,H34R,H34Y,V35E,V35G,Q37P,Q37R,L39P,L40R,R41G,R41T,R43P,R43W,R45Q,R45W,L47I,L47P,R48G,R48P,A50V,P51S,P52L,G53A,P54L,F55C,F55I,F55S,F55V,A56V,W57R,P58Q,L59P,L59V,I60M,G61A,G61R,N62K,A63P,A64T,A64V,A65T,A65V,G67D,G67S,G67V,Q68R,A69V,A70T,A70V,H71Q,L72F,L72H,L72V,S73L,S73W,F74L,A75G,A75S,A75T,A75V,R76C,R76S,L77V,A78E,R79Q,R80G,R80H,R80L,R80P,Y81C,G82D,G82S,D83E,V84A,V84I,Q86R,I87N,R88G,R88L,R88S,L89P,L89Q,G90D,G90R,S91G,P93S,I94L,I94M,I94T,I94V,V95A,V95L,V95M,V96E,V96M,L97P,R101G,I103S,I103V,Q105R,A106D,A106G,L107V,V108E,Q109R,Q110H,Q110P,G111C,G111D,G111V,S112L,A113D,A113S,F114L,A115T,A115V,D116E,D116H,R117G,R117Q,R117W,P118L,P118S,A119S,A121T,S122F,F123L,R124H,R124L,V125G,S127A,G128D,G128R,G128S,G129S,R130C,R130L,A133P,A133S,A133T,A133V,F134L,H136N,Y137C,Y137N,S138L,S138W,E139A,E139K,W141G,K142N,V143L,V143M,Q144H,R146C,R146G,A147G,A147P,A148T,A148V,H149Y,S150G,S150N,M151I,M151L,M151R,M151T,M152I,R153C,R153H,R153S,N154K,N154S,F156V,T157M,Q159H,P160L,R161L,S162N,R163C,Q164H,L166P,E167G,G168D,G168S,V170M,L171M,L171P,S172G,E173K,R175G,E176K,L177P,V178L,V178M,A179V,L181P,V182L,R183C,R183S,G184C,G184R,G184S,S185R,A186G,A186T,D187Y,G188C,G188R,G188S,A189D,A189P,F190C,F190L,L191R,D192A,P193R,P193S,R194K,R194M,P195L,P195S,T197N,V199L,V199M,A202T,N203D,V204A,M205V,S206G |

| | |
|---|---|
| | ,S206N,A207D,V208L,V208M,G211D,R213C,R213H,R213S,S215T,H216N,H216Q,D217E,D217H,D218A,D218V,D218Y,P219S,R222G,R222P,H227Y,F231Y,R233L,T234M,V235A,V235M,G236R,G238C,S239G,L240P,V241L,D242E,D242H,V243L,M244I,P245L,P245S,Q248R,Y249C,Y249D,N252I,N252K,N252S,P253A,P253L,P253R,V254M,R255C,R255H,R255L,V257I,V257L,R259H,R259L,E260K,E260Q,F261L,E262K,N265D,N265I,N265S,R266C,R266H,R266L,N267S,F268L,S269G,S269R,N270S,L273R,K275T,L277W,C280W,C280Y,S282R,L283F,R284W,P285S,G286R,G286W,A287T,A288P,A288T,A288V,P289L,P289S,R290C,R290P,D291G,D291N,D291Y,M292I,M292L,M292V,M293I,M293V,D294Y,A295T,F296C,F296L,I297T,L298H,S299C,S299P,A300E,A300T,E301D,K302T,K303T,A304E,G306E,G306R,G306V,D307A,S308P,H309R,H309Y,G310D,G311A,G311S,G312D,A313G,A313V,R314Q,L315P,D316E,D316N,D316V,L317F,E318D,N319S,V320A,V320I,P321Q,P321R,T323S,T325I,G329S,A330P,A330S,A330V,D333E,L335Q,S336Y,T337I,T337N,A338E,A338T,A338V,L339M,Q340R,W341R,L343P,L345V,F346L,R348T,Y349D,Y349H,D351E,D351V,V352A,Q353P,T354S,R355P,A358T,A358V,E359A,L360S,D361H,D361N,V363A,V363I,V364A,V364G,G365R,R366G,R366K,D367Y,R368C,R368G,L369Q,C371S,M372I,M372L,M372V,G373D,G373V,Q375H,N377S,L378P,P379L,L382M,A383D,F384V,L385F,E387Q,M389I,S393I,V395A,V395G,V395L,V397D,V397L,I399V,P400S,H401R,A402P,A402V,T404S,A405T,A405V,N406T,T407A,T407I,Y412C,Y412N,I414F,I414V,P415L,K416E,D417G,T418I,V419G,V419M,F421L,V422I,V422G,N423I,Q424R,W425C,W425R,N428S,H429D,H429Y,D430E,P431L,P431S,L432P,L432Q,L432V,K433M,W434C,W434R,P435L,P435R,N436H,N436K,P437A,P437Q,P437S,P437T,E438D,N439D,F440C,F440L,F440S,D441E,D441H,P442A,A443P,A443V,R444P,F445L,L446F,K448N,D449E,D449H,D449N,D449Y,G450S,N453S,N453T,K454Q,D455E,L456P,L456V,S458I,M461I,M461T,M461V,I462T,S464F,V465M,K467R,R469Q,C470R,C470S,C470Y,I471L,I471N,I471T,I471V,G472R,E473K,E474V,S476P,K477E,K477R,K477T,M478I,F483L,I484M,I484T,S485C,I486V,L487P,H489R,H489Y,Q490L,C491R,C491S,D492N,A495T,N496D,P497S,N498I,E499K,P500H,A501E,A501T,A501V,S506N,L509I,T510P,I511S,P513T,F516L,V518L,V518A,N519S,N519T,T521I,T521P,L522P,E524D,M526T,E527Q,L528I,L528P,L529P,D530E,D530H,D530N,S531R,S531T,Q534K,Q537K,K539E,K539R,K539T,T541S. |
| Pathogenic | M1T,S28W,W57C,G61E,L77P,Y81N,A115P,M132R,Q144P,Q144R,R145W,D192V,P193L,V198I,N203S,S215I,E229K,G232R,S239R,V320L,A330F,L345F,V364M,G365W,R368H,D374N,E387K,A388T,R390C,R390H,R390S,I399S,V409F,N423Y,P437L,A443G,R44Q,F445C,G466D,R469W,E499G,S515L,R523T,D530G |
| **CYP2A6** | |
| Non-pathogenic | L2Q,A3S,A3V,G5R,M6I,M6L,M6T,L7F,L8V,V9M,L11W,L12M,V13A,V13F,C14Y,T16S,V17A,V17G,M18T,V19I,L20F,M21I,S22T,Q25K,Q25R,R27G,R27K,K28E,S29I,S29N,K30N,K30R,G31E,K32R,P34A,P35L,P35R,G36A,G36V,P37A,P37L,P37R,P37T,T38N,P39L,L40W,P41L,P41S,I43L,I43T,G44A,G44E,L47P,Q48P,N50I,N50S,E52Q,M54K,N56K,S57F,L58P,I61F,I61L,I61N,E63G,R64C,R64H,R64S,P67S,V68A,V68M,P75L,R76P,R76Q,R76W,R77L,R77Q,R77W,V78I,V79M,V80M,G83A,G83R,H84P,H84Q,H84R,H84Y,D85G,A86V,V87I,R88K,R88S,R88W,E89Q,A90V,L91V,V92A,V92M,D93N,Q94E,Q94K,Q94R,A95V,E96K,E97K,S99G,G100E,G100R,R101G,R101L,R101Q,E103K,A105V,F107L,D108N,W109C,V110F,V110I,V110L,F111L,G113S,Y114N,G115D,V116L,V116M,V117A,V117L,F118L,F118Y,S119R,S119T,N120H,N120K,N120S,G121E,G121R,G121W,E122K,A124V,K125E,K125Q,Q126H,Q126L,L127F,R128L,R128P,R128Q,R128W,R129C,R129H,R129P,S131A,I132N,I132T,T134P,R136W,F138I,F138L,G139A,G139E,G139R,V140A,V140M,G141R,K142Q,R143G,R143P,R143Q,G144S,I145S,E146D,E146G,E146K,E146Q,E147D,E147K,R148C,I149N,E151K,E152G,A153P,A153S,G154D,I157F,I157M,D158E,D158N,D158Y,L160H,L160I,R161P,R161W,G162C,G162D,G162S,G162V,G164S,G164V,G165C,G165R,A166S,A166T,N167H,N167K,N167S,I168N,D169E,D169H,P170S,F173L,N180 |

| | |
|---|---|
| | S,V181G,V181I,V181L,I182L,I185T,V186L,G188E,G188V,D189E,R190C,R190H,R190L,R190S,F191S,F191V,K194E,K196N,E197D,E197G,F198C,L199P,L202F,R203C,R203G,R203H,R203P,R203S,M204V,M205I,M205T,F209C,F211V,T212M,T212R,T214A,T214P,S215F,T216R,G217A,Q218E,Y220H,E221K,M222I,M222T,S224P,S225L,V226E,H229P,P231S,G232A,G232E,P233T,Q234E,Q234P,Q235K,A237D,A237G,A237T,A237V,Q239K,L240V,G243E,G243R,L244P,E245K,D246A,D246E,D246N,D246Y,F247L,I248T,K251N,E253A,E253Q,H254N,H254Q,N255D,Q256K,R257C,R257G,R257H,R257L,R257P,R257S,T258K,T258M,T258R,T258S,L259M,N262S,S263A,S263T,P264T,R265G,R265P,R265Q,R265W,I268L,I268M,I268T,S270C,S270F,I273M,I273T,R274C,R274H,R274P,R274S,M275I,M275L,Q276K,Q276P,E278G,E279A,E279K,E279Q,K280E,K280N,N281K,T284K,T284M,F286I,F286L,Y287H,Y287S,L288F,K289R,V292G,V292M,M293I,M293L,M293T,T294A,T294I,T294S,T295M,L296M,N297K,N297S,N297T,I300N,I300T,G301A,G301R,G301V,G302A,G302D,E304K,T305N,V306I,T308I,T308N,T309I,R311C,R311G,R311H,R311P,R311S,Y312C,F314L,L317F,K319R,H320N,V323G,V323M,E324K,A325P,K326N,K326R,H328R,E329K,E330D,I331T,R333S,V334G,V334M,I335M,G336R,G336S,R339P,R339Q,R339W,P341L,R346Q,R346W,A347T,M349I,M349K,Y351H,M352I,M352T,M352V,E353K,A354S,V355M,H357D,H357Q,H357Y,E358D,E358K,R361I,F362C,F362S,D364H,D364N,V365M,I366F,I366L,I366N,P367A,M368I,M368R,M368T,M368V,L370F,L370S,L370V,A371V,R372C,R372H,R372P,K375N,K376R,K379T,R381Q,R381W,D382Y,F384S,L385F,P386S,K387N,T389I,E390D,E390K,E390Q,V391L,V391M,Y392C,Y392F,Y392H,Y392S,M394T,M394V,L395P,G396A,V398M,L399R,R400K,P402S,S403I,S403R,S406C,N407K,P408S,Q409R,D410Y,F411V,F411Y,N412S,P413L,P413S,P413T,Q414R,L417V,N418D,E419D,E419K,K420E,K420M,Q422K,K425N,K425T,S426G,S426M,S426N,D427G,A428G,A428V,I434M,I434V,G435E,G435R,N438Y,C439S,F440L,G441R,G443A,G443D,L444P,L444V,A445V,R446G,M447I,E448D,L451I,F452Y,F453L,V456F,V456I,M457I,M457T,R461G,R461H,R461L,R461P,L462F,S465C,S465P,S465T,Q466L,Q466P,P468S,D470Y,I471T,D472V,P475H,P475S,K476R,G479V,F480S,A481V,T482K,R485L,R485P,N486I,Y487S,M489I,M489K,M489R,M489T,M489V,S490G,S490N,F491C,P493L,R494G,R494S. |
| **CYP2A13** | |
| Non-pathogenic | A3T,A3V,S4A,S4T,G5R,L7F,T10A,A13S,A13V,C14G,T16S,M18R,M18I,M18T,M18V,L20F,S22P,W24R,R25Q,R25L,R25W,K28N,S29R,S29N,S29I,R30K,K32R,P35R,P35L,P35S,G36V,P37R,T38N,T38P,P41L,Q48H,Q48K,L49P,T51I,E52Q,E52K,Q53K,M54K,N56S,S57F,L58F,M59I,K60E,I61L,I61F,S62I,R64C,R64H,R64S,Y65N,Y65C,Y65H,G66S,P67R,T70A,T70I,L73F,G74E,P75T,R76Q,R76G,R76W,R77Q,R77P,R77W,V79A,V79L,V79M,V80A,V80M,L81M,L81V,C82Y,G83R,G83E,H84R,H84L,H84Y,D85V,A86P,A86T,A86V,V87I,K88R,K88N,E89Q,A90S,A90V,V92M,D93H,D93V,G100R,R101Q,R101G,G102D,G102S,E103K,E103V,F107L,D108N,W109R,W109G,L110P,L110V,F111L,F111V,Y114C,G115R,G115S,G115V,V116E,V116L,V116M,A117S,A117V,S119T,G121R,G121E,E122K,R123H,R123S,A124S,Q126H,R129H,R129P,R129S,F130I,I132T,I132V,A133V,T134N,T134S,L135R,L135Q,R136K,R136S,G137D,G137C,V140G,V140L,G141D,G141S,R143G,R143H,R143P,R143S,G144R,G144C,G144S,G144V,I145V,E146Q,E146K,E147K,R148H,R148S,I149F,Q150R,Q150E,Q150H,E151Q,E151V,A153E,A153V,G154S,G154V,F155L,D158N,D158E,D158H,A159T,A159V,L160I,R161Q,R161W,T163M,H164R,H164D,G165R,G165D,G165S,N167H,N167S,I168M,I168V,D169E,D169H,D169V,P170S,T171I,T171S,F172V,F173I,F173L,L174R,S175N,R176C,R176H,T177R,V178A,V178G,S179F,N180S,V181I,I182F,S184P,V186F,F187L,D189N,R190C,R190G,E194K,E194V,D195G,K196E,E197G,L199Q,S200L,L202F,R203C,R203H,R203L,R203S,M204V,M205T,S208G,S208I,F209Y,Q210R,T212M,A213S,A213V,T214P,T216A,T216M,G217R,G217V,L219H,Y220H,M222I,M222V,F223I,S224F,S225L,M227I,M227T,M227V,H229R,H229Q,H229P,H229Y,G232R,G232E,Q234E,Q236R,A237V,K239Q,E240A,E240V,E245A,E245K,D246N,I248T,I248V,A249T,K250R,K250N,N255S,Q256E,R257C,R257H,R257S,T258A,T258R,T258K,T258M,P261A,N262K,S263A,P264T,R265Q,R265W,D266E,I268L,D269N,D269G,S270Y,R27 |

| | |
|---|---|
| | 4C,R274H,R274L,R274S,M275V,Q276K,E279D,E279Q,E279G,E279K,K280R,T284A,T284R,E285Q,F286S,K289E,N290K,L291M,V292A,V292M,M293T,T294N,T294I,L296P,N297K,L298H,L298V,F300S,A301E,A301V,G302D,G302V,T303A,T303P,E304D,T305A,T305I,V306M,S307R,S307C,T308P,L310P,R311C,R311H,R311S,G313A,G313S,L315R,L316R,L316M,L316P,M318V,K319R,H320N,P321L,V323L,V323M,E324D,E324G,A325P,A325T,K326E,V327A,V327I,V327F,E329A,I331T,V334L,I335M,I335F,I335T,G336S,K337R,N338I,R339Q,R339W,R346Q,R346W,A347T,M349T,P350S,P350T,Y351H,T352A,T352I,A354G,V355A,I356N,E358K,I359L,I359V,Q360K,R361K,F362S,M365I,M365T,M365V,L366I,M368K,L370M,L370F,L370W,L370V,H372R,H372P,V374I,V374F,N375D,N375K,D377N,K379Q,F380I,R381Q,R381W,F384L,P386R,P386T,G388D,T389A,T389I,V391E,V391M,F392Y,P393L,M394I,M394L,L395V,G396D,S397F,V398M,D401E,D401G,D401H,P402R,P402H,P402L,P402S,R403G,R403M,R403S,F404S,N407S,P408L,P408S,R409Q,R409G,R409W,D410A,D410E,D410Y,F411L,N412H,P413L,D418N,K419N,K419E,K419T,K420N,G421R,F423L,K424N,K425M,S426R,S426C,S426G,D427G,F429S,V430A,P431L,P431S,F432L,I434T,G435R,R437Q,R437P,R437W,Y438N,Y438C,C439R,C439Y,G441A,G443V,L444P,A445T,M447T,M447V,E448K,L449F,L449V,F450L,F450S,L451F,F452V,F453Y,I456V,Q458H,F460C,F460L,R461C,R461H,R461P,F462I,F462L,S467L,S467P,S467T,P468R,D470V,I471S,I471T,I471V,D472N,D472E,V473A,V473E,V473L,P475A,P475L,H477Q,V478A,V478L,V478M,G479A,G479D,G479V,F480S,A481G,A481V,T482M,I483S,P484R,P484L,P484S,R485Q,N486K,Y487C,Y487S,T488A,S490R,L492V,R494C,R494L |
| **CYP2B6** | |
| Non-pathogenic | E2D,E2K,L3I,V5D,V5I,V5L,L6H,L9I,L11F,L12I,L12R,G14E,L15I,L18V,L19M,L19R,L19V,Q21L,R22C,R22H,R22L,R22P,P24A,T26S,H27L,D28G,R29C,R29H,R29P,R29S,R29T,L30F,P31S,G33R,P34A,R35C,R35H,R35S,L37M,P38S,L39F,L40S,L40W,G41E,N42I,N42K,L44Q,L44V,M46L,M46T,M46V,D47N,R48K,L51P,L52P,K53Q,F55L,R57W,F58L,R59Q,K61I,K61T,Y62C,D64N,V65D,V65I,T67M,V68I,V68L,L70P,G71A,G71R,P72L,P72R,P74A,V75L,V75M,V76F,V76I,C79R,G80V,V81A,E82D,A83V,R85P,R85Q,R85W,E86D,E86G,A87V,L88F,L88P,K91N,E93D,A94D,A94S,A94T,S96C,S96L,R98G,R98Q,R98W,G99E,I101S,A102S,A102T,M103K,M103T,D105N,P106Q,F108L,R109Q,R109W,G110A,G110R,G110V,Y111C,I114F,I114T,N117D,N117S,G118R,R120C,R120H,V123M,L124P,L124V,R125Q,R125W,R126Q,F127I,S128F,T131A,M132V,R133K,D134Y,G136R,M137I,M137V,G138E,K139E,K139M,K139N,R140P,R140Q,R140W,S141N,S141R,R145Q,R145W,I146S,I146T,E148D,E149K,A150D,A150P,Q151E,C152S,I154T,E155Q,E156K,L157I,R158Q,R158W,K161E,A163G,A163T,L164F,L164P,M165I,M165L,M165V,D166A,P167A,P167S,T168I,T168P,L170F,F171S,Q172H,S173C,I174F,T175A,T175P,A176T,N177Y,I179L,I182M,V183G,V183I,F184L,G185E,R187Q,H189Y,Q191E,Q191K,Q191R,D192E,D192G,Q193E,Q193K,F195I,F195L,L196P,L196Q,M198I,M198T,L201C,F202L,Y203H,Y203S,F206S,S207L,S207P,I209F,I209V,S210G,F213C,F213L,F213S,G214V,L216P,L224S,K225N,Y226H,P228S,A230P,A230S,R232K,Q239R,E240D,E240K,E240V,N242H,N242S,A243P,I245F,I245L,I245T,I245V,S248R,V249M,E250D,E250G,E250K,H252N,R253C,R253G,R253H,T255S,D257N,S259N,S259R,A260S,A260T,A260V,K262R,D263G,L264I,I265M,D266N,D266V,T267N,Y268H,L269R,L269V,L270I,H271L,M272L,E273A,E273K,E275D,S277F,N278D,N278H,N278S,A279P,A279S,A279T,H280Q,H280R,S281G,E282K,F283L,H285R,Q286R,N289H,N289I,N289K,L290I,T292M,L293F,S294L,A298V,G299V,T302N,T305N,T306I,T306S,L307F,R308C,R308H,R308S,Y309C,Y309D,Y309H,G310S,L313I,L313V,M314R,Y317S,P318L,E322D,E322K,R323G,R323S,R323T,V324I,Y325F,E327D,E327G,I328T,E329A,E329K,E329V,V331E,V331L,I332F,I332T,I332V,G333S,P334T,H335R,R336C,R336H,R336L,R336P,P337L,P338T,E339A,E339D,H341D,H341Q,H341R,D342E,R343P,R343Q,P347S,T349R,E350D,E350G,A351T,I353M,Y354C,Y354H,E355K,F359L,S360A,S360C,S360Y,D361N,D361V,D361Y,L362H,P364T,M365R,M365V,G366D,V367L,P368L,H369Y,V371F,T372A,H374N,T375N,S376G,S376N,S376R,R378Q,G379A,G379 |

| | |
|---|---|
| | W,Y380H,I381L,I381T,I381V,I382F,I382N,K384R,D385G,D385Y,E387G,V388I,L390I,I391N,A395T,A395V,L396V,H397R,H397Y,P399T,H400Y,P405Q,D406E,A407T,N409D,N409H,N409K,D411Y,H412Q,H412R,F413L,D415E,A416V,N417S,G418E,G418W,A419S,T423N,E424K,A425D,I427M,I427T,P428T,F429L,L431S,G432R,R434Q,R434W,I435V,G438D,G438V,G440A,G440V,I441T,A442P,A442T,A442V,R443C,R443H,R443S,A444E,A444V,E445Q,F447L,F447Y,L448I,F449V,F450C,F450L,T451I,T452I,M459T,M459V,A460T,S461R,V463A,V463M,A464D,A464T,D467Y,I468M,D469E,D469N,P472L,P472S,Q473P,C475G,C475S,G476A,G476D,G476V,P481L,P481T,T483I,Y484S,Q485H,R487C,R487H,R487L,R487S,P490H,P490L,R491C,R491H,R491L |
| **CYP2C8** | |
| Non-pathogenic | P3R,P3T,F4S,F4V,V5E,V8L,V8M,C10R,L11V,S12Y,M14I,M14K,M14L,M14V,L15F,L16F,L16P,L16V,L19R,R21G,R21S,S23I,R26K,L29H,P30T,P31A,P31S,P33L,P33S,T34I,T34S,P35L,I38V,I39T,I39V,G40E,G40V,M42I,Q44H,Q44K,Q44P,I45K,I45V,D46N,V47I,D49N,I50T,C51G,K52Q,K52R,S53Y,T55A,N56S,F57L,K59E,K59N,V60A,Y61C,Y61D,G62R,P63A,V64G,V64L,V64M,F65Y,V67M,Y68C,M71I,M71T,N72K,P73A,P73L,I74M,V76G,V76L,V76M,F77V,H78R,H78Y,G79R,E81Q,A82S,A82T,E85G,A86T,A86V,L87Q,I88S,I88T,D89N,N90D,E93Q,G98V,N99I,N99S,S100P,P101L,I102V,S103T,R105I,T107N,K108E,K108N,L110F,S114C,S114F,S115I,S115R,N116I,G117A,K118T,K121E,E122K,I123N,R124Q,R124W,R125C,R125G,R125H,L128H,T129K,T130I,T130N,R132Q,R132W,M136I,M136L,M136V,R139K,R139M,I141T,I141V,E142K,D143N,R144C,R144G,R144H,V145L,Q146R,E147D,H150R,E154A,E154K,E154D,E155K,L156F,K158E,K158I,K158N,K158R,K160N,A161T,S162A,P163L,C164R,C164Y,P166L,P166S,T167I,T167S,I169T,I169V,L170M,L170P,G171S,A173S,A173T,P174S,C175R,N176D,N176T,V177A,V177G,I178F,C179S,V181A,V181D,V181I,V182I,K185T,R186G,R186L,R186Q,D188G,K190R,D191H,D191V,D191Y,N193H,N193K,F194Y,T196I,T196N,M198L,M198V,R200K,F201L,F201Y,N202S,E203D,I207T,S210T,W212C,V215I,C216R,N217K,N218K,N218T,P220T,I223M,C225F,C225S,P227L,T229S,H230P,H230Q,N231K,K232R,V233L,K235E,K235I,N236M,V237F,V237I,A238G,A238P,A238V,L239F,L239P,T240I,R241G,R241Q,S242N,Y243C,Y243D,I244F,I244T,I244V,E246Q,K247R,K249R,E250D,E250K,H251L,H251P,H251Q,A253P,A253T,A253V,D256G,D256V,D256Y,N259K,N259S,P260S,R261L,R261Q,R261W,D262E,D262G,I264M,I264V,D265N,C266F,I269F,K270R,E272D,Q273H,Q273R,E274G,E274Q,E274V,K275Q,N277K,F282L,N283S,N283T,L287V,V288F,V288I,T290A,V291A,V296A,G298V,T299K,T301I,T301K,S303G,T304A,T304S,T305A,L306M,L306R,R307G,Y308H,L311V,L312I,H316R,P317S,V319I,V319L,T320I,A321P,A321S,V323F,Q324H,E326K,I327T,D328V,H329L,H329Y,V330A,I331M,I331S,I331T,G332C,H334Q,R335G,S336C,S336G,P337T,R342K,S343I,S343N,H344R,M345V,P346A,P346S,P346T,T348I,T348S,A350V,V351A,E354K,I355F,I355N,Q356H,Q356K,S359R,L361F,L361I,P363H,P363S,P363T,T364A,T364H,G365C,G365D,G365S,V366L,V366M,A369S,T371I,T371S,T372A,Y379C,P382S,K383R,K383N,G384S,T385I,I387T,I387V,M388V,A389S,A389T,A389V,L390S,L391M,L391P,L391V,S393C,S393F,V394C,V394M,H396P,D397N,D398G,K399R,P402T,N403S,P404A,N405T,I406M,F407S,D408E,P409S,H411Y,H411L,D414G,N416H,G417A,G417D,G417V,N418Y,S422N,D423G,Y424D,F425L,F425V,M426I,M426T,M426V,P427H,P427S,P427T,G431A,R433Q,A436G,A436S,A436T,A436V,E438G,E438Q,R442C,R442H,R442L,M443I,M443L,L445V,T450N,T451A,I452V,Q454H,Q454K,N455K,K459T,V461F,D462A,N466Y,N468S,T469N,T470A,T470I,A471S,V472I,T473A,T473N,K474R,G475E,G475R,I476T,V477A,V477I,S478P,L479M,P480T,P481H,P481S,S482L,Y483D,Q484H,Q484K,I485M,C486F,F487L,I488N,P489L,P489R. |
| **CYP2C9** | |
| Non-pathogenic | D2N,D2V,S3C,S3P,S3Y,L4F,V5A,V5L,V6F,L7F,L7H,L9F,C13Y,L17F,S18L,L19I,L19R,W20C,S23G,S23N,S24P,R26K,L29R,P30L,P31T,G32D,G32R,P33L,T34A,T34S,P35L,L36R,P37A,P37L,P37S,V38E,I39F,G40R,G40V,N41I,I42N,I42V,L43P,Q44H,I45M,G46D,G46 |

| | |
|---|---|
| | V,I47T,I47V,D49G,D49V,I50V,S51N,S51R,S53F,T55I,N56S,L57P,S58L,V60A,V60I,G62D,G62S,V64M,F65L,L67P,Y68C,G70D,G70R,G70S,L71R,L71V,K72E,P73L,P73T,I74T,I74V,V76L,H78P,H78Y,Y80F,A82V,V83E,V83M,K84E,E85A,A86D,A86P,A86T,A86V,I88T,I88V,D89N,D89Y,L90P,G91R,E92D,S95Y,G96V,R97G,R97K,R97T,G98C,G98D,G98R,G98V,I99L,I99N,I99T,P101L,A103T,E104D,A106D,A106T,R108G,G109E,G109R,G111E,V113A,S115R,K118N,K119N,K119R,K119T,W120L,W120R,R124L,R124Q,R124W,R125C,R125H,R125L,R125P,R125S,F126L,S127F,S127P,L128P,L128R,M129I,M129K,M129T,T130K,T130M,R132Q,R132W,F134L,G135R,G135V,M136I,M136L,M136V,K138N,S140N,I141N,I141T,I141V,D143H,R144C,R144H,V145F,V145I,A149P,A149T,A149V,R150C,R150H,R150L,R150P,C151Y,L152P,V153A,V153G,E154D,E154G,E154K,E155D,L156V,T159N,A161P,S162L,P163L,D165G,P166A,P166L,P166S,L170M,G171C,G171R,C172S,A173D,A173T,A173V,P174A,P174L,C175F,I178S,I178V,S180F,S180P,I181V,F183S,H184P,H184Q,K185T,R186C,R186H,R186L,R186S,F187L,D188Y,K190E,K190N,D191E,Q193R,N196I,N196K,L197V,M198I,M198T,E199K,E199V,K200N,L201V,N202D,N204S,N204H,I205F,I205M,K206N,I207F,I207T,L208F,L208V,S210N,S210R,P211L,P211S,P211T,W212C,I213F,I213L,I213T,Q214K,C216F,C216Y,N218I,F219C,P221S,I223T,I223V,Y225H,P227L,P227Q,G228E,G228R,T229A,T229P,H230D,H230Q,K235R,N236K,N236T,V237A,V237F,V237I,A238S,A238V,M240T,M240V,K241E,S242R,I244N,I244T,I244V,E246K,H251R,H251Y,M255I,M255R,M255T,D256N,M257I,N258S,N259I,N259K,N259S,P260H,P260L,P260T,Q261P,F263L,I264S,I264T,L268P,M269I,M269R,M269T,M271I,M271T,E272G,E272K,K273E,K273M,K273T,E274K,E274Q,K275E,H276L,H276R,N277S,Q278H,Q278K,P279T,S280P,S286N,L287M,E288G,N289K,N289S,T290A,V292A,D293H,L294F,F295L,A297V,G298R,T299A,T299I,T299K,T299R,T301A,T301M,T302S,S303G,S303I,S303N,T304A,T304P,T305A,T305S,L306M,L306P,R307K,A309V,L311F,L314R,L314V,P317S,T320I,A321P,V323A,Q324H,Q324R,E325K,E326K,I327T,E328G,R329C,R329G,R329H,R329S,V330A,I331T,R333G,R333K,N334H,N334K,N334T,R335Q,R335W,S336R,P337L,P337R,P337T,C338G,C338R,C338Y,M339L,Q340R,D341G,R342G,R342K,S343R,H344L,M345I,M345T,P346L,P346S,Y347F,Y347H,Y347N,T348A,D349E,D349N,A350V,V352M,E354K,E354Q,V355I,Q356R,R357G,R357S,Y358C,Y358S,I359L,I359N,I359T,I359V,D360A,D360E,D360N,L361F,L361I,L362V,P363H,P363L,P363S,T364A,T364I,T364P,S365N,S365R,P367H,P367S,H368Y,A369V,V370A,C372W,I374V,R377I,Y379C,Y379H,L380F,P382L,K383E,G384S,T385S,T386N,I387L,I387V,L388V,I389T,S390C,S390F,T392I,T392N,V394M,H396R,D397A,D397N,N398I,E400G,E400K,N403K,P404L,P404T,E405V,M406I,F407S,D408E,D408N,P409H,H410R,H410Y,F412C,L413P,E415K,G416C,G416S,G417D,N418T,F419I,F419S,K420M,S422G,K423N,K423R,Y424H,M426K,P427L,P427S,F428L,F428S,G431R,R433Q,R433W,I434M,I434F,C435G,C435S,V436A,V436M,G437R,E438G,A441V,G442S,M443I,M443V,L445P,L445Q,L445V,F446L,L447F,L447S,T450I,T450N,S451F,S451T,Q454H,Q454R,N455K,N455T,N457S,L458V,K459T,L461V,K465E,N466K,N466T,L467F,L467P,D468H,T469N,T470I,P471A,V472A,V473A,N474S,G475R,F476C,F476L,A477T,A477V,S478T,V479L,P480L,P480T,P481S,P481T,F482L,C486Y,I488F,I488V,P489S,V490F. |
| **CYP2C19** | |
| Non-pathogenic | M1V,D2H,P3L,P3S,F4L,F4S,V6F,L7F,L7I,L7P,V8A,V8L,L9P,L9V,C10R,C10S,C13S,L16F,L17H,L17P,I19L,W20C,R21I,R21T,Q22R,S23G,S23R,G25E,G25R,G25V,G25W,R26T,G27E,K28E,K28I,L29F,P30L,P31H,G32D,G32S,P35T,V38M,G40R,I42V,Q44H,I45V,D46E,D46G,D46V,K48N,D49V,V50I,S51G,K52Q,T55N,T55S,L57P,L57V,K59E,I60N,G62A,G62D,P63L,V64M,F65L,T66I,T66S,G70A,G70V,L71V,R73C,R73H,R73S,M74T,V75M,V76L,H78L,H78Q,H78Y,E81K,V82A,V82M,A86V,L87Q,L87V,I88T,D89H,G91R,E92D,E92G,E93D,F94L,G98D,G98S,H99L,H99R,H99Y,F100Y,L102P,L102V,R105G,A106G,N107S,R108G,G109A,G109R,F110Y,G111E,G111R,I112L,I112T,V113I,F114C,S115G,S115I,N116S,G117A,R119K,W120L,W120R,E122A,E122K,I123M,I123T,I123V,R124G,R124Q,R124W,R125C,R125G,R125H,R125P,S127L,L128F,L128P,T130A,T130K,T130M,T130P,L131P,R132Q,R132W,N133K,N133Y,F134L,G135E,M136K,M136T,G137E,G137R,R139T,S |

| | |
|---|---|
| | 140N,I141T,E142D,E142G,D143E,D143Y,R144C,R144H,R144P,R144S,V145I,E147G,E147Q,E148K,A149T,R150C,R150H,V153A,V153L,E154G,E154Q,E155K,E155V,K158R,K160E,A161P,A161T,S162P,P163L,C164F,C164G,P166H,P166L,T167I,F168L,F168S,G171D,A173D,A173V,P174A,P174H,P174L,C175F,N176S,C179F,C179G,C179W,Q184R,K185Q,R186C,R186H,R186L,R186P,D188N,Y189C,Y189F,K190E,K190R,Q192H,Q193R,L195F,N196H,N196S,L197F,M198L,M198R,M198T,L201W,E203G,E203K,N204I,I205S,V208L,S209N,T210N,P211S,P211T,C216W,C216Y,F219Y,P220S,T221I,T221S,I222L,I223T,D224V,F226I,F226Y,P227L,P227S,N231K,L233F,K235E,N236K,N236S,N236T,A238T,A238V,E241D,E241K,S242R,D243G,D243V,L245F,E246D,E246Q,V248I,E250A,E250K,H251Q,S254L,S254P,M255K,M255T,D256G,D256H,D256N,I257F,N258D,N258S,N259K,P260L,P260S,P260T,R261Q,R261W,D262E,D262N,D262Y,I264F,I264T,D265G,F267L,M271I,K275M,K275N,Q276R,N277I,N277K,Q279H,Q279L,Q279P,S280C,S280F,S280Y,E281K,T283I,T283P,I284T,I284V,E285K,N286K,V288I,I289S,I289V,D293N,L295F,L295P,L295V,G296A,G296R,A297T,A297V,G298A,T299A,T299I,T299Q,T299R,T302A,T302I,T302R,S303G,S303N,S303R,T304R,T305A,T305I,Y308C,A309T,L310F,L310I,L313P,L314P,L314R,H316L,E318D,E318Q,V319A,V319I,T320A,T320I,A321S,A321T,K322I,K322R,Q324R,E325K,E326G,E326V,I327T,R329C,R329H,R329S,I331L,I331M,I331T,I331V,R333I,N334I,N334K,R335Q,R335W,S336I,S336N,S336R,P337H,P337S,C338F,C338Y,M339R,M339V,Q340K,D341H,D341N,R342K,G343D,G343S,H344Y,M345I,M345K,P346L,P346S,P346T,T348A,D349A,A350G,A350T,A350V,V351M,H353R,E354A,E354K,E354Q,V355A,Q356P,R357G,Y358H,I359N,I359V,D360A,D360E,D360H,D360N,D360V,D360Y,L361I,L361R,I362F,I362T,P363S,T364S,S365T,L366M,P367S,A369V,T371A,T371N,T371S,C372S,D373E,D373G,V374I,F376Y,R377G,L380V,K383N,K383R,G384R,G384S,G384V,T386A,I387L,I387T,I387V,T389I,S390F,L391F,L391V,T392A,T392S,S393F,S393P,V394A,V394M,L395P,H396R,H396Y,D397G,D397N,D397Y,N398K,N398S,N398Y,K399R,E400V,P402A,P402L,P402S,N403I,E405G,E405K,E405Q,M406I,M406R,M406V,F407S,D408E,D408G,P409S,P409T,R410C,R410G,R410H,R410L,R410P,H411Q,L413V,E415D,G416D,G416S,G416V,G417E,N418D,K421N,K421R,S422N,S422V,N423H,N423K,Y424C,M426I,P427S,P427T,A430P,A430S,G431E,G431R,K432I,R433Q,R433W,I434M,C435Y,G437R,E438V,G439D,G439V,L440V,A441T,A441V,R442C,R442G,R442H,R442L,M443I,E444Q,L445M,L445P,L445V,L447S,F448L,F448S,F448Y,T450I,T450N,T450P,F451I,F451S,I452T,Q454R,N455K,N455Y,N457K,L458R,L458V,I462M,D463G,P464S,P464T,K465E,D466E,D466G,D466N,D466V,T470P,V472I,N474S,G475R,A477T,V479F,V479I,P480L,P481L,Y483C,F487L,P489S,V490A,V490L. |
| **CYP2D6** | |
| Non-pathogenic | A5T,A5V,L6V,V7A,V7M,P8H,P8L,P8S,A10V,V11M,I12L,A14V,F16V,L18F,L19V,L22P,M23R,H24R,R25Q,R25W,R26C,R26H,Q27R,R28C,R28H,R28L,R28S,W29G,A30V,A31E,A31T,A31V,R32C,R32H,P34A,P34L,P34Q,P34R,P34S,P35L,P35S,G36C,G36D,P37H,P37S,P39L,L40Q,L40R,P41L,P41S,G42E,G42R,L43R,G44S,H48Y,V49M,D50G,F51L,Q52R,P55L,P55S,Y56C,Y56H,C57R,C57Y,F58L,D59H,Q60R,R62G,R62P,R62Q,R62W,R63C,R63H,R64C,R64H,F65S,G66R,D67N,D67Y,V68E,V68G,F69L,F69S,S70I,S70N,S70R,L71V,A74V,W75C,W75R,T76M,T76S,P77L,V78A,V78L,V78M,V80M,L81F,N82S,N82T,A85T,A85V,A86V,V87M,R88C,R88G,R88H,R88P,E89G,E89K,E89Q,A90V,L91M,L91P,V92L,V92M,T93I,T93P,T93S,H94D,H94L,H94R,H94Y,G95R,G95S,E96D,E96K,D97E,D97G,D97Y,T98I,A99D,A99T,D100A,D100E,D100Y,R101C,P102L,V104A,V104E,V104M,P105H,P105L,P105S,I106N,T107I,T107N,T107S,Q108H,Q108R,I109T,I109V,G111C,G111S,G113E,G113R,P114A,P114L,P114R,R115C,R115H,S116C,G118A,R119C,V119L,V119M,F120I,F120L,P120L,P120S,R122C,A122E,R122H,A122S,A122V,R123C,R123H,R123L,P123L,P123S,P123T,Y124H,N124S,N124T,G125E,G125R,G125S,L126F,P126T,L126V,L127F,A127T,A127V,L127W,D128N,W128R,R129C,R129H,R129L,K129N,R129P,E130D,E130G,E130K,V131A,Q131L,V131M,R132K,R132M,S132R,R132T,R133C,N133D,R133G,R133H,N133H,V134L,V134M,S135F,V136M,A136T,S137A,S137F,S137P,L138F,L138R,T139A,T139I,T139N,L139S,R140C,R140H,R140S,C140Y,N141K,G141R,N141S,R1 |

| | |
|---|---|
| | 42C,R142H,L142S,G143A,R143C,R143H,F144L,E145A,G145D,E145K,E145Q,G145S,D147E,D147G,D147N,K147R,D148A,D148H,S148L,D148Y,P149A,P149H,L149M,P149T,R150C,R150G,R150H,E150K,R150P,R150S,Q151E,Q151R,L152F,W152G,L152V,R153G,T154A,T154I,E155A,E155K,E156A,E156D,E156K,D156N,D156V,L157P,L157V,A158T,A158V,L160F,E160G,E160K,G161A,G161E,C161F,G161R,C161S,L162P,A162T,K163E,A163T,A163V,E164K,F164L,E165A,E165D,A165G,E165K,A165P,A165T,A165V,N166D,S166L,S166P,S166W,G167D,H167Q,G167R,G167V,S168A,F168S,S168Y,G169A,G169R,R170C,R170H,E171D,E171K,P171L,E171Q,P171S,V172G,V172L,V172M,R173C,R173H,L173P,P174L,P174S,N174T,P174T,N175S,A175T,N175T,A175V,G176R,G176S,L177F,P177L,P177S,L177V,V178A,L178F,V178I,L178W,L179F,L179I,D179N,K180N,L180P,L180V,H181L,H181R,V182A,I182L,V182M,I182N,P183L,S183R,N184D,A184G,N184H,A184T,A184V,V185L,L185M,V185M,L185P,A186P,A186S,G187S,A187T,S188A,S188F,S188P,L189F,V189I,L189R,T190A,T190I,T190N,L190R,R191C,R191H,C191Y,G192R,R193C,R193H,R194C,R194H,F195L,E196A,E196K,E196Q,D198E,D198G,D198N,T198S,D199A,D199H,D199Y,P200A,P200H,P200T,R201C,D201E,D201G,R201G,R201H,D201N,R201P,R201S,L203F,L203P,L203V,R204G,L204I,L204Q,E206Q,H207D,D207N,D207V,L208P,R208S,L208V,M209R,A209V,T210I,T210S,E211G,E211K,W211R,G212A,D212A,G212E,D212N,G212R,P213L,L213P,P213T,K214E,A214P,A214V,Q215H,E215K,Q215P,Q215R,P216A,E216A,E216D,P216H,E216K,P216L,P216R,P217H,P217L,S217L,S217P,P217R,P217S,S217W,G218D,R218G,R218P,R218Q,G218R,G218V,D219G,D219H,F219S,L220P,R221C,R221H,T221I,E222D,E222K,E222Q,E222V,V223G,V223L,V223M,A223V,L224P,N225T,A226E,A226T,A226V,E227D,E227K,M228I,M228K,P228L,P228S,M228T,V229A,E229D,V229I,K230E,L230F,L230I,L231P,L231V,H232L,H232R,I233L,I233N,G233R,P234L,N234S,N234Y,A235G,P235L,A235T,P235T,A235V,L236M,L236P,A237P,S237R,A237S,S238R,G238S,F239I,V240I,N240S,D241N,L241R,R242C,R242H,N243D,L244V,C245R,C245S,C245Y,I246L,I246M,I246T,V247A,V248L,A249D,A249G,T249S,T249P,D250H,D250N,L251P,D252E,D252G,F252L,D252N,F252S,S253C,S253F,S253P,L254P,A254V,L254V,L255I,L255Q,G255R,M256I,M256T,M256V,E257Q,H258D,T258N,R259S,S260L,M260R,T261I,T261S,T262M,T262R,W262R,D263A,D263N,L263P,P264L,P264T,A265P,A265V,G266D,Q266H,Q266P,Q266R,G266S,P267A,L267H,P267H,P267L,P267R,P268H,P268L,P268R,P268S,L269F,R269G,R269P,R269Q,D270G,D270H,M270T,L271P,T272I,H273P,E273V,H273Y,P274L,P274R,A274V,D275A,D275N,L276P,A277E,A277T,A277V,R278C,E278D,R278G,E278K,R278L,R279C,R279H,M279I,M279K,R279P,M279T,E280D,K281E,Q282H,Q282R,E283A,G284R,D285G,D285N,N285S,N285Y,D286G,P286L,D286N,P286T,V287M,I288L,S288R,I288T,I288V,G289E,G289R,S289R,Q290E,F290I,V291A,V291L,V291M,N291S,R292G,D292N,R292Q,R292W,R293Q,P294A,N294D,E295Q,L295V,R296C,R296H,M296I,M296K,R296S,M296T,M296V,I297L,I297M,I297T,V298A,D298E,D298N,V299L,A300D,A300G,D301H,D301N,H301R,M302I,L302P,M302R,F303L,P303L,F303S,P303S,Y304C,S304C,Y304D,S304F,S304P,T305I,A305V,G306R,M307I,A307S,M307T,A307V,M307V,V308L,V308M,T309N,H310Y,E311D,S311L,E311Q,V312E,T312I,V312L,V312M,T313M,T313R,Q313R,R314C,R314H,R314P,L314P,R314S,F315L,F315S,G316R,G317D,G317S,D317Y,I318F,L318H,I318M,I318N,I318S,V319I,L320F,P320L,P320S,L321R,M321T,G322D,G322S,V323L,V323M,T324A,H324P,H324Y,P325L,H325R,P325R,D326A,M326K,D326N,M326V,T327I,V327M,R329C,R329G,R329H,R329L,R330C,D330G,R330H,D330N,R330P,I331V,E332D,E332K,E332Q,Q333H,Q333R,E334A,D336G,D336N,R337C,D337G,R337H,D337N,R337P,I338F,V338M,I338M,I339L,I339T,I339V,G340E,G340R,Q341E,G341E,V342A,V342L,V342M,T342M,T342P,T342R,R343G,T343I,R343Q,T343R,R343W,L344F,R344Q,P345A,T346I,T346N,E346Q,M347I,M347K,M347T,M347V,D349E,S349L,D349N,S349P,S350L,V351L,H352R,L352V,M353I,M353R,P354L,P354S,Y355C,Y355D,E355K,T356I,A356V,V357I,V357L,A358S,A358V,E359K,V359L,V359M,K360Q,P361L,P361S,H361Y,E362D,E362Q,F362S,R363C,V363E,R363H,V363L,V363M,Q364R,R365C,R365H,R365P,H365Q,H365R,R365S,H365Y,P366L,F366L,F366S,E367A,E367K,E367Q,G367R,H368Q,H368R,H368Y,D368Y,I369F,I369M,I369N,I369R,F369S,I369S,I369T,V370I,L370P,P371L,P371S,L372R,G373D,Q373P,G373S,G374D,V374L,V374M,T375A,H376R,F376S,M377K,V377L,V377M,M377V,K378E,T378I |

| | |
|---|---|
| | ,K378M,P379L,P379S,R380C,R380H,D381G,D381N,I382V,E383D,E383K,E383Q,L383Q,P384L,P384S,G386D,A387E,R388C,G388D,R388H,R388P,G388R,R389C,I389F,R389H,I389M,R390C,R390H,A391T,G392E,C392R,T393M,T393P,T393R,G394E,T394I,G394R,T394R,G394V,G394W,L395F,P396S,T397I,T397N,A398D,R399C,R399H,R399S,M400I,S400L,S400P,S401L,V402L,F403L,L403V,F403V,L404V,F405L,E406K,T407A,T407N,A407V,V408I,V408L,L409M,L409V,E410K,L410P,Q411H,K411Q,P412L,H412Q,P412S,F413S,R414C,R414H,S414T,S416L,H416Q,H416R,H416Y,P417L,P418A,E418A,E418K,E418Q,P418T,T419A,H419Q,H419R,H419Y,G420R,F420S,L421P,Q421R,P422H,P422S,R423Q,R423W,Q424P,P424S,G425D,S425T,H426N,H426P,H426Y,H427N,H427P,H427R,F427S,H427Y,G428A,G428C,G428D,V428L,V428M,K429E,V429I,K429M,F430L,P430L,P430S,F430S,A431G,A431P,A431T,F432I,F432L,L433R,V434L,L434Q,P435L,T435S,P435S,P436L,P436S,S437F,S437Y,A438E,P438L,Y439C,G439D,G439R,R440C,R440H,E440K,R441C,R441H,A442T,A443G,C443R,V444A,V444G,G445E,G445R,P445S,G445V,G445W,R446C,R446H,R446P,P447S,A449D,R450C,R450H,R450S,M451I,F454L,F454V,L455V,F456L,F457L,T458A,T458N,L460M,L460V,L461P,Q462H,H463Q,H463D,S465N,S465T,S467L,P469A,P469T,T470A,G471R,Q472R,P473H,P473S,R474Q,R474W,P475S,S476T,H477N,H477P,H477Y,H478N,H478P,H478R,H478Y,G479A,G479C,G479D,V480I,F481L,F481S,A482G,A482P,A482T,F483I,F483L,L484R,V485L,S486R,S486T,P487L,P487S,S488F,S488Y,P489L,Y490C,E491K,A494G,V495A,V495G,P496S,R497C,R497H,R497P. |
| **CYP2E1** | |
| Non-pathogenic | L4F,L4R,G5R,V8M,L11M,V12L,V12M,W13L,A14V,A15V,L17F,L19P,V20M,M22I,M22T,M22V,V26G,V26M,S28G,S28N,S29R,S29T,N31I,P33S,P34T,P36A,F37L,P38L,P38R,P40R,G43R,N44K,Q47E,N52S,N52T,I53T,P54L,P54S,S56C,S56F,F57L,T58S,R59P,R59Q,R59W,Q62E,Q62R,R63C,R63G,R63H,F64L,G65R,T69M,V72L,V72M,G73A,G73D,S74L,Q75P,R76C,R76H,R76L,R76P,M77V,V78M,M80I,H81D,H81L,H81Q,G82D,G82S,Y83H,K84R,A85V,V86L,A89V,E96D,S98L,G99A,G99D,L103F,P104L,P104R,P104S,A105T,A108E,A108V,H109L,H109R,R110G,R110K,D111A,G113R,F116C,P120L,P120S,T121I,K123E,R126L,R126Q,R126W,R127Q,R127W,L130V,T132I,L133F,L133I,R134Q,R134W,Y136C,M138L,G139E,G139R,K140R,Q141H,Q141P,G142A,G142S,E144D,R146Q,R146W,E150A,A151T,H152Q,H152R,F153V,L154R,E156V,L158I,R159G,R159S,T161S,Q162R,P165A,P165R,P165S,F166I,D167A,D167E,D167G,D167N,T169N,T169P,T169S,L171F,I172M,I172V,G173C,G173S,C174S,A175T,A175V,P176S,N178S,V179F,V179I,I180M,D182G,D182N,I183T,F185L,R186C,R186H,D190N,N192S,E194D,K195M,R198K,M200I,M200L,M200R,H208P,H208Q,H208R,H208Y,L209Q,S211G,T212A,P213R,P213S,Q216K,N219D,N219S,N220T,F224S,H226Y,L228S,L228W,G230E,G230R,S231N,R233S,R233T,K234R,V235I,I236L,I236T,N238M,N238Y,V242I,E244D,V246L,V246M,R249S,K251N,E252D,H254Y,L257Q,L257R,N260S,P262L,P262S,R263Q,R263W,D264H,L265F,D267N,D267Y,C268S,C268W,V271M,M273V,E274Q,K277N,K277R,H278Q,H278R,S279R,A280V,R282C,R282H,R282S,T285A,T285I,M286V,G288S,T290I,T290S,V291M,V293L,A294V,D295N,A299S,A299V,G300R,T301S,T303N,T304S,S305I,S305N,T307I,L308V,R309I,G311R,L312F,L313Q,I314S,M316L,Y318H,P319A,P319H,I321M,I321V,E322K,K324E,L325V,E328D,I329T,D330Y,V332L,I333M,I333T,I333V,G334R,P335A,P335T,R337L,R337Q,A340V,I341M,K342E,D343N,R344K,Q345R,E346D,M347L,M347T,Y349C,M350R,M350V,V354L,V354M,R359P,R359Q,R359W,I361F,T362I,T362P,V364A,V364M,P365S,S366C,N367D,L368P,P369L,E371G,T373A,R374G,R374Q,D375E,D375G,F378L,G380E,G380V,Y381H,L382F,I383M,K385R,V388A,V388I,V389I,V389L,L393V,D394E,D394G,V396L,Y398F,N400S,Q401P,D405E,P406A,E407G,K408E,F409L,H413R,N416S,N418S,K420E,K420M,F421L,Y423H,S424G,P429A,V436A,C437F,G439R,E440G,A443D,R444C,R444H,R444L,R444P,M445I,L447F,L450S,C452S,A453T,A453V,Q456K,H457L,H457Y,P462S,V464A,V464I,D465V,P466L,P466S,K467N,D468G,D468N,D470N,D470Y,L471F,I474L,H475P,H475Y,I476V,G477R,G477V,P483S,R484C,R484H,R484S,K486E,C488W,C488Y,P491L,R492C,R492H,S493L. |

| | |
|---|---|
| **CYP2R1** | |
| Non-pathogenic | W2C,W2G,W2R,K3N,L4I,W5C,W5G,W5S,R6K,R6S,R6T,A7V,E8G,E8K,E9K,G10A,G10D,G10V,A11E,A11V,A12P,A12V,A13E,A13G,L14P,G16S,A17T,A17V,L20P,L20V,L21M,L21R,F23Y,A24V,L25V,G26E,V27I,R28S,Q29H,Q29P,Q29R,L30P,R34K,P36L,M37I,M37T,G38S,G38V,P40H,P40S,P40T,P41A,P41L,P41R,P41S,P41T,G42E,G42R,P44A,P44L,L46V,P47Q,I49V,G50S,N51S,S54F,L55V,A57S,S58A,S59P,E60K,P62H,H63R,H63Y,M66I,M66L,M66R,R67K,Q71H,V72L,Y73C,E75K,S78N,S78R,L79I,G82E,G83R,I84L,I84M,S85L,S85P,S85T,V87M,G91D,Y92C,D93G,V94A,V95A,V95I,E97G,L99F,V100A,H101D,H101N,H101R,H101Y,Q102H,Q102R,E104K,I105T,D108G,R109G,R109S,P110L,M116I,M116K,M118I,M118K,M118V,T119K,G122R,G123R,L124F,L124S,L125P,N126I,S127C,S127F,Y129F,R131Q,W133S,V134A,V134I,D135H,R138L,R138Q,L139F,A140T,V141I,V141L,S143C,S143G,S143R,F144L,R145Q,G148V,G150D,K152T,S156C,I158M,L159F,T162N,F165L,F165S,N166H,N166S,A168T,I169M,I169T,I169V,T171I,Y172C,Y172N,P176R,D178E,F179S,Q181H,Q181R,L182S,T184M,V187F,V187I,V187L,N192S,N192T,L193M,I194M,R199L,R199Q,F200L,T201A,T201I,D206G,D206N,M210V,F214C,S215G,L220I,L220V,A221T,A221V,A222T,A224V,F227S,Y229D,I237M,I237T,P239T,G241E,G241R,K242N,H243Q,R248G,R248S,A250V,V252A,D255A,L257F,S258F,R259K,I261T,K269R,Q274R,V277I,A279P,E283K,M284I,M284T,M284V,D285G,Q286E,Q286P,G287D,G287S,G287V,N289M,S293C,T294A,T294P,F295L,K297R,E298D,L300R,V304L,E306G,I309V,T312N,T316A,T316S,N317D,N317S,V318G,R320Q,R320W,W321R,A322V,L324V,F325L,F325S,M326V,A327V,L328I,P330A,P330R,N331D,I332T,Q335R,V336A,Q337H,Q337L,E339Q,I340L,I340T,D341G,I343T,M344I,M344T,G345C,N347I,N347Y,G348R,K349T,D354E,D354N,M358I,P359L,P359S,H366Q,E367Q,C372F,C372S,I374R,I374T,I374V,V375L,P376A,P376S,G378E,G378R,G378V,I379F,H381R,A382E,T383A,T383I,S384C,D386G,A387G,V389E,R390C,R390H,S393F,I394L,I394T,I394V,G397D,T399P,T399S,V400L,T402I,H408N,H408Q,H408R,F409L,Y413H,Y413N,V419L,R424Q,D427G,Y431D,Y431H,A433G,K434R,K435E,E436G,A437S,V439I,P440R,P440T,F441S,S442P,S442T,G444R,R445G,H447D,H447N,H447R,C448R,L449R,G450R,E451K,R455Q,R455W,M456T,E457G,M458I,M458R,L460S,F461C,T463Y,L466P,R468K,F469L,F469V,L471S,H475Q,P479A,P479L,P479Q,M487I,M487V,P491T,P493A,P493T,Y494N,I496V,C497R,A498P,R500S,R500T,R501C,R501H. |
| Pathogenic | L99P,G42R,P43R,P44R,G45R,L46R. |
| **CYP3A4** | |
| Non-pathogenic | A2D,A2G,A2T,A2V,L3P,L3V,I4T,P5S,D6E,A8P,E10Q,T11I,W12C,L15P,V17I,S18R,L19Q,V20A,V20L,V20M,L21V,L22V,T27I,H28Q,H28R,S29T,H30D,H30R,G31E,K34E,K35R,L36F,L36R,L36V,I38V,P39L,G40E,P41L,T42A,T42I,T42K,P43L,P43R,P43S,P45L,G48E,N49D,N49T,I50T,Y53H,H54R,K55Q,G56D,F60L,M62K,E63K,C64S,K66R,K67N,K67R,K67T,Y68C,Y68H,G69R,W72G,W72L,G77R,G77V,Q78H,Q79R,V81A,V81L,L82V,M89V,I90T,I90V,K91Q,K96E,E97D,C98R,C98Y,Y99H,S100F,V101F,R105L,R105Q,R105W,R106K,R106T,P107L,P107R,G109D,G109R,G109S,G112R,F113I,F113L,F113S,M114K,S116C,S116T,A117T,A117V,I118F,I118V,S119T,I120T,A121G,E122D,D123E,D123Y,E124G,K127N,L129I,R130G,R130P,R130Q,L132W,S134C,P135L,P135S,T136A,T136I,S139G,S139T,M145V,P147L,I148V,A150S,Q151R,V155A,L160P,R162Q,R162W,T166I,G167A,G167D,G167R,K168N,P169A,P169S,P169T,V170I,K173E,K173R,D174E,D174H,D174N,V175D,V175I,F176L,A178S,A178V,Y179S,S180N,M181T,I184T,T185A,T185S,S186G,S186N,S186R,S188L,F189L,F189S,V191A,V191E,N192K,N192S,I193M,I193V,D194N,D194Y,N198D,Q200E,Q200H,D201N,P202L,P202R,P202S,E205D,E205G,E205K,N206Y,T207I,T207N,K208E,L210P,F213Y,D214G,D214N,L216F,D217N,P218R,F219L,F220C,F220V,S222P,S222T,I223L,I223M,I223V,F226L,P227L,P227T,F228L,L229F,L229R,I230V,P231L,I232V,L233P,E234G,V235A,I238T,C239F,P242L,R243T,N247D,N247I,K251T,S252A,S252T,V253A,R255K,R255T,E258V,S259G,R260C,R260H,E262K,D263N,H267N,R26 |

| | |
|---|---|
| | 8Q,L272P,Q273R,D277N,S278P,K282E,E285G,S286P,K288R,A289S,A289V,S291C,S291F,D292E,D292N,L293P,V296L,V296M,S299P,S299T,I300V,I301L,A305S,T309A,T309I,T310K,T310M,S311N,V313G,L314F,L314P,S315A,S315F,Y319C,T323P,H324Q,Q328R,Q332R,E333K,E333Q,E334K,I335T,D336E,D336H,A337E,A337V,K342N,P345L,P345S,Y347C,T349N,M353I,M353L,E354K,Y355C,Y355H,D357E,M358I,M358V,V359E,T363K,T363M,L364F,I369V,A370S,A370V,M371R,R372G,R372I,R372T,L373F,E374D,R375K,R375M,K379R,E382D,M386I,I388F,P389L,P389S,V392M,M395I,M395V,I396T,P397L,S398N,S398R,Y399C,L401F,L401P,R403C,R403H,R403P,P405T,W408R,P411A,P411L,E412K,P416L,P416R,R418T,F419L,S420G,S420I,K421R,K424R,D425E,D425N,N426K,N426S,I427V,D428E,D428H,D428N,P429L,P429R,P429S,I431T,Y432F,T433A,T433I,P434A,G436R,S437T,G438V,P439S,N441D,G444A,M445I,M445T,M445V,R446K,L449F,M450T,M450V,M452L,M452T,M452V,K453N,L454F,L454I,L454P,I457V,R458I,V459L,L460F,N462K,F463C,S464F,S464T,P467A,P467S,C468W,C468Y,E470K,Q472H,Q472R,I473F,I473M,I473N,S478C,S478G,S478R,G480E,G481R,Q484R,P485R,K487E,P488H,V489I,V490F,L491P,L491Q,L491V,V493A,S495T,V500A,V500I,S501G,S501N,G502E. |
| Pathogenic | I301T |
| **CYP3A5** | |
| Non-pathogenic | L3I,P5Q,L7S,A8V,V9E,V9L,E10K,L15R,Y25H,T27N,R28C,R28H,T29I,H30Y,F33L,F33S,L36R,I38V,T42I,P43H,P43L,L46F,L46M,V50I,L51F,S52F,Y53C,Y53H,R54C,R54G,R54H,R54S,G56C,G56D,W58S,K59T,F60C,D61E,C64F,C64S,K67R,K70N,G73R,T74K,T74M,G77A,P80R,L82R,L82V,T85R,D88H,D88N,V89M,R91K,V93L,L94Q,S100C,S100Y,T103I,T103K,N104I,R105G,R105Q,R106S,S107P,G112V,M114V,S116T,L120S,D123E,D123V,E124D,W126R,K127R,I129T,R130Q,L132V,L133P,T136A,T136N,T138S,G140R,K141T,K143R,E144G,E144K,M145T,F146L,I148M,I148N,I148T,I149T,I149V,A150D,A150S,Q151L,Q151R,Y152H,D154N,R158I,N159H,L160W,R161G,R162Q,R162W,A164V,E165G,E165Q,K166N,K168R,P169S,V170I,K173R,D174G,I175S,Y179F,S180I,D182N,V183G,T185I,G186A,T187I,S188T,F189L,V191M,I193L,I193V,D194H,D194N,S195C,S195F,S195P,N197K,Q200R,D201H,D201N,P202L,P202S,F203C,F203S,E205D,S206R,F210S,K212N,F213V,G214V,L219S,F220C,I223V,L225F,P227S,P227T,F228L,F228S,L229F,L229P,T230I,P231T,F233L,F233S,V238A,V238D,L240P,K243E,T245A,T245I,I246K,I246V,F248C,F248L,S250N,K251Q,R255G,M256K,K257M,S259G,R260H,R260P,N262S,D263N,H267Q,R268L,R268Q,Q273R,L274P,M275I,M275R,I276S,I276T,D277E,D277G,Q279R,N280H,S281L,K282N,E285K,H287L,A289P,A289V,E294K,L295F,L295I,A296T,A297V,S299A,S299T,I303L,F304S,Y307H,T309I,T309N,T310I,S311R,S311T,L314I,S315C,T317I,T317S,Y319H,A322D,P325S,D326N,V327I,K330E,L331P,Q332L,Q332P,I335T,D336Y,A337T,L339F,P340S,N341S,A343S,A343V,P344L,P344Q,P345A,P345L,P345T,T346A,Y347C,V350A,V350M,Q352P,Y355C,D357N,M358L,M358V,V360M,N361S,E362G,T363I,L364H,A370P,I371T,I371V,L373I,R375G,C377R,K378E,D380G,V381I,N384H,G385R,V386I,F387I,F387Y,I388T,P389S,S392P,S392T,M393I,M393V,V394L,V395L,T398N,L401P,H403R,D404Y,K406E,T409S,P411R,P411S,E412K,R415C,R415H,R415L,R415P,P416S,R418M,F419L,S420I,S420R,K423E,D424N,S425N,I426M,D427H,D427V,P428A,Y429S,I430T,I430V,Y431N,T432I,T432K,G435E,G435R,T436I,T436N,T436S,R439K,I442T,G443D,G443S,M444V,F446S,A447V,M449K,M449T,L453I,L453R,L453V,A454V,L455P,L455V,V458A,N461H,P466H,C467R,I472T,P473A,P473L,P473S,L476S,T478M,P484Q,P484S,K486E,P487S,I488T,V489I,V492M,D493N,R495T,G497V,T498N,L499P,L499V,S500G,G501E. |
| **CYP3A7** | |
| Non-pathogenic | D2G,D2Y,A8S,A8V,V9M,E10G,T11I,W12C,W12G,W12L,L13I,L13R,V17I,S18R,L21F,L21V,L22F,Y23C,R28C,R28H,R28L,R28P,H30R,G31R,L32V,F33V,L36P,G37R,P39S,G40R,P45S,L47F,N49D,A50S,A50V,S52F,S52Y,R54C,R54H,R54L,G56D,Y57H,T59M,M62I,E63D,C64Y,Y65C,Y65H,K67S,K67T,V71A,I74V,Y75F,C77G,C77R,Q78H,Q78L,Q79R,P |

| | |
|---|---|
| | 80L,M81V,A83G,I84V,D88N,D88Y,M89I,M89T,I90V,K91R,V93L,L94I,L94P,E97D,C98R,C98S,Y99F,S100P,V101A,R105Q,R105W,P107L,G109R,P110L,V111A,G112E,F113Y,N116H,A117S,A117V,S119T,E122K,D123H,D123N,E125K,W126C,I129K,I129L,I129T,R130Q,S134P,T136A,T136I,S139R,G140R,E144K,M145R,M145T,V146A,P147L,I149T,I149V,A150V,Q151E,G153E,L156F,R158K,R162G,R162Q,R162W,E163A,E163K,E165Q,G167A,G167S,K168M,P169A,V170D,T171S,K173I,H174D,V175A,V175I,F176C,A178S,A178V,M181T,V183E,T185S,S186I,S186N,T187R,S188T,F189L,G190A,V191A,V191E,V191G,S192N,I193V,D194N,N197D,N197S,P199L,D201E,D201N,D201Y,F203L,E205K,T207A,K209N,L210F,R212I,F213L,N214D,N214S,P215L,P215S,P215T,L216S,D217H,P218A,P218Q,F219C,F219L,V220F,V220I,S222P,I223L,I223V,K224I,V225G,V225I,P227L,F228L,L229P,T230A,P231L,I232V,L233P,A235V,L236I,V240A,R243G,R243I,K244Q,V245L,I246L,I246V,S247R,K251E,K251I,S252T,E258G,G259C,G259S,G259V,R260C,R260H,R260L,L261F,L261I,T264I,Q265E,Q265K,H267Y,R268Q,V269L,D270N,D270Y,L272P,L272R,M275L,M275T,I276N,I276T,D277N,Q279R,S281T,K282E,K282N,D283E,D283V,S284P,T286N,H287Y,K288E,K288T,L290P,L290R,L290V,E294D,L295V,M296I,M296T,M296V,A297V,F302S,I303L,F304V,G306V,E308A,T310K,T310M,T310R,S311G,L314F,L314P,S315F,F316L,I317V,I318L,I318M,I318T,Y319C,A322T,T323I,T323P,H324R,P325T,D326H,Q328R,V331M,K333Q,K333R,E334D,E334Q,E334V,I335N,I335T,D336G,T337A,T337K,T337R,T337S,V338I,P340S,N341S,P344A,P344L,Y347H,D348N,T349I,Q352H,L353M,E354Q,L356F,L356R,L356V,D357H,M358L,M358V,L364P,R365S,F367L,P368Q,P368S,P368T,V369I,A370T,R372G,R372S,R375G,C377Y,K378E,D380M,N384K,G385E,M386I,M386V,F387L,I388T,I388V,P389L,G391E,G391R,V392M,V393A,V394L,V394M,M395I,M395T,I396L,S398T,V400A,V400I,L401F,L401P,L401V,H402L,H402Y,H403D,H403R,K406N,K406T,Y407H,T409R,P411A,P411H,E412Q,P416L,E417D,S420G,K422R,N423I,K424E,D425E,D425G,N426K,I427T,I427V,D428V,I431M,I431T,P434S,G438E,G438V,P439S,P439T,R440G,R440K,N441K,I443T,G444D,M445L,V450M,N451K,M452K,M452V,L454F,A455D,L456P,V457I,R458G,R458K,S464F,K466T,P467A,P467L,P467S,I473N,P474A,L475M,K476I,K476T,R478C,R478G,R478H,G480R,L482R,L483H,L484R,T485R,K487N,P488S,P488T,I489V,V490F,E494K,S495A,S495L,S495P,S495T,D497N,E498D,E498K,T499N,V500I,G502E,G502R,A503D. |
| **CYP7A1** | |
| Non-pathogenic | T4I,T4K,S5C,G9R,I10V,A11V,A13T,A13V,C15R,C16S,C16Y,C17S,C17Y,L18V,W19R,R27M,Q28K,T29M,T29R,G30S,P32T,P33S,L34P,L34V,N36K,P40S,Y41H,L42V,A45S,F48V,G49D,G49S,G49V,A50V,P52R,L53F,E54D,E54Q,R57G,A58V,Q60K,H63N,H63Q,H63R,G64D,H65D,H65Y,F67S,T68I,C69F,G73E,G73V,Y75C,Y75F,V76I,H77L,H77R,F78I,I79V,P82S,S84P,Y85C,Y85F,H86N,H86Y,K87E,G92E,G92R,Y94D,D96A,D96G,D96H,D96Y,K99I,K99N,K99T,F100C,F100I,F100S,T104S,S105A,S105C,A106E,A106V,K107N,K107T,A108S,A108T,A108V,H111Q,R112G,R112K,S113N,S113T,I114T,P116L,P116S,M117T,T121I,N124K,N126K,D127G,D127N,F129L,I130F,I130V,Q134K,Q134L,G135D,L138S,T142K,T142M,E143K,S144N,M145R,M145V,M146T,L149I,Q150E,R151C,R151G,R151H,R151P,I152T,M153T,R154K,P155L,V157A,S158C,S158T,N160K,S161A,T163I,T163S,A164D,A164S,A164T,A165T,W166C,W166R,E169D,G170V,M171L,Y172C,S173A,R177Q,V178M,M179I,A182S,G183R,T186I,I187V,R190G,D191G,L192I,T193I,R194K,R194S,R195Q,R195W,D196G,K199R,H201Q,H201R,H201Y,N204H,N204K,D207E,D207H,Q211R,F212I,F212L,D213G,D213H,D213N,K214E,V215A,A218D,A221V,G222D,G222R,P224H,P224L,P224S,I225F,H226Q,F228L,F228V,A231S,A231V,H232Y,N233S,A234V,R235Q,R235W,A239T,S241N,R243K,H244Q,E245A,E245K,Q248L,K249N,R250M,S252C,I253V,S254A,E255K,I257S,S258I,S258N,S258R,R260H,R260L,R260S,M261T,F262L,N264S,D265Y,T269I,D272A,L273P,A276D,A276P,V281M,L283F,W284L,A285T,S286L,I291T,I291V,A293E,A293V,F295S,W296C,W296R,S297C,L298F,Q300K,M301T,R303K,R303W,P305L,E306A,E306D,A307S,A307V,M308T,A310S,E313K,V315M,R317K,R317T,L319S,N321T,A322D,A322V,Q324R,S327G,S327R,L328S,G330D,G330S,P332L,P332S,I333M,L335F,A338T,D342A,P344Q,P344T,L346F,D347N,I349K,I350V,S353L,S353W,L356F,S357P,S358G,S358R,A359S,S360P,L361F,L361V,R364Q,R364W,A366V,K367E,K367M, |

| | |
|---|---|
| | D369E,D369Y,H373Q,L374H,L374P,D376N,G377S,I381L,I381S,R382Q,I386F,I387T,I387V,A388T,Y390H,Q392L,Q392R,M394I,M394V,H395N,L396S,P398A,P398T,E399G,P402L,P402T,D403V,P404A,P404S,L405F,L405W,T406I,T406N,T406S,F407L,F407V,D410G,D410H,Y412H,N416D,G417E,G417R,T419A,T419I,T419K,T422I,Y424C,C425R,N426K,L428F,K429Q,K431N,K431R,Y433C,Y433H,Y434C,Y434F,Y434H,M435T,M435V,F437L,S439L,S439T,G440A,G440E,T442A,I443L,I443V,C444R,P445L,G446R,R447S,L448F,L448W,F449I,A450P,A450S,A450T,A450V,I451L,I451V,E453A,E453G,E453K,I454S,Q456E,L458F,I459F,I459N,M461I,M461T,L462F,L462V,Y464H,I470L,I470T,G472S,A474P,A474V,K475R,P477S,D480E,Q481E,Q481K,R483L,R483P,R483Q,R483W,A484V,G487D,G487S,I488T,P490L,P490R,P491S,N493S,D494V,I495M,I495T,F497L,K498N,K498Q,H503R. |
| **CYP8A1** | |
| Non-pathogenic | W3S,A4T,A5T,G8R,L10V,A11D,A11V,A12S,L16P,R21C,R22C,R22H,R23C,T24A,R25L,R25Q,R26L,R26Q,G28D,P30L,P30S,P31S,D33E,L34P,G35D,G35S,I37F,P38A,P38L,P38R,W39C,W39L,W39S,A43D,D45E,D45Y,A50P,S52R,F53L,T55K,T55M,R56K,M57I,K58R,G62D,G62S,D63N,I64F,I64V,T66I,V69F,V69G,G70W,G71S,R72K,R72S,V74I,V74L,T75I,V76I,L78P,H81Q,S82F,Y83C,D84N,D84Y,A85T,A85V,V86A,V86G,V86M,W88S,E89D,E89K,P90T,R91C,R91H,R91S,T92N,L94P,L94R,D95H,D95N,H97Y,I101T,I101V,E105D,E105K,E105V,R106K,I107S,D109Y,V110M,Q111H,P113S,S118R,D119V,K121R,A122T,R123S,L126P,T127N,L129F,L129I,Q134H,M140T,Y141C,L144H,H145Y,A146E,V147A,D151E,D151G,D151N,A152V,T153I,T153S,E154A,E154K,A155T,G158D,H160D,E161D,E161K,G163C,D166E,D166N,D166Y,Y169N,F171L,A175V,G176D,G176S,Y177S,T179I,T179P,L180F,G182R,E184D,A185G,A185S,A185V,P187L,P187T,R188C,R188G,R188H,H190R,Q193R,A194V,Q195E,Q195R,D196E,D196N,R197C,R197H,V198I,A201T,D202E,V203D,T206A,R208C,R208H,R208P,R208S,L210F,L210I,D211A,D211E,D211N,R212Q,R212W,L214F,L214V,P215S,K216Q,A218P,A218V,R219C,R219G,R219H,R219P,G220D,S221Y,G225E,D226A,D226E,D226G,D226H,K227T,D228H,D228V,H229D,M230I,M230V,C231Y,S232I,V233A,V233I,K234R,S235I,R236C,R236H,R236L,L237M,K239T,L240Q,S242F,S242P,P243T,R245G,R245S,A247P,A247S,A247V,R248W,R249L,R249Q,R249W,A250G,R252G,R252P,R252Q,R252W,W255C,E257V,L261Q,L263V,M266I,M266T,M266V,G267A,G267C,V268E,V268M,S269L,S269P,E271K,M272I,M272T,A274P,R275W,V278M,Q280R,W282R,T284I,T284K,G286E,N287I,N287S,M288I,M288V,P290L,A291T,F293S,L295F,F298L,F298Y,L300F,P303A,P303L,A305T,A305V,A308V,V309D,R310C,R310G,R310H,R310L,R310P,G311E,G311R,L313F,L313V,E314K,S315N,I316V,W318C,Q319K,A320G,A320T,A320V,E321D,E321K,E321Q,Q322H,P323S,S325L,S325W,T327K,T327M,T329A,L330I,L330V,P331A,Q332H,K333T,V334I,L335V,S337G,P339L,S343R,V344A,V344L,V344M,L345M,S346N,S346R,E347Q,L349F,R350S,L351I,T352A,T352R,A353V,A354S,A354V,P355L,P355R,F356I,F356S,T358N,R359C,R359H,R359S,E360K,V361D,V362L,V363L,D364E,A366V,M367V,P368S,M369V,G372R,R373Q,N376H,N376S,R378Q,R379C,R379G,R379H,R379S,D381Y,R382C,R382H,R382L,L384I,L385H,L385V,P387A,F388L,L389M,L389V,S390G,S390N,P391L,P395A,P395S,I397T,P401L,P401T,E402G,V403A,V403I,V403L,F409L,L410R,N411K,P412A,D413E,D413G,D413N,G414E,G414R,E416Q,K422E,R426Q,R426W,L427Q,K428M,K428N,K428R,N429D,N431T,M432I,M432T,M432V,P433L,P433S,G435E,A436R,A436T,A436V,H438N,H438Q,H438Y,N439D,N439S,N439T,H440N,C441S,G443R,R444K,S445R,S445T,Y446F,A447G,A447S,A447T,A447V,V448I,I451T,K452R,K452T,Q453H,F454L,F454S,V455M,F456L,F456S,F456V,L457F,V460M,H461P,L462F,N468I,N468S,A469T,A469V,V471L,V471M,I473N,P474S,F476S,F476Y,D477N,L478F,L478I,Y481C,G482D,G482S,F483L,G484R,G484S,G484V,M486L,M486T,P488L,E489K,H490R,D491E,D491N,V492L,V492M,P493A,V494I,V494L,R495C,R495H,R497C,R497H,R499C,R499H,P500S |
| Pathogenic | R275Q |

| | |
|---|---|
| **CYP8B1** | |
| Non-pathogenic | R26Q,V40M,R59C,R59H,V67M,D82N,S88P,L106T,D134M,L139R,W164C,H165Y,S168I,F172V,F206I,F213L,Y218H,R231Q,R234C,R234H,H237S,K238R,L240F,V267Y,P289L,R306Q,R306W,R349Q,R349W,L357F,S369Y,P400S,V402I,R407C,R415Q,F419L,S435L,G442R,F444L,L447H,D462N,E464Q,P473S,W480C,D490N,R492C,R494C,R494H. |
| **CYP11A1** | |
| Non-pathogenic | A3V,K4Q,L6F,L6R,P7S,P8L,P8S,R9C,R9H,R9L,R9S,L12P,V13L,G15A,G15C,G15S,C16Y,L20M,A22D,A22T,A22V,R24K,E25K,L27P,G28E,G28R,R29C,R29H,R29L,V32M,T34I,E36K,G37E,G37V,A38T,A38V,G39D,I40T,S41F,T42I,T42N,R43C,R43H,R46C,R46H,N49H,N49I,E50D,E50G,S53F,G55C,G55V,D56G,D56N,G58V,L60V,N61K,H64N,H64Y,R67K,E68K,E68Q,T69M,T71K,H72N,V74A,V74I,L76F,H77Q,V79I,F82L,Q83R,K84N,P87L,P87Q,P87R,E91K,G94S,N95I,N95K,N95S,V96A,V96M,V99I,Y100C,I102F,D103N,D106N,V107A,V107M,A108D,K112N,K112Q,E114D,E114K,N117I,L122F,I123M,P124L,P124T,A128T,H130Q,Y133C,I137K,G138R,L141F,S144L,S144T,S144W,A145E,W147G,K148R,D150E,R151Q,R151W,L154P,Q156R,E157K,A160T,E162G,A163T,T164N,K165R,N166S,P169S,L171F,L171M,A173T,V174A,R176Q,R176W,D177E,F178L,F178S,V179I,L182P,R184S,R185C,R185H,R185S,K187R,G190D,G192R,N193I,S195L,G196R,D197H,D197N,S199N,D201V,F203L,R204C,R204G,R204H,F205V,A206S,E208K,T211S,V213I,I214V,F215S,E217A,E217K,R218C,R218H,R218L,Q219H,G220V,M221L,E223V,V225I,V225L,V226A,N227K,P228L,P228R,E229K,A230V,R232Q,F233L,I234V,D235G,I237V,M240L,M240T,H242Y,V245I,P246R,L248P,N249S,L250F,L250H,P251H,P252L,P252Q,P252S,D253H,D253N,R256C,R256G,R256H,R256P,F258L,T262A,K264R,H266D,H266N,H266Y,V267G,V267M,A268S,A269V,V272L,V272M,I279M,Y280C,Y280H,T281N,F284S,Y285D,W286R,R289I,G292R,G292V,V294I,H295N,D297H,D297N,Y298S,R299C,R299H,G300S,I301M,L302V,R304G,L305F,L306M,L306V,S309R,M311V,F313L,D315G,V320I,V320L,T321I,M323K,L324V,A325E,V328A,V328G,V328L,V328M,D329G,T330M,T331M,M333I,M333T,T334I,T334S,Q336H,Y340F,E341K,M342I,R344C,R344G,R344H,R344S,N345K,L346M,Q349K,M351T,M351V,R353Q,A354S,L357S,A358T,A358V,R360Q,R360W,H361Y,A363T,Q364K,Q364R,G365E,D366N,D366Y,M367L,M367T,A368T,T369A,T369M,M370V,Q372R,L373V,P375L,P375R,P375S,L377V,K378R,A379T,A379V,K382N,L385V,H388Q,H388Y,S391C,S391P,V392M,Q395K,L398I,L398V,V399A,V399L,D401E,D401N,V403F,L404I,R405Q,D406H,Y407C,M408I,I409T,A411T,V415M,Q416E,V417G,V417M,Y420C,A421T,L422V,R424Q,E425K,E425Q,P426S,P426T,T427I,T427P,F428I,F428S,F430L,D431N,P432L,P432S,E433G,P437S,T438N,R439L,R439Q,L441P,S442N,K443Q,I447V,T448A,T448N,Y449C,R451Q,N452I,G454D,R460Q,R460W,C462S,L463P,R465Q,R465T,R465W,R466Q,R466W,A468T,L470V,M472V,T473A,I474V,M479L,M479T,V485I,E486D,H489Y,S491I,S491N,S491R,D492N,V493E,T496A,M502T,P503L,P503T,E504A,I507V,T510I,T510P,P513S,F514L,N515I,E517K,Q520H,Q520K,Q521R. |
| Pathogenic | L141W,A189V,L222P,R353W,A359V,V415E,E314K,R451W. |
| **CYP11B1** | |
| Non-pathogenic | A2T,L3F,R4K,R4M,K6Q,A7T,E8D,V9A,V9M,C10G,C10R,C10Y,M11V,P14L,P14S,S17F,S17P,L18P,Q19K,Q19P,A21T,A23T,G25S,T26K,T26M,T26R,R27I,R27T,A28D,A28T,A28V,A29T,A29V,R30Q,R30W,V31F,V31I,V31L,P32A,P32S,P32T,R33G,V35A,L36Q,L36R,P37S,F38I,R43W,R44C,R44H,R51K,R51S,R51T,L52V,L53Q,I55N,W56C,W56S,E58A,E58K,Q59E,Q59H,Q59L,Y61C,Y61H,D63H,L64P,L66M,L66Q,E67K,V68I,H69Q,H69R,T71N,Q73P,Q73R,L75P,P77A,P77S,P77T,D82N,D82V,L83F,A86P,G87D,G87R,M88L,M88T,V89L,C90F,C90Y,M92I,P94Q,P94S,P94T,D96N,V97M,E98D,E98Q,L100M,Q101P,Q102H,V103G,V103L,D104A,S105G,S105R,H107L,H107Q,H107Y,P108T,H109L,H109R,H109Y,R110S,M111I,S112G,S112I,S112R,L113R,E114Q,W116R,V117M,Y119C,R120K,Q121E,H122N,H122Y,R123C,R123G,R123H,R123L,G124E,H125Q,K126E,C127R,C127S,G1 |

| | |
|---|---|
| | 28D,V129A,V129E,V129L,F130L,L132R,N133D,N133K,P135L,P135T,R138C,R138H,L142S,R143L,E147D,E147Q,L149P,P151S,N152H,N152K,N152S,A153G,A153T,Q155H,Q155K,R156K,R156M,M160I,M160T,V161L,R166K,D167G,D167N,Q170R,A171T,A171V,K173N,K173R,K175Q,V176A,N179S,A180D,A180P,A180T,R181P,R181Q,R181W,S183A,S183N,T185I,T185P,T185S,L186V,D187N,V188A,V188I,P190T,S191I,S191N,S191R,I192V,F193L,H194Q,H194R,Y195C,T196S,I197T,E198K,E198V,S200R,A203P,L204F,L204V,G206V,R208Q,R208W,G210D,L211Q,L211P,V212I,G213D,G213R,G213S,H214R,S215G,S215N,S215R,P216L,S217N,S218Y,A219V,N222K,L224F,H225R,L227M,L227P,L227V,E228K,V229G,V229I,M230I,S233T,T234A,T234P,V235A,V235I,M238K,M240T,M240V,P241R,R242S,S243R,R246C,R246H,T248I,T248S,S249N,P250S,P250T,V252A,K254R,E255Q,H256Y,F257L,F257S,F257Y,E258Q,D261E,C262Y,Q265H,Y266H,G267R,G267S,D268G,D268N,C270R,I271M,I271T,Q272K,Y275H,Q276H,E277K,L278M,L278V,A279T,S281N,R282C,R282H,P283S,Q285H,Q285L,Q285R,T287N,S288G,I289T,V290L,V290M,A291S,A291V,L293V,L295M,L295W,N296K,A297E,A297V,E298A,E298Q,L299Q,S300A,S300L,S300W,P301L,P301R,D302E,D302G,D302N,D302V,A303D,A303V,I304M,K305R,K305T,A306T,N307K,S308C,M309I,M309T,M309V,E310D,L311F,L311V,S315R,V316L,V316M,T318K,V320A,V320L,P322L,L323F,L324M,L324R,M325L,T326M,E329Q,A331G,R332W,N333T,P334A,P334T,N335D,N335S,V336A,V336M,A339I,A339S,A339T,A339V,L340R,R341C,R341G,R341H,R341L,Q342H,Q342K,E343K,L345Q,A346T,A347T,A347V,A348T,A349T,S352T,H354L,P355L,P355S,A358E,T359N,T360I,E361G,E361K,P363A,P363T,L364F,R366H,R366L,A367S,A367T,A367V,A368S,L369V,E371A,T372S,R374G,R374P,R374W,L375F,L375I,L375V,P377L,P377S,V378L,F381L,L382R,E383Q,V385L,A386E,A386G,A386V,S387N,S387R,D389G,D389H,D389N,Q393P,Y395H,I397N,P398L,P398T,A399P,A399V,G400E,T401I,T401R,L402S,L402W,V403L,R404C,R404H,R404L,R404S,V405M,L407P,S409T,L410V,G411D,R412C,N413T,P414A,P414S,A415T,L416S,F417L,F417V,P418A,P418S,E421K,R422C,R422H,R422L,R422S,Y423H,N424K,P425S,P425T,Q426S,R427C,R427S,D430G,D430H,I431F,I431L,R432S,R432T,G433D,G433S,G435S,R436S,Y439H,V441M,F443L,F443S,G444C,F445C,F445I,F445S,F445Y,M447I,R448P,L451P,R453W,R454H,L455Q,A456T,A456V,E457Q,E459G,E459K,E459Q,M460I,L463P,H465R,H465Y,H466P,H470P,H470Y,L471R,Q472K,V473L,L476I,T477N,T477P,Q478R,E479D,E479K,D480G,D480N,I481M,K482N,M483I,V484D,S486G,F487S,L489V,S492C,S492N,M493I,F494C,T498P,I502V,N503S. |
| Pathogenic | P42L,P42S,R43Q,F79I,L83S,M88I,P94L,W116C,W116G,H125R,V129M,N133H,P135S,F139L,R141Q,R143W,S150L,L158P,P159L,A165D,T196A,G267D,L299P,A306V,E310K,G314R,T318M,T318P,T318R,T319M,F321V,A331V,R332G,R332Q,R341S,R366C,A368D,E371G,R374Q,G379V,R384G,R384Q,T401A,4427H,V441G,G444D,R448C,R448H,R453Q,R454C,L489S |
| **CYP11B2** | |
| Non-pathogenic | A2P,A2T,R4S,R4T,K6N,E8G,V9M,C10W,V11M,A13V,W15L,L18P,Q19K,R20K,A21T,R22Q,R22W,A23V,L24P,R27G,A28P,A28V,A29T,R30P,R30Q,R30W,A31D,A31T,A31V,P32L,R33S,T34M,V35G,V35L,L36M,L36R,P37L,F38L,A40D,P42R,P42S,P42T,Q43R,H44R,P45L,N47D,R48K,R51K,R51S,R51W,I55F,I55N,I55V,R57K,R57T,E58G,E58Q,G60D,Y61F,E62K,H63D,H65L,H65Y,L66V,M68I,M68R,H69Y,T71N,E74K,G76W,P77S,I78F,Y81C,N82D,N82K,P86A,P86S,R87C,R87G,R87H,R87L,M88K,M88V,C90F,V91A,V91M,M92I,M92T,P94L,P94S,V97L,E98K,Q101K,Q102R,L106P,L106V,H107N,P108L,P108S,C109R,C109S,C109Y,M111I,I112S,P115S,V117M,A118G,A118T,Q121K,R123C,R123H,G124V,H125N,H125R,C127Y,V129L,V129M,F130L,F130S,G134A,G134R,P135S,R138C,R138H,R141P,R141Q,R143Q,R143W,D147E,V148M,K152N,K152Q,A153V,V154A,V154L,V154M,R156G,R156S,L158V,P159L,M160L,M160T,A163V,A165G,A165S,A165T,R166W,D167V,F168L,S169F,Q170H,Q170R,A171T,A171V,L172P,K173E,K173R,K174E,K174R,K175E,K175Q,V176G,V176L,V176M,Q178L,A180T,R181P,R181Q,G182E,S183N,L186R,V188A,V188I,P190H,S191I,F193L,H194Y,T196I,T196N,I197M,I197T,E198G,S200I,S200R,N201D,N201I,N201K,A203T,L204V,F205L,E207D,R208P,R208Q,R208W,G210 |

| | |
|---|---|
| | S,L211P,L211Q,V212A,V212G,G213S,H214D,H214R,S218F,A219T,A219V,L221R,N222T,F223L,H225Q,H225R,A226V,E228G,V229F,M230T,F231V,F231Y,T234P,V235I,L237V,M238I,M238T,M240V,P241S,R242K,R242S,R242T,S243N,S245L,S245P,R246C,R246H,W247R,I248T,S249N,S249R,P250L,P250S,K251R,V252G,V252M,E255D,E255G,H256N,F257V,A259T,A259V,W260R,C262Y,I263N,Q265K,Y266D,G267S,D268N,C270S,C270Y,Q272H,I274T,Y275C,Q276K,Q276R,E277K,E277Q,L278P,L278V,A279V,N281K,N281S,R282C,R282H,R282P,P283S,H285Q,H285Y,Y286C,T287R,G288R,G288S,I289N,V290M,A291E,E292D,E292K,L294M,A297G,L299P,E302D,I304T,N307S,N307T,S308P,G314E,G314R,S315G,S315I,S315N,V316M,D317E,T318R,T319R,A320P,A320V,F321I,L323F,L323S,M325K,T326M,T326R,L327F,L327P,A331V,R332P,R332Q,R332W,P334H,P334T,D335N,V336A,V336L,V336M,Q338E,Q338P,Q338R,I339N,I339T,I339V,R341C,R341H,R341S,Q342H,E343K,S344G,L345P,L345V,A346T,A347T,A348E,A348T,S352I,P355T,K357E,K357N,K357T,T359N,T360N,E361K,E361Q,L364F,R366L,R366P,R366Q,R366W,A367E,A367V,T372N,L373V,R374Q,R374W,L375F,L375I,Y376C,Y376H,P377H,G379D,G379S,G379V,L382V,E383V,R384P,V386G,S387I,L390F,I397N,I397T,A399S,T401I,V403A,V403E,Q404H,Q404R,F406V,L407F,L407I,L407R,Y408F,Y408S,R412C,R412G,R412H,R412L,R412P,N413D,A414P,A415T,F417S,P418L,P418R,P420T,E421D,E421K,R422L,R422Q,R422W,N424D,P425L,R427C,R427H,D430A,I431S,I431T,R432S,R432W,G433D,G433S,S434F,S434P,S434Y,G435S,R436K,N437K,N437T,H439Y,V441M,P442L,F443C,F445S,G446S,M447I,M447T,R448G,R448H,L451H,G452A,G452E,G452R,R453Q,R453W,R454C,R454H,R454L,A458S,E459D,E459V,M460R,M460T,L461V,L464M,L464Q,H465Q,H465R,H466D,H466Q,V467M,H470Y,L472P,E474G,E474K,T477I,Q478L,E479G,D480E,D480N,I481L,K482N,M483V,V484A,Y485C,S486I,F487L,F487V,I488M,I488R,I488T,P491S,G492D,T493M,S494F,P495H,L496P,L497I,T498I,F499C,A501G,A501V,I502T,N503I,N503S,N503T. |
| Pathogenic | T498A,L461P,V386A,T318M,E198D,T185I,R181W,S150W. |
| **CYP17A1** | |
| Non-pathogenic | M1T,M1V,E3D,E3K,V5G,V5M,L10P,T11I,A13V,R21G,R21K,R21S,C22R,C22W,P23H,P23L,K26E,Y27S,P28S,K29Q,K29R,S30N,L31F,L36R,V37G,G38A,S39R,S39T,P41T,P44S,H46Y,G47S,M49T,H50N,F53L,F53V,F54L,K59I,K59N,Y60I,P62R,P62S,S65L,R67C,R67H,R67L,M68L,M68V,T70A,T73I,V74L,I75M,G77S,H78Q,H79D,H79N,H79Q,L81V,K83R,E84D,V85A,V85G,I87N,I87V,G90D,G90V,G95V,Q98L,M99I,A100T,L102P,D103A,D103H,A105T,A105V,N108I,R109C,I112M,A113G,A113T,A115T,G118D,A119S,A119T,H120P,H120Q,R125Q,A128E,A128V,M129T,M129V,A130T,A130V,D137N,G138S,G138V,D139H,D139N,I145T,E149G,T152I,T152R,D155N,M156T,T159A,T159I,N161H,N161K,G162R,G162V,Q163E,S164P,I165V,D166G,S168Y,F169S,V173L,V173M,A174V,V175G,V178D,V178I,F184L,N185K,T186S,S187T,D192E,D192N,P193T,N196K,V197D,V197I,I198L,Q199H,Y201D,Y201N,E203D,E203K,G204A,I206T,D207E,N208S,L209M,S210G,S210I,S210T,K211T,D212G,S213G,S213N,V215G,D216E,D216H,P219L,P219S,K222N,I223V,F224S,E230K,K231N,S234G,S234R,K237E,I238T,R239Q,D241Y,N244K,N244S,E248A,E248K,E252D,R255Q,R255W,I259L,I259T,M262I,M262L,M262R,L266M,S273A,D274Y,D283H,D283Y,E285K,L286R,L287F,L293F,T294S,I296M,I296T,I296V,G297A,D298G,I299T,V304M,T306A,T308I,K312R,L315P,A316D,A316S,A316V,F317S,L318M,H320Q,H320R,N321K,N321S,P322L,Q323R,K327Q,L328F,L328V,Y329C,Y329N,E330K,E331Q,I332T,D333N,S339G,R340C,R340H,T341A,T341I,I344V,R347S,R349C,R349H,A355T,R362H,L363F,R364K,V366M,A367V,M369T,M369V,I371T,P372A,P372H,P372L,H373N,K374T,A375V,N376K,N376S,V377I,D378A,S379F,S380G,S380R,I381V,G382A,G382S,D387G,D387N,K388E,K388N,T390R,A398V,H400Q,H401Q,N402D,E403D,K404M,H407P,P409L,P409R,P409S,F412L,M413V,P414L,E415D,R416C,R416H,L418V,N419K,P420A,P420L,P420S,A421V,G422E,P428S,P428T,Y432S,G436A,G436E,G436R,A437E,A437T,P439S,R440C,R440S,C442R,G444D,E445Q,A448D,A448G,A448V,R449C,R449H,E451A,F453S,L454F,I455F,I455L,I455T,I455V,M456I,M456V,A457T,F463C,D464N,E466 |

| | |
|---|---|
| | K,V467A,V467G,P468S,D470H,D470Y,G471E,Q472K,Q472S,P474S,G478D,G478S,G478V,P480T,V482M,V483A,I486M,D487N,D487Y,K490E,V495A,V495M,R496L,A498S,A498T,A498V,E501K,A502P,A504T,E505K,G506D,G506S,S507N,S507R,T508I. |
| Pathogenic | R496H,R496C,R440H,P428L,F417C,W406R,W406L,H373N,H373L,R362C,R358Q,R347C,R347H,P342T,Y329D,N177D,A174E,W121R,D116V,F114V,S106P,R96W,R96Q,F93C,Y64S,P35L. |
| **CYP19A1** | |
| Non-pathogenic | E4K,M5I,N7T,P8A,P8L,P8S,H10R,Y11C,N12H,N12K,I13L,I13M,I13V,T14I,T14N,T14P,S15G,V17M,A20T,M21I,M21T,M21V,P22S,P22T,A23P,A23S,T25A,T25N,T25S,P27S,V28L,L30V,L37S,V38M,W39R,E42K,G43V,T44A,S45F,I47M,G49D,Y52S,M54I,M54T,M54V,G57V,H62N,H62R,G63S,G71S,S72R,A73T,A73V,Y77C,N78S,R79G,R79P,R79Q,R79W,V80G,Y81C,F84L,M85I,M85R,M85V,R86Q,I89M,I89V,G91A,E92K,T94I,L95F,I96N,I96Y,I97F,I97M,S100Y,S102I,S102N,S102T,M107I,N110D,S113G,R115Q,G117S,K119N,K119Q,K119R,I125L,I125V,G126R,G126S,M127I,M127T,H128R,E129A,E129K,G131D,G131S,I132N,N136K,N137H,E139Q,L140F,L140V,R145Q,P146S,F147L,F148S,F148V,M149R,M149V,K150E,A151G,A151V,P155L,P155S,G156D,G156R,G156S,L157I,R159C,R159H,R159L,M160I,V161A,T162I,C164Y,A165T,K169R,T170I,R174G,R174M,E177K,E177Q,V178M,T179N,N180D,N180S,S182L,Y184F,Y184H,Y184S,D186H,V187M,T189A,R192C,R193C,R193H,M195I,M195L,M195T,D197H,T198P,T201K,T201M,L202R,F203L,L204W,R205K,P207S,P207T,L208F,L208W,D209G,D209N,S211I,S211N,A212D,A212T,I213N,I213V,V214M,K216N,G219R,Y220D,F221S,A223G,A223T,A223V,I229T,I233F,K236E,S238P,L240P,L240Q,Y241C,Y241F,Y241N,Y244H,E245G,E245K,S247P,V248A,D250V,D250Y,I255M,I259R,E261D,R263G,R263K,R264C,R264H,R264L,T268I,E274K,C275G,C275R,M276V,D277E,D277Y,T280A,T280I,T280N,E281A,E281Q,L282M,I283N,I283T,K287Q,R288C,R288G,R288H,G289V,D290N,R293T,N295H,V296L,Q298R,I300M,I300V,L301V,E302K,M303V,L304R,A306T,A307T,D309G,D309N,T310I,T310S,M311I,M311T,M311V,V313I,S314P,F316L,F317L,M318L,M318R,M318T,L321I,I322T,P326S,N327S,E330G,E330Q,I332M,I333M,I336N,T338A,V339I,G341C,G341R,D344H,I345K,I347T,D349Y,Q351P,K354E,V355M,M356T,M356V,N358D,N358S,F359L,F359Y,I360F,I360V,E362D,S363R,S363T,M364T,R365W,Y366S,Q367L,V370A,V370M,V373F,M374L,R375H,R375L,L378S,L378V,E379D,D381V,V382I,I383V,D384N,D384Y,G385C,G385V,Y386H,P387A,P387L,P387R,P387S,K389R,K390E,K390N,G391E,T392A,T392K,N393H,N393T,I394V,N397H,R403T,L404F,E405K,F406L,F406V,F407S,P408H,P410S,P410T,N411S,E412K,F418S,F418V,K420N,N421S,V422I,Y424F,Y424H,R425K,R425M,Y426C,Y426H,G431C,G433E,R435H,G436C,C437W,A438V,Y441C,I442T,I442V,A443S,A443T,M444V,V445G,L451P,V452I,V452S,T453I,R456I,R457L,R457Q,F458L,V460L,V460M,T462A,T462I,Q464H,G465R,C467Y,V468A,V468I,E469K,I471M,I471V,K473E,I474V,H475Q,D476A,D476H,D476N,S478F,L479W,H480P,P481L,D482E,T484I,K485R,M487V,E489K,F492V,T493N,T493P,P494S,R495G,R495T,N496K,D498E,D498N,R499G,R499T,R499W,C500Y,L501P,E502Q. |
| Pathogenic | R192H,R365Q,R375C,R435C,C437Y,E210K. |
| **CYP21A2** | |
| Non-pathogenic | L2Q,L3F,L3I,L6P,L7P,L9V,L9LL,P10S,P10T,L10V,P11L,P11S,P11T,L12P,L13P,A13P,A13S,A14P,A14S,G14S,G15D,A15D,G15S,A15V,R16C,A16D,R16H,R16L,A16S,A16T,A16V,R17C,R17H,R17L,W19C,W20C,N21S,W22C,W23C,L24F,L25F,R25Q,R25W,S26G,R26Q,R26W,S27G,S27R,H28P,H28Q,H28Y,L29F,L29H,H29N,H29P,H29Q,H29Y,L30F,L30H,P30S,P31L,P31S,P31T,P32H,P32R,P32S,P32T,A33V,P34L,A34T,A34V,P35L,P35R,G36S,F37I,L38F,H38L,H38P,L38S,H39D,H39L,H39P,L39V,L40P,L40V,L41P,Q42L,Q42P,D43G,D43N,L44F,D44G,D44N,P45A,L45F,P45L,P45R,P45S,P46A,P46L,I46L,P46R,P46S,Y47C,I47L,I47M,Y47S,Y48C,Y48S,L50P,G51D,G51S,T52I,Q53E,T53I,Q53K,Q54E,Q54K,G57R,P57S,P58S,I58V,I59T,I59V,Y60C,R60T,R61T,H62R,H62Y,H63L,H63R,H63Y,G65R, |

| | |
|---|---|
| | Q66K,Q67K,Q67L,D68G,K69E,V69L,K69N,K69R,L70V,V71M,S72T,S72Y,S73C,R73K,R73T,N74K,K74R,Y75C,Y75H,R75K,Y75N,P76L,P76Q,P76R,P76S,P76T,D77E,D77N,E78D,L78Q,L78R,L78V,S79F,S79P,A80V,G81E,M81T,V82D,S84F,S84Y,W85C,L85V,A86S,D87G,D87N,A89P,A89S,A89V,G90D,H90P,H90Q,H90Y,K91E,R91G,R91K,K91Q,K91R,P92H,K92Q,K92R,P92T,E93G,L93P,E93Q,T94I,T94S,R95C,R95H,L95P,T96A,T96I,S96L,Y97C,Y97N,A97S,A97V,K98E,K98N,K98R,G101A,G101D,G101S,S101T,S101Y,I102M,K102R,K102T,I102V,R103C,R103H,N103K,Y104C,Y104H,D104N,D104Y,S105F,P105Q,P105T,M106I,D106N,M106V,L107Q,E107V,L107V,S108P,P108T,V109A,V110A,G110E,E111A,E111G,E111K,E111Q,E111V,S113Y,F117I,C118S,A118V,H119Q,R120C,R120H,R120L,R120P,K120Q,K120R,R120S,M121I,M121L,K121R,M121V,R122G,R122K,R122T,T123I,R124C,R124CQ124HQ124H,Q124H,Q124P,S125L,P125S,G126D,G126S,T127I,V129M,G130A,G130D,G130S,A130T,I131M,I131V,R132C,E132D,E132G,E132V,E133G,D133N,D133Y,E134A,S134F,E134K,F135V,M135V,S136C,S136F,E136V,L137R,L137V,L138F,V139A,T139I,E140G,E140K,E140Q,C140S,E140V,C140Y,S141G,S141I,S141NI143N,I143N,Y145C,L146F,L146H,F146I,L146P,L146V,T147N,C147S,F148L,G149A,R149C,R149H,R149L,R149P,G149R,R149S,D150E,M150L,M150V,R151G,R151K,K151N,I152M,I152T,Q153H,Q153P,D154E,P154S,D155H,D155N,G155S,N156K,L157S,L157V,M158I,M158T,P159A,P159L,P159R,A159T,A160P,A160V,E161D,E161G,Y161H,Y161N,E163A,E163K,F164V,C164Y,S165F,I165T,L166R,L166V,E167K,V168L,V168M,L169F,L169S,S170G,S170I,T171A,I172N,S173G,S173I,S173N,S173R,H174Q,H174Y,L175F,L175H,L175P,W175S,T176N,F177L,I177V,Q178R,Q178RD179E,D179E,I179L,I179T,I179V,D181E,I181M,I181T,V182A,V182L,V182M,D183E,I183T,D184H,D184N,P184T,F185S,L186S,L186V,M187I,M187T,R187W,P188A,P188L,F188L,A189P,A189V,F189Y,Y190H,P190L,Y190N,N191D,N191H,N191S,Y191H,G193C,G193R,C193Y,L194F,L194P,R195G,R195Q,R195W,E196K,V197L,V197M,K198Q,L198S,Q199E,T200A,A200G,I201K,I201T,E202G,S202G,S202N,K203N,K203Q,H203Q,H203Y,R204M,D205E,D205N,H206N,I206V,H206Y,I207F,I207M,I207N,Q207R,I207S,I207V,V208E,I208L,V208L,V208M,E209D,E209K,E209Q,M210K,M210T,M210V,V211A,Q211E,V211M,L212M,I212T,L212V,R213S,F214S,H215R,H215Y,K216N,S218N,S218R,S218T,L219F,P219L,V220A,N220D,V220M,N220S,A221T,G222C,G222S,L223F,L223P,Q223R,W224C,R224Q,W224R,R224W,R225K,R225S,R225T,D226E,D226G,M227T,M227V,M228R,D229G,D229N,M231I,M231L,M231R,M231T,M231V,L232P,Q233E,D234E,G234E,D234N,G234V,V235A,I236M,I236MA236V,A236V,I236V,V237M,E238D,E238K,P238L,P238R,S239G,S239N,S239T,M239T,M239V,Q240E,M240I,M240V,E241K,L241V,R242S,G243V,S244C,H244R,S244Y,G245A,L247F,S247R,L247V,L248F,V249A,V249M,G250E,A250T,G251S,H251Y,V252L,V252M,W253C,W253R,H253Y,R254K,R254T,M254V,D255E,D255G,A255P,A255T,A256S,A256T,M256T,V257A,V257G,D258G,D258N,L259F,L259V,M260R,M260T,M260V,G262R,G262S,G263S,G263V,V264A,E265K,E265Q,A265V,T266A,P267L,P267R,S268G,S268N,S268T,T270A,T270I,T270S,L271F,L271P,S272F,G272V,S273Y,A274T,V275L,V275M,V276A,V276F,L276F,L276V,F277V,L278F,L278S,L278V,L279F,H280P,H280Q,H280Y,H281D,V281M,V281MH281Y,P282A,P282L,P282S,H282N,E283D,E283G,M283V,I284L,A284P,Q285H,A285S,A285T,V286A,V286G,D287G,R287G,R287Q,L288F,E290K,L292I,L292R,G292S,D293H,D293V,E294K,H294Q,H294Y,T295A,E295G,E295K,E295Q,L296Q,G297D,G297S,P298R,P298S,G299R,G299S,L300P,A300V,S302G,S302N,S303F,A303T,V304M,R304Q,R304W,V305A,V305F,P306S,F306V,Y307H,L307S,L307V,H309P,H309Y,R310C,H310D,R310G,R310H,H310N,R310S,P311A,P311L,P311S,R312G,E312G,R312P,R312Q,R312W,I313L,Q314H,L316F,R316Q,A318T,A318V,T319A,E319K,T319S,I320M,A321T,A321V,D322H,E322K,E324G,E324K,E324Q,R325C,R325H,G326S,R327Q,P327S,R327W,P328H,P328T,V329I,A329V,S331G,P331H,P331S,S332F,R333Q,R333W,V334F,P335H,P335L,P335R,P335T,Y336H,H336Y,R337C,R337G,R337H,R337S,R339C,R339G,R339S,R340L,R340P,R340Q,R340W,R341G,R341Q,P341R,P341S,S343N,S345Y,G346D,G346R,G346S,A347V,T348A,D348N,T348S,D348Y,I349M,I349N,I349S,I349T,P350R,A350T,A350V,E351D,E351K,G352S,T353I,V354D,V354L,I355V,P357L,P357Q,P357S,V358I,N358K,L359F,L359R,Q360H,P360S,A362P,A362T,A362TH363N,H363Q,R366C,R366G,R366H,R366S,T367A,T367M,V368I,V368L,R369P,R369Q,W369S,E370D,E370G,P370S,R |

| | |
|---|---|
| | 371G,R371K,S372N,H373R,H373Y,E374D,E374G,G375D,G375R,G375S,D377N,D377Y,I378N,I378S,I378T,R379C,R379G,R379H,P379R,R379S,L381M,G381S,T382I,P383A,V383D,V383L,P383S,G384D,G384R,K385N,P386L,P386Q,S387C,S387F,N387K,L388F,R388K,L388R,Q389H,L390P,A391G,A391P,A391T,A391V,F392C,F392L,H392Q,F392S,G393S,G395S,T396M,A396T,A396V,R397C,R397H,V397I,V397L,V398M,W398S,E399D,E399G,R400G,G401S,E402K,H402R,H402Y,E403D,P403L,P403R,L404P,L404V,A405V,R406C,R406H,R408G,R408H,R408S,L409F,L409R,V411L,V411M,V412L,V412M,P412S,G413D,G413R,T414I,T414N,K414N,T414S,R415P,R415Q,S416C,S416F,R417K,L419P,A419T,A419V,A420G,F421L,T421M,G422S,P424HP424SP424S,P424S,A425T,G426R,D427E,V427M,A428D,A428T,G430S,P430S,S431F,E431K,P432L,L432P,P432R,L433P,L433V,P434L,P434R,A434V,R435H,P436S,L438F,L438R,C438Y,S439G,S439T,V440I,V440L,V440M,V441L,V441M,L442F,L442V,T443I,T443N,T443S,M444I,R444P,R444Q,M444R,M444V,Q445R,P446L,P446S,P446T,F447L,F447S,F447V,A448T,A448V,R450G,R450L,T450M,R450Q,R450W,L451R,Q452E,P453A,P453T,R454P,R454Q,R454W,G455E,G455R,D456E,M456I,M456L,A457D,G457R,G457W,A458D,A458P,A458T,A458V,H459Q,H459R,P459S,S460F,S460G,S460I,S460T,P461L,L461P,G462D,G462S,G462V,Q463H,P463L,P463R,Q463R,S464N,S464R,S464T,Q465H,P465S,S468G,V469I,L471F,L471V,M473I,M473R,P475T,F476L,F476S,R479G,R479Q,R479W,P482A,P482T,G484E,M485I,M485L,G486R,A487T,H488Q,S489G,S489T,P490L,G491D,G491S,G491V,Q492H,N493K,N493S,N493T,Q494H. |
| Pathogenic | R483W,R483Q,R483P,P482S,R479L,P453S,T450P,R435C,R426H,R426C,G424S,R408C,E380D,R369W,H365Y,L363W,A362V,R356W,R356Q,R356P,R354H,R354C,R341W,R341P,R339H,E320K,L317M,W302R,S301Y,L300F,G292D,G291S,G291R,G291C,M283L,H282N,V281L,V281G,L261P,M239K,V237E,I236N,R233K,I230T,V211L,L198F,Y191H,G178A,G178R,I172N,C169Y,C169R,L167P,L142P,R124H,K121Q,S113F,L107R,P105L,G90V,I77T,G64E,H62L,G56R,P30Q,P30L,A15T,L12M. |
| **CYP46A1** | |
| Non-pathogenic | L5M,L8F,G9C,A11T,V12G,L13P,L13V,A15S,A15T,F16L,A26S,A26T,R29G,R29H,E31G,P37Q,P39H,F41Y,V55A,R58C,R58G,R58H,R58P,F64L,K70N,P73S,V75A,R76Q,R76W,N78S,V79I,H81D,H81Q,I86V,V87I,S89I,P90L,E91G,V93F,V93I,M98T,N103S,D105N,R110C,R110G,R110H,R110S,A111V,T114S,E128K,E132D,W134C,H135Y,R138Q,V140G,I141M,D142E,D142N,D142Y,F145S,R147Q,R147W,S148N,L150S,L153F,M154L,E159K,K160E,V165M,G174E,Q175K,T176N,P177T,S179Y,Q181E,Q181L,D182N,T185A,T185I,T185P,T187A,T187P,A188T,M189V,I191M,A196T,M199I,T201N,T201S,M203V,G206C,G206S,A207T,A207V,Q208H,P210A,P210L,P210S,S212F,Q213L,V215G,K216N,M218I,M218T,T223P,A224V,R226C,R226H,T228P,T228S,A230T,F232L,G235A,K238M,R241Q,R241W,E242K,V243I,R244Q,E245K,R248C,R248H,R251C,Q252R,V253L,V258I,Q259R,R260C,R260H,R260P,R262Q,A264D,A264S,R267M,G268D,G268S,E269K,V271D,V271F,V271I,P272S,A273V,D274N,I275V,T277P,G285E,A286V,Q287H,Q287K,D289N,G291V,V297I,T298I,F299S,F300Y,H304Q,E305K,T306P,A312V,T314A,R320H,E323D,I324F,V325M,A326T,Q329L,E331G,V332G,V332L,I336V,G337D,F344S,E345G,D346N,Y352C,S360L,L361V,G369D,T370N,R372C,R372H,E377D,E377K,V383I,G387S,P390L,P390S,G399V,D402H,Y404H,E406G,D407E,D407N,T410N,D414N,R415H,G417R,G417S,P418T,A420E,A420S,A420T,P421H,K422R,P423L,R424Q,R424W,F428L,R435G,S436F,C437G,G439R,Q444E,Q444H,M445V,K448Q,A452T,A452V,R457G,R457K,R457S,L458V,E459V,R461Q,R461W,V463A,V463G,V463M,Q466K,R467C,R467H,R467S,G469R,Q473K,T475I,L479M,D480N,P481A,P481L,P481S,V482L,V482M,T485N,L486V,R487Q,R487W,P488L,R489C,G490D,G490R,G490S,W491C,A494T,P495A,P495T,P496L,P496T,P497S,P498L,P498R,P498S,P499A,P499S,C500Y. |
| **CYP51A1** | |

| | |
|---|---|
| Non-pathogenic | A5P,A5T,L8V,L9P,Q14L,A15V,G16R,G16V,G17A,G17E,S18A,S18W,V19A,A23T,M24I,M24L,K26N,K26R,V27E,N31K,L32H,L33S,M35I,M35L,L36R,L37P,L37V,I38T,A39S,C40G,C40S,A41T,T43P,S45T,L46Q,R51C,R51H,R51S,L52V,A54T,A54V,H56Q,H56Y,L60M,P61L,A62S,G63E,G63W,V64M,K65R,S66G,P68T,I70V,F71V,S72F,I74F,I74L,I74V,F76L,F76Y,L77F,G78V,H79Y,F83L,G84E,I88T,I88V,E92G,E96G,Y98C,S103G,F104C,T105N,T105S,M106V,V107A,V107I,T110A,T112S,L114P,L115R,D118A,A119G,A120V,A121T,A121V,N125H,S126N,D130E,D130H,A133V,D135G,Y137C,S138N,R139C,R139H,R139L,P143S,V144M,Y151D,D152E,D152G,D152N,P154L,P154T,N155S,P156L,V157A,V157G,V157I,F158V,M164L,S167G,L169V,I171V,A172V,H177Q,H177R,V178I,S179P,I180T,I180V,T185A,Y188C,E190D,E190G,E197Q,K198R,N199D,N199S,V200L,V200M,A203T,A203V,I208L,I208V,I209V,T211I,A212G,A212S,A212V,H214R,L216F,H217L,K219R,I221V,L225V,N226S,E227G,K228Q,V229L,Y233C,D237G,G238E,G239D,S241C,S241T,H242R,A244G,L247S,P248S,G249D,G249S,P252A,P252R,L253S,L253W,S255G,R258C,R258H,R258L,R261K,H263R,H263Y,R264G,R264Q,R264W,E265G,I266S,D268G,I269V,F270L,Y271C,K272E,A273P,I274T,Q275R,R277C,R277H,R277L,Q281P,I284V,D285G,D285Y,I287V,T290A,D293V,A294T,R300C,R300H,D304V,D305Y,A308V,L311F,L311R,I312F,L315F,A317E,A317V,H320P,H320R,T321I,T324A,T324I,T325A,S326I,S326R,M329V,G330A,L333S,A334S,D336E,K337R,T338I,Q340K,K341E,Y344H,E346D,Q347K,P356L,P356S,T359A,T359S,D361E,Q362E,D365N,D365V,D370E,R371C,R371G,R371H,R371L,R371S,I373V,L379I,P381H,P381T,I383L,I383V,M384L,I385F,M386I,M386V,M387I,M387V,V396M,Y399D,T400A,P403L,P403S,H405Y,Q406R,V409I,S410F,P411S,L417P,L417R,D419Y,S420L,E423D,R424C,R424H,R424P,D426N,D430H,R431C,R431H,R431L,Y432D,Q434E,Q434H,Q434R,D435E,P437S,E441K,A444V,Y445C,A450V,R452C,R452H,R452S,H453N,R454C,R454H,I456T,E458K,Y462C,Y462D,Y462H,Y462N,V463I,Q464H,K466E,K466R,T467I,T467K,I468V,W469C,S470F,M472V,R474H,R474P,L480F,I481F,I481T,D482V,G483R,F485L,V488L,T492A,T492I,M493V,T496I,P497H,N499Y,P500R,P500S,I502M,R503C,R503H,R506Q,R507K,S508T. |

**Table S3.** Summary of benchmarked methods.

| **Method** | **Description** | **Input** | **Threshold** |
|---|---|---|---|
| SNPMUSIC | Stability-driven, knowledge-based classifier that uses statistical potential | 3D structure | G < 0 neutral, G> 0 deleterious |
| mCSM-Stability | Graph-based structural signatures | 3D structure | G > 0 stabilizing, G < 0 destabilizing |
| POPMUSIC | Linear combination of statistical potential | 3D structure | G < 0 stabilizing, G > 0 destabilizing |
| I-Mutant3.0 | Support vector machine-based protein stability predictor | 3D structure | G > 0 stabilizing, G < 0 destabilizing |
| SIMBA | A straightforward multilinear regression model that was trained using an experimental dataset balanced for substitution types | 3D structure | G > 0 stabilizing, G <0 destabilizing |

| | | | |
|---|---|---|---|
| SNAP2 | A classifier based on neural networks, trained using experimentally derived variant data to predict functional effects | Sequence | Score > 50 strong signal for effect, score <-50 strong signal for neutral/no effect. |
| METASNP | A meta predictor employing a random forest approach that integrates SNAP, SIFT, PANTHER, and PHD-SNP predictions | Sequence | Score < 0.5 neutral, score > 0.5 disease-causing |
| SIFT | Employing sequence similarity (utilizing PSI-BLAST) in conjunction with established probabilities of amino acid substitutions | Sequence | Score > 0.5 tolerated, score < 0.5 deleterious |
| FATHMM | A classifier based on Hidden Markov Models (HMMs) | Sequence | Score > 0 neutral, score < 0 pathogenic |
| Envision | High-throughput laboratory methods like deep mutational scanning has generated extensive quantitative datasets on mutational effects for diverse proteins across various organisms | Sequence | Score = 0 most disruptive mutations, score ~ 1 wild-type-like mutation |

**Table S4. 48 properties of the twenty standard amino acids.**

| **Property** | **A** | **D** | **C** | **E** | **F** | **G** | **H** | **I** | **K** | **L** |
|---|---|---|---|---|---|---|---|---|---|---|
| **K** | -25.5 | -33.12 | -32.82 | -36.17 | -34.54 | -27 | -31.84 | -31.78 | -32.4 | -31.78 |
| **Ht** | 0.87 | 0.66 | 1.52 | 0.67 | 2.87 | 0.1 | 0.87 | 3.15 | 1.64 | 2.17 |
| **Hs** | 13.05 | 11.1 | 14.3 | 11.41 | 13.89 | 12.2 | 12.42 | 15.34 | 11.01 | 14.19 |
| **P** | 0 | 49.7 | 1.48 | 49.9 | 0.35 | 0 | 51.6 | 0.1 | 49.5 | 0.13 |
| **pHi** | 6 | 2.77 | 5.05 | 5.22 | 5.48 | 5.97 | 7.59 | 6.02 | 9.74 | 5.98 |
| **pK** | 2.34 | 2.01 | 1.65 | 2.19 | 1.89 | 2.34 | 1.82 | 1.36 | 2.18 | 2.36 |
| **Mw** | 89 | 133 | 121 | 147 | 165 | 75 | 155 | 131 | 146 | 131 |
| **B1** | 11.5 | 11.68 | 13.46 | 13.57 | 19.8 | 3.4 | 13.67 | 21.4 | 15.71 | 21.4 |
| **Rf** | 9.9 | 2.8 | 2.8 | 3.2 | 18.8 | 5.6 | 8.2 | 17.1 | 3.5 | 17.6 |
| **u** | 14.34 | 12 | 35.77 | 17.26 | 29.4 | 0 | 21.81 | 19.06 | 21.29 | 18.78 |
| **Hnc** | 0.62 | 0.9 | 0.29 | -0.74 | 1.19 | 0.48 | -0.4 | 1.38 | -1.5 | 1.06 |
| **Esm** | 1.4 | 1.16 | 1.37 | 1.16 | 1.14 | 1.36 | 1.22 | 1.19 | 1.07 | 1.32 |
| **El** | 0.49 | 0.35 | 0.67 | 0.37 | 0.72 | 0.53 | 0.54 | 0.76 | 0.3 | 0.65 |
| **Et** | 1.9 | 1.52 | 2.04 | 1.54 | 1.86 | 1.9 | 1.76 | 1.95 | 1.37 | 1.97 |
| **Pa** | 1.42 | 1.01 | 0.7 | 1.51 | 1.13 | 0.57 | 1 | 1.08 | 1.16 | 1.21 |

| | | | | | | | | | | |
|---|---|---|---|---|---|---|---|---|---|---|
| **Pb** | 0.83 | 0.54 | 1.19 | 0.37 | 1.38 | 0.75 | 0.87 | 1.6 | 0.74 | 1.3 |
| **Pt** | 0.66 | 1.46 | 1.19 | 0.74 | 0.6 | 1.56 | 0.95 | 0.47 | 1.01 | 0.59 |
| **Pc** | 0.71 | 1.21 | 1.19 | 0.84 | 0.71 | 1.52 | 1.07 | 0.66 | 0.99 | 0.69 |
| **Ca** | 20 | 26 | 25 | 33 | 46 | 13 | 37 | 39 | 46 | 35 |
| **F** | 0.96 | 1.14 | 0.87 | 1.07 | 0.69 | 1.16 | 0.8 | 0.76 | 1.14 | 0.79 |
| **Br** | 0.38 | 0.14 | 0.57 | 0.09 | 0.51 | 0.38 | 0.31 | 0.56 | 0.04 | 0.5 |
| **Ra** | 3.7 | 2.6 | 3.03 | 3.3 | 6.6 | 3.13 | 3.57 | 7.69 | 1.79 | 5.88 |
| **Ns** | 6.05 | 4.95 | 7.86 | 5.1 | 6.62 | 6.16 | 5.8 | 7.51 | 4.88 | 7.37 |
| **an** | 1.59 | 0.53 | 0.33 | 1.45 | 1.14 | 0.53 | 0.89 | 1.22 | 1.13 | 1.91 |
| **ac** | 1.44 | 2.13 | 0.76 | 2.01 | 1.01 | 0.62 | 0.56 | 0.68 | 0.59 | 0.58 |
| **am** | 1.22 | 0.56 | 1.53 | 1.28 | 1.13 | 0.4 | 2.23 | 0.77 | 1.65 | 1.05 |
| **V0** | 60.46 | 73.83 | 67.7 | 85.88 | 121.48 | 43.25 | 98.79 | 107.72 | 108.5 | 107.75 |
| **Nm** | 2.11 | 1.8 | 1.88 | 2.09 | 1.98 | 1.53 | 1.98 | 1.77 | 1.96 | 2.19 |
| **Nl** | 3.92 | 2.85 | 5.55 | 2.72 | 4.53 | 4.31 | 3.77 | 5.58 | 2.79 | 4.59 |
| **Hgm** | 13.85 | 11.61 | 15.37 | 11.38 | 13.93 | 13.34 | 13.82 | 15.28 | 11.58 | 14.13 |
| **ASAD** | 104 | 132.2 | 132.5 | 161.9 | 182 | 73.4 | 165.8 | 171.5 | 195.2 | 161.4 |
| **ASAN** | 33.2 | 62.4 | 17.9 | 81 | 33.1 | 29.2 | 57.7 | 28.3 | 107.5 | 31.1 |
| **ΔASA** | 70.9 | 69.6 | 114.3 | 80.5 | 148.4 | 44 | 107.9 | 142.7 | 87.5 | 129.8 |
| **ΔGh** | -0.54 | -2.97 | -1.64 | -3.71 | -1.06 | -0.59 | -3.38 | 0.32 | -2.19 | 0.27 |
| **GhD** | -0.58 | -6.1 | -1.91 | 7.37 | -1.35 | -0.82 | -5.57 | 0.4 | -5.97 | 0.35 |
| **GhN** | -0.06 | -3.11 | -0.27 | -3.62 | -0.28 | -0.23 | -2.18 | 0.07 | -1.7 | 0.07 |
| **ΔHh** | -2.24 | -4.54 | -3.43 | -5.63 | -5.11 | -1.46 | -6.83 | -3.84 | -5.02 | -3.52 |
| **TΔSh** | 1.7 | 1.57 | 1.79 | 1.92 | 4.05 | 0.87 | 3.45 | 4.16 | 2.83 | 3.79 |
| **ΔCh** | 14.22 | 2.73 | 9.41 | 3.17 | 39.06 | 4.88 | 20.05 | 41.98 | 17.68 | 38.26 |
| **ΔGc** | 0.51 | 2.89 | 2.71 | 3.58 | 3.22 | 0.68 | 3.95 | -0.4 | 1.87 | -0.35 |
| **ΔHc** | 2.77 | 4.72 | 8.64 | 5.69 | 11.93 | 1.23 | 7.64 | 4.03 | 3.57 | 3.69 |
| **ΔSc** | -2.25 | -1.83 | -5.92 | -2.11 | -8.71 | -0.55 | -3.69 | -4.42 | -1.7 | -4.04 |
| **ΔG** | -0.02 | -0.08 | 1.08 | -0.13 | 2.16 | 0.09 | 0.56 | -0.08 | -0.32 | -0.08 |
| **ΔH** | 0.51 | 0.18 | 5.21 | 0.05 | 6.82 | -0.23 | 0.79 | 0.19 | -1.45 | 0.17 |
| **TΔS** | -0.54 | -0.26 | -4.14 | -0.19 | -4.66 | 0.31 | -0.23 | -0.27 | 1.13 | -0.24 |
| **v** | 1 | 4 | 2 | 5 | 7 | 0 | 6 | 4 | 5 | 4 |
| **s** | 0 | 2 | 0 | 3 | 2 | 0 | 2 | 1 | 0 | 2 |
| **f** | 0 | 2 | 1 | 3 | 2 | 0 | 2 | 2 | 4 | 2 |

| **Property** | **M** | **N** | **P** | **Q** | **R** | **S** | **T** | **V** | **W** | **Y** |
|---|---|---|---|---|---|---|---|---|---|---|
| **K** | -31.18 | -30.9 | -23.25 | -32.6 | -26.62 | -29.88 | -31.23 | -30.62 | -30.24 | -35.01 |
| **Ht** | 1.67 | 0.09 | 2.77 | 0 | 0.85 | 0.07 | 0.07 | 1.87 | 3.77 | 2.67 |
| **Hs** | 13.62 | 11.72 | 11.06 | 11.78 | 12.4 | 11.68 | 12.12 | 14.73 | 13.96 | 13.57 |
| **P** | 1.43 | 3.38 | 1.58 | 3.53 | 52 | 1.67 | 1.66 | 0.13 | 2.1 | 1.61 |
| **pHi** | 5.74 | 5.41 | 6.3 | 5.65 | 10.76 | 5.68 | 5.66 | 5.96 | 5.89 | 5.66 |
| **pK** | 2.28 | 2.02 | 1.99 | 2.17 | 1.81 | 2.21 | 2.1 | 2.32 | 2.38 | 2.2 |
| **Mw** | 149 | 132 | 115 | 146 | 174 | 105 | 119 | 117 | 204 | 181 |
| **B1** | 16.25 | 12.82 | 17.43 | 14.45 | 14.28 | 9.47 | 15.77 | 21.57 | 21.61 | 18.03 |
| **Rf** | 14.7 | 5.4 | 14.8 | 9 | 4.6 | 6.9 | 9.5 | 14.3 | 17 | 15 |
| **u** | 21.64 | 13.28 | 10.93 | 17.56 | 26.66 | 6.35 | 11.01 | 13.92 | 42.53 | 31.55 |

| | | | | | | | | | | |
|---|---|---|---|---|---|---|---|---|---|---|
| **Hnc** | 0.64 | -0.78 | 0.12 | -0.85 | -2.53 | -0.18 | -0.05 | 1.08 | 0.81 | 0.26 |
| **Esm** | 1.3 | 1.18 | 1.24 | 1.12 | 0.92 | 1.3 | 1.25 | 1.25 | 1.03 | 1.03 |
| **El** | 0.65 | 0.38 | 0.46 | 0.4 | 0.55 | 0.45 | 0.52 | 0.73 | 0.83 | 0.65 |
| **Et** | 1.96 | 1.56 | 1.7 | 1.52 | 1.48 | 1.75 | 1.77 | 1.98 | 1.87 | 1.69 |
| **Pa** | 1.45 | 0.67 | 0.57 | 1.11 | 0.98 | 0.77 | 0.83 | 1.06 | 1.08 | 0.69 |
| **Pb** | 1.05 | 0.89 | 0.55 | 1.1 | 0.93 | 0.75 | 1.19 | 1.7 | 1.37 | 1.47 |
| **Pt** | 0.6 | 1.56 | 1.52 | 0.98 | 0.95 | 1.43 | 0.96 | 0.5 | 0.96 | 1.14 |
| **Pc** | 0.59 | 1.37 | 1.61 | 0.87 | 1.07 | 1.34 | 1.08 | 0.63 | 0.76 | 1.07 |
| **Ca** | 43 | 28 | 22 | 36 | 55 | 20 | 28 | 33 | 61 | 46 |
| **F** | 0.78 | 1.04 | 1.16 | 1.07 | 1.05 | 1.13 | 0.96 | 0.79 | 0.77 | 1.01 |
| **Br** | 0.42 | 0.15 | 0.18 | 0.11 | 0.07 | 0.23 | 0.23 | 0.48 | 0.4 | 0.26 |
| **Ra** | 5.21 | 2.12 | 2.12 | 2.7 | 2.53 | 2.43 | 2.6 | 7.14 | 6.25 | 3.03 |
| **Ns** | 6.39 | 5.04 | 5.65 | 5.45 | 5.7 | 5.53 | 5.81 | 7.62 | 6.98 | 6.73 |
| **an** | 1.25 | 0.53 | 0 | 0.98 | 0.67 | 0.7 | 0.75 | 1.42 | 1.33 | 0.58 |
| **ac** | 0.73 | 0.93 | 2.19 | 1.2 | 0.39 | 0.81 | 1.25 | 0.63 | 1.4 | 0.72 |
| **am** | 1.47 | 0.93 | 0 | 1.63 | 1.59 | 0.87 | 0.46 | 1.2 | 0.46 | 0.52 |
| **V0** | 105.35 | 78.01 | 82.83 | 93.9 | 127.34 | 60.62 | 76.83 | 90.78 | 143.91 | 123.6 |
| **Nm** | 2.27 | 1.84 | 1.32 | 2.03 | 1.94 | 1.57 | 1.57 | 1.63 | 1.9 | 1.67 |
| **Nl** | 4.14 | 3.64 | 3.57 | 3.06 | 3.78 | 3.75 | 4.09 | 5.43 | 4.83 | 4.93 |
| **Hgm** | 13.86 | 13.02 | 12.35 | 12.61 | 13.1 | 13.39 | 12.7 | 14.56 | 15.48 | 13.88 |
| **ASAD** | 189.8 | 134.9 | 135.1 | 164.9 | 210.2 | 111.4 | 130.4 | 143.9 | 208.8 | 196.4 |
| **ASAN** | 41.3 | 60.5 | 60.7 | 71.5 | 94.5 | 48.7 | 52 | 28.1 | 39.5 | 50.4 |
| **ΔASA** | 147.9 | 74 | 73.5 | 93.3 | 116 | 62.8 | 78 | 115.6 | 167.8 | 145.9 |
| **ΔGh** | -0.6 | -3.55 | 0.32 | -3.92 | -5.96 | -3.82 | -1.97 | 0.13 | -3.8 | -5.64 |
| **GhD** | -0.71 | -6.63 | 0.56 | -7.12 | -12.78 | -6.18 | -3.66 | 0.18 | -4.71 | -8.45 |
| **GhN** | -0.1 | -3.03 | 0.23 | -3.15 | -6.85 | -2.36 | -1.69 | 0.04 | -0.88 | -2.82 |
| **ΔHh** | -4.16 | -5.68 | -1.95 | -6.23 | -10.43 | -5.94 | -4.39 | -3.15 | -8.99 | -10.67 |
| **TΔSh** | 3.56 | 2.13 | 2.27 | 2.31 | 4.47 | 2.12 | 2.42 | 3.28 | 5.19 | 5.03 |
| **ΔCh** | 31.67 | 3.91 | 23.69 | 3.74 | 16.66 | 6.14 | 16.11 | 32.58 | 37.69 | 30.54 |
| **ΔGc** | 1.13 | 3.26 | -0.39 | 3.69 | 5.25 | 3.42 | 1.74 | -0.19 | 5.59 | 6.56 |
| **ΔHc** | 7.06 | 3.64 | 1.97 | 4.47 | 6.03 | 5.8 | 4.42 | 3.45 | 13.46 | 14.41 |
| **ΔSc** | -5.93 | -0.39 | -2.36 | -0.78 | -0.78 | -2.38 | -2.68 | -3.64 | -7.87 | -7.95 |
| **ΔG** | 0.53 | -0.3 | -0.06 | -0.23 | -0.71 | -0.4 | -0.24 | -0.06 | 1.78 | 0.91 |
| **ΔH** | 2.89 | -2.03 | 0.02 | -1.76 | -4.4 | -0.16 | 0.04 | 0.3 | 4.47 | 3.73 |
| **TΔS** | -2.36 | 1.74 | -0.08 | 1.53 | 3.69 | -0.24 | -0.28 | -0.36 | -2.69 | -2.82 |
| **v** | 4 | 4 | 3 | 5 | 7 | 2 | 3 | 3 | 10 | 8 |
| **s** | 0 | 2 | 0 | 3 | 5 | 0 | 1 | 1 | 2 | 2 |
| **f** | 3 | 2 | 0 | 3 | 5 | 1 | 1 | 1 | 2 | 2 |

**Table S5.** Combined ANOVA results (A: 'all mutations', N: 'Non-pathogenic', P: 'Pathogenic', DIFF: Difference between means, LWR: Lower end of 95% confidence interval, UPR: Upper end of 95% confidence interval) for structure-based methods.

| Category | Method | Group Pair | DIFF | LWR | UPR | P-value |
|---|---|---|---|---|---|---|
| 26 CYP450 | SNPMUSIC | N-P | 0.27 | 0.26 | 0.28 | 0 |

| | | | | | | |
|---|---|---|---|---|---|---|
| | | A-P | 0.12 | 0.12 | 0.12 | 0 |
| | | A-N | -0.15 | -0.15 | -0.15 | 0 |
| | mCSM | N-P | 0.26 | 0.25 | 0.28 | 0 |
| | | A-P | 0.17 | 0.17 | 0.17 | 0 |
| | | A-N | -0.1 | -0.1 | -0.09 | 0 |
| | POPMUSIC | N-P | 0.26 | 0.25 | 0.28 | 0 |
| | | A-P | 0.1 | 0.1 | 0.1 | 0.35 |
| | | A-N | -0.16 | -0.16 | -0.15 | 0 |
| | I-MUTANT3.0 | N-P | 0.17 | 0.16 | 0.18 | 0 |
| | | A-P | 0.1 | 0.1 | 0.1 | 0.13 |
| | | A-N | -0.07 | -0.07 | -0.07 | 0 |
| | SIMBA | N-P | 0.03 | 0.01 | 0.05 | 1 |
| | | A-P | -0.11 | -0.12 | -0.1 | 0.72 |
| | | A-N | -0.14 | -0.14 | -0.13 | 0 |
| 10 Pathogenic mutation causing CYP450 | SNPMUSIC | N-P | 0.28 | 0.27 | 0.29 | 0 |
| | | A-P | 0.13 | 0.13 | 0.13 | 0 |
| | | A-N | -0.15 | -0.16 | -0.16 | 0 |
| | mCSM | N-P | 0.26 | 0.23 | 0.28 | 0 |
| | | A-P | 0.15 | 0.15 | 0.16 | 0.05 |
| | | A-N | -0.1 | -0.11 | -0.1 | 0 |
| | POPMUSIC | N-P | 0.26 | 0.24 | 0.29 | 0 |
| | | A-P | 0.12 | 0.11 | 0.12 | 0.25 |
| | | A-N | -0.15 | -0.15 | -0.14 | 0 |
| | I-MUTANT3.0 | N-P | 0.16 | 0.14 | 0.17 | 0 |
| | | A-P | 0.08 | 0.08 | 0.09 | 0.3 |
| | | A-N | -0.08 | -0.08 | -0.07 | 0 |
| | SIMBA | N-P | 0 | -0.03 | 0.03 | 1 |
| | | A-P | -0.12 | -0.13 | -0.12 | 0.55 |
| | | A-N | -0.13 | -0.14 | -0.12 | 0 |
| Dominant mutation containing 4 CYP450 | SNPMUSIC | N-P | 0.31 | 0.29 | 0.33 | 0 |
| | | A-P | 0.14 | 0.13 | 0.14 | 0 |
| | | A-N | -0.17 | -0.17 | -0.16 | 0 |
| | mCSM | N-P | 0.27 | 0.23 | 0.3 | 0 |
| | | A-P | 0.17 | 0.16 | 0.18 | 0.04 |
| | | A-N | -0.1 | -0.11 | -0.09 | 0 |
| | POPMUSIC | N-P | 0.28 | 0.24 | 0.31 | 0 |
| | | A-P | 0.14 | 0.14 | 0.13 | 0.12 |
| | | A-N | -0.14 | -0.15 | -0.13 | 0 |
| | I-MUTANT3.0 | N-P | 0.15 | 0.12 | 0.17 | 0 |
| | | A-P | 0.08 | 0.07 | 0.09 | 0.37 |
| | | A-N | -0.06 | -0.07 | -0.06 | 0 |
| | SIMBA | N-P | 0.05 | 0 | 0.09 | 1 |
| | | A-P | -0.06 | -0.07 | -0.05 | 1 |
| | | A-N | -0.11 | -0.12 | -0.1 | 0 |

**Table S6.** SNPMUSIC ANOVA results (A: 'all mutations', N: 'Non-pathogenic', P: 'Pathogenic', DIFF: Difference between means, LWR: Lower end of 95% confidence interval, UPR: Upper end of 95% confidence interval).

| Name | Group Pair | DIFF | LWR | UPR | P-value |
|---|---|---|---|---|---|
| CYP1A1 | A-N | -0.09 | -0.1 | -0.08 | **0** |
| CYP1A2 | A-N | -0.14 | -0.15 | -0.13 | **0** |
| CYP1B1 | A-N | -0.14 | -0.15 | -0.13 | **0** |
| CYP1B1 | A-P | 0.05 | 0.04 | 0.05 | 1 |
| CYP1B1 | N-P | 0.19 | 0.16 | 0.22 | **0.02** |
| CYP2A6 | A-N | -0.19 | -0.2 | -0.18 | **0** |
| CYP2A13 | A-N | -0.16 | -0.17 | -0.16 | **0** |
| CYP2B6 | A-N | -0.17 | -0.18 | -0.16 | **0** |
| CYP2C8 | A-N | -0.17 | -0.18 | -0.16 | **0** |
| CYP2C9 | A-N | -0.13 | -0.13 | -0.12 | **0** |
| CYP2C19 | A-N | -0.14 | -0.15 | -0.13 | **0** |
| CYP2D6 | A-N | -0.14 | -0.15 | -0.13 | **0** |
| CYP2E1 | A-N | -0.18 | -0.19 | -0.18 | **0** |
| CYP2R1 | A-N | -0.14 | -0.15 | -0.13 | **0** |
| CYP2R1 | A-P | -0.17 | -0.18 | -0.16 | 0.82 |
| CYP2R1 | N-P | -0.03 | -0.07 | 0.01 | 1 |
| CYP3A4 | A-N | -0.16 | -0.16 | -0.15 | **0** |
| CYP3A4 | A-P | -0.14 | -0.15 | -0.13 | 1 |
| CYP3A4 | N-P | -0.02 | -0.02 | 0.06 | 1 |
| CYP3A5 | A-N | -0.15 | -0.15 | -0.14 | **0** |
| CYP3A7 | A-N | -0.19 | -0.19 | -0.18 | **0** |
| CYP7A1 | A-N | -0.13 | -0.13 | -0.12 | **0** |
| CYP8A1 | A-N | -0.16 | -0.17 | -0.16 | **0** |
| CYP8A1 | A-P | -0.31 | -0.32 | -0.3 | 1 |
| CYP8A1 | N-P | -0.15 | -0.18 | -0.11 | 1 |
| CYP8B1 | A-N | -0.07 | -0.08 | -0.06 | 0.73 |
| CYP11A1 | A-N | -0.1 | -0.11 | -0.09 | **0** |
| CYP11A1 | A-P | 0.17 | 0.17 | 0.18 | 0.78 |
| CYP11A1 | N-P | 0.27 | 0.23 | 0.31 | 0.16 |
| CYP11B1 | A-N | -0.2 | -0.21 | -0.2 | **0** |
| CYP11B1 | A-P | 0.11 | 0.1 | 0.12 | 0.17 |
| CYP11B1 | N-P | 0.31 | 0.28 | 0.35 | **0** |
| CYP11B2 | A-N | -0.16 | -0.17 | -0.15 | **0** |
| CYP11B2 | A-P | 0.14 | 0.13 | 0.15 | 0.9 |
| CYP11B2 | N-P | 0.3 | 0.26 | 0.33 | 0.1 |
| CYP17A1 | A-N | -0.19 | -0.2 | -0.18 | **0** |
| CYP17A1 | A-P | 0.37 | 0.36 | 0.38 | **0** |
| CYP17A1 | N-P | 0.55 | 0.51 | 0.6 | **0** |
| CYP19A1 | A-N | -0.16 | -0.17 | -0.16 | **0** |
| CYP19A1 | A-P | 0.18 | 0.17 | 0.18 | 0.81 |
| CYP19A1 | N-P | 0.34 | 0.31 | 0.38 | 0.07 |
| CYP21A2 | A-N | -0.13 | -0.14 | -0.12 | **0** |

| | | | | | |
|---|---|---|---|---|---|
| CYP21A2 | A-P | 0.11 | 0.1 | 0.12 | 0.06 |
| CYP21A2 | N-P | 0.24 | 0.21 | 0.28 | **0** |
| CYP46A1 | A-N | -0.22 | -0.23 | -0.21 | **0** |
| CYP51A1 | A-N | -0.18 | -0.18 | -0.17 | **0** |

**Table S7.** mCSM ANOVA results (A: 'all mutations', N: 'Non-pathogenic', P: 'Pathogenic', DIFF: Difference between means, LWR: Lower end of 95% confidence interval, UPR: Upper end of 95% confidence interval).

| Name | Group Pair | DIFF | LWR | UPR | P-value |
|---|---|---|---|---|---|
| CYP1A1 | A - N | -0.05 | -0.07 | -0.03 | 0.28 |
| CYP1A2 | A - N | -0.08 | -0.09 | -0.06 | 0.09 |
| CYP1B1 | N - P | 0.02 | -0.04 | 0.08 | 0.87 |
| CYP1B1 | A - P | -0.07 | -0.09 | -0.05 | 0.64 |
| CYP1B1 | A - N | -0.09 | -0.11 | -0.07 | 0.03 |
| CYP2A6 | A - N | -0.12 | -0.14 | -0.1 | 0 |
| CYP2A13 | A - N | -0.09 | -0.1 | -0.07 | 0.05 |
| CYP2B6 | A - N | -0.08 | -0.1 | -0.06 | 0.08 |
| CYP2C8 | A - N | -0.07 | -0.09 | -0.06 | 0.1 |
| CYP2C9 | A - N | -0.06 | -0.08 | -0.04 | 0.14 |
| CYP2C19 | A - N | -0.11 | -0.12 | -0.09 | 0.01 |
| CYP2D6 | A - N | -0.11 | -0.13 | -0.1 | 0 |
| CYP2E1 | A - N | -0.18 | -0.2 | -0.16 | 0 |
| CYP2R1 | N - P | -0.28 | -0.37 | -0.2 | 0.39 |
| CYP2R1 | A - P | -0.34 | -0.36 | -0.32 | 0.37 |
| CYP2R1 | A - N | -0.06 | -0.08 | -0.04 | 0.27 |
| CYP3A4 | N - P | 1.43 | 1.35 | 1.5 | 0.05 |
| CYP3A4 | A - P | 1.32 | 1.3 | 1.34 | 0.15 |
| CYP3A4 | A - N | -0.1 | -0.12 | -0.08 | 0.03 |
| CYP3A5 | A - N | -0.04 | -0.06 | -0.02 | 0.45 |
| CYP3A7 | A - N | -0.09 | -0.11 | -0.07 | 0.06 |
| CYP7A1 | A - N | -0.1 | -0.12 | -0.08 | 0.04 |
| CYP8A1 | N - P | 0.05 | -0.02 | 0.13 | 0.95 |
| CYP8A1 | A - P | -0.04 | -0.07 | -0.02 | 0.96 |
| CYP8A1 | A - N | -0.1 | -0.12 | -0.08 | 0.04 |
| CYP8B1 | A - N | -0.12 | -0.14 | -0.1 | 0.41 |
| CYP11A1 | N - P | 0.2 | 0.12 | 0.28 | 0.45 |
| CYP11A1 | A - P | 0.03 | 0.01 | 0.05 | 0.92 |
| CYP11A1 | A - N | -0.17 | -0.19 | -0.15 | 0 |
| CYP11B1 | N - P | 0.32 | 0.26 | 0.39 | 0 |
| CYP11B1 | A - P | 0.2 | 0.18 | 0.22 | 0.13 |
| CYP11B1 | A - N | -0.12 | -0.14 | -0.1 | 0.01 |
| CYP11B2 | N - P | 0.2 | 0.13 | 0.27 | 0.48 |
| CYP11B2 | A - P | 0.08 | 0.06 | 0.1 | 0.81 |
| CYP11B2 | A - N | -0.12 | -0.14 | -0.1 | 0.01 |
| CYP17A1 | N - P | 0.74 | 0.65 | 0.83 | 0 |
| CYP17A1 | A - P | 0.61 | 0.59 | 0.63 | 0 |

| | | | | | |
|---|---|---|---|---|---|
| CYP17A1 | A - N | -0.13 | -0.15 | -0.11 | 0.02 |
| CYP19A1 | N - P | 0.58 | 0.5 | 0.65 | 0.07 |
| CYP19A1 | A - P | 0.46 | 0.44 | 0.48 | 0.23 |
| CYP19A1 | A - N | -0.11 | -0.13 | -0.09 | 0.02 |
| CYP21A2 | N - P | 0.16 | 0.09 | 0.23 | 0.13 |
| CYP21A2 | A - P | 0.12 | 0.1 | 0.14 | 0.3 |
| CYP21A2 | A - N | -0.04 | -0.06 | -0.02 | 0.4 |
| CYP46A1 | A - N | -0.22 | -0.24 | -0.2 | 0 |
| CYP51A1 | A - N | -0.31 | -0.33 | -0.29 | 0.23 |

**Table S8.** POPMUSIC ANOVA results (A: 'all mutations', N: 'Non-pathogenic', P: 'Pathogenic', DIFF: Difference between means, LWR: Lower end of 95% confidence interval, UPR: Upper end of 95% confidence interval).

| Name | Group Pair | DIFF | LWR | UPR | P-value |
|---|---|---|---|---|---|
| CYP1A1 | A - N | -0.07 | -0.08 | -0.05 | 0.14 |
| CYP1A2 | A - N | -0.14 | -0.16 | -0.13 | 0 |
| CYP1B1 | N - P | 0.07 | 0 | 0.13 | 0.62 |
| CYP1B1 | A - P | -0.01 | -0.03 | 0.01 | 0.93 |
| CYP1B1 | A - N | -0.08 | -0.1 | -0.06 | 0.05 |
| CYP2A6 | A - N | -0.16 | -0.18 | -0.14 | 0 |
| CYP2A13 | A - N | -0.14 | -0.16 | -0.12 | 0 |
| CYP2B6 | A - N | -0.17 | -0.19 | -0.15 | 0 |
| CYP2C8 | A - N | -0.13 | -0.15 | -0.11 | 0 |
| CYP2C9 | A - N | -0.14 | -0.16 | -0.12 | 0 |
| CYP2C19 | A - N | -0.16 | -0.17 | -0.14 | 0 |
| CYP2D6 | A - N | -0.16 | -0.17 | -0.14 | 0 |
| CYP2E1 | A - N | -0.23 | -0.25 | -0.21 | 0 |
| CYP2R1 | N - P | -0.11 | -0.2 | -0.02 | 0.74 |
| CYP2R1 | A - P | -0.21 | -0.22 | -0.19 | 0.6 |
| CYP2R1 | A - N | -0.09 | -0.11 | -0.07 | 0.08 |
| CYP3A4 | N - P | 0.73 | 0.64 | 0.82 | 0.41 |
| CYP3A4 | A - P | 0.62 | 0.6 | 0.65 | 0.52 |
| CYP3A4 | A - N | -0.11 | -0.13 | -0.09 | 0.04 |
| CYP3A5 | A - N | -0.12 | -0.14 | -0.1 | 0.03 |
| CYP3A7 | A - N | -0.14 | -0.16 | -0.12 | 0.01 |
| CYP7A1 | A - N | -0.14 | -0.16 | -0.12 | 0.01 |
| CYP8A1 | N - P | -0.28 | -0.35 | -0.21 | 0.72 |
| CYP8A1 | A - P | -0.52 | -0.54 | -0.5 | 0.6 |
| CYP8A1 | A - N | -0.24 | -0.26 | -0.22 | 0 |
| CYP8B1 | A - N | -0.27 | -0.3 | -0.25 | 0.06 |
| CYP11A1 | N - P | 0.32 | 0.24 | 0.4 | 0.25 |
| CYP11A1 | A - P | 0.18 | 0.17 | 0.2 | 0.59 |
| CYP11A1 | A - N | -0.14 | -0.16 | -0.12 | 0.01 |
| CYP11B1 | N - P | 0.44 | 0.37 | 0.51 | 0 |
| CYP11B1 | A - P | 0.24 | 0.23 | 0.26 | 0.08 |
| CYP11B1 | A - N | -0.2 | -0.22 | -0.18 | 0 |

| | | | | | |
|---|---|---|---|---|---|
| CYP11B2 | N - P | 0.5 | 0.42 | 0.58 | 0.1 |
| CYP11B2 | A - P | 0.33 | 0.31 | 0.35 | 0.34 |
| CYP11B2 | A - N | -0.17 | -0.19 | -0.15 | 0 |
| CYP17A1 | N - P | 0.39 | 0.3 | 0.48 | 0.02 |
| CYP17A1 | A - P | 0.18 | 0.16 | 0.2 | 0.32 |
| CYP17A1 | A - N | -0.2 | -0.22 | -0.18 | 0 |
| CYP19A1 | N - P | -0.06 | -0.14 | 0.02 | 0.86 |
| CYP19A1 | A - P | -0.21 | -0.23 | -0.19 | 0.61 |
| CYP19A1 | A - N | -0.15 | -0.17 | -0.13 | 0 |
| CYP21A2 | N - P | 0.18 | 0.11 | 0.25 | 0.1 |
| CYP21A2 | A - P | 0.12 | 0.1 | 0.14 | 0.3 |
| CYP21A2 | A - N | -0.06 | -0.08 | -0.04 | 0.2 |
| CYP46A1 | A - N | -0.34 | -0.36 | -0.31 | 0 |
| CYP51A1 | A - N | -0.39 | -0.41 | -0.37 | 0.16 |

**Table S9.** I-Mutant3.0 ANOVA results (A: 'all mutations', N: 'Non-pathogenic', P: 'Pathogenic', DIFF: Difference between means, LWR: Lower end of 95% confidence interval, UPR: Upper end of 95% confidence interval).

| Name | Group Pair | DIFF | LWR | UPR | P-value |
|---|---|---|---|---|---|
| CYP1A1 | A - N | 0 | -0.02 | 0.01 | 0.93 |
| CYP1A2 | A - N | -0.04 | -0.05 | -0.03 | 0.25 |
| **CYP1B1** | **N - P** | **0.12** | **0.07** | **0.16** | **0.22** |
| CYP1B1 | A - P | 0.05 | 0.04 | 0.07 | 0.63 |
| CYP1B1 | A - N | -0.06 | -0.08 | -0.05 | 0.03 |
| CYP2A6 | A - N | -0.08 | -0.1 | -0.07 | 0.01 |
| CYP2A13 | A - N | -0.07 | -0.08 | -0.05 | 0.03 |
| CYP2B6 | A - N | -0.09 | -0.1 | -0.07 | 0.01 |
| CYP2C8 | A - N | -0.03 | -0.04 | -0.01 | 0.42 |
| CYP2C9 | A - N | -0.04 | -0.05 | -0.02 | 0.24 |
| CYP2C19 | A - N | -0.08 | -0.1 | -0.07 | 0.01 |
| CYP2D6 | A - N | -0.1 | -0.11 | -0.09 | 0 |
| CYP2E1 | A - N | -0.12 | -0.13 | -0.1 | 0 |
| **CYP2R1** | **N - P** | **0.3** | **0.24** | **0.36** | **0.19** |
| CYP2R1 | A - P | 0.3 | 0.28 | 0.31 | 0.3 |
| CYP2R1 | A - N | 0 | -0.02 | 0.01 | 0.96 |
| **CYP3A4** | **N - P** | **0.23** | **0.17** | **0.29** | **0.7** |
| CYP3A4 | A - P | 0.12 | 0.1 | 0.14 | 0.87 |
| CYP3A4 | A - N | -0.11 | -0.13 | -0.1 | 0 |
| CYP3A5 | A - N | 0 | -0.02 | 0.01 | 0.97 |
| CYP3A7 | A - N | -0.07 | -0.09 | -0.06 | 0.07 |
| CYP7A1 | A - N | -0.08 | -0.09 | -0.07 | 0.02 |
| **CYP8A1** | **N - P** | **0.08** | **0.03** | **0.14** | **0.88** |
| CYP8A1 | A - P | -0.02 | -0.03 | 0 | 0.98 |
| CYP8A1 | A - N | -0.1 | -0.11 | -0.09 | 0 |
| CYP8B1 | A - N | 0.05 | 0.04 | 0.07 | 0.59 |
| **CYP11A1** | **N - P** | **0.27** | **0.2** | **0.33** | **0.22** |

| | | | | | |
|---|---|---|---|---|---|
| CYP11A1 | A - P | 0.16 | 0.14 | 0.17 | 0.56 |
| CYP11A1 | A - N | -0.11 | -0.13 | -0.1 | 0.01 |
| **CYP11B1** | **N - P** | **0.21** | **0.16** | **0.26** | **0.01** |
| CYP11B1 | A - P | 0.14 | 0.13 | 0.16 | 0.16 |
| CYP11B1 | A - N | -0.07 | -0.08 | -0.05 | 0.04 |
| **CYP11B2** | **N - P** | **0.2** | **0.15** | **0.26** | **0.34** |
| CYP11B2 | A - P | 0.11 | 0.09 | 0.12 | 0.66 |
| CYP11B2 | A - N | -0.09 | -0.11 | -0.08 | 0.01 |
| **CYP17A1** | **N - P** | **0.28** | **0.22** | **0.34** | **0.02** |
| CYP17A1 | A - P | 0.19 | 0.17 | 0.2 | 0.2 |
| CYP17A1 | A - N | -0.09 | -0.1 | -0.07 | 0.04 |
| **CYP19A1** | **N - P** | **0.17** | **0.12** | **0.23** | **0.44** |
| CYP19A1 | A - P | 0.13 | 0.11 | 0.14 | 0.67 |
| CYP19A1 | A - N | -0.05 | -0.06 | -0.03 | 0.21 |
| **CYP21A2** | **N - P** | **0.05** | **-0.01** | **0.1** | **0.57** |
| CYP21A2 | A - P | 0.01 | 0 | 0.03 | 0.91 |
| CYP21A2 | A - N | -0.03 | -0.05 | -0.02 | 0.34 |
| CYP46A1 | A - N | -0.15 | -0.17 | -0.14 | 0 |
| CYP51A1 | A - N | -0.07 | -0.09 | -0.06 | 0.08 |

**Table S10.** SIMBA ANOVA results (A: 'all mutations', N: 'Non-pathogenic', P: 'Pathogenic', DIFF: Difference between means, LWR: Lower end of 95% confidence interval, UPR: Upper end of 95% confidence interval).

| Name | Group Pair | DIFF | LWR | UPR | P-value |
|---|---|---|---|---|---|
| CYP1A1 | A - N | -0.08 | -0.1 | -0.05 | 0.23 |
| CYP1A2 | A - N | -0.11 | -0.14 | -0.08 | 0.1 |
| **CYP1B1** | **N - P** | **-0.4** | **-0.49** | **-0.32** | **0.01** |
| CYP1B1 | A - P | -0.48 | -0.51 | -0.46 | 0.02 |
| CYP1B1 | A - N | -0.08 | -0.1 | -0.05 | 0.18 |
| CYP2A6 | A - N | -0.14 | -0.16 | -0.11 | 0.03 |
| CYP2A13 | A - N | -0.13 | -0.16 | -0.1 | 0.04 |
| CYP2B6 | A - N | -0.16 | -0.18 | -0.13 | 0.01 |
| CYP2C8 | A - N | -0.17 | -0.19 | -0.14 | 0 |
| CYP2C9 | A - N | -0.16 | -0.18 | -0.13 | 0 |
| CYP2C19 | A - N | -0.08 | -0.11 | -0.06 | 0.16 |
| CYP2D6 | A - N | -0.16 | -0.19 | -0.14 | 0 |
| CYP2E1 | A - N | -0.29 | -0.31 | -0.26 | 0 |
| **CYP2R1** | **N - P** | **-0.3** | **-0.41** | **-0.19** | **0.48** |
| CYP2R1 | A - P | -0.32 | -0.35 | -0.29 | 0.57 |
| CYP2R1 | A - N | -0.02 | -0.05 | 0.01 | 0.78 |
| **CYP3A4** | **N - P** | **1.9** | **1.8** | **1.99** | **0.05** |
| CYP3A4 | A - P | 1.74 | 1.71 | 1.77 | 0.19 |
| CYP3A4 | A - N | -0.16 | -0.18 | -0.13 | 0.02 |
| CYP3A5 | A - N | -0.05 | -0.07 | -0.02 | 0.54 |
| CYP3A7 | A - N | -0.08 | -0.1 | -0.05 | 0.28 |
| CYP7A1 | A - N | -0.12 | -0.15 | -0.09 | 0.08 |

| | | | | | |
|---|---|---|---|---|---|
| **CYP8A1** | **N - P** | **0.36** | **0.27** | **0.44** | **0.71** |
| CYP8A1 | A - P | 0.15 | 0.13 | 0.18 | 0.91 |
| CYP8A1 | A - N | -0.2 | -0.23 | -0.18 | 0 |
| CYP8B1 | A - N | -0.21 | -0.24 | -0.19 | 0.28 |
| **CYP11A1** | **N - P** | **-0.36** | **-0.46** | **-0.26** | **0.3** |
| CYP11A1 | A - P | -0.54 | -0.56 | -0.51 | 0.27 |
| CYP11A1 | A - N | -0.18 | -0.21 | -0.15 | 0.02 |
| **CYP11B1** | **N - P** | **0.19** | **0.12** | **0.27** | **0.16** |
| CYP11B1 | A - P | 0.05 | 0.02 | 0.07 | 0.81 |
| CYP11B1 | A - N | -0.15 | -0.17 | -0.12 | 0.02 |
| **CYP11B2** | **N - P** | **-0.35** | **-0.45** | **-0.26** | **0.34** |
| CYP11B2 | A - P | -0.48 | -0.51 | -0.45 | 0.32 |
| CYP11B2 | A - N | -0.13 | -0.15 | -0.1 | 0.06 |
| **CYP17A1** | **N - P** | **0.39** | **0.29** | **0.5** | **0.05** |
| CYP17A1 | A - P | 0.26 | 0.23 | 0.29 | 0.33 |
| CYP17A1 | A - N | -0.13 | -0.16 | -0.11 | 0.08 |
| **CYP19A1** | **N - P** | **-0.35** | **-0.45** | **-0.24** | **0.4** |
| CYP19A1 | A - P | -0.49 | -0.52 | -0.46 | 0.38 |
| CYP19A1 | A - N | -0.15 | -0.17 | -0.12 | 0.05 |
| **CYP21A2** | **N - P** | **0.02** | **-0.07** | **0.11** | **0.88** |
| CYP21A2 | A - P | -0.05 | -0.08 | -0.02 | 0.78 |
| CYP21A2 | A - N | -0.07 | -0.09 | -0.04 | 0.32 |
| CYP46A1 | A - N | -0.28 | -0.31 | -0.26 | 0 |
| CYP51A1 | A - N | -0.17 | -0.2 | -0.14 | 0.65 |

**Table S11.** METASNP ANOVA results (A: 'all mutations', N: 'Non-pathogenic', P: 'Pathogenic', DIFF: Difference between means, LWR: Lower end of 95% confidence interval, UPR: Upper end of 95% confidence interval).

| Name | Group Pair | DIFF | LWR | UPR | P-value |
|---|---|---|---|---|---|
| CYP1A1 | A - N | 0.02 | 0.02 | 0.03 | 0.01 |
| CYP1A2 | A - N | 0.06 | 0.05 | 0.06 | 0 |
| CYP1B1 | N - P | -0.16 | -0.17 | -0.14 | 0 |
| CYP1B1 | A - P | -0.12 | -0.12 | -0.12 | 0 |
| CYP1B1 | A - N | 0.04 | 0.03 | 0.04 | 0 |
| CYP2A6 | A - N | 0.05 | 0.05 | 0.06 | 0 |
| CYP2A13 | A - N | 0.05 | 0.05 | 0.06 | 0 |
| CYP2B6 | A - N | 0.06 | 0.05 | 0.06 | 0 |
| CYP2C8 | A - N | 0.06 | 0.06 | 0.07 | 0 |
| CYP2C9 | A - N | 0.05 | 0.05 | 0.06 | 0 |
| CYP2C19 | A - N | 0.05 | 0.05 | 0.06 | 0 |
| CYP2D6 | A - N | 0.05 | 0.05 | 0.05 | 0 |
| CYP2E1 | A - N | 0.07 | 0.07 | 0.08 | 0 |
| CYP2R1 | N - P | -0.22 | -0.24 | -0.2 | 0.01 |
| CYP2R1 | A - P | -0.16 | -0.16 | -0.15 | 0.04 |
| CYP2R1 | A - N | 0.06 | 0.06 | 0.07 | 0 |
| CYP3A4 | N - P | -0.12 | -0.14 | -0.1 | 0.52 |

| | | | | | |
|---|---|---|---|---|---|
| CYP3A4 | A - P | -0.05 | -0.05 | -0.05 | 0.79 |
| CYP3A4 | A - N | 0.07 | 0.07 | 0.08 | 0 |
| CYP3A5 | A - N | 0.06 | 0.06 | 0.06 | 0 |
| CYP3A7 | A - N | 0.08 | 0.07 | 0.08 | 0 |
| CYP7A1 | A - N | 0.05 | 0.05 | 0.06 | 0 |
| CYP8A1 | N - P | 0.05 | 0.03 | 0.07 | 0.83 |
| CYP8A1 | A - P | 0.12 | 0.11 | 0.12 | 0.61 |
| CYP8A1 | A - N | 0.07 | 0.07 | 0.07 | 0 |
| CYP8B1 | A - N | -0.01 | -0.01 | 0 | 0.79 |
| CYP11A1 | N - P | -0.14 | -0.16 | -0.12 | 0.05 |
| CYP11A1 | A - P | -0.07 | -0.08 | -0.07 | 0.31 |
| CYP11A1 | A - N | 0.07 | 0.06 | 0.07 | 0 |
| CYP11B1 | N - P | -0.19 | -0.21 | -0.18 | 0 |
| CYP11B1 | A - P | -0.13 | -0.13 | -0.12 | 0 |
| CYP11B1 | A - N | 0.07 | 0.06 | 0.07 | 0 |
| CYP11B2 | N - P | -0.13 | -0.15 | -0.11 | 0.07 |
| CYP11B2 | A - P | -0.07 | -0.08 | -0.07 | 0.28 |
| CYP11B2 | A - N | 0.06 | 0.05 | 0.06 | 0 |
| CYP17A1 | N - P | -0.27 | -0.29 | -0.25 | 0 |
| CYP17A1 | A - P | -0.2 | -0.21 | -0.2 | 0 |
| CYP17A1 | A - N | 0.06 | 0.06 | 0.07 | 0 |
| CYP19A1 | N - P | -0.3 | -0.31 | -0.28 | 0 |
| CYP19A1 | A - P | -0.22 | -0.23 | -0.22 | 0 |
| CYP19A1 | A - N | 0.07 | 0.07 | 0.08 | 0 |
| CYP21A2 | N - P | -0.15 | -0.17 | -0.14 | 0 |
| CYP21A2 | A - P | -0.09 | -0.09 | -0.09 | 0 |
| CYP21A2 | A - N | 0.06 | 0.06 | 0.07 | 0 |
| CYP46A1 | A - N | 0.09 | 0.09 | 0.09 | 0 |
| CYP51A1 | A - N | 0.07 | 0.07 | 0.08 | 0 |

**Table S12.** SNAP2 ANOVA results (A: ‘all mutations’, N: ‘Non-pathogenic’, P: ‘Pathogenic’, DIFF: Difference between means, LWR: Lower end of 95% confidence interval, UPR: Upper end of 95% confidence interval).

| Name | Group Pair | DIFF | LWR | UPR | P-value |
|---|---|---|---|---|---|
| CYP1A1 | A - N | 1.65 | 0.53 | 2.76 | 0.54 |
| CYP1A2 | A - N | 10.42 | 9.3 | 11.53 | 0 |
| CYP1B1 | N - P | -54.2 | -58.86 | -49.54 | 0 |
| CYP1B1 | A - P | -42.28 | -43.4 | -41.17 | 0 |
| CYP1B1 | A - N | 11.92 | 10.83 | 13.01 | 0 |
| CYP2A6 | A - N | 10.5 | 9.4 | 11.6 | 0 |
| CYP2A13 | A - N | 10.77 | 9.68 | 11.86 | 0 |
| CYP2B6 | A - N | 15.62 | 14.51 | 16.74 | 0 |
| CYP2C8 | A - N | 20.45 | 19.35 | 21.55 | 0 |
| CYP2C9 | A - N | 15.65 | 14.57 | 16.74 | 0 |
| CYP2C19 | A - N | 13.38 | 12.26 | 14.51 | 0 |
| CYP2D6 | A - N | 10.5 | 9.38 | 11.61 | 0 |

| | | | | | |
|---|---|---|---|---|---|
| CYP2E1 | A - N | 17.22 | 16.1 | 18.33 | 0 |
| CYP2R1 | N - P | -41.54 | -47.55 | -35.53 | 0.09 |
| CYP2R1 | A - P | -28.06 | -29.22 | -26.9 | 0.25 |
| CYP2R1 | A - N | 13.48 | 12.34 | 14.62 | 0 |
| CYP3A4 | N - P | 29.19 | 23.53 | 34.85 | 0.62 |
| CYP3A4 | A - P | 45.21 | 44.09 | 46.32 | 0.43 |
| CYP3A4 | A - N | 16.02 | 14.92 | 17.11 | 0 |
| CYP3A5 | A - N | 13.68 | 12.57 | 14.79 | 0 |
| CYP3A7 | A - N | 20.27 | 19.17 | 21.37 | 0 |
| CYP7A1 | A - N | 12.02 | 10.95 | 13.1 | 0 |
| CYP8A1 | N - P | 25.89 | 20.84 | 30.93 | 0.64 |
| CYP8A1 | A - P | 39.01 | 37.9 | 40.13 | 0.49 |
| CYP8A1 | A - N | 13.13 | 12.04 | 14.22 | 0 |
| CYP8B1 | A - N | -9.34 | -10.48 | -8.21 | 0.27 |
| CYP11A1 | N - P | -55.07 | -60.55 | -49.6 | 0.01 |
| CYP11A1 | A - P | -38.27 | -39.38 | -37.15 | 0.06 |
| CYP11A1 | A - N | 16.81 | 15.71 | 17.91 | 0 |
| CYP11B1 | N - P | -61.16 | -66.08 | -56.25 | 0 |
| CYP11B1 | A - P | -40.28 | -41.4 | -39.16 | 0 |
| CYP11B1 | A - N | 20.88 | 19.78 | 21.97 | 0 |
| CYP11B2 | N - P | -40.57 | -45.67 | -35.47 | 0.05 |
| CYP11B2 | A - P | -26.08 | -27.21 | -24.96 | 0.2 |
| CYP11B2 | A - N | 14.48 | 13.38 | 15.58 | 0 |
| CYP17A1 | N - P | -79.47 | -85.35 | -73.59 | 0 |
| CYP17A1 | A - P | -61.41 | -62.54 | -60.28 | 0 |
| CYP17A1 | A - N | 18.06 | 16.95 | 19.17 | 0 |
| CYP19A1 | N - P | -88.99 | -94.34 | -83.64 | 0 |
| CYP19A1 | A - P | -72.46 | -73.59 | -71.33 | 0 |
| CYP19A1 | A - N | 16.53 | 15.43 | 17.64 | 0 |
| CYP21A2 | N - P | -48.65 | -53.57 | -43.73 | 0 |
| CYP21A2 | A - P | -32.27 | -33.42 | -31.12 | 0 |
| CYP21A2 | A - N | 16.38 | 15.25 | 17.5 | 0 |
| CYP46A1 | A - N | 20.22 | 19.1 | 21.35 | 0 |
| CYP51A1 | A - N | 18.79 | 17.65 | 19.93 | 0 |

**Table S13.** SIFT ANOVA results (A: 'all mutations', N: 'Non-pathogenic', P: 'Pathogenic', DIFF: Difference between means, LWR: Lower end of 95% confidence interval, UPR: Upper end of 95% confidence interval).

| Name | Group Pair | DIFF | LWR | UPR | P-value |
|---|---|---|---|---|---|
| CYP1A1 | A - N | -0.04 | -0.04 | -0.03 | 0 |
| CYP1A2 | A - N | -0.07 | -0.08 | -0.07 | 0 |
| CYP1B1 | N - P | 0.11 | 0.08 | 0.13 | 0.02 |
| CYP1B1 | A - P | 0.04 | 0.04 | 0.05 | 0.27 |
| CYP1B1 | A - N | -0.06 | -0.07 | -0.06 | 0 |
| CYP2A6 | A - N | -0.07 | -0.07 | -0.06 | 0 |
| CYP2A13 | A - N | -0.06 | -0.06 | -0.06 | 0 |

| | | | | | |
|---|---|---|---|---|---|
| CYP2B6 | A - N | -0.08 | -0.09 | -0.08 | 0 |
| CYP2C8 | A - N | -0.06 | -0.06 | -0.05 | 0 |
| CYP2C9 | A - N | -0.06 | -0.07 | -0.06 | 0 |
| CYP2C19 | A - N | -0.05 | -0.06 | -0.05 | 0 |
| CYP2D6 | A - N | -0.05 | -0.06 | -0.05 | 0 |
| CYP2E1 | A - N | -0.1 | -0.1 | -0.09 | 0 |
| CYP2R1 | N - P | 0.1 | 0.07 | 0.13 | 0.39 |
| CYP2R1 | A - P | 0.03 | 0.02 | 0.04 | 0.78 |
| CYP2R1 | A - N | -0.07 | -0.07 | -0.06 | 0 |
| CYP3A4 | N - P | 0.08 | 0.05 | 0.11 | 0.81 |
| CYP3A4 | A - P | -0.04 | -0.05 | -0.04 | 0.86 |
| CYP3A4 | A - N | -0.12 | -0.13 | -0.12 | 0 |
| CYP3A5 | A - N | -0.09 | -0.09 | -0.08 | 0 |
| CYP3A7 | A - N | -0.12 | -0.12 | -0.11 | 0 |
| CYP7A1 | A - N | -0.1 | -0.1 | -0.09 | 0 |
| CYP8A1 | N - P | 0.02 | -0.01 | 0.04 | 0.95 |
| CYP8A1 | A - P | -0.05 | -0.05 | -0.04 | 0.85 |
| CYP8A1 | A - N | -0.06 | -0.07 | -0.06 | 0 |
| CYP8B1 | A - N | 0.05 | 0.05 | 0.06 | 0.15 |
| CYP11A1 | N - P | 0.04 | 0.01 | 0.06 | 0.7 |
| CYP11A1 | A - P | -0.02 | -0.03 | -0.02 | 0.8 |
| CYP11A1 | A - N | -0.06 | -0.07 | -0.06 | 0 |
| CYP11B1 | N - P | 0.11 | 0.09 | 0.14 | 0.01 |
| CYP11B1 | A - P | 0.02 | 0.01 | 0.02 | 0.6 |
| CYP11B1 | A - N | -0.09 | -0.1 | -0.09 | 0 |
| CYP11B2 | N - P | 0.18 | 0.16 | 0.21 | 0.09 |
| CYP11B2 | A - P | 0.11 | 0.1 | 0.11 | 0.24 |
| CYP11B2 | A - N | -0.08 | -0.08 | -0.07 | 0 |
| CYP17A1 | N - P | 0.2 | 0.17 | 0.23 | 0 |
| CYP17A1 | A - P | 0.12 | 0.12 | 0.13 | 0.02 |
| CYP17A1 | A - N | -0.08 | -0.09 | -0.08 | 0 |
| CYP19A1 | N - P | 0.16 | 0.13 | 0.18 | 0.12 |
| CYP19A1 | A - P | 0.1 | 0.09 | 0.1 | 0.22 |
| CYP19A1 | A - N | -0.06 | -0.06 | -0.06 | 0 |
| CYP21A2 | N - P | 0.09 | 0.07 | 0.11 | 0.01 |
| CYP21A2 | A - P | 0.03 | 0.02 | 0.03 | 0.42 |
| CYP21A2 | A - N | -0.06 | -0.07 | -0.06 | 0 |
| CYP46A1 | A - N | -0.08 | -0.08 | -0.07 | 0 |
| CYP51A1 | A - N | -0.11 | -0.11 | -0.1 | 0 |

**Table S14.** Envision ANOVA results (A: ‘all mutations’, N: ‘Non-pathogenic’, P: ‘Pathogenic’, DIFF: Difference between means, LWR: Lower end of 95% confidence interval, UPR: Upper end of 95% confidence interval).

| Name | Group Pair | DIFF | LWR | UPR | P-value |
|---|---|---|---|---|---|
| CYP1A1 | A - N | -0.03 | -0.04 | -0.03 | 0 |
| CYP1A2 | A - N | -0.05 | -0.05 | -0.04 | 0 |

| | | | | | |
|---|---|---|---|---|---|
| CYP1B1 | N - P | 0.09 | 0.08 | 0.1 | 0 |
| CYP1B1 | A - P | 0.03 | 0.03 | 0.04 | 0.09 |
| CYP1B1 | A - N | -0.05 | -0.06 | -0.05 | 0 |
| CYP2A6 | A - N | -0.06 | -0.07 | -0.06 | 0 |
| CYP2A13 | A - N | -0.06 | -0.06 | -0.05 | 0 |
| CYP2B6 | A - N | -0.07 | -0.07 | -0.07 | 0 |
| CYP2C8 | Al - N | -0.07 | -0.07 | -0.07 | 0 |
| CYP2C9 | A - N | -0.06 | -0.06 | -0.06 | 0 |
| CYP2C19 | A - N | -0.06 | -0.07 | -0.06 | 0 |
| CYP2D6 | A - N | -0.05 | -0.06 | -0.05 | 0 |
| CYP2E1 | A - N | -0.05 | -0.05 | -0.05 | 0 |
| CYP2R1 | N - P | 0.06 | 0.05 | 0.07 | 0.25 |
| CYP2R1 | A - P | 0 | 0 | 0 | 0.99 |
| CYP2R1 | A - N | -0.06 | -0.06 | -0.06 | 0 |
| CYP3A4 | N - P | -0.01 | -0.03 | 0 | 0.91 |
| CYP3A4 | A - P | -0.08 | -0.08 | -0.07 | 0.54 |
| CYP3A4 | A - N | -0.06 | -0.07 | -0.06 | 0 |
| CYP3A5 | A - N | -0.06 | -0.06 | -0.06 | 0 |
| CYP3A7 | A - N | -0.07 | -0.07 | -0.07 | 0 |
| CYP7A1 | A - N | -0.05 | -0.05 | -0.05 | 0 |
| CYP8A1 | N - P | -0.05 | -0.06 | -0.04 | 0.69 |
| CYP8A1 | A - P | -0.11 | -0.11 | -0.11 | 0.44 |
| CYP8A1 | A - N | -0.06 | -0.06 | -0.06 | 0 |
| CYP8B1 | A - N | 0 | 0 | 0 | 0.77 |
| CYP11A1 | N - P | 0.12 | 0.11 | 0.13 | 0.01 |
| CYP11A1 | A - P | 0.05 | 0.05 | 0.05 | 0.3 |
| CYP11A1 | A - N | -0.07 | -0.07 | -0.06 | 0 |
| CYP11B1 | N - P | 0.06 | 0.06 | 0.07 | 0 |
| CYP11B1 | A - P | 0.02 | 0.02 | 0.02 | 0.04 |
| CYP11B1 | A - N | -0.04 | -0.04 | -0.04 | 0 |
| CYP11B2 | N - P | 0.06 | 0.05 | 0.06 | 0.03 |
| CYP11B2 | A - P | 0.02 | 0.02 | 0.02 | 0.44 |
| CYP11B2 | A - N | -0.04 | -0.04 | -0.03 | 0 |
| CYP17A1 | N - P | 0.09 | 0.08 | 0.1 | 0 |
| CYP17A1 | A - P | 0.06 | 0.05 | 0.06 | 0 |
| CYP17A1 | A - N | -0.04 | -0.04 | -0.03 | 0 |
| CYP19A1 | N - P | 0.29 | 0.27 | 0.3 | 0 |
| CYP19A1 | A - P | 0.23 | 0.22 | 0.23 | 0 |
| CYP19A1 | A - N | -0.06 | -0.06 | -0.06 | 0 |
| CYP21A2 | N - P | 0.06 | 0.06 | 0.07 | 0 |
| CYP21A2 | A - P | 0.03 | 0.03 | 0.03 | 0 |
| CYP21A2 | A - N | -0.03 | -0.04 | -0.03 | 0 |
| CYP46A1 | A - N | -0.06 | -0.07 | -0.06 | 0 |
| CYP51A1 | A - N | -0.04 | -0.04 | -0.04 | 0.29 |

**Table S15.** FATHMM ANOVA results (A: 'all mutations', N: 'Non-pathogenic', P: 'Pathogenic', DIFF: Difference between means, LWR: Lower end of 95% confidence interval, UPR: Upper end of 95% confidence interval).

| Name | Group Pair | DIFF | LWR | UPR | P-value |
|---|---|---|---|---|---|
| CYP1A1 | A - N | 0.05 | 0.04 | 0.07 | 0.14 |
| CYP1A2 | A - N | 0 | -0.01 | 0.02 | 0.95 |
| CYP1B1 | N - P | 0.7 | 0.64 | 0.77 | 0 |
| CYP1B1 | A - P | 0.67 | 0.66 | 0.69 | 0 |
| CYP1B1 | A - N | -0.03 | -0.05 | -0.02 | 0.3 |
| CYP2A6 | A - N | -0.11 | -0.16 | -0.06 | 0.3 |
| CYP2A13 | A - N | 0 | -0.05 | 0.05 | 0.99 |
| CYP2B6 | A - N | -0.12 | -0.17 | -0.07 | 0.3 |
| CYP2C8 | A - N | -0.11 | -0.14 | -0.08 | 0.13 |
| CYP2C9 | A - N | 0 | -0.03 | 0.03 | 0.98 |
| CYP2C19 | A - N | -0.14 | -0.17 | -0.11 | 0.04 |
| CYP2D6 | A - N | 0.06 | 0.04 | 0.08 | 0.17 |
| CYP2E1 | A - N | -0.14 | -0.19 | -0.09 | 0.28 |
| CYP2R1 | N - P | 0.39 | 0.29 | 0.48 | 0.3 |
| CYP2R1 | A - P | 0.37 | 0.35 | 0.39 | 0.27 |
| CYP2R1 | A - N | -0.02 | -0.03 | 0 | 0.73 |
| CYP3A4 | N - P | 0.03 | -0.09 | 0.14 | 0.98 |
| CYP3A4 | A - P | -0.08 | -0.11 | -0.06 | 0.94 |
| CYP3A4 | A - N | -0.11 | -0.13 | -0.09 | 0.07 |
| CYP3A5 | A - N | -0.01 | -0.03 | 0.01 | 0.89 |
| CYP3A7 | A - N | -0.03 | -0.05 | 0 | 0.66 |
| CYP7A1 | A - N | -0.09 | -0.11 | -0.06 | 0.24 |
| CYP8A1 | N - P | 1.41 | 1.18 | 1.64 | 0.57 |
| CYP8A1 | A - P | 1.26 | 1.21 | 1.31 | 0.61 |
| CYP8A1 | A - N | -0.15 | -0.2 | -0.1 | 0.19 |
| CYP8B1 | A - N | 0.07 | 0.02 | 0.12 | 0.84 |
| CYP11A1 | N - P | -0.16 | -0.24 | -0.07 | 0.61 |
| CYP11A1 | A - P | -0.22 | -0.23 | -0.2 | 0.52 |
| CYP11A1 | A - N | -0.06 | -0.08 | -0.04 | 0.23 |
| CYP11B1 | N - P | 0.37 | 0.33 | 0.42 | 0 |
| CYP11B1 | A - P | 0.31 | 0.3 | 0.33 | 0 |
| CYP11B1 | A - N | -0.06 | -0.07 | -0.05 | 0.03 |
| CYP11B2 | N - P | 0 | -0.05 | 0.06 | 0.99 |
| CYP11B2 | A - P | -0.01 | -0.03 | 0 | 0.95 |
| CYP11B2 | A - N | -0.02 | -0.03 | 0 | 0.56 |
| CYP17A1 | N - P | 0.73 | 0.63 | 0.83 | 0 |
| CYP17A1 | A - P | 0.72 | 0.71 | 0.74 | 0 |
| CYP17A1 | A - N | -0.01 | -0.02 | 0.01 | 0.84 |
| CYP19A1 | N - P | 2.06 | 1.99 | 2.12 | 0 |
| CYP19A1 | A - P | 2.02 | 2.01 | 2.03 | 0 |
| CYP19A1 | A - N | -0.04 | -0.05 | -0.02 | 0.24 |
| CYP21A2 | N - P | 0.65 | 0.58 | 0.73 | 0 |

| | | | | | |
|---|---|---|---|---|---|
| CYP21A2 | A - P | 0.61 | 0.6 | 0.63 | 0 |
| CYP21A2 | A - N | -0.04 | -0.06 | -0.03 | 0.28 |
| CYP46A1 | A - N | -0.08 | -0.09 | -0.07 | 0.04 |
| CYP51A1 | A - N | -0.31 | -0.33 | -0.29 | 0.27 |

**Table S16.** Combined ANOVA results (A: 'all mutations', N: 'Non-pathogenic', P: 'Pathogenic', DIFF: Difference between means, LWR: Lower end of 95% confidence interval, UPR: Upper end of 95% confidence interval) for sequence-based methods.

| Category | Method | Group Pair | DIFF | LWR | UPR | P-value |
|---|---|---|---|---|---|---|
| 26 CYP450 | SNAP2 | N-P | -49.15 | -50.2 | -48.09 | 0 |
| | | A-P | -35.38 | -35.6 | -35.15 | 0 |
| | | A-N | 13.76 | 13.55 | 13.98 | 0 |
| | METASNP | N-P | -0.15 | -0.15 | -0.15 | 0 |
| | | A-P | -0.1 | -0.1 | -0.09 | 0 |
| | | A-N | 0.05 | 0.05 | 0.05 | 0 |
| | SIFT | N-P | -0.02 | -0.03 | -0.02 | 0.39 |
| | | A-P | -0.08 | -0.08 | -0.07 | 0 |
| | | A-N | -0.05 | -0.05 | -0.04 | 0 |
| | FATHMM | N-P | 1.22 | 1.19 | 1.25 | 0 |
| | | A-P | 1.15 | 1.15 | 1.16 | 0 |
| | | A-N | -0.06 | -0.07 | -0.05 | 0 |
| | ENVISION | N-P | 0.02 | 0.01 | 0.02 | 0.02 |
| | | A-P | -0.02 | -0.02 | -0.02 | 0 |
| | | A-N | -0.05 | -0.05 | -0.04 | 0 |
| 10 Pathogenic mutation causing CYP450 | SNAP2 | N-P | -53.78 | -55.45 | -52.12 | 0 |
| | | A-P | -38.27 | -38.63 | -37.92 | 0 |
| | | A-N | 15.5 | 15.15 | 15.85 | 0 |
| | METASNP | N-P | -0.18 | -0.18 | -0.17 | 0 |
| | | A-P | -0.11 | -0.11 | -0.11 | 0 |
| | | A-N | 0.06 | 0.06 | 0.06 | 0 |
| | SIFT | N-P | 0.02 | 0.01 | 0.03 | 0.71 |
| | | A-P | 0 | 0 | 0 | 1 |
| | | A-N | -0.02 | -0.02 | -0.02 | 0 |
| | FATHMM | N-P | 0.84 | 0.81 | 0.88 | 0 |
| | | A-P | 0.78 | 0.77 | 0.79 | 0 |
| | | A-N | -0.06 | -0.07 | -0.05 | 0 |
| | ENVISION | N-P | 0.04 | 0.04 | 0.04 | 0 |
| | | A-P | -0.01 | -0.01 | -0.01 | 1 |
| | | A-N | -0.05 | -0.05 | -0.05 | 0 |
| Dominant mutation containing 4 CYP450 | SNAP2 | N-P | -56.06 | -58.59 | -53.54 | 0 |
| | | A-P | -40.04 | -40.6 | -39.47 | 0 |
| | | A-N | 16.02 | 15.47 | 16.58 | 0 |
| | METASNP | N-P | -0.17 | -0.18 | -0.16 | 0 |
| | | A-P | -0.11 | -0.12 | -0.11 | 0 |

| | | | | | | |
|---|---|---|---|---|---|---|
| | | A-N | 0.05 | 0.05 | 0.05 | 0 |
| | SIFT | N-P | 0.1 | 0.09 | 0.12 | 0 |
| | | A-P | 0.03 | 0.03 | 0.04 | 0.14 |
| | | A-N | -0.07 | -0.07 | -0.06 | 0 |
| | FATHMM | N-P | 0.6 | 0.57 | 0.6 | 0 |
| | | A-P | 0.56 | 0.55 | 0.56 | 0 |
| | | A-N | -0.04 | -0.05 | -0.03 | 0.03 |
| | ENVISION | N-P | 0.06 | 0.06 | 0.06 | 0 |
| | | A-P | 0.02 | 0.02 | 0.03 | 0 |
| | | A-N | -0.03 | -0.03 | -0.03 | 0 |

**Table S17.** CYP1B1: ANOVA results for change in 48 properties (N: 'Non-pathogenic', P: 'Pathogenic', DIFF: Difference between means, LWR: Lower end of 95% confidence interval, UPR: Upper end of 95% confidence interval).

| Name | Group Pair | DIFF | LWR | UPR | p-value |
|---|---|---|---|---|---|
| K | N - P | 0.32 | -0.02 | 0.67 | 0.64 |
| Ht | N - P | -0.43 | -0.54 | -0.32 | 0.05 |
| Hs | N - P | -0.18 | -0.3 | -0.06 | 0.47 |
| P | N - P | 3.7 | 1.55 | 5.86 | 0.4 |
| pHi | N - P | 0.02 | -0.17 | 0.21 | 0.95 |
| pK | N - P | 0.04 | 0.01 | 0.07 | 0.51 |
| Mw | N - P | -5.89 | -8.86 | -2.92 | 0.33 |
| B1 | N - P | -1.15 | -1.63 | -0.68 | 0.23 |
| Rf | N - P | -1.49 | -1.95 | -1.03 | 0.11 |
| u | N - P | -3.32 | -4.2 | -2.44 | 0.06 |
| Hnc | N - P | -0.21 | -0.32 | -0.1 | 0.34 |
| Esm | N - P | 0 | -0.01 | 0.02 | 0.99 |
| El | N - P | -0.03 | -0.04 | -0.02 | 0.17 |
| Et | N - P | -0.03 | -0.05 | -0.01 | 0.4 |
| Pa | N - P | 0.04 | 0.01 | 0.07 | 0.54 |
| Pb | N - P | -0.07 | -0.1 | -0.03 | 0.36 |
| Pt | N - P | 0 | -0.04 | 0.03 | 0.97 |
| Pc | N - P | 0 | -0.03 | 0.03 | 0.99 |
| Ca | N - P | -2.22 | -3.5 | -0.95 | 0.39 |
| F | N - P | -0.04 | -0.06 | -0.03 | 0.18 |
| Br | N - P | -0.04 | -0.06 | -0.03 | 0.18 |
| Ra | N - P | -0.24 | -0.4 | -0.07 | 0.48 |
| Ns | N - P | -0.16 | -0.24 | -0.07 | 0.36 |
| an | N - P | 0.01 | -0.05 | 0.07 | 0.92 |
| ac | N - P | 0.01 | -0.05 | 0.07 | 0.93 |
| am | N - P | 0.06 | 0 | 0.11 | 0.62 |
| V0 | N - P | -5.83 | -8.42 | -3.25 | 0.27 |
| Nm | N - P | -0.01 | -0.04 | 0.03 | 0.92 |
| Nl | N - P | -0.15 | -0.22 | -0.07 | 0.34 |
| Hgm | N - P | -0.25 | -0.35 | -0.15 | 0.21 |

| | | | | | |
|---|---|---|---|---|---|
| ASAD | N - P | -5.42 | -9.25 | -1.59 | 0.49 |
| ASAN | N - P | 3.72 | 1.44 | 6 | 0.42 |
| ΔASA | N - P | -8.96 | -11.99 | -5.92 | 0.15 |
| ΔGh | N - P | -0.38 | -0.59 | -0.18 | 0.35 |
| GhD | N - P | -0.12 | -0.59 | 0.35 | 0.9 |
| GhN | N - P | -0.65 | -0.87 | -0.42 | 0.16 |
| ΔHh | N - P | -0.17 | -0.44 | 0.1 | 0.76 |
| TΔSh | N - P | -0.22 | -0.33 | -0.1 | 0.37 |
| ΔCh | N - P | -4.2 | -5.37 | -3.02 | 0.08 |
| ΔGc | N - P | 0.03 | -0.16 | 0.22 | 0.94 |
| ΔHc | N - P | -0.9 | -1.19 | -0.62 | 0.12 |
| ΔSc | N - P | 0.93 | 0.73 | 1.13 | 0.02 |
| ΔG | N - P | -0.35 | -0.42 | -0.29 | 0.01 |
| ΔH | N - P | -1.08 | -1.32 | -0.83 | 0.03 |
| TΔS | N - P | 0.72 | 0.53 | 0.91 | 0.06 |
| v | N - P | 0.24 | 0.07 | 0.41 | 0.48 |
| s | N - P | 0.24 | 0.07 | 0.41 | 0.48 |
| ff | N - P | 0.13 | -0.02 | 0.29 | 0.67 |

**Table S18.** CYP2R1: ANOVA results for change in 48 properties (N: 'Non-pathogenic', P: 'Pathogenic', DIFF: Difference between means, LWR: Lower end of 95% confidence interval, UPR: Upper end of 95% confidence interval).

| Name | Group Pair | DIFF | LWR | UPR | p-value |
|---|---|---|---|---|---|
| K | N - P | -1.6 | -2.05 | -1.16 | 0.38 |
| Ht | N - P | 0.33 | 0.18 | 0.48 | 0.59 |
| Hs | N - P | 0.31 | 0.16 | 0.47 | 0.62 |
| P | N - P | -41.79 | -44.62 | -38.96 | 0 |
| pHi | N - P | -3.81 | -4.05 | -3.57 | 0 |
| pK | N - P | 0.36 | 0.32 | 0.41 | 0.04 |
| Mw | N - P | -58.52 | -62.57 | -54.47 | 0 |
| B1 | N - P | -0.8 | -1.43 | -0.17 | 0.76 |
| Rf | N - P | 5.42 | 4.83 | 6 | 0.02 |
| u | N - P | -14.26 | -15.48 | -13.05 | 0 |
| Hnc | N - P | 2.51 | 2.38 | 2.65 | 0 |
| Esm | N - P | 0.33 | 0.31 | 0.35 | 0 |
| El | N - P | 0.01 | -0.01 | 0.02 | 0.93 |
| Et | N - P | 0.33 | 0.31 | 0.35 | 0 |
| Pa | N - P | -0.15 | -0.19 | -0.11 | 0.37 |
| Pb | N - P | 0.03 | -0.01 | 0.07 | 0.87 |
| Pt | N - P | 0.18 | 0.13 | 0.22 | 0.36 |
| Pc | N - P | 0.12 | 0.07 | 0.16 | 0.5 |
| Ca | N - P | -26.35 | -28.05 | -24.66 | 0 |
| F | N - P | -0.03 | -0.05 | -0.01 | 0.72 |
| Br | N - P | 0.26 | 0.24 | 0.28 | 0 |
| Ra | N - P | 1.14 | 0.92 | 1.36 | 0.2 |
| Ns | N - P | 0.72 | 0.61 | 0.82 | 0.09 |

| | | | | | |
|---|---|---|---|---|---|
| an | N - P | 0.25 | 0.18 | 0.32 | 0.37 |
| ac | N - P | 0.36 | 0.28 | 0.44 | 0.25 |
| am | N - P | -0.78 | -0.86 | -0.71 | 0.01 |
| V0 | N - P | -43.42 | -46.89 | -39.94 | 0 |
| Nm | N - P | -0.17 | -0.21 | -0.13 | 0.3 |
| Nl | N - P | 0.45 | 0.35 | 0.54 | 0.26 |
| Hgm | N - P | 0.3 | 0.18 | 0.43 | 0.56 |
| ASAD | N - P | -75.44 | -80.66 | -70.23 | 0 |
| ASAN | N - P | -47.24 | -50.15 | -44.34 | 0 |
| ΔASA | N - P | -28.76 | -32.89 | -24.63 | 0.09 |
| ΔGh | N - P | 4.73 | 4.47 | 4.99 | 0 |
| GhD | N - P | 10.05 | 9.45 | 10.65 | 0 |
| GhN | N - P | 5.46 | 5.17 | 5.75 | 0 |
| ΔHh | N - P | 6.57 | 6.21 | 6.93 | 0 |
| TΔSh | N - P | -1.84 | -2 | -1.69 | 0 |
| ΔCh | N - P | 3.01 | 1.49 | 4.54 | 0.63 |
| ΔGc | N - P | -4.24 | -4.48 | -3.99 | 0 |
| ΔHc | N - P | -3.2 | -3.58 | -2.82 | 0.04 |
| ΔSc | N - P | -1.04 | -1.31 | -0.77 | 0.34 |
| ΔG | N - P | 0.49 | 0.4 | 0.58 | 0.17 |
| ΔH | N - P | 3.37 | 3.05 | 3.7 | 0.01 |
| TΔS | N - P | -2.89 | -3.13 | -2.64 | 0 |
| v | N - P | -4.13 | -4.44 | -3.83 | 0 |
| s | N - P | -3.44 | -3.66 | -3.21 | 0 |
| ff | N - P | -3.4 | -3.6 | -3.2 | 0 |

**Table S19.** CYP3A4: ANOVA results for change in 48 properties (N: 'Non-pathogenic', P: 'Pathogenic', DIFF: Difference between means, LWR: Lower end of 95% confidence interval, UPR: Upper end of 95% confidence interval).

| Name | Group Pair | DIFF | LWR | UPR | p-value |
|---|---|---|---|---|---|
| K | N - P | -0.89 | -1.3 | -0.48 | 0.84 |
| Ht | N - P | 2.96 | 2.82 | 3.09 | 0.03 |
| Hs | N - P | 3.37 | 3.22 | 3.52 | 0.03 |
| P | N - P | -0.15 | -2.67 | 2.36 | 1 |
| pHi | N - P | 0.53 | 0.31 | 0.76 | 0.82 |
| pK | N - P | -0.76 | -0.8 | -0.72 | 0.06 |
| Mw | N - P | 14.42 | 11.07 | 17.76 | 0.68 |
| B1 | N - P | 5.83 | 5.31 | 6.35 | 0.28 |
| Rf | N - P | 7.13 | 6.6 | 7.66 | 0.2 |
| u | N - P | 9.27 | 8.28 | 10.26 | 0.37 |
| Hnc | N - P | 1.31 | 1.19 | 1.44 | 0.31 |
| Esm | N - P | -0.07 | -0.09 | -0.05 | 0.68 |
| El | N - P | 0.25 | 0.24 | 0.26 | 0.09 |
| Et | N - P | 0.18 | 0.16 | 0.2 | 0.4 |
| Pa | N - P | 0.26 | 0.23 | 0.3 | 0.49 |
| Pb | N - P | 0.46 | 0.42 | 0.51 | 0.29 |

| | | | | | |
|---|---|---|---|---|---|
| Pt | N - P | -0.54 | -0.59 | -0.5 | 0.25 |
| Pc | N - P | -0.46 | -0.5 | -0.42 | 0.26 |
| Ca | N - P | 12.3 | 10.85 | 13.76 | 0.42 |
| F | N - P | -0.22 | -0.24 | -0.21 | 0.21 |
| Br | N - P | 0.34 | 0.32 | 0.35 | 0.09 |
| Ra | N - P | 5.18 | 4.97 | 5.38 | 0.02 |
| Ns | N - P | 1.76 | 1.67 | 1.86 | 0.08 |
| an | N - P | 0.52 | 0.45 | 0.59 | 0.48 |
| ac | N - P | -0.66 | -0.74 | -0.57 | 0.44 |
| am | N - P | 0.41 | 0.35 | 0.48 | 0.56 |
| V0 | N - P | 32.55 | 29.6 | 35.5 | 0.29 |
| Nm | N - P | 0.23 | 0.2 | 0.27 | 0.56 |
| Nl | N - P | 1.56 | 1.47 | 1.65 | 0.09 |
| Hgm | N - P | 2.67 | 2.55 | 2.8 | 0.04 |
| ASAD | N - P | 43.25 | 38.84 | 47.66 | 0.35 |
| ASAN | N - P | -23.55 | -26.2 | -20.91 | 0.39 |
| ΔASA | N - P | 66.74 | 63.13 | 70.34 | 0.07 |
| ΔGh | N - P | 2.1 | 1.86 | 2.34 | 0.4 |
| GhD | N - P | 3.75 | 3.15 | 4.34 | 0.54 |
| GhN | N - P | 1.52 | 1.26 | 1.77 | 0.57 |
| ΔHh | N - P | 0.3 | -0.03 | 0.62 | 0.93 |
| TΔSh | N - P | 1.81 | 1.67 | 1.94 | 0.21 |
| ΔCh | N - P | 25.56 | 24.19 | 26.93 | 0.07 |
| ΔGc | N - P | -1.94 | -2.17 | -1.7 | 0.42 |
| ΔHc | N - P | -0.24 | -0.59 | 0.1 | 0.95 |
| ΔSc | N - P | -1.68 | -1.91 | -1.45 | 0.48 |
| ΔG | N - P | 0.17 | 0.1 | 0.25 | 0.83 |
| ΔH | N - P | 0.04 | -0.25 | 0.34 | 0.99 |
| TΔS | N - P | 0.13 | -0.09 | 0.35 | 0.95 |
| v | N - P | 1.18 | 0.93 | 1.44 | 0.65 |
| s | N - P | 0.3 | 0.1 | 0.5 | 0.89 |
| ff | N - P | 1.15 | 0.97 | 1.32 | 0.53 |

**Table S20.** CYP8A1: ANOVA results for change in 48 properties (N: 'Non-pathogenic', P: 'Pathogenic', DIFF: Difference between means, LWR: Lower end of 95% confidence interval, UPR: Upper end of 95% confidence interval).

| Name | Group Pair | DIFF | LWR | UPR | p-value |
|---|---|---|---|---|---|
| K | N - P | 4.79 | 4.42 | 5.16 | 0.24 |
| Ht | N - P | 0.87 | 0.75 | 1 | 0.52 |
| Hs | N - P | 0.83 | 0.69 | 0.96 | 0.58 |
| P | N - P | 44.2 | 41.62 | 46.78 | 0.12 |
| pHi | N - P | 4.76 | 4.53 | 5 | 0.07 |
| pK | N - P | -0.4 | -0.43 | -0.37 | 0.28 |
| Mw | N - P | 27.72 | 24.21 | 31.23 | 0.47 |
| B1 | N - P | 0.3 | -0.22 | 0.82 | 0.96 |
| Rf | N - P | -4.13 | -4.64 | -3.62 | 0.46 |

| | | | | | |
|---|---|---|---|---|---|
| u | N - P | 9.71 | 8.7 | 10.71 | 0.38 |
| Hnc | N - P | -1.52 | -1.65 | -1.39 | 0.29 |
| Esm | N - P | -0.18 | -0.2 | -0.16 | 0.36 |
| El | N - P | 0.17 | 0.15 | 0.18 | 0.28 |
| Et | N - P | -0.01 | -0.03 | 0.01 | 0.97 |
| Pa | N - P | -0.16 | -0.2 | -0.13 | 0.66 |
| Pb | N - P | -0.09 | -0.13 | -0.05 | 0.84 |
| Pt | N - P | -0.03 | -0.07 | 0.01 | 0.94 |
| Pc | N - P | 0.19 | 0.16 | 0.23 | 0.62 |
| Ca | N - P | 18.22 | 16.69 | 19.74 | 0.28 |
| F | N - P | -0.05 | -0.06 | -0.03 | 0.79 |
| Br | N - P | -0.01 | -0.02 | 0.01 | 0.98 |
| Ra | N - P | 0.02 | -0.17 | 0.2 | 0.99 |
| Ns | N - P | 0.41 | 0.31 | 0.5 | 0.7 |
| an | N - P | -0.31 | -0.38 | -0.25 | 0.65 |
| ac | N - P | -0.9 | -0.97 | -0.83 | 0.25 |
| am | N - P | 0 | -0.07 | 0.06 | 1 |
| V0 | N - P | 32.01 | 28.94 | 35.09 | 0.34 |
| Nm | N - P | -0.11 | -0.14 | -0.07 | 0.78 |
| Nl | N - P | 0.9 | 0.81 | 0.99 | 0.35 |
| Hgm | N - P | 0.73 | 0.62 | 0.85 | 0.56 |
| ASAD | N - P | 44.1 | 39.54 | 48.67 | 0.38 |
| ASAN | N - P | 18.43 | 15.68 | 21.19 | 0.54 |
| ΔASA | N - P | 26.03 | 22.6 | 29.47 | 0.49 |
| ΔGh | N - P | -1.97 | -2.21 | -1.74 | 0.44 |
| GhD | N - P | -5.3 | -5.88 | -4.73 | 0.4 |
| GhN | N - P | -3.35 | -3.62 | -3.08 | 0.26 |
| Hh | N - P | -4.14 | -4.46 | -3.82 | 0.24 |
| TΔSh | N - P | 2.16 | 2.03 | 2.3 | 0.15 |
| ΔCh | N - P | 13.48 | 12.18 | 14.79 | 0.35 |
| ΔGc | N - P | 1.6 | 1.38 | 1.81 | 0.5 |
| ΔHc | N - P | 2.09 | 1.76 | 2.42 | 0.56 |
| ΔSc | N - P | -0.5 | -0.73 | -0.26 | 0.85 |
| ΔG | N - P | -0.38 | -0.46 | -0.3 | 0.68 |
| ΔH | N - P | -2.04 | -2.36 | -1.73 | 0.56 |
| TΔS | N - P | 1.67 | 1.43 | 1.91 | 0.53 |
| v | N - P | 1.93 | 1.67 | 2.2 | 0.51 |
| s | N - P | 1.69 | 1.49 | 1.9 | 0.45 |
| ff | N - P | 1.86 | 1.68 | 2.04 | 0.36 |

**Table S21.** CYP11A1: ANOVA results for change in 48 properties (N: 'Non-pathogenic', P: 'Pathogenic', DIFF: Difference between means, LWR: Lower end of 95% confidence interval, UPR: Upper end of 95% confidence interval).

| Name | Group Pair | DIFF | LWR | UPR | p-value |
|---|---|---|---|---|---|
| K | N - P | 0.37 | -0.05 | 0.79 | 0.81 |
| Ht | N - P | -1.19 | -1.32 | -1.05 | 0.02 |

| | | | | | |
|---|---|---|---|---|---|
| Hs | N - P | 0.28 | 0.13 | 0.42 | 0.6 |
| P | N - P | 1.32 | -1.45 | 4.09 | 0.9 |
| pHi | N - P | 0.55 | 0.29 | 0.8 | 0.56 |
| pK | N - P | -0.11 | -0.14 | -0.07 | 0.44 |
| Mw | N - P | -26 | -29.48 | -22.52 | 0.04 |
| B1 | N - P | -2.82 | -3.38 | -2.25 | 0.17 |
| Rf | N - P | -2.19 | -2.74 | -1.64 | 0.28 |
| u | N - P | -5.98 | -7 | -4.96 | 0.11 |
| Hnc | N - P | -0.35 | -0.48 | -0.22 | 0.47 |
| Esm | N - P | 0.1 | 0.08 | 0.11 | 0.13 |
| El | N - P | -0.06 | -0.08 | -0.05 | 0.24 |
| Et | N - P | 0.03 | 0.01 | 0.06 | 0.67 |
| Pa | N - P | 0.13 | 0.09 | 0.16 | 0.32 |
| Pb | N - P | -0.05 | -0.09 | 0 | 0.78 |
| Pt | N - P | -0.19 | -0.23 | -0.15 | 0.22 |
| Pc | N - P | -8.28 | -9.8 | -6.77 | 0.14 |
| Ca | N - P | -8.28 | -9.8 | -6.77 | 0.14 |
| F | N - P | 0 | -0.02 | 0.02 | 0.99 |
| Br | N - P | 0.03 | 0.01 | 0.05 | 0.64 |
| Ra | N - P | -0.53 | -0.73 | -0.32 | 0.48 |
| Ns | N - P | 0.03 | -0.07 | 0.13 | 0.94 |
| an | N - P | 0.24 | 0.18 | 0.3 | 0.29 |
| ac | N - P | -0.47 | -0.54 | -0.39 | 0.09 |
| am | N - P | 0.5 | 0.43 | 0.57 | 0.05 |
| V0 | N - P | -16.23 | -19.29 | -13.16 | 0.15 |
| Nm | N - P | 0.24 | 0.2 | 0.27 | 0.08 |
| Nl | N - P | -0.06 | -0.15 | 0.03 | 0.86 |
| Hgm | N - P | -0.07 | -0.2 | 0.05 | 0.87 |
| ASAD | N - P | -19.18 | -23.8 | -14.57 | 0.26 |
| ASAN | N - P | -3.56 | -6.39 | -0.73 | 0.73 |
| ΔASA | N - P | -14.96 | -18.65 | -11.27 | 0.27 |
| ΔGh | N - P | 0.23 | -0.01 | 0.47 | 0.79 |
| GhD | N - P | -0.8 | -1.07 | -0.52 | 0.43 |
| GhN | N - P | -0.8 | -1.07 | -0.52 | 0.43 |
| Hh | N - P | 0.73 | 0.41 | 1.05 | 0.53 |
| TΔSh | N - P | -0.5 | -0.64 | -0.36 | 0.34 |
| ΔCh | N - P | -5.4 | -6.83 | -3.97 | 0.3 |
| ΔGc | N - P | -0.96 | -1.19 | -0.74 | 0.24 |
| ΔHc | N - P | -2.75 | -3.1 | -2.4 | 0.03 |
| ΔSc | N - P | 1.79 | 1.53 | 2.04 | 0.06 |
| ΔG | N - P | -0.73 | -0.81 | -0.64 | 0.02 |
| ΔH | N - P | -2.02 | -2.35 | -1.7 | 0.09 |
| TΔS | N - P | 1.29 | 1.05 | 1.54 | 0.15 |
| v | N - P | -2.23 | -2.49 | -1.97 | 0.02 |
| s | N - P | 0.51 | 0.3 | 0.72 | 0.51 |
| ff | N - P | 0.29 | 0.11 | 0.48 | 0.67 |

**Table S22.** CYP11B1: ANOVA results for change in 48 properties (N: 'Non-pathogenic', P: 'Pathogenic', DIFF: Difference between means, LWR: Lower end of 95% confidence interval, UPR: Upper end of 95% confidence interval).

| Name | Group Pair | DIFF | LWR | UPR | p-value |
|---|---|---|---|---|---|
| K | N - P | 1.05 | 0.68 | 1.42 | 0.11 |
| Ht | N - P | 0.04 | -0.07 | 0.16 | 0.82 |
| Hs | N - P | -0.16 | -0.27 | -0.04 | 0.46 |
| P | N - P | 4.86 | 2.45 | 7.27 | 0.26 |
| pHi | N - P | 1.24 | 1.03 | 1.45 | 0 |
| pK | N - P | -0.06 | -0.09 | -0.03 | 0.27 |
| Mw | N - P | 11.42 | 8.47 | 14.37 | 0.03 |
| B1 | N - P | 0.62 | 0.16 | 1.08 | 0.45 |
| Rf | N - P | -0.04 | -0.51 | 0.43 | 0.96 |
| u | N - P | 2.07 | 1.24 | 2.89 | 0.16 |
| Hnc | N - P | -0.6 | -0.71 | -0.48 | 0 |
| Esm | N - P | -0.08 | -0.09 | -0.06 | 0.01 |
| El | N - P | 0.02 | 0 | 0.03 | 0.45 |
| Et | N - P | -0.06 | -0.08 | -0.04 | 0.08 |
| Pa | N - P | -0.05 | -0.08 | -0.02 | 0.31 |
| Pb | N - P | 0.02 | -0.02 | 0.06 | 0.77 |
| Pt | N - P | -0.02 | -0.05 | 0.02 | 0.81 |
| Pc | N - P | 0.04 | 0.01 | 0.07 | 0.46 |
| Ca | N - P | 6.16 | 4.87 | 7.44 | 0.01 |
| F | N - P | 0 | -0.01 | 0.02 | 0.92 |
| Br | N - P | -0.06 | -0.08 | -0.04 | 0.05 |
| Ra | N - P | -0.34 | -0.5 | -0.17 | 0.25 |
| Ns | N - P | -0.19 | -0.27 | -0.1 | 0.22 |
| an | N - P | -0.09 | -0.15 | -0.04 | 0.36 |
| ac | N - P | -0.1 | -0.16 | -0.03 | 0.39 |
| am | N - P | -0.11 | -0.17 | -0.05 | 0.3 |
| V0 | N - P | 11.23 | 8.66 | 13.81 | 0.01 |
| Nm | N - P | -0.05 | -0.08 | -0.02 | 0.36 |
| Nl | N - P | 0 | -0.07 | 0.08 | 0.99 |
| Hgm | N - P | -0.15 | -0.25 | -0.05 | 0.4 |
| ASAD | N - P | 14.3 | 10.47 | 18.12 | 0.04 |
| ASAN | N - P | 10.47 | 7.99 | 12.96 | 0.02 |
| ΔASA | N - P | 3.82 | 0.97 | 6.67 | 0.45 |
| ΔGh | N - P | -0.72 | -0.94 | -0.51 | 0.06 |
| GhD | N - P | -1.7 | -2.21 | -1.18 | 0.06 |
| GhN | N - P | -0.95 | -1.19 | -0.71 | 0.03 |
| ΔHh | N - P | -1.27 | -1.55 | -0.99 | 0.01 |
| TΔSh | N - P | 0.54 | 0.43 | 0.66 | 0.01 |
| Ch | N - P | 2.52 | 1.35 | 3.7 | 0.23 |
| ΔGc | N - P | 0.67 | 0.47 | 0.87 | 0.06 |
| ΔHc | N - P | 0.56 | 0.28 | 0.84 | 0.26 |

| | | | | | |
|---|---|---|---|---|---|
| ΔSc | N - P | 0.11 | -0.1 | 0.31 | 0.77 |
| ΔG | N - P | -0.06 | -0.13 | 0.01 | 0.62 |
| ΔH | N - P | -0.71 | -0.98 | -0.43 | 0.15 |
| TΔS | N - P | 0.65 | 0.43 | 0.86 | 0.09 |
| v | N - P | 0.97 | 0.75 | 1.2 | 0.01 |
| s | N - P | 0.64 | 0.46 | 0.83 | 0.05 |
| ff | N - P | 0.49 | 0.33 | 0.65 | 0.09 |

**Table S23.** CYP11B2: ANOVA results for change in 48 properties (N: ‘Non-pathogenic’, P: ‘Pathogenic’, DIFF: Difference between means, LWR: Lower end of 95% confidence interval, UPR: Upper end of 95% confidence interval).

| Name | Group Pair | DIFF | LWR | UPR | p-value |
|---|---|---|---|---|---|
| K | N - P | -3 | -3.38 | -2.62 | 0.05 |
| Ht | N - P | -1.47 | -1.6 | -1.34 | 0 |
| Hs | N - P | -0.63 | -0.77 | -0.49 | 0.26 |
| P | N - P | 3.48 | 0.95 | 6.02 | 0.73 |
| pHi | N - P | 0.31 | 0.08 | 0.53 | 0.74 |
| pK | N - P | 0.03 | -0.01 | 0.06 | 0.85 |
| Mw | N - P | -10.71 | -13.9 | -7.51 | 0.41 |
| B1 | N - P | -0.5 | -1.03 | 0.02 | 0.81 |
| Rf | N - P | -3.47 | -4.01 | -2.92 | 0.12 |
| u | N - P | -8.18 | -9.1 | -7.26 | 0.03 |
| Hnc | N - P | -0.76 | -0.89 | -0.63 | 0.15 |
| Esm | N - P | 0 | -0.01 | 0.02 | 0.97 |
| El | N - P | -0.07 | -0.09 | -0.06 | 0.2 |
| Et | N - P | -0.08 | -0.1 | -0.05 | 0.37 |
| Pa | N - P | -0.19 | -0.23 | -0.16 | 0.15 |
| Pb | N - P | 0.09 | 0.05 | 0.13 | 0.58 |
| Pt | N - P | 0.04 | 0 | 0.08 | 0.79 |
| Pc | N - P | 0.14 | 0.1 | 0.17 | 0.33 |
| Ca | N - P | -4.93 | -6.34 | -3.52 | 0.39 |
| F | N - P | 0.05 | 0.04 | 0.07 | 0.43 |
| Br | N - P | -0.1 | -0.12 | -0.08 | 0.2 |
| Ra | N - P | -1.11 | -1.32 | -0.91 | 0.18 |
| Ns | N - P | -0.24 | -0.33 | -0.14 | 0.54 |
| an | N - P | -0.12 | -0.19 | -0.06 | 0.61 |
| ac | N - P | -0.38 | -0.45 | -0.31 | 0.18 |
| am | N - P | 0.05 | -0.01 | 0.12 | 0.84 |
| V0 | N - P | -11.38 | -14.21 | -8.54 | 0.32 |
| Nm | N - P | -0.18 | -0.22 | -0.15 | 0.19 |
| Nl | N - P | -0.11 | -0.2 | -0.03 | 0.75 |
| Hgm | N - P | -0.91 | -1.02 | -0.79 | 0.05 |
| ASAD | N - P | -11.88 | -16.1 | -7.65 | 0.49 |
| ASAN | N - P | 10.83 | 8.16 | 13.5 | 0.32 |
| ASA | N - P | -22.4 | -25.84 | -18.97 | 0.11 |
| ΔGh | N - P | -0.86 | -1.09 | -0.62 | 0.37 |

| | | | | | |
|---|---|---|---|---|---|
| GhD | N - P | -0.77 | -1.33 | -0.2 | 0.74 |
| GhN | N - P | -1.34 | -1.61 | -1.07 | 0.22 |
| ΔHh | N - P | -0.46 | -0.77 | -0.15 | 0.71 |
| TΔSh | N - P | -0.4 | -0.53 | -0.27 | 0.45 |
| ΔCh | N - P | -7.66 | -9.04 | -6.29 | 0.17 |
| ΔGc | N - P | 0.2 | -0.02 | 0.42 | 0.82 |
| ΔHc | N - P | -1.26 | -1.57 | -0.94 | 0.32 |
| ΔSc | N - P | 1.45 | 1.21 | 1.68 | 0.13 |
| ΔG | N - P | -0.67 | -0.74 | -0.59 | 0.03 |
| ΔH | N - P | -1.71 | -2.01 | -1.4 | 0.16 |
| TΔS | N - P | 1.04 | 0.81 | 1.27 | 0.26 |
| v | N - P | -0.92 | -1.16 | -0.68 | 0.35 |
| s | N - P | 0.54 | 0.34 | 0.74 | 0.5 |
| ff | N - P | 0.34 | 0.16 | 0.51 | 0.64 |

**Table S24.** CYP17A1: ANOVA results for change in 48 properties (N: 'Non-pathogenic', P: 'Pathogenic', DIFF: Difference between means, LWR: Lower end of 95% confidence interval, UPR: Upper end of 95% confidence interval).

| Name | Group Pair | DIFF | LWR | UPR | p-value |
|---|---|---|---|---|---|
| K | N - P | 1.58 | 1.14 | 2.02 | 0.07 |
| Ht | N - P | 0.26 | 0.12 | 0.4 | 0.35 |
| Hs | N - P | -0.28 | -0.44 | -0.13 | 0.37 |
| P | N - P | 5.6 | 2.76 | 8.44 | 0.32 |
| pHi | N - P | 1.51 | 1.25 | 1.76 | 0 |
| pK | N - P | -0.01 | -0.05 | 0.02 | 0.84 |
| Mw | N - P | 23.66 | 20.2 | 27.13 | 0 |
| B1 | N - P | 0.45 | -0.17 | 1.06 | 0.72 |
| Rf | N - P | 1.25 | 0.68 | 1.83 | 0.27 |
| u | N - P | 3.32 | 2.33 | 4.31 | 0.09 |
| Hnc | N - P | -0.7 | -0.83 | -0.57 | 0.01 |
| Esm | N - P | -0.14 | -0.16 | -0.13 | 0 |
| El | N - P | 0.02 | 0.01 | 0.04 | 0.47 |
| Et | N - P | -0.12 | -0.14 | -0.1 | 0.01 |
| Pa | N - P | -0.05 | -0.09 | -0.01 | 0.54 |
| Pb | N - P | 0.02 | -0.03 | 0.07 | 0.82 |
| Pt | N - P | 0.03 | -0.02 | 0.08 | 0.77 |
| Pc | N - P | 0.03 | -0.01 | 0.07 | 0.67 |
| Ca | N - P | 10.93 | 9.41 | 12.45 | 0 |
| F | N - P | 0.03 | 0.01 | 0.04 | 0.49 |
| Br | N - P | -0.1 | -0.12 | -0.08 | 0.01 |
| Ra | N - P | -0.24 | -0.46 | -0.02 | 0.58 |
| Ns | N - P | -0.13 | -0.2 | -0.07 | 0.31 |
| an | N - P | -0.13 | -0.2 | -0.07 | 0.31 |
| ac | N - P | -0.12 | -0.2 | -0.04 | 0.46 |
| am | N - P | -0.11 | -0.18 | -0.04 | 0.42 |
| V0 | N - P | 20.85 | 17.71 | 23.98 | 0 |

| | | | | | |
|---|---|---|---|---|---|
| Nm | N - P | -0.06 | -0.1 | -0.03 | 0.4 |
| Nl | N - P | -0.14 | -0.24 | -0.04 | 0.48 |
| Hgm | N - P | -0.14 | -0.28 | 0 | 0.61 |
| ASAD | N - P | 26.52 | 21.83 | 31.21 | 0 |
| ASAN | N - P | 15.49 | 12.6 | 18.38 | 0.01 |
| ΔASA | N - P | 11.09 | 7.28 | 14.9 | 0.15 |
| ΔGh | N - P | -1.64 | -1.88 | -1.41 | 0 |
| GhD | N - P | -4.06 | -4.66 | -3.46 | 0 |
| GhN | N - P | -1.55 | -1.81 | -1.28 | 0 |
| ΔHh | N - P | -2.67 | -2.99 | -2.35 | 0 |
| TΔSh | N - P | 1.03 | 0.88 | 1.17 | 0 |
| ΔCh | N - P | 4.07 | 2.55 | 5.6 | 0.18 |
| ΔGc | N - P | 1.72 | 1.49 | 1.94 | 0 |
| ΔHc | N - P | 1.98 | 1.62 | 2.35 | 0.01 |
| Sc | N - P | -0.28 | -0.54 | -0.01 | 0.6 |
| ΔG | N - P | 0.07 | -0.01 | 0.15 | 0.67 |
| ΔH | N - P | -0.69 | -1.01 | -0.37 | 0.29 |
| TΔS | N - P | 0.76 | 0.51 | 1.01 | 0.13 |
| v | N - P | 0.98 | 0.77 | 1.19 | 0.02 |
| s | N - P | 0.98 | 0.77 | 1.19 | 0.02 |
| ff | N - P | 0.82 | 0.63 | 1 | 0.02 |

**Table S25.** CYP19A1: ANOVA results for change in 48 properties (N: 'Non-pathogenic', P: 'Pathogenic', DIFF: Difference between means, LWR: Lower end of 95% confidence interval, UPR: Upper end of 95% confidence interval).

| Name | Group Pair | DIFF | LWR | UPR | p-value |
|---|---|---|---|---|---|
| K | N - P | 3.57 | 3.18 | 3.95 | 0.03 |
| Ht | N - P | -0.53 | -0.67 | -0.39 | 0.37 |
| Hs | N - P | -0.3 | -0.44 | -0.15 | 0.63 |
| P | N - P | 22.83 | 20.32 | 25.33 | 0.03 |
| pHi | N - P | 2.43 | 2.19 | 2.67 | 0.01 |
| pK | N - P | -0.08 | -0.12 | -0.04 | 0.65 |
| Mw | N - P | 13.54 | 10.08 | 17 | 0.36 |
| B1 | N - P | -0.68 | -1.23 | -0.12 | 0.77 |
| Rf | N - P | -2.89 | -3.45 | -2.33 | 0.22 |
| u | N - P | -0.7 | -1.73 | 0.34 | 0.87 |
| Hnc | N - P | -1.43 | -1.56 | -1.3 | 0.01 |
| Esm | N - P | -0.15 | -0.16 | -0.13 | 0.05 |
| El | N - P | 0 | -0.01 | 0.02 | 0.95 |
| Et | N - P | -0.14 | -0.16 | -0.11 | 0.14 |
| Pa | N - P | 0.11 | 0.07 | 0.14 | 0.47 |
| Pb | N - P | -0.17 | -0.21 | -0.13 | 0.34 |
| Pt | N - P | -0.11 | -0.16 | -0.07 | 0.52 |
| Pc | N - P | -0.01 | -0.05 | 0.02 | 0.94 |
| Ca | N - P | 9.61 | 8.15 | 11.07 | 0.12 |
| F | N - P | 0.05 | 0.03 | 0.07 | 0.49 |

| | | | | | |
|---|---|---|---|---|---|
| Br | N - P | -0.14 | -0.16 | -0.12 | 0.09 |
| Ra | N - P | -0.1 | -0.31 | 0.11 | 0.91 |
| Ns | N - P | -0.41 | -0.51 | -0.31 | 0.34 |
| an | N - P | 0.06 | 0 | 0.12 | 0.8 |
| ac | N - P | -0.13 | -0.19 | -0.06 | 0.67 |
| am | N - P | 0.1 | 0.03 | 0.17 | 0.73 |
| V0 | N - P | 15.12 | 12.1 | 18.15 | 0.24 |
| Nm | N - P | 0.05 | 0.01 | 0.08 | 0.76 |
| Nl | N - P | -0.31 | -0.41 | -0.22 | 0.42 |
| Hgm | N - P | -0.41 | -0.53 | -0.29 | 0.43 |
| ASAD | N - P | 21.24 | 16.68 | 25.8 | 0.27 |
| ASAN | N - P | 23.52 | 20.8 | 26.24 | 0.04 |
| ASA | N - P | -1.99 | -5.65 | 1.67 | 0.9 |
| ΔGh | N - P | -1.78 | -2.03 | -1.52 | 0.1 |
| GhD | N - P | -2.8 | -3.41 | -2.18 | 0.28 |
| GhN | N - P | -3.3 | -3.57 | -3.03 | 0 |
| ΔHh | N - P | -2.46 | -2.8 | -2.12 | 0.08 |
| TΔSh | N - P | 0.68 | 0.54 | 0.82 | 0.26 |
| ΔCh | N - P | -2.38 | -3.83 | -0.93 | 0.7 |
| ΔGc | N - P | 0.96 | 0.72 | 1.21 | 0.35 |
| ΔHc | N - P | -1.5 | -1.87 | -1.12 | 0.35 |
| Sc | N - P | 2.48 | 2.21 | 2.74 | 0.03 |
| ΔG | N - P | -0.81 | -0.9 | -0.73 | 0.02 |
| ΔH | N - P | -3.95 | -4.28 | -3.63 | 0 |
| TΔS | N - P | 3.14 | 2.89 | 3.39 | 0 |
| v | N - P | 1.03 | 0.77 | 1.29 | 0.35 |
| s | N - P | 2.49 | 2.29 | 2.7 | 0 |
| ff | N - P | 1.68 | 1.51 | 1.85 | 0.02 |

**Table S26.** CYP21A2: ANOVA results for change in 48 properties (N: ‘Non-pathogenic’, P: ‘Pathogenic’, DIFF: Difference between means, LWR: Lower end of 95% confidence interval, UPR: Upper end of 95% confidence interval).

| Name | Group Pair | DIFF | LWR | UPR | p-value |
|---|---|---|---|---|---|
| K | N - P | 0.84 | 0.46 | 1.22 | 0.15 |
| Ht | N - P | -0.36 | -0.48 | -0.23 | 0.06 |
| Hs | N - P | 0.25 | 0.12 | 0.39 | 0.21 |
| P | N - P | 0.8 | -1.6 | 3.2 | 0.83 |
| pHi | N - P | 0.75 | 0.54 | 0.95 | 0.02 |
| pK | N - P | -0.03 | -0.06 | 0 | 0.49 |
| Mw | N - P | -8.79 | -11.99 | -5.6 | 0.07 |
| B1 | N - P | -1.5 | -2.01 | -0.99 | 0.05 |
| Rf | N - P | -1.17 | -1.67 | -0.68 | 0.12 |
| u | N - P | -3.1 | -4.03 | -2.18 | 0.03 |
| Hnc | N - P | -0.23 | -0.35 | -0.11 | 0.2 |
| Esm | N - P | 0 | -0.02 | 0.01 | 0.89 |
| El | N - P | 0.02 | 0.01 | 0.03 | 0.25 |

| | | | | | |
|---|---|---|---|---|---|
| Et | N - P | 0.02 | 0 | 0.04 | 0.5 |
| Pa | N - P | -0.05 | -0.08 | -0.02 | 0.31 |
| Pb | N - P | 0.06 | 0.02 | 0.1 | 0.28 |
| Pt | N - P | -0.02 | -0.06 | 0.02 | 0.75 |
| Pc | N - P | 0.02 | -0.02 | 0.05 | 0.73 |
| Ca | N - P | -0.55 | -1.92 | 0.81 | 0.79 |
| F | N - P | 0.01 | 0 | 0.03 | 0.59 |
| Br | N - P | 0.02 | 0 | 0.03 | 0.55 |
| Ra | N - P | 0.17 | -0.02 | 0.35 | 0.55 |
| Ns | N - P | 0.15 | 0.06 | 0.24 | 0.27 |
| an | N - P | -0.01 | -0.08 | 0.05 | 0.89 |
| ac | N - P | -0.36 | -0.43 | -0.29 | 0 |
| am | N - P | 0.05 | -0.02 | 0.11 | 0.66 |
| V0 | N - P | -3.66 | -6.42 | -0.91 | 0.38 |
| Nm | N - P | -0.04 | -0.07 | 0 | 0.47 |
| Nl | N - P | 0.23 | 0.15 | 0.31 | 0.06 |
| Hgm | N - P | 0.16 | 0.05 | 0.27 | 0.32 |
| ASAD | N - P | -5.74 | -9.87 | -1.6 | 0.36 |
| ASAN | N - P | -0.85 | -3.32 | 1.63 | 0.82 |
| ΔASA | N - P | -4.69 | -7.99 | -1.39 | 0.35 |
| ΔGh | N - P | 0 | -0.23 | 0.23 | 0.99 |
| GhD | N - P | -0.93 | -1.47 | -0.39 | 0.26 |
| GhN | N - P | -0.44 | -0.69 | -0.18 | 0.26 |
| ΔHh | N - P | 0.01 | -0.3 | 0.32 | 0.98 |
| TΔSh | N - P | -0.02 | -0.14 | 0.11 | 0.93 |
| ΔCh | N - P | -0.86 | -2.1 | 0.39 | 0.65 |
| ΔGc | N - P | -0.28 | -0.5 | -0.07 | 0.39 |
| ΔHc | N - P | -1.17 | -1.49 | -0.85 | 0.02 |
| ΔSc | N - P | 0.89 | 0.67 | 1.11 | 0.01 |
| ΔG | N - P | -0.29 | -0.36 | -0.21 | 0.01 |
| ΔH | N - P | -1.15 | -1.43 | -0.88 | 0.01 |
| TΔS | N - P | 0.87 | 0.65 | 1.08 | 0.01 |
| v | N - P | -0.7 | -0.94 | -0.46 | 0.05 |
| s | N - P | 0.2 | 0.01 | 0.39 | 0.48 |
| ff | N - P | 0.22 | 0.05 | 0.39 | 0.39 |

**Table S27.** Correlation coefficient values for pathogenic mutations containing CYPs (48 properties vs SNAP2 score of non-pathogenic mutations). Correlation coefficient is represented here, more closer to +1 represents positive correlation, more closer to -1 represents negative correlation, and 0 means no correlation.

| Property | CYP1B1 | CYP2R1 | CYP3A4 | CYP8A1 | CYP11A1 | CYP11B1 | CYP11B2 | CYP17A1 | CYP19A1 | CYP21A2 |
|---|---|---|---|---|---|---|---|---|---|---|
| K | 0.08 | 0 | -0.07 | -0.06 | -0.08 | 0.08 | 0 | -0.1 | -0.03 | 0.11 |
| Ht | -0.02 | -0.11 | 0 | -0.07 | 0.03 | 0.05 | 0.05 | 0.01 | 0.11 | -0.02 |
| Hs | -0.07 | -0.07 | -0.05 | -0.02 | 0 | -0.02 | 0.01 | 0.06 | -0.02 | -0.12 |

| | | | | | | | | | | |
|---|---|---|---|---|---|---|---|---|---|---|
| P | -0.08 | 0 | 0.01 | -0.04 | -0.11 | -0.11 | -0.08 | -0.03 | 0.02 | -0.02 |
| pHi | -0.08 | 0.01 | -0.1 | -0.16 | -0.1 | -0.07 | -0.12 | -0.09 | 0 | -0.01 |
| pK | 0.04 | 0.01 | 0 | 0.05 | 0.17 | 0.11 | 0.03 | -0.07 | -0.02 | 0.02 |
| Mw | -0.12 | 0.03 | -0.02 | -0.08 | -0.1 | -0.11 | -0.09 | -0.03 | 0.1 | -0.1 |
| B1 | -0.09 | -0.07 | 0.03 | -0.04 | 0.02 | -0.01 | 0.02 | 0.02 | 0.13 | -0.09 |
| Rf | -0.03 | -0.11 | -0.03 | -0.04 | 0.03 | 0.03 | 0.01 | 0.01 | 0.03 | -0.08 |
| u | -0.05 | 0.05 | 0.01 | -0.01 | 0.02 | -0.01 | 0.03 | 0.01 | 0.12 | -0.07 |
| Hnc | 0.03 | -0.11 | 0.03 | 0.09 | 0.13 | 0.07 | 0.1 | 0.08 | -0.03 | -0.03 |
| Esm | 0.11 | -0.05 | 0.04 | 0.14 | 0.18 | 0.11 | 0.11 | 0.08 | -0.05 | 0.06 |
| El | -0.13 | -0.12 | -0.12 | -0.08 | -0.05 | -0.06 | -0.04 | -0.01 | 0 | -0.14 |
| Et | 0.01 | -0.12 | -0.05 | 0.06 | 0.1 | 0.06 | 0.06 | 0.06 | -0.04 | -0.04 |
| Pa | 0.02 | 0.07 | 0.12 | 0.07 | 0.09 | 0.03 | 0.02 | 0.09 | -0.02 | -0.06 |
| Pb | -0.12 | -0.04 | -0.07 | -0.08 | 0 | -0.06 | -0.02 | 0 | -0.01 | -0.13 |
| Pt | 0.07 | 0 | -0.04 | 0.02 | -0.03 | 0.04 | 0.03 | -0.07 | 0.01 | 0.12 |
| Pc | 0.05 | -0.04 | -0.07 | -0.02 | -0.08 | 0 | -0.01 | -0.08 | 0.02 | 0.12 |
| Ca | -0.14 | 0.03 | -0.05 | -0.14 | -0.09 | -0.11 | -0.12 | -0.05 | 0.05 | -0.12 |
| F | 0.1 | 0.08 | 0.02 | 0 | -0.05 | 0.05 | 0 | -0.07 | 0 | 0.15 |
| Br | -0.01 | -0.11 | -0.04 | 0.04 | 0.1 | 0.03 | 0.07 | 0.08 | -0.06 | -0.08 |
| Ra | -0.1 | -0.11 | -0.01 | -0.04 | 0.02 | -0.06 | 0.01 | 0.07 | -0.03 | -0.14 |
| Ns | -0.05 | -0.11 | -0.07 | 0 | 0.04 | 0.04 | 0.05 | 0.04 | 0.01 | -0.09 |
| an | -0.02 | 0.01 | 0.05 | 0.06 | 0.13 | 0 | 0.05 | 0.1 | -0.03 | -0.08 |
| ac | 0.12 | 0.03 | 0.13 | 0.12 | 0.04 | -0.11 | 0.14 | 0.03 | 0.12 | 0.09 |
| am | -0.03 | 0.09 | 0.05 | 0.04 | 0.05 | 0.05 | -0.01 | 0.05 | -0.05 | -0.04 |
| V0 | -0.13 | -0.01 | -0.05 | -0.13 | -0.09 | 0.03 | -0.1 | -0.04 | 0.08 | -0.11 |
| Nm | 0.03 | 0.1 | 0.06 | 0.06 | 0.07 | -0.03 | 0.02 | 0.1 | -0.04 | -0.06 |
| Nl | -0.08 | -0.12 | -0.11 | -0.05 | -0.02 | 0.01 | 0.01 | 0.01 | 0 | -0.08 |
| Hgm | -0.04 | -0.05 | -0.07 | 0.04 | 0.03 | 0.03 | 0.07 | 0.06 | -0.01 | -0.07 |
| ASAD | -0.12 | 0.02 | -0.01 | -0.12 | -0.09 | -0.02 | -0.11 | -0.03 | 0.08 | -0.09 |
| ASAN | -0.04 | 0.09 | 0.03 | -0.1 | -0.09 | -0.06 | -0.1 | -0.07 | 0.06 | 0.03 |
| ΔASA | -0.12 | -0.03 | -0.03 | -0.08 | -0.04 | -0.08 | -0.07 | 0.02 | 0.05 | -0.14 |
| ΔGh | 0.06 | -0.12 | 0.02 | -0.01 | 0.16 | 0.1 | 0.09 | 0.05 | -0.02 | 0.01 |
| GhD | 0.08 | -0.12 | 0.09 | 0.06 | 0.14 | 0.1 | 0.11 | 0.12 | 0.08 | 0.03 |
| GhN | 0.07 | -0.1 | 0.04 | 0.04 | 0.22 | 0.13 | 0.12 | 0.06 | 0.01 | 0.03 |
| ΔHh | 0.11 | -0.07 | 0.06 | 0.04 | 0.18 | 0.13 | 0.12 | 0.06 | -0.04 | 0.06 |
| TΔSh | -0.15 | -0.04 | -0.09 | -0.11 | -0.13 | -0.13 | -0.13 | -0.04 | 0.05 | -0.12 |
| ΔCh | -0.11 | -0.15 | -0.08 | -0.12 | -0.01 | -0.04 | -0.06 | -0.01 | 0.04 | -0.11 |
| ΔGc | -0.07 | 0.12 | -0.03 | 0.01 | -0.13 | -0.1 | -0.07 | -0.06 | 0.03 | -0.02 |
| ΔHc | -0.09 | 0.01 | -0.03 | 0.03 | -0.02 | -0.08 | -0.06 | -0.03 | 0.05 | -0.07 |
| ΔSc | 0.06 | 0.09 | 0.01 | -0.02 | -0.09 | 0 | 0.01 | 0 | -0.05 | 0.08 |
| ΔG | 0 | -0.05 | -0.01 | 0.01 | 0.12 | 0.03 | 0.06 | -0.01 | 0.02 | -0.04 |
| ΔH | 0.03 | -0.06 | 0.03 | 0.07 | 0.16 | 0.05 | 0.07 | 0.02 | 0.02 | -0.02 |
| TΔS | -0.04 | 0.07 | -0.04 | -0.09 | -0.16 | -0.06 | -0.07 | -0.03 | -0.02 | 0.01 |
| v | -0.12 | 0.02 | -0.03 | -0.08 | -0.1 | -0.11 | -0.07 | -0.04 | 0.1 | -0.09 |
| s | -0.1 | 0.06 | -0.04 | -0.05 | -0.21 | -0.14 | -0.08 | -0.02 | 0 | -0.09 |
| ff | -0.11 | 0.08 | -0.01 | -0.11 | -0.13 | -0.13 | -0.15 | -0.02 | -0.01 | -0.09 |

**Table S28.** p-values for pathogenic mutations containing CYPs (48 properties vs SNAP2 score of non-pathogenic mutations).

| Property | CYP1B1 | CYP2R1 | CYP3A4 | CYP8A1 | CYP11A1 | CYP11B1 | CYP11B2 | CYP17A1 | CYP19A1 | CYP21A2 |
|---|---|---|---|---|---|---|---|---|---|---|
| K | 0.05 | 0.96 | 0.15 | 0.23 | 0.1 | 0.07 | 0.91 | 0.05 | 0.54 | 0.02 |
| Ht | 0.68 | 0.03 | 0.99 | 0.16 | 0.49 | 0.29 | 0.28 | 0.88 | 0.02 | 0.7 |
| Hs | 0.09 | 0.19 | 0.31 | 0.71 | 0.99 | 0.57 | 0.74 | 0.23 | 0.69 | 0.01 |
| P | 0.05 | 0.93 | 0.85 | 0.44 | 0.03 | 0.01 | 0.06 | 0.54 | 0.67 | 0.67 |
| pHi | 0.04 | 0.85 | 0.04 | 0 | 0.06 | 0.1 | 0.01 | 0.08 | 0.97 | 0.89 |
| pK | 0.29 | 0.92 | 0.94 | 0.26 | 0 | 0.01 | 0.45 | 0.22 | 0.67 | 0.6 |
| Mw | 0 | 0.56 | 0.73 | 0.09 | 0.05 | 0.01 | 0.05 | 0.54 | 0.04 | 0.04 |
| B1 | 0.03 | 0.16 | 0.6 | 0.37 | 0.67 | 0.75 | 0.65 | 0.64 | 0.01 | 0.05 |
| Rf | 0.43 | 0.03 | 0.51 | 0.34 | 0.5 | 0.51 | 0.91 | 0.87 | 0.57 | 0.1 |
| u | 0.22 | 0.34 | 0.81 | 0.81 | 0.71 | 0.75 | 0.49 | 0.8 | 0.02 | 0.15 |
| Hnc | 0.4 | 0.03 | 0.52 | 0.05 | 0.01 | 0.1 | 0.03 | 0.16 | 0.54 | 0.47 |
| Esm | 0.01 | 0.36 | 0.38 | 0 | 0 | 0.01 | 0.01 | 0.12 | 0.29 | 0.23 |
| El | 0 | 0.02 | 0.01 | 0.09 | 0.28 | 0.16 | 0.38 | 0.86 | 0.96 | 0 |
| Et | 0.83 | 0.02 | 0.3 | 0.16 | 0.05 | 0.2 | 0.17 | 0.29 | 0.39 | 0.34 |
| Pa | 0.7 | 0.15 | 0.02 | 0.14 | 0.08 | 0.45 | 0.68 | 0.1 | 0.69 | 0.22 |
| Pb | 0 | 0.4 | 0.13 | 0.08 | 0.93 | 0.18 | 0.65 | 0.99 | 0.83 | 0 |
| Pt | 0.08 | 0.93 | 0.43 | 0.66 | 0.54 | 0.42 | 0.58 | 0.22 | 0.88 | 0.01 |
| Pc | 0.2 | 0.42 | 0.18 | 0.69 | 0.11 | 0.95 | 0.87 | 0.14 | 0.64 | 0.01 |
| Ca | 0 | 0.58 | 0.34 | 0 | 0.06 | 0.01 | 0.01 | 0.33 | 0.28 | 0.01 |
| F | 0.01 | 0.14 | 0.64 | 0.93 | 0.37 | 0.22 | 0.97 | 0.19 | 0.93 | 0 |
| Br | 0.82 | 0.04 | 0.44 | 0.34 | 0.05 | 0.49 | 0.14 | 0.14 | 0.26 | 0.11 |
| Ra | 0.02 | 0.03 | 0.8 | 0.45 | 0.75 | 0.48 | 0.8 | 0.19 | 0.58 | 0 |
| Ns | 0.19 | 0.03 | 0.18 | 0.97 | 0.41 | 0.75 | 0.26 | 0.45 | 0.88 | 0.05 |
| an | 0.58 | 0.85 | 0.29 | 0.17 | 0.01 | 0.52 | 0.27 | 0.05 | 0.55 | 0.09 |
| ac | 0 | 0.63 | 0.01 | 0.01 | 0.38 | 0.01 | 0 | 0.55 | 0.01 | 0.06 |
| am | 0.52 | 0.09 | 0.29 | 0.42 | 0.3 | 0.51 | 0.77 | 0.37 | 0.29 | 0.36 |
| V0 | 0 | 0.9 | 0.36 | 0.01 | 0.06 | 0.02 | 0.02 | 0.46 | 0.13 | 0.02 |
| Nm | 0.54 | 0.05 | 0.19 | 0.16 | 0.18 | 0.85 | 0.73 | 0.07 | 0.39 | 0.19 |
| Nl | 0.05 | 0.02 | 0.02 | 0.3 | 0.66 | 0.54 | 0.86 | 0.8 | 0.93 | 0.09 |
| Hgm | 0.32 | 0.34 | 0.19 | 0.44 | 0.53 | 0.57 | 0.11 | 0.25 | 0.8 | 0.15 |
| ASAD | 0 | 0.64 | 0.82 | 0.01 | 0.09 | 0.02 | 0.01 | 0.6 | 0.11 | 0.04 |
| ASAN | 0.34 | 0.09 | 0.57 | 0.03 | 0.06 | 0.15 | 0.03 | 0.19 | 0.22 | 0.52 |
| ΔASA | 0 | 0.57 | 0.48 | 0.1 | 0.44 | 0.07 | 0.15 | 0.76 | 0.29 | 0 |
| ΔGh | 0.12 | 0.02 | 0.63 | 0.84 | 0 | 0.02 | 0.05 | 0.34 | 0.71 | 0.84 |
| GhD | 0.04 | 0.02 | 0.07 | 0.19 | 0 | 0.03 | 0.05 | 0.02 | 0.1 | 0.59 |
| GhN | 0.07 | 0.05 | 0.48 | 0.42 | 0 | 0 | 0.01 | 0.28 | 0.78 | 0.55 |
| ΔHh | 0.01 | 0.16 | 0.25 | 0.36 | 0 | 0 | 0.01 | 0.27 | 0.07 | 0.23 |
| TΔSh | 0 | 0.41 | 0.06 | 0.01 | 0.01 | 0 | 0 | 0.42 | 0.3 | 0.01 |

| | | | | | | | | | | |
|---|---|---|---|---|---|---|---|---|---|---|
| ΔCh | 0.01 | 0.01 | 0.12 | 0.01 | 0.89 | 0.32 | 0.2 | 0.8 | 0.4 | 0.02 |
| ΔGc | 0.09 | 0.03 | 0.59 | 0.77 | 0.01 | 0.02 | 0.1 | 0.3 | 0.6 | 0.61 |
| ΔHc | 0.04 | 0.81 | 0.6 | 0.59 | 0.71 | 0.08 | 0.21 | 0.55 | 0.3 | 0.14 |
| ΔSc | 0.17 | 0.1 | 0.81 | 0.64 | 0.09 | 0.91 | 0.9 | 0.94 | 0.3 | 0.09 |
| ΔG | 0.98 | 0.35 | 0.91 | 0.82 | 0.02 | 0.54 | 0.22 | 0.89 | 0.67 | 0.37 |
| ΔH | 0.5 | 0.22 | 0.52 | 0.14 | 0 | 0.21 | 0.15 | 0.67 | 0.62 | 0.67 |
| TΔS | 0.37 | 0.2 | 0.37 | 0.06 | 0 | 0.16 | 0.14 | 0.55 | 0.62 | 0.81 |
| v | 0 | 0.75 | 0.6 | 0.08 | 0.04 | 0.01 | 0.1 | 0.46 | 0.04 | 0.04 |
| s | 0.02 | 0.28 | 0.41 | 0.28 | 0 | 0 | 0.07 | 0.75 | 0.95 | 0.04 |
| ff | 0.01 | 0.11 | 0.81 | 0.02 | 0.01 | 0 | 0 | 0.7 | 0.83 | 0.05 |

**Table S29.** Correlation coefficient values for pathogenic mutations containing CYPs (48 properties vs SNAP2 score of pathogenic mutations). Correlation coefficient is represented here, more closer to +1 represents positive correlation, more closer to -1 represents negative correlation, and 0 means no correlation.

| Property | CYP1B1 | CYP2R1 | CYP3A4 | CYP8A1 | CYP11A1 | CYP11B1 | CYP11B2 | CYP17A1 | CYP19A1 | CYP21A2 |
|---|---|---|---|---|---|---|---|---|---|---|
| K | -0.11 | 0.53 | nan | nan | 0.04 | -0.01 | -0.62 | 0.08 | -0.7 | 0.01 |
| Ht | 0.13 | 0.2 | nan | nan | 0.2 | -0.02 | 0.83 | 0.25 | -0.08 | 0.19 |
| Hs | 0.03 | -0.54 | nan | nan | -0.19 | -0.23 | 0.62 | 0.15 | 0.38 | -0.03 |
| P | 0.01 | -0.43 | nan | nan | -0.15 | 0.13 | -0.3 | -0.15 | -0.67 | -0.12 |
| pHi | -0.11 | -0.42 | nan | nan | -0.25 | -0.04 | -0.19 | 0.15 | -0.67 | -0.2 |
| pK | 0.15 | -0.2 | nan | nan | 0.45 | -0.19 | 0.24 | 0.01 | 0.24 | -0.01 |
| Mw | 0.07 | -0.43 | nan | nan | 0.3 | 0.04 | 0.73 | 0.1 | -0.12 | 0.09 |
| B1 | 0.24 | -0.21 | nan | nan | -0.39 | -0.22 | 0.82 | 0.23 | -0.07 | 0.12 |
| Rf | 0.04 | 0.05 | nan | nan | -0.01 | -0.12 | 0.81 | 0.21 | 0.15 | 0.11 |
| u | 0.16 | -0.45 | nan | nan | 0.69 | 0.05 | 0.61 | 0.11 | 0.08 | 0.16 |
| Hnc | 0.08 | 0.29 | nan | nan | 0.17 | -0.01 | 0.5 | -0.04 | 0.5 | 0.1 |
| Esm | 0.05 | 0.35 | nan | nan | 0.31 | 0.05 | -0.47 | -0.09 | 0.21 | -0.03 |
| El | -0.04 | -0.54 | nan | nan | -0.16 | -0.2 | 0.81 | 0.22 | 0.36 | 0 |
| Et | 0.01 | -0.12 | nan | nan | 0.02 | -0.07 | 0.58 | 0.08 | 0.28 | -0.03 |
| Pa | 0.18 | -0.53 | nan | nan | 0.66 | -0.05 | 0.14 | -0.23 | 0.21 | -0.01 |
| Pb | 0.04 | -0.54 | nan | nan | -0.35 | -0.31 | 0.71 | 0.16 | 0.14 | -0.01 |
| Pt | -0.1 | 0.53 | nan | nan | 0.14 | 0.2 | -0.41 | -0.05 | -0.17 | 0.06 |
| Pc | -0.16 | 0.54 | nan | nan | -0.22 | 0.18 | -0.47 | 0.06 | -0.35 | 0.01 |
| Ca | 0.01 | -0.49 | nan | nan | 0.12 | -0.04 | 0.73 | 0.13 | -0.35 | -0.02 |
| F | -0.11 | 0.53 | nan | nan | 0.01 | 0.05 | -0.69 | -0.06 | -0.01 | 0.03 |
| Br | -0.01 | -0.37 | nan | nan | 0.03 | -0.03 | 0.55 | 0.06 | 0.21 | -0.04 |
| Ra | 0.07 | -0.53 | nan | nan | -0.19 | -0.24 | 0.74 | 0.07 | 0.49 | -0.02 |
| Ns | 0.03 | -0.52 | nan | nan | -0.31 | -0.24 | 0.73 | 0.15 | 0.33 | -0.02 |
| an | 0.17 | -0.54 | nan | nan | 0.39 | -0.24 | 0.27 | -0.06 | 0.1 | -0.11 |
| ac | 0.16 | 0.48 | nan | nan | 0.4 | 0.07 | -0.25 | -0.13 | 0.76 | 0.39 |
| am | 0.18 | -0.52 | nan | nan | -0.23 | 0.14 | 0.01 | -0.16 | -0.69 | -0.15 |
| V0 | 0.03 | -0.44 | nan | nan | -0.03 | -0.05 | 0.75 | 0.19 | -0.29 | 0.01 |
| Nm | 0.05 | -0.54 | nan | nan | 0.69 | 0.07 | 0.04 | -0.23 | -0.02 | -0.12 |

| | | | | | | | | | | |
|---|---|---|---|---|---|---|---|---|---|---|
| Nl | -0.03 | -0.47 | nan | nan | -0.35 | -0.26 | 0.73 | 0.23 | 0.27 | -0.01 |
| Hgm | 0.11 | -0.52 | nan | nan | 0.11 | -0.18 | 0.62 | 0.11 | 0.11 | 0.01 |
| ASAD | 0.05 | -0.42 | nan | nan | -0.13 | 0 | 0.74 | 0.16 | -0.32 | 0.01 |
| ASAN | -0.01 | -0.06 | nan | nan | -0.1 | 0.05 | -0.4 | -0.02 | -0.41 | -0.04 |
| ΔASA | 0.06 | -0.49 | nan | nan | 0 | -0.03 | 0.79 | 0.19 | 0.05 | 0.04 |
| ΔGh | -0.03 | 0.43 | nan | nan | -0.22 | 0 | 0.55 | 0.02 | -0.05 | -0.02 |
| GhD | 0.07 | 0.44 | nan | nan | 0.07 | -0.08 | 0.39 | 0.08 | 0.54 | 0.08 |
| GhN | 0.1 | 0.44 | nan | nan | 0.25 | 0.04 | 0.55 | 0.04 | 0.09 | 0.07 |
| ΔHh | -0.04 | 0.48 | nan | nan | -0.13 | 0.05 | -0.41 | -0.11 | 0.08 | -0.01 |
| TΔSh | 0.03 | -0.47 | nan | nan | -0.18 | -0.11 | 0.76 | 0.27 | -0.25 | -0.02 |
| ΔCh | 0.03 | -0.31 | nan | nan | -0.17 | -0.12 | 0.81 | 0.28 | -0.41 | -0.01 |
| ΔGc | 0.05 | -0.42 | nan | nan | 0.48 | 0.12 | 0.05 | 0.02 | 0.19 | 0.09 |
| ΔHc | 0.13 | -0.5 | nan | nan | 0.56 | 0.19 | 0.65 | 0.14 | 0.57 | 0.2 |
| ΔSc | -0.14 | 0.24 | nan | nan | -0.37 | -0.17 | -0.76 | -0.16 | -0.51 | -0.17 |
| ΔG | 0.06 | 0.44 | nan | nan | 0.56 | 0.29 | 0.64 | 0.06 | 0.35 | 0.18 |
| ΔH | 0.12 | 0.42 | nan | nan | 0.51 | 0.24 | 0.59 | 0.06 | 0.42 | 0.18 |
| TΔS | -0.14 | -0.42 | nan | nan | -0.49 | -0.22 | -0.55 | -0.06 | -0.44 | -0.17 |
| v | 0.07 | -0.42 | nan | nan | 0.38 | 0 | 0.69 | 0.12 | -0.13 | 0.11 |
| s | -0.12 | -0.51 | nan | nan | -0.38 | -0.12 | 0.17 | -0.02 | 0.09 | -0.07 |
| ff | -0.09 | -0.51 | nan | nan | -0.25 | 0.01 | 0.33 | -0.03 | -0.38 | -0.15 |

**Table S30.** p-values for pathogenic mutations containing CYPs (48 properties vs SNAP2 score of pathogenic mutations).

| Property | CYP1B1 | CYP2R1 | CYP3A4 | CYP8A1 | CYP11A1 | CYP11B1 | CYP11B2 | CYP17A1 | CYP19A1 | CYP21A2 |
|---|---|---|---|---|---|---|---|---|---|---|
| K | 0.49 | 0.28 | Nan | Nan | 0.92 | 0.93 | 0.1 | 0.68 | 0.12 | 0.94 |
| Ht | 0.39 | 0.71 | Nan | Nan | 0.64 | 0.87 | 0.01 | 0.22 | 0.88 | 0.12 |
| Hs | 0.84 | 0.27 | Nan | Nan | 0.65 | 0.1 | 0.1 | 0.46 | 0.45 | 0.78 |
| P | 0.96 | 0.39 | Nan | Nan | 0.71 | 0.37 | 0.47 | 0.48 | 0.14 | 0.35 |
| pHi | 0.47 | 0.41 | Nan | Nan | 0.55 | 0.8 | 0.66 | 0.45 | 0.15 | 0.1 |
| pK | 0.32 | 0.71 | Nan | Nan | 0.26 | 0.19 | 0.57 | 0.98 | 0.65 | 0.97 |
| Mw | 0.65 | 0.39 | Nan | Nan | 0.47 | 0.8 | 0.04 | 0.61 | 0.82 | 0.45 |
| B1 | 0.12 | 0.68 | Nan | Nan | 0.34 | 0.12 | 0.01 | 0.27 | 0.9 | 0.32 |
| Rf | 0.79 | 0.92 | Nan | Nan | 0.98 | 0.4 | 0.02 | 0.3 | 0.77 | 0.36 |
| u | 0.29 | 0.37 | Nan | Nan | 0.06 | 0.7 | 0.11 | 0.59 | 0.88 | 0.19 |
| Hnc | 0.59 | 0.58 | Nan | Nan | 0.69 | 0.95 | 0.2 | 0.85 | 0.31 | 0.41 |
| Esm | 0.75 | 0.5 | Nan | Nan | 0.46 | 0.72 | 0.24 | 0.66 | 0.69 | 0.78 |
| El | 0.78 | 0.27 | Nan | Nan | 0.71 | 0.17 | 0.01 | 0.29 | 0.48 | 0.99 |
| Et | 0.97 | 0.82 | Nan | Nan | 0.95 | 0.63 | 0.13 | 0.71 | 0.59 | 0.83 |
| Pa | 0.25 | 0.28 | Nan | Nan | 0.07 | 0.71 | 0.75 | 0.26 | 0.69 | 0.94 |
| Pb | 0.79 | 0.27 | Nan | Nan | 0.4 | 0.03 | 0.05 | 0.43 | 0.79 | 0.93 |
| Pt | 0.51 | 0.28 | Nan | Nan | 0.73 | 0.16 | 0.31 | 0.82 | 0.75 | 0.61 |
| Pc | 0.29 | 0.27 | Nan | Nan | 0.6 | 0.21 | 0.24 | 0.77 | 0.49 | 0.96 |
| Ca | 0.92 | 0.33 | Nan | Nan | 0.79 | 0.76 | 0.04 | 0.53 | 0.49 | 0.9 |

| | | | | | | | | | | |
|---|---|---|---|---|---|---|---|---|---|---|
| F | 0.47 | 0.28 | Nan | Nan | 0.98 | 0.7 | 0.06 | 0.78 | 0.99 | 0.8 |
| Br | 0.97 | 0.48 | Nan | Nan | 0.95 | 0.82 | 0.15 | 0.78 | 0.69 | 0.76 |
| Ra | 0.67 | 0.28 | Nan | Nan | 0.65 | 0.09 | 0.03 | 0.74 | 0.33 | 0.87 |
| Ns | 0.83 | 0.29 | Nan | Nan | 0.46 | 0.1 | 0.04 | 0.47 | 0.52 | 0.85 |
| an | 0.28 | 0.27 | Nan | Nan | 0.34 | 0.09 | 0.52 | 0.76 | 0.85 | 0.37 |
| ac | 0.29 | 0.34 | Nan | Nan | 0.32 | 0.65 | 0.55 | 0.54 | 0.08 | 0 |
| am | 0.23 | 0.29 | Nan | Nan | 0.59 | 0.31 | 0.98 | 0.45 | 0.13 | 0.21 |
| V0 | 0.86 | 0.39 | Nan | Nan | 0.94 | 0.72 | 0.03 | 0.36 | 0.58 | 0.94 |
| Nm | 0.73 | 0.27 | Nan | Nan | 0.06 | 0.62 | 0.93 | 0.27 | 0.97 | 0.35 |
| Nl | 0.85 | 0.35 | Nan | Nan | 0.39 | 0.06 | 0.04 | 0.25 | 0.61 | 0.92 |
| Hgm | 0.48 | 0.29 | Nan | Nan | 0.79 | 0.21 | 0.1 | 0.6 | 0.84 | 0.91 |
| ASAD | 0.75 | 0.4 | Nan | Nan | 0.75 | 0.98 | 0.04 | 0.43 | 0.54 | 0.93 |
| ASAN | 0.97 | 0.91 | Nan | Nan | 0.81 | 0.73 | 0.33 | 0.9 | 0.41 | 0.77 |
| ΔASA | 0.68 | 0.33 | Nan | Nan | 1 | 0.83 | 0.02 | 0.35 | 0.93 | 0.73 |
| ΔGh | 0.84 | 0.4 | Nan | Nan | 0.6 | 1 | 0.16 | 0.93 | 0.93 | 0.88 |
| GhD | 0.63 | 0.39 | Nan | Nan | 0.87 | 0.6 | 0.34 | 0.7 | 0.27 | 0.51 |
| GhN | 0.53 | 0.38 | Nan | Nan | 0.56 | 0.8 | 0.16 | 0.86 | 0.87 | 0.57 |
| ΔHh | 0.82 | 0.34 | Nan | Nan | 0.76 | 0.71 | 0.31 | 0.58 | 0.87 | 0.97 |
| TΔSh | 0.85 | 0.35 | Nan | Nan | 0.67 | 0.43 | 0.03 | 0.19 | 0.63 | 0.86 |
| ΔCh | 0.84 | 0.55 | Nan | Nan | 0.68 | 0.4 | 0.01 | 0.17 | 0.41 | 0.94 |
| ΔGc | 0.72 | 0.41 | Nan | Nan | 0.23 | 0.42 | 0.9 | 0.92 | 0.72 | 0.47 |
| ΔHc | 0.4 | 0.32 | Nan | Nan | 0.15 | 0.18 | 0.08 | 0.5 | 0.24 | 0.1 |
| ΔSc | 0.37 | 0.65 | Nan | Nan | 0.37 | 0.24 | 0.03 | 0.45 | 0.3 | 0.16 |
| ΔG | 0.72 | 0.38 | Nan | Nan | 0.14 | 0.04 | 0.09 | 0.77 | 0.49 | 0.13 |
| ΔH | 0.44 | 0.4 | Nan | Nan | 0.19 | 0.08 | 0.12 | 0.78 | 0.4 | 0.15 |
| TΔS | 0.38 | 0.41 | Nan | Nan | 0.22 | 0.11 | 0.15 | 0.79 | 0.38 | 0.16 |
| v | 0.65 | 0.41 | Nan | Nan | 0.36 | 0.99 | 0.06 | 0.57 | 0.8 | 0.38 |
| s | 0.43 | 0.31 | Nan | Nan | 0.35 | 0.4 | 0.69 | 0.91 | 0.86 | 0.56 |
| ff | 0.58 | 0.31 | Nan | Nan | 0.55 | 0.95 | 0.43 | 0.87 | 0.46 | 0.23 |

**Table S31.** Changes in properties that indicate significant differences between Non-pathogenic and pathogenic mutations.

| **CYP450** | **Change in amino acid properties** |
|---|---|
| CYP1B1 | Ht, ΔSc, ΔG, ΔH. |
| CYP2R1 | P, pHi, pK, Mw, Rf, u, Hnc, Esm, Et, Pc, Ca, Br, am, V0, ASAD, ASAN, ΔGh, GhD, GhN, ΔHh, TΔSh, ΔGc, ΔHc, ΔH, TΔS, v, s, ff. |
| CYP3A4 | Ht, Hs, Ra, Hgm. |
| CYP8A1 | - |
| CYP11A1 | Ht, Mw, am, ΔHc, ΔG, v. |
| CYP11B1 | pHi, Mw, Hnc, Esm, Ca, V0, ASAD, ASAN, GhN, ΔHh, TΔSh, v. |
| CYP11B2 | K, Ht, u, Hgm, ΔG. |

| | |
|---|---|
| CYP17A1 | pHi, Mw, Hnc, Esm, Et, Ca, Br, V0, ASAD, ASAN, ΔGh, GhD, GhN, ΔHh, TΔSh, ΔGc, ΔHc, v, s, ff. |
| CYP19A1 | K, P, pHi, Hnc, Esm, ASAN, GhN, ΔSc, ΔG, ΔH, TΔS, s, ff. |
| CYP21A2 | pHi, B1, u, ac, ΔHc, ΔSc, ΔG, ΔH, TΔS,v. |

**Table S32.** CYPs and conservation score of their wild-type residues of pathogenic mutations**.**

| CYP1B1 | | CYP2R1 | | CYP3A4 | | CYP8A1 | | CYP11A1 | |
|---|---|---|---|---|---|---|---|---|---|
| **Mutation** | **CS** | **Mutation** | **CS** | **Mutation** | **CS** | **Mutation** | **CS** | **Mutation** | **CS** |
| A115P | 0.21 | G42R | 0.43 | I301T | 0.27 | R275Q | 0.48 | A189V | 0.12 |
| A330F | 0.61 | P43R | 0.51 | | | | | A359V | 0.26 |
| A388T | 0.38 | P44R | 0.23 | | | | | E314K | 0.31 |
| A443G | 0.16 | G45R | 0.22 | | | | | L141W | 0.22 |
| D192V | 0.4 | L46R | 0.29 | | | | | L222P | 0.29 |
| D374N | 0.54 | L99P | 0.26 | | | | | R353W | 0.36 |
| D530G | 0.2 | | | | | | | R451W | 0.37 |
| E229K | 0.17 | | | | | | | V415E | 0.38 |
| E387K | 0.82 | | | | | | | | |
| E499G | 0.08 | | | | | | | | |
| F445C | 0.63 | | | | | | | | |
| G61E | 0.59 | | | | | | | | |
| G232R | 0.41 | | | | | | | | |
| G365W | 0.3 | | | | | | | | |
| G466D | 0.7 | | | | | | | | |
| I399S | 0.42 | | | | | | | | |
| L77P | 0.35 | | | | | | | | |
| L345F | 0.15 | | | | | | | | |
| M1T | 0.52 | | | | | | | | |
| M132R | 0.26 | | | | | | | | |
| N203S | 0.69 | | | | | | | | |
| N423Y | 0.54 | | | | | | | | |
| P193L | 0.41 | | | | | | | | |
| P437L | 0.56 | | | | | | | | |
| Q144R | 0.3 | | | | | | | | |
| Q144P | 0.3 | | | | | | | | |
| R44Q | 0.21 | | | | | | | | |
| R145W | 0.61 | | | | | | | | |
| R368H | 0.56 | | | | | | | | |
| R390C | 0.74 | | | | | | | | |
| R390H | 0.74 | | | | | | | | |
| R390S | 0.74 | | | | | | | | |
| R469W | 0.06 | | | | | | | | |
| R523T | 0.58 | | | | | | | | |
| S28W | 0.34 | | | | | | | | |
| S215I | 0.18 | | | | | | | | |

| | | | | | | | | | |
|---|---|---|---|---|---|---|---|---|---|
| S239R | 0.22 | | | | | | | | |
| S515L | 0.02 | | | | | | | | |
| V198I | 0.13 | | | | | | | | |
| V320L | 0.41 | | | | | | | | |
| V364M | 0.33 | | | | | | | | |
| V409F | 0.35 | | | | | | | | |
| W57C | 0.36 | | | | | | | | |
| Y81N | 0.58 | | | | | | | | |
| **CYP11B1** | | **CYP11B2** | | **CYP17A1** | | **CYP19A1** | | **CYP21A2** | |
| **Mutation** | **CS** | **Mutation** | **CS** | **Mutation** | **CS** | **Mutation** | **CS** | **Mutation** | **CS** |
| A165D | 0.33 | E198D | 0.58 | A174E | 0.2 | C437Y | 0.91 | A15T | 0.22 |
| A306V | 0.48 | L461P | 0.44 | D116V | 0.16 | E210K | 0.19 | A362V | 0.29 |
| A331V | 0.58 | R181W | 0.26 | F93C | 0.63 | R192H | 0.3 | C169R | 0.43 |
| A368D | 0.43 | S150W | 0.37 | F114V | 0.34 | R365Q | 0.85 | C169Y | 0.43 |
| E310K | 0.6 | T185I | 0.31 | F417C | 0.56 | R375C | 0.56 | E320K | 0.4 |
| E371G | 0.84 | T318M | 0.58 | H373N | 0.6 | R435C | 0.72 | E380D | 0.13 |
| F79I | 0.65 | T498A | 0.26 | H373L | 0.6 | | | G56R | 0.64 |
| F139L | 0.15 | V386A | 0.39 | N177D | 0.55 | | | G64E | 0.38 |
| F321V | 0.47 | | | P35L | 0.74 | | | G90V | 0.28 |
| G267D | 0.52 | | | P342T | 0.58 | | | G178A | 0.45 |
| G314R | 0.55 | | | P428L | 0.24 | | | G178R | 0.45 |
| G379V | 0.37 | | | R96Q | 0.6 | | | G291R | 0.45 |
| G444D | 0.57 | | | R96W | 0.6 | | | G291C | 0.45 |
| H125R | 0.26 | | | R347C | 0.33 | | | G291S | 0.45 |
| L83S | 0.14 | | | R347H | 0.33 | | | G292D | 0.71 |
| L158P | 0.22 | | | R358Q | 0.15 | | | G424S | 0.72 |
| L299P | 0.37 | | | R362C | 0.82 | | | H62L | 0.15 |
| L489S | 0.42 | | | R440H | 0.77 | | | H282N | 0.18 |
| M88I | 0.03 | | | R496C | 0.52 | | | H365Y | 0.54 |
| N133H | 0.32 | | | R496H | 0.52 | | | I77T | 0.5 |
| P42L | 0.68 | | | S106P | 0.22 | | | I172N | 0.42 |
| P42S | 0.68 | | | W121R | 0.68 | | | I230T | 0.09 |
| P94L | 0.58 | | | W406R | 0.71 | | | I236N | 0.28 |
| P135S | 0.36 | | | W406L | 0.71 | | | K121Q | 0.28 |
| P159L | 0.32 | | | Y64S | 0.55 | | | L12M | 0.32 |
| R43Q | 0.44 | | | Y329D | 0.3 | | | L107R | 0.32 |
| R141Q | 0.69 | | | | | | | L142P | 0.11 |
| R143W | 0.21 | | | | | | | L167P | 0.22 |
| R332Q | 0.53 | | | | | | | L198F | 0.07 |
| R332G | 0.53 | | | | | | | L261P | 0.39 |
| R341S | 0.37 | | | | | | | L300F | 0.4 |
| R366C | 0.32 | | | | | | | L317M | 0.33 |
| R374Q | 0.95 | | | | | | | L363W | 0.25 |
| R384Q | 0.64 | | | | | | | M239K | 0.14 |
| R384G | 0.64 | | | | | | | M283L | 0.23 |
| R448C | 0.77 | | | | | | | P30Q | 0.65 |

| | | | | | | | | | |
|---|---|---|---|---|---|---|---|---|---|
| R448H | 0.77 | | | | | | | P30L | 0.65 |
| R453Q | 0.54 | | | | | | | P105L | 0.12 |
| R454C | 0.46 | | | | | | | P453S | 0.41 |
| R427H | 0.9 | | | | | | | P482S | 0.26 |
| S150L | 0.31 | | | | | | | R124H | 0.25 |
| T196A | 0.59 | | | | | | | R233K | 0.43 |
| T318R | 0.6 | | | | | | | R339H | 0.41 |
| T318M | 0.6 | | | | | | | R341P | 0.19 |
| T318P | 0.6 | | | | | | | R341W | 0.19 |
| T319M | 0.49 | | | | | | | R354C | 0.84 |
| T401A | 0.32 | | | | | | | R354H | 0.84 |
| V129M | 0.29 | | | | | | | R356Q | 0.21 |
| V441G | 0.34 | | | | | | | R356P | 0.21 |
| W116C | 0.68 | | | | | | | R369W | 0.14 |
| W116G | 0.68 | | | | | | | R408C | 0.77 |
| | | | | | | | | R426C | 0.78 |
| | | | | | | | | R426H | 0.78 |
| | | | | | | | | R435C | 0.36 |
| | | | | | | | | R479L | 0.36 |
| | | | | | | | | R483Q | 0.49 |
| | | | | | | | | R483P | 0.49 |
| | | | | | | | | R483W | 0.49 |
| | | | | | | | | R356W | 0.21 |
| | | | | | | | | S113F | 0.25 |
| | | | | | | | | S301Y | 0.22 |
| | | | | | | | | T450P | 0.25 |
| | | | | | | | | V211L | 0.14 |
| | | | | | | | | V237E | 0.19 |
| | | | | | | | | V281G | 0.43 |
| | | | | | | | | V281L | 0.43 |
| | | | | | | | | W302R | 0.81 |
| | | | | | | | | Y191H | 0.27 |

**Table S33.** Heme and substrate binding site residues of 10 CYPs were identified using CDD database. Pathogenic mutation causing residue positions were showing in bold.

| **CYPs** | **Heme binding site residues** | **Mutated number for heme-binding site** | **Substrate binding pocket residues** | **Mutated number for substrate-binding pocket** |
|---|---|---|---|---|
| CYP1B1 | R117, **M132**, A133, W141, **R145**, | 25 | V126, S127, S131, A133, F134, N228, | 19 |

| | | | | |
|---|---|---|---|---|
| | M152, I327, **A330**, S331, T334, L335, F394, V395, T398, I399, H401, Q424, I462, F463, S464, R468, C470, I471, G472, S476 | | F231, L264, N265, F268, D326, G329, A330, D333, T334, V395, **I399**, L509, T510 | |
| CYP2R1 | R109, L124, L125, W133, R137, F144, T191, L307, A310, G311, T314, T315, V318, V375, I379, H381, L404, P440, F441, S442, R446, C448, L449, G450, A454 | 25 | L114, F115, M118, T119, L125, N126, F214, V218, A221, A250, V253, Y254, L257, F302, G305, E306, I309, T314, V375, I379, M487, T488 | 22 |
| CYP3A4 | R105, I118, S119, W126, R130, F137, F302, A305, G306, T309, A370, L373, R375, P434, F435, G436, S437, R440, N441, C442, I443, G444, A448 | 23 | D76, R105, R106, P107, F108, S119, I120, L210, R212, F213, F215, F220, F241, **I301**, F304, A305, T309, A370, M371, R372, E374, G481, L482 | 23 |
| CYP8A1 | K121, L128, L180, A283, T284, N287, M288, A291, L349, P355, P433, W434, G435, N439, C441, L442, G443, Y446, A447, I451, L485 | 21 | F96, Y99, A100, L103, W282, A283, T284, N287, P355, F356, I357, T358, R359, G482 | 14 |
| CYP11A1 | R120, V139, L140, W147, R151, M323, G326, G327, T330, T334, L385, I390, S391, L394, R396, | 26 | R120, F121, I123, L140, M240, F241, E322, A325, G326, T330, I390, S391, | 18 |

| | | | | |
|---|---|---|---|---|
| | G454, F455, G456, W457, R460, Q461, C462, L463, G464, A468, M472 | | V392, T393, Q395, F497, L499, I500 | |
| CYP11B1 | R110, **V129**, F130, W137, **R141**, L311, **G314**, S315, **T318**, **T319**, L382, **R384**, P442, F443, **G444**, F445, **R448**, Q449, C450, G452, A456 | 21 | **W116**, F130, F231, W260, **E310**, **G314**, T318, V378, **G379**, L380, F381, L382, L407, F487, I488 | 15 |
| CYP11B2 | R110, V129, F130, W137, R141, L311, G314, S315, **T318**, T319, L382, R384, P442, F443, G444, F445, R448, Q449, C450, G452, A456 | 21 | W116, F130, F231, W260, E310, G314, **T318**, V378, G379, L380, F381, L382, L407, F487, I488 | 15 |
| CYP17A1 | L86, R96, I112, A113, **W121**, R125, D298, I299, A302, G303, T306, T307, V310, V366, L370, **H373**, P434, F435, G436, R440, S441, C442, I443, G444, A448 | 25 | A105, A113, **F114**, Y201, N202, I205, I206, L209, R239, G297, D298, G301, A302, E305, T306, V366, A367, I371, C442, V482, V483 | 21 |
| CYP19A1 | M107, R115, I132, I133, W141, R145, L152, A306, T310, M311, M364, V370, V373, **R375**, P429, F430, G431, **R435**, G436, **C437**, A438, G439, M446, M447 | 24 | R115, I133, F134, W224, I305, A306, D309, T310, V370, L372, V373, M374, L477 | 13 |

| | | | | |
|---|---|---|---|---|
| CYP21A2 | R91, S108, W116, K120, **I172**, D287, L288, **G291**, **G292**, T295, T296, T299, L353, V358, V359, **A362**, L363, **H365**, L388, A420, F421, **R426**, C428, L429, G430, L433, A434 | 27 | V100, D106, S108, L109, V197, L198, W201, **I230**, **R233**, V286, D287, I290, G291, T295, V359, **L363**, V469 | 17 |

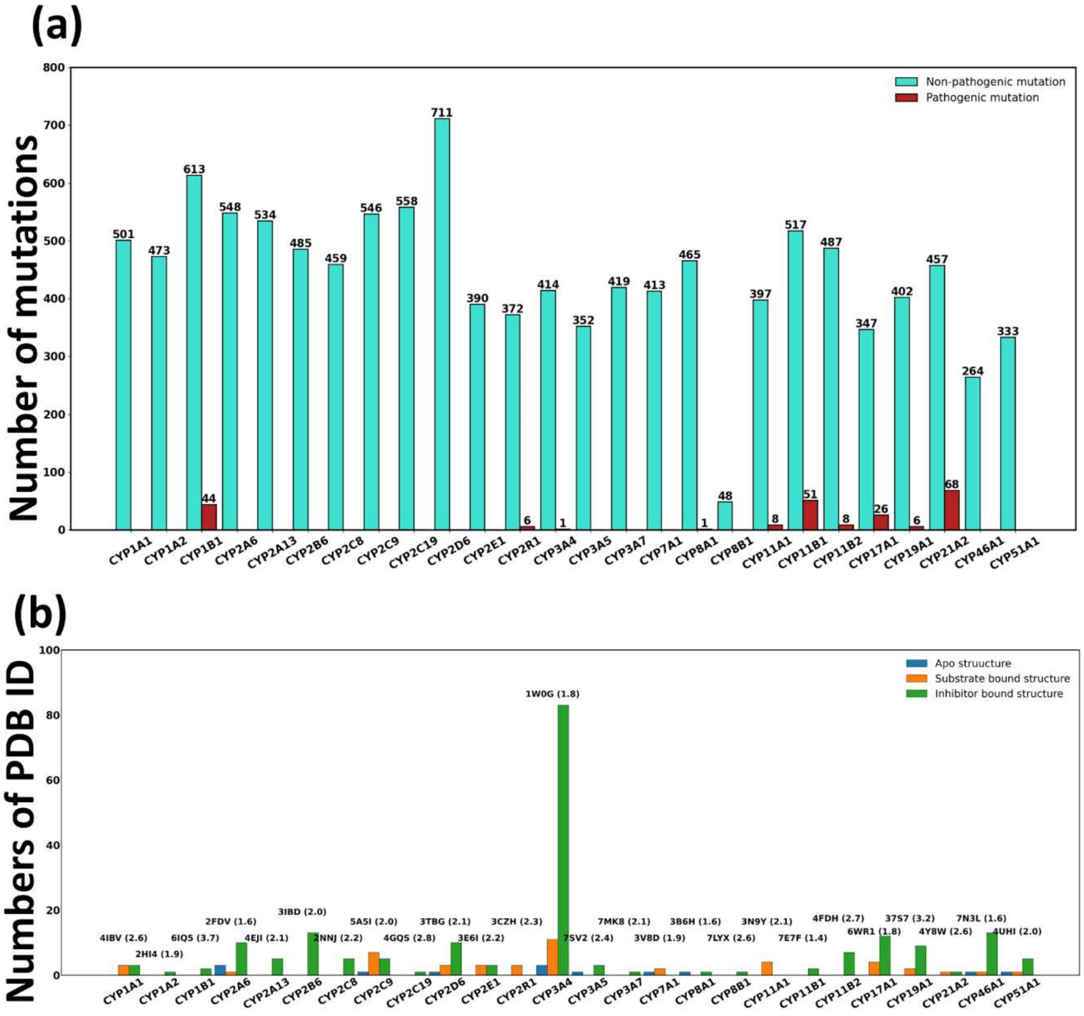


**Figure S1.** (a) Bar plot for 26 CYP450 showing total number of non-pathogenic and pathogenic mutations. (b) X-ray crystallographic structure (apo, substrate bound, inhibitor bound) for 26 CYP450; PDB Id and respective resolution (in Å) of the CYPs are shown above each bar.

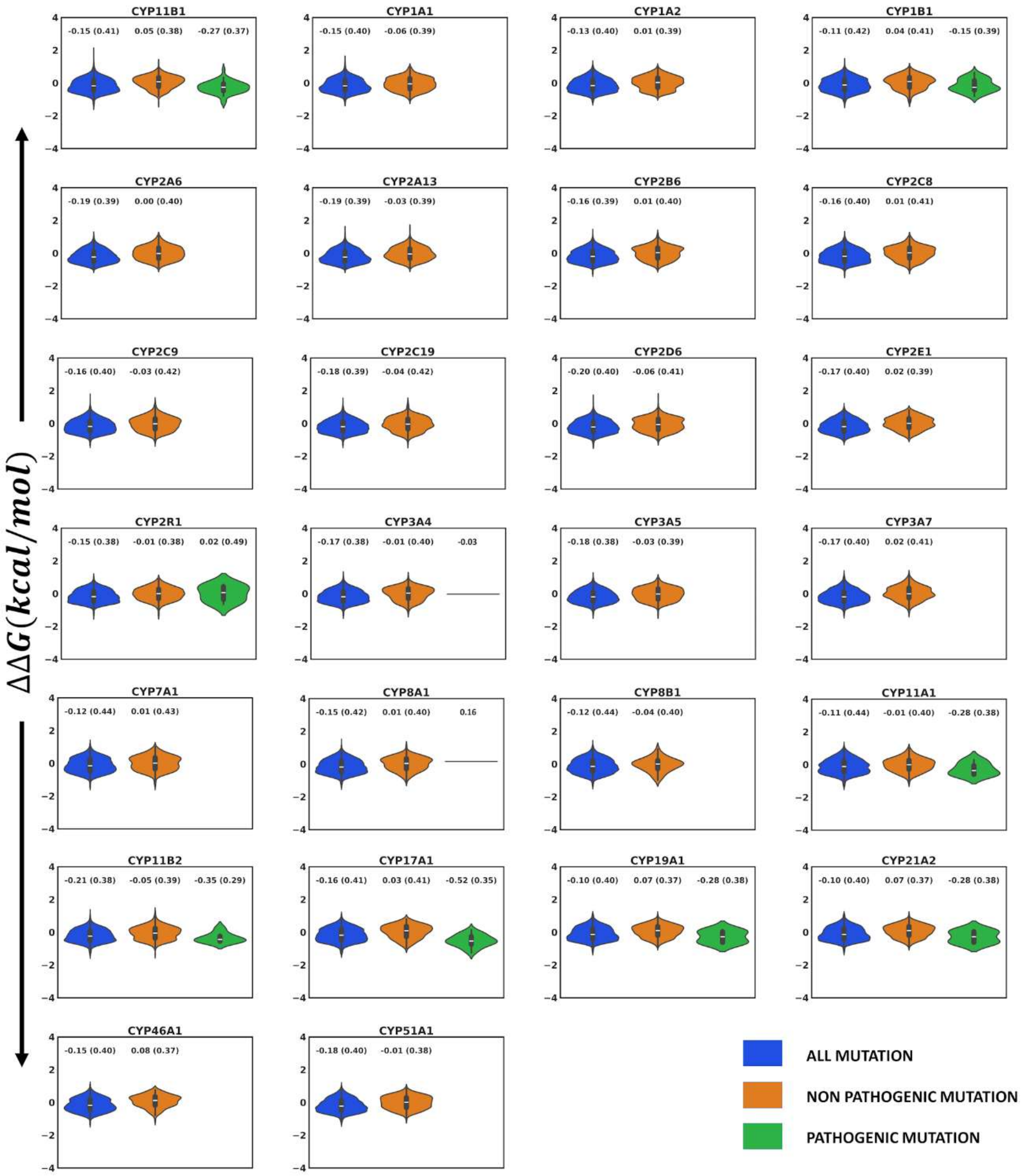


**Figure S2.** Violin plots between all possible mutation categories ('all mutation', 'non-pathogenic mutation', and 'pathogenic mutation') using SNPMUSIC. Y-axis represents ΔΔG value and given colour codes inside the figure depict different Groups.

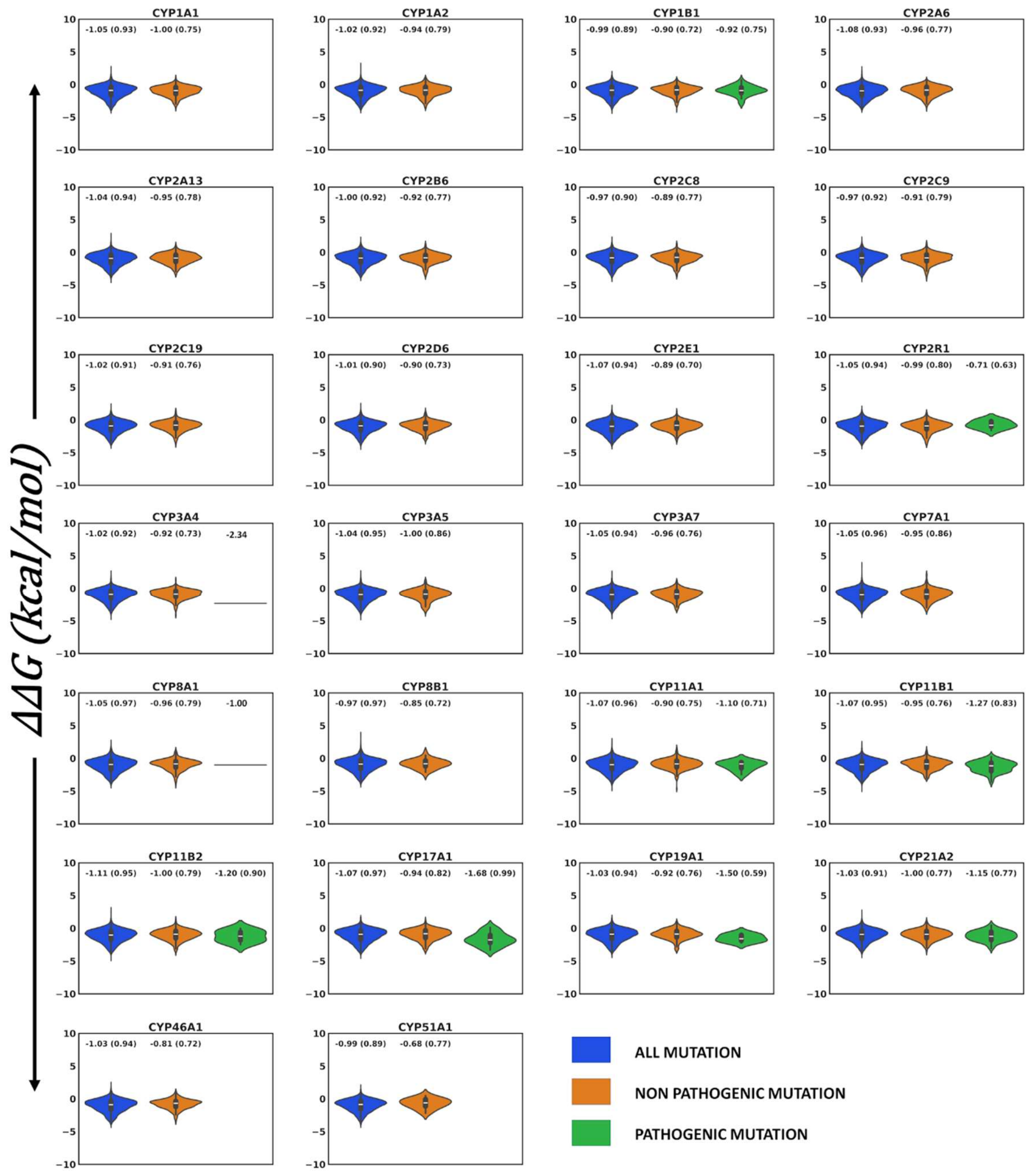


**Figure S3.** Violin plots between all possible mutation categories ('all mutation', 'non-pathogenic mutation', and 'pathogenic mutation') using mCSM. Y-axis represents ΔΔG value and given colour codes inside the figure depict different Groups.

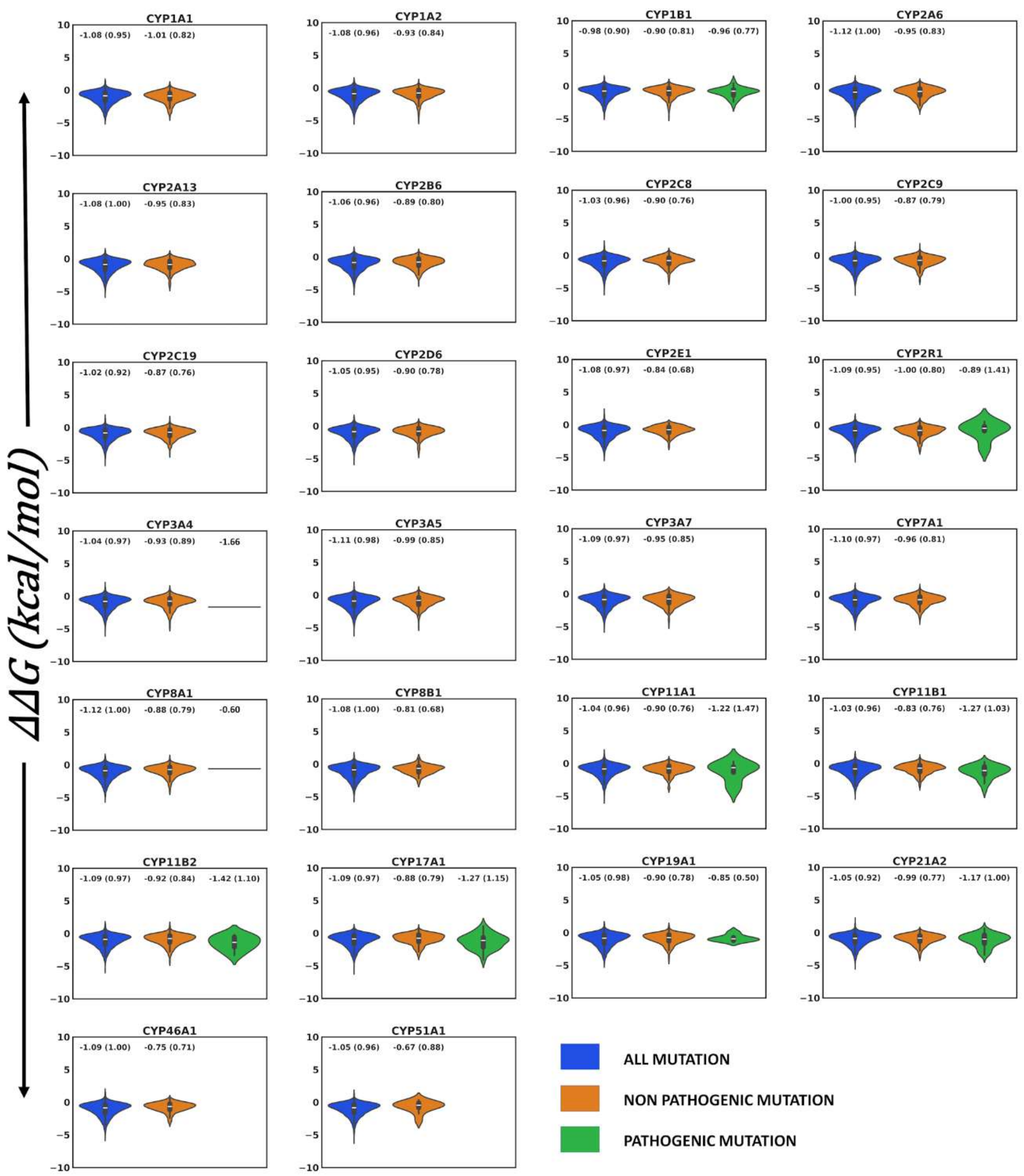


**Figure S4.** Violin plots between all possible mutation categories ('all mutation', 'non-pathogenic mutation', and 'pathogenic mutation') using POPMUSIC. Y-axis represents ΔΔG value and given colour codes inside the figure depict different Groups.

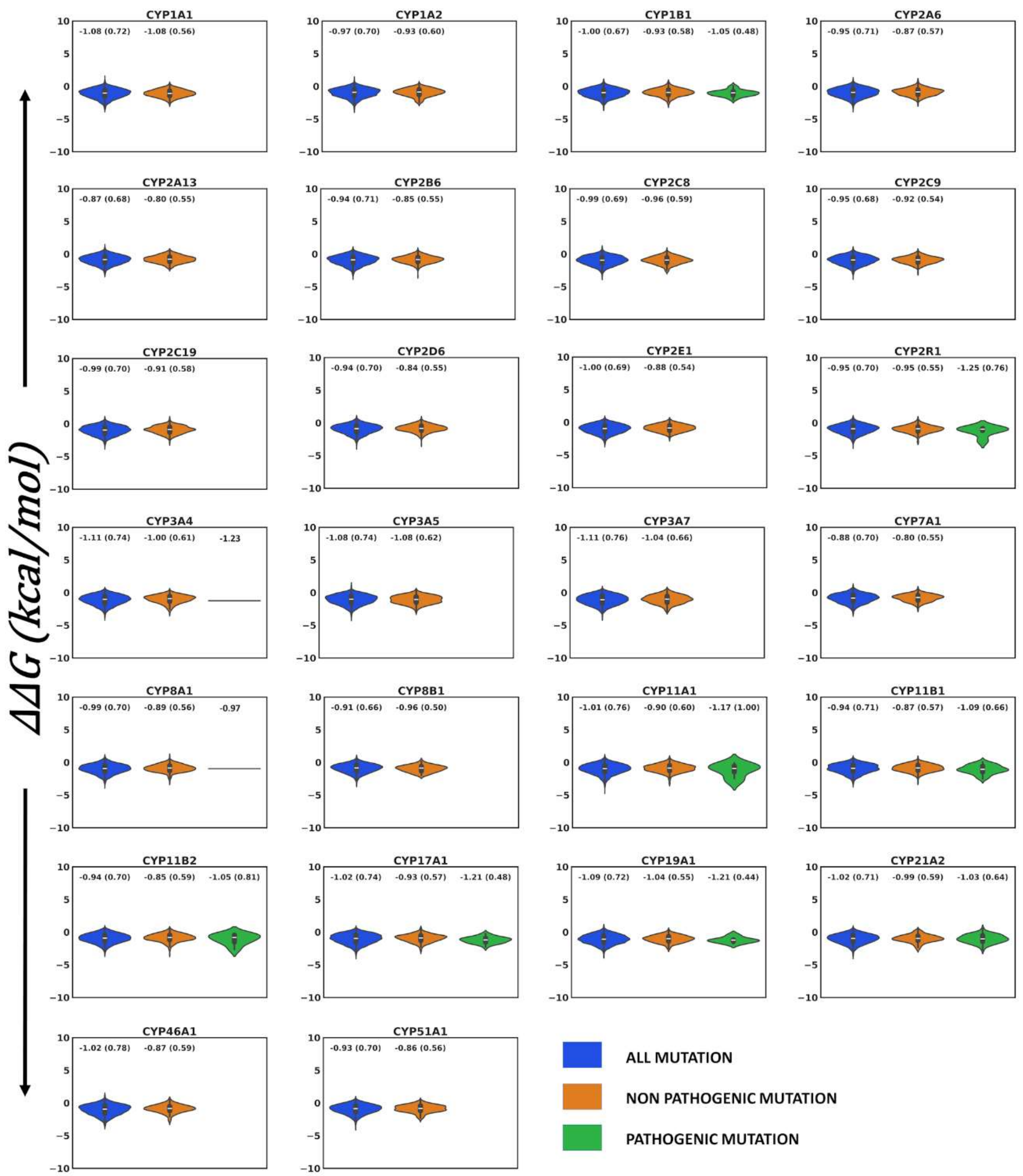


**Figure S5.** Violin plots between all possible mutation categories ('all mutation', 'non-pathogenic mutation', and 'pathogenic mutation') using IMUTANT3.0. Y-axis represents ΔΔG value and given colour codes inside the figure depict different Groups.

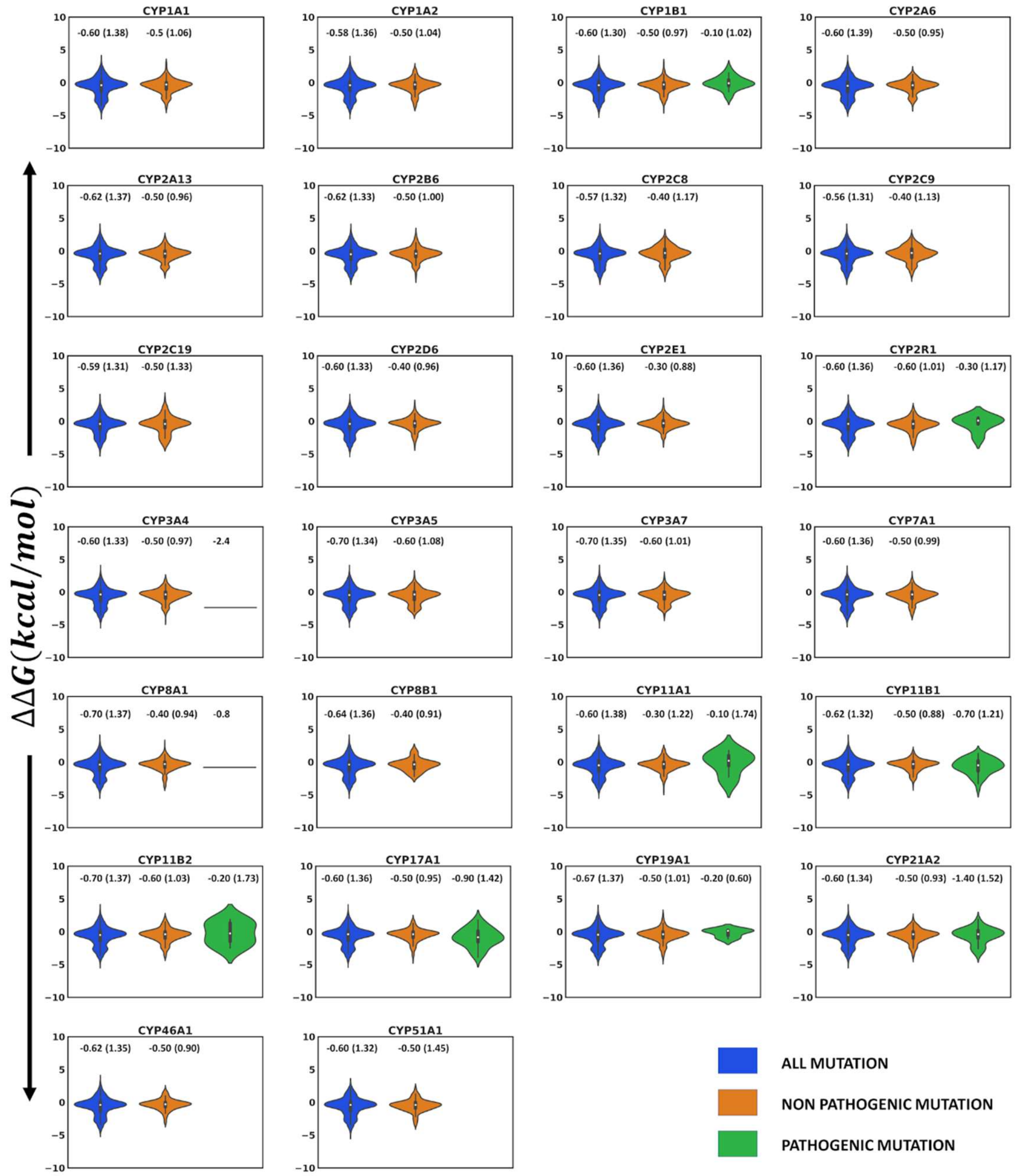


**Figure S6. Violin plots of the mutation groups computed using SIMBA**. The average and standard deviation (in bracket) for each group is shown on the top of each violin.

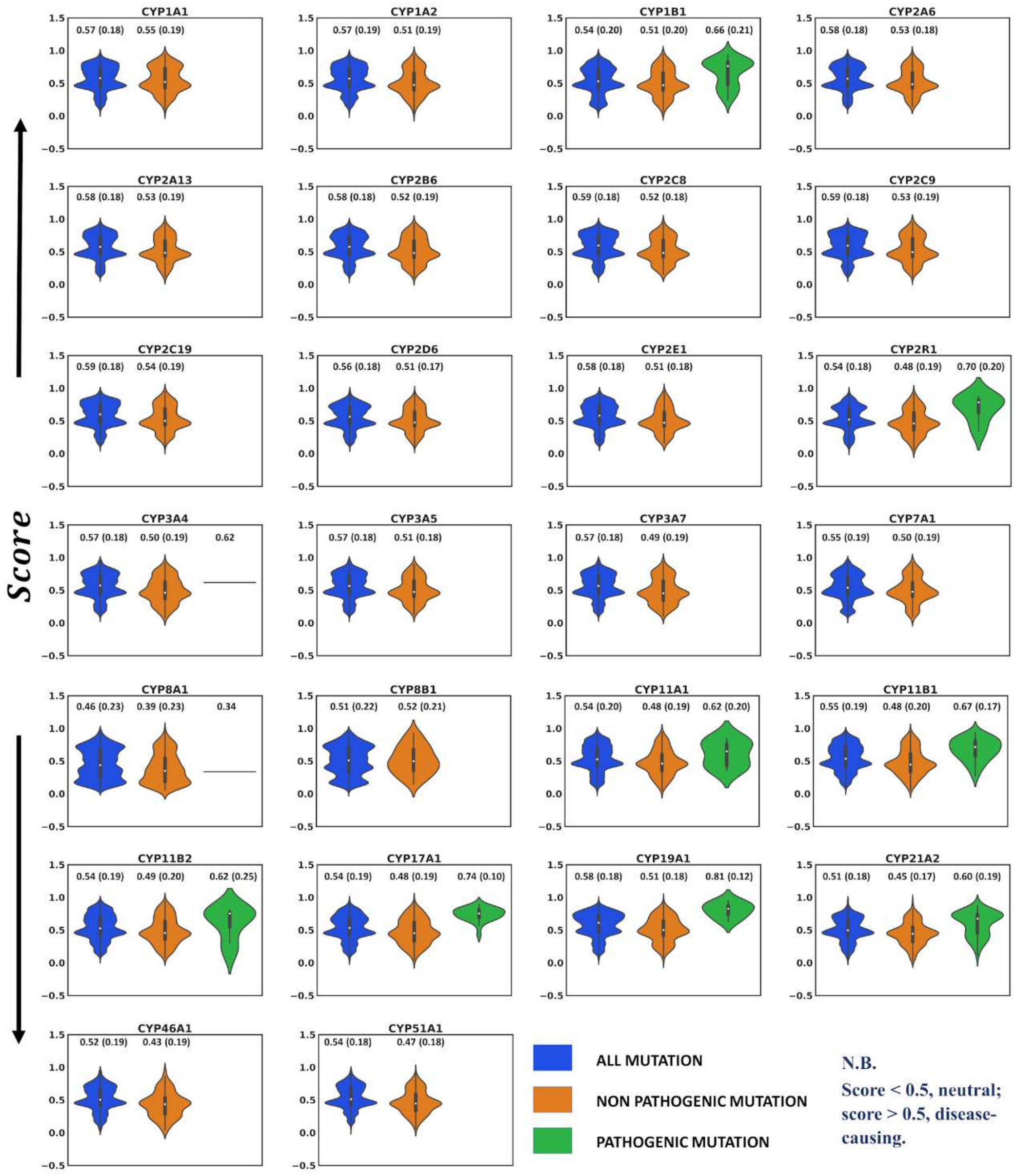


**Figure S7.** Violin plots between all possible mutation categories ('all mutation', 'non-pathogenic mutation', and 'pathogenic mutation') using METASNP. The numeric value over each violin represents the average (standard deviation) of score.

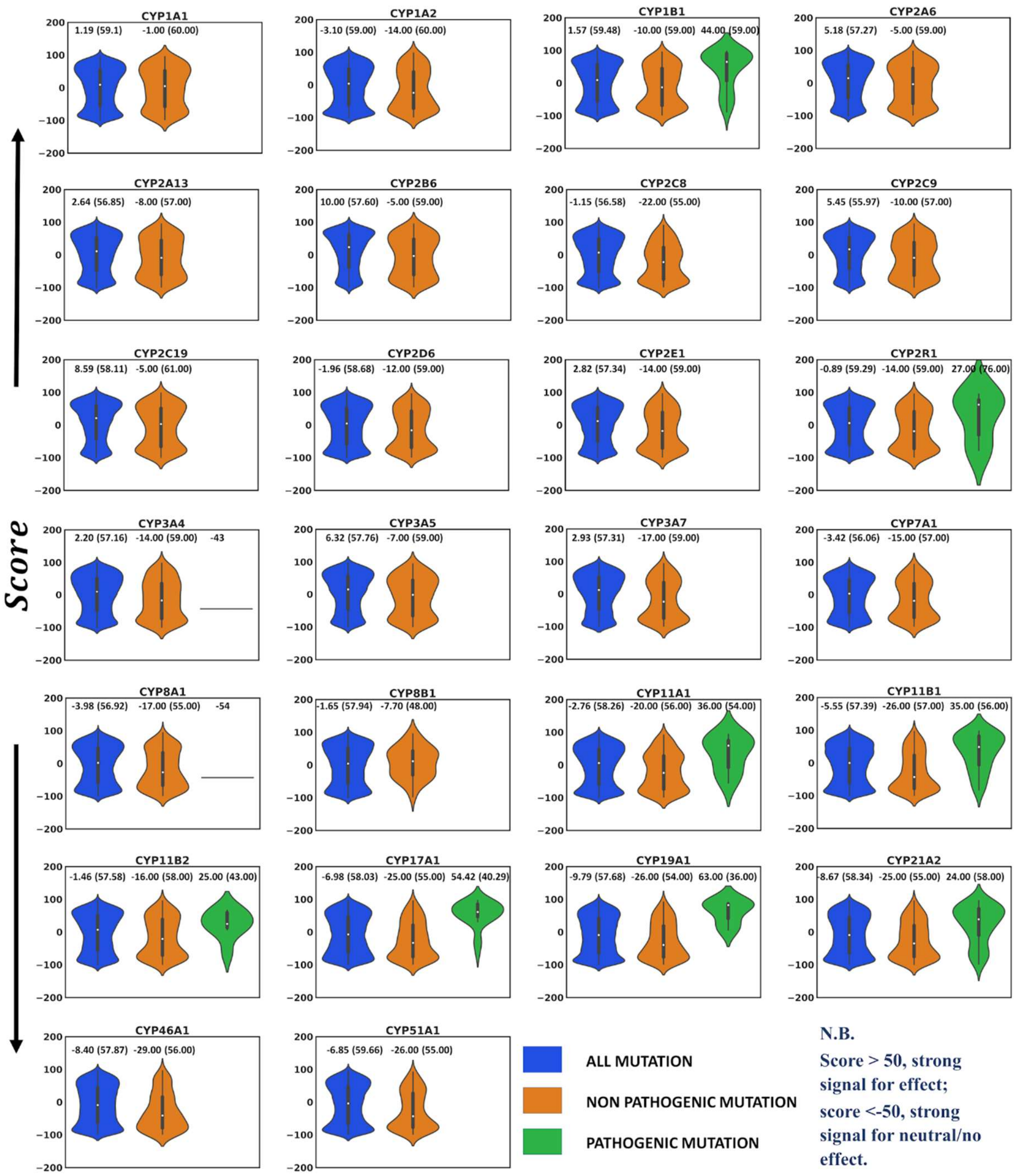


**Figure S8.** Violin plots between all possible mutation categories ('all mutation', 'non-pathogenic mutation', and 'pathogenic mutation') using SNAP2. The numeric value over each violin represents the average (standard deviation) of score.

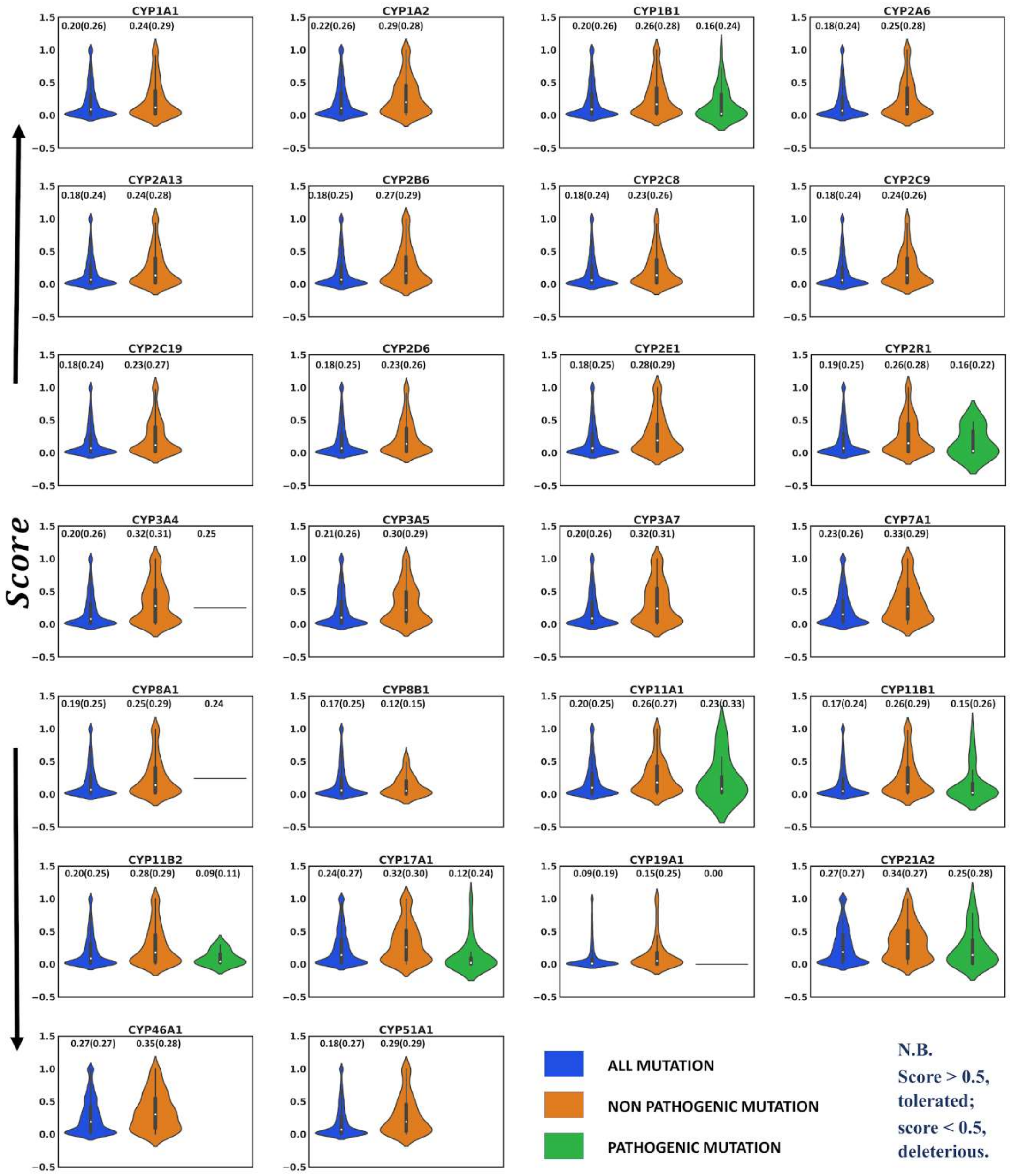


**Figure S9.** Violin plots between all possible mutation categories ('all mutation', 'non-pathogenic mutation', and 'pathogenic mutation') using SIFT. The numeric value over each violin represents the average (standard deviation) of score.

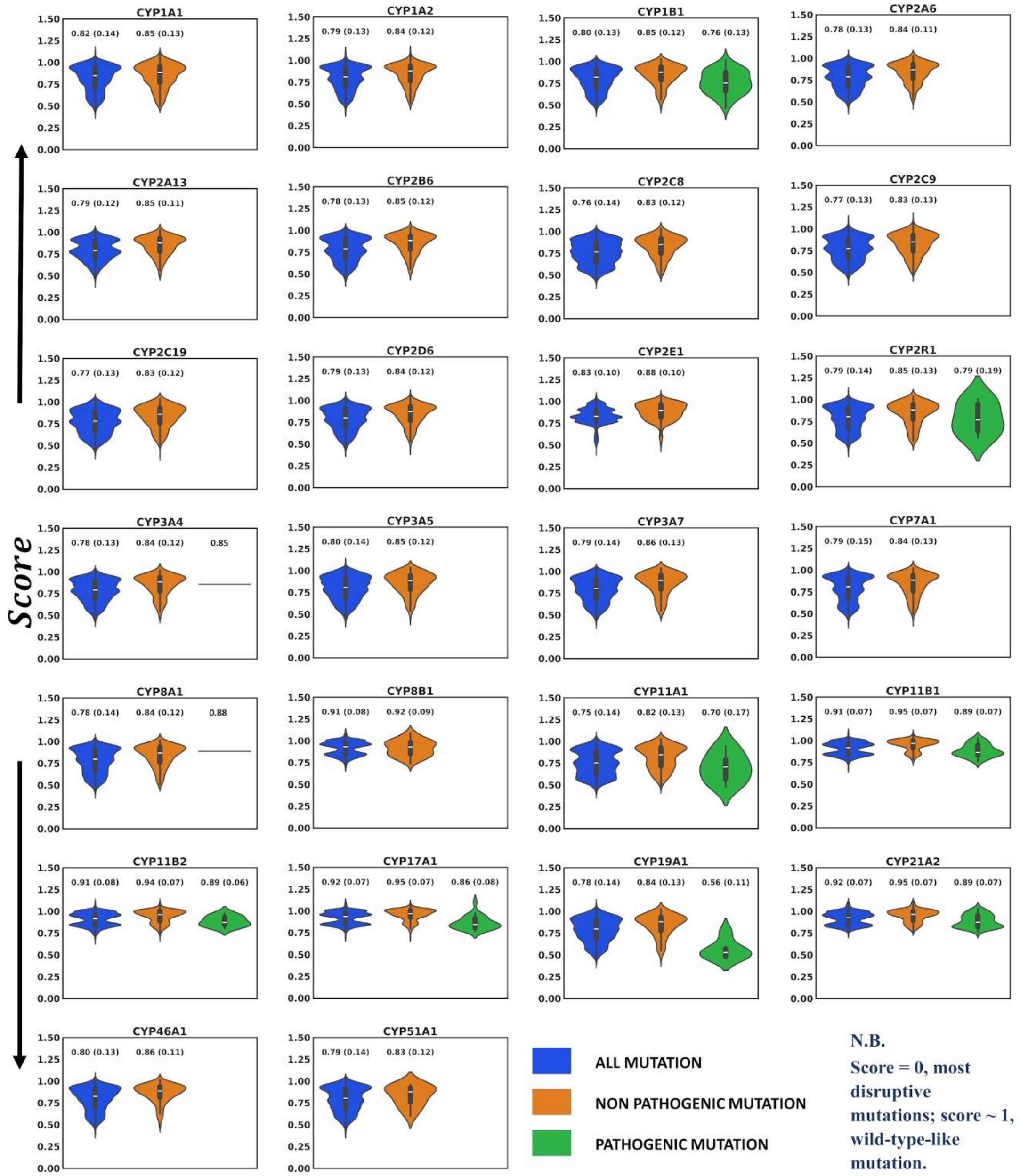


**Figure S10.** Violin plots between all possible mutation categories ('all mutation', 'non-pathogenic mutation', and 'pathogenic mutation') using ENVISION. The numeric value over each violin represents the average (standard deviation) of score.

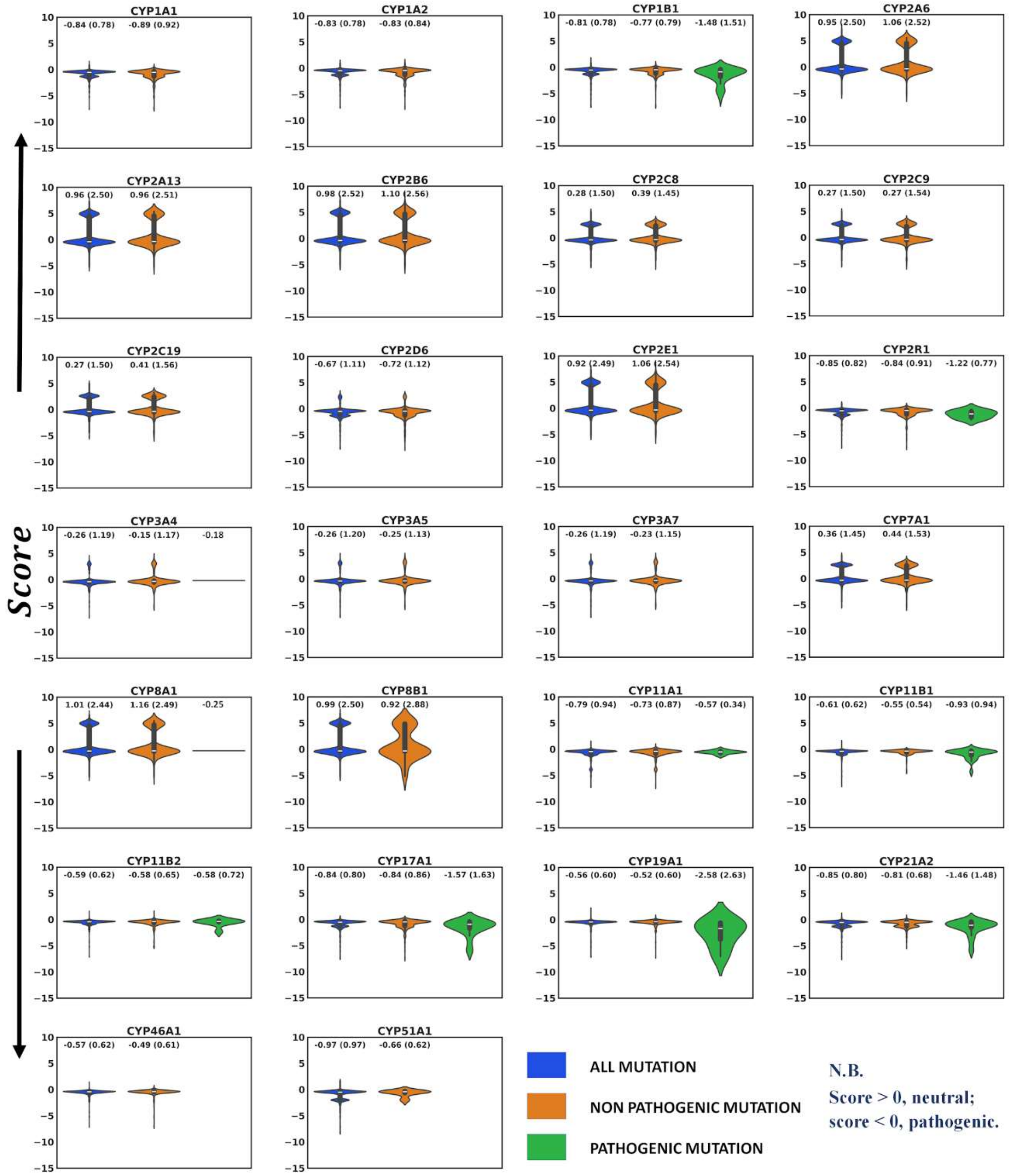


**Figure S11.** Violin plots between all possible mutation categories ('all mutation', 'non-pathogenic mutation', and 'pathogenic mutation') using FATHMM. The numeric value over each violin represents the average (standard deviation) of score.

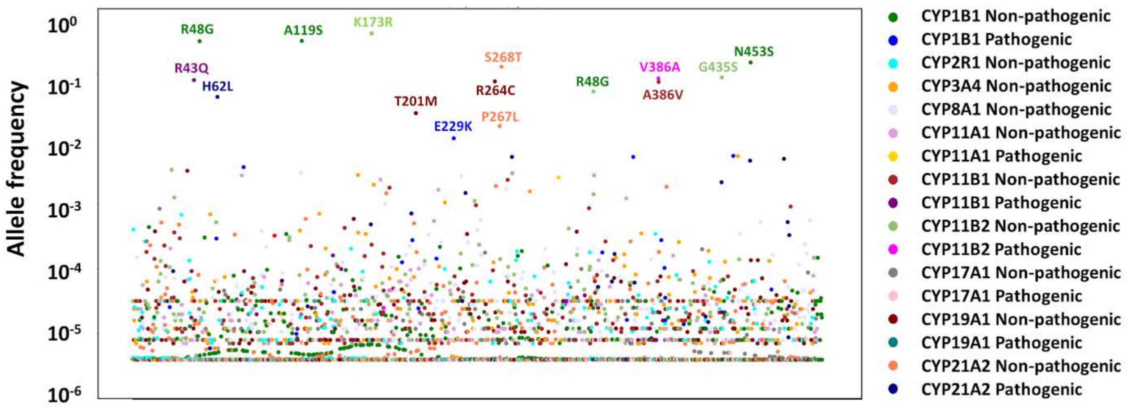


**Figure S12. Analysis of allele frequency of ten CYPs (with pathogenic dataset) for non-pathogenic and pathogenic mutations.** The y-axis represents allele frequence, which is allele count divided by allele number. The data is collected from gnomAD database. Some higher frequency mutations are labelled in the plot. Allele frequency of CYP2R1 (Pathogenic), CYP3A4 (Pathogenic), and CYP8A1 (Pathogenic) are missing in gnomAD database.

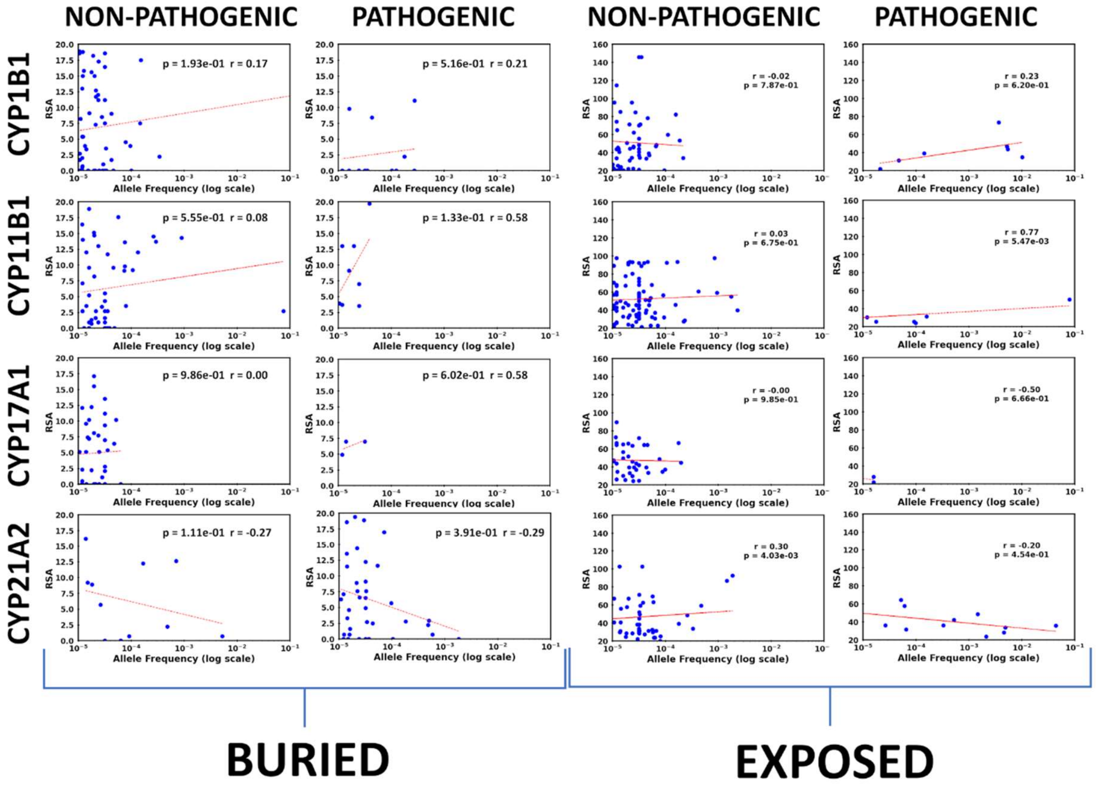


**Figure S13.** Correlation between allele frequency and relative solvent accessibility for CYP1B1, CYP11B1, CYP17A1, and CYP21A2.

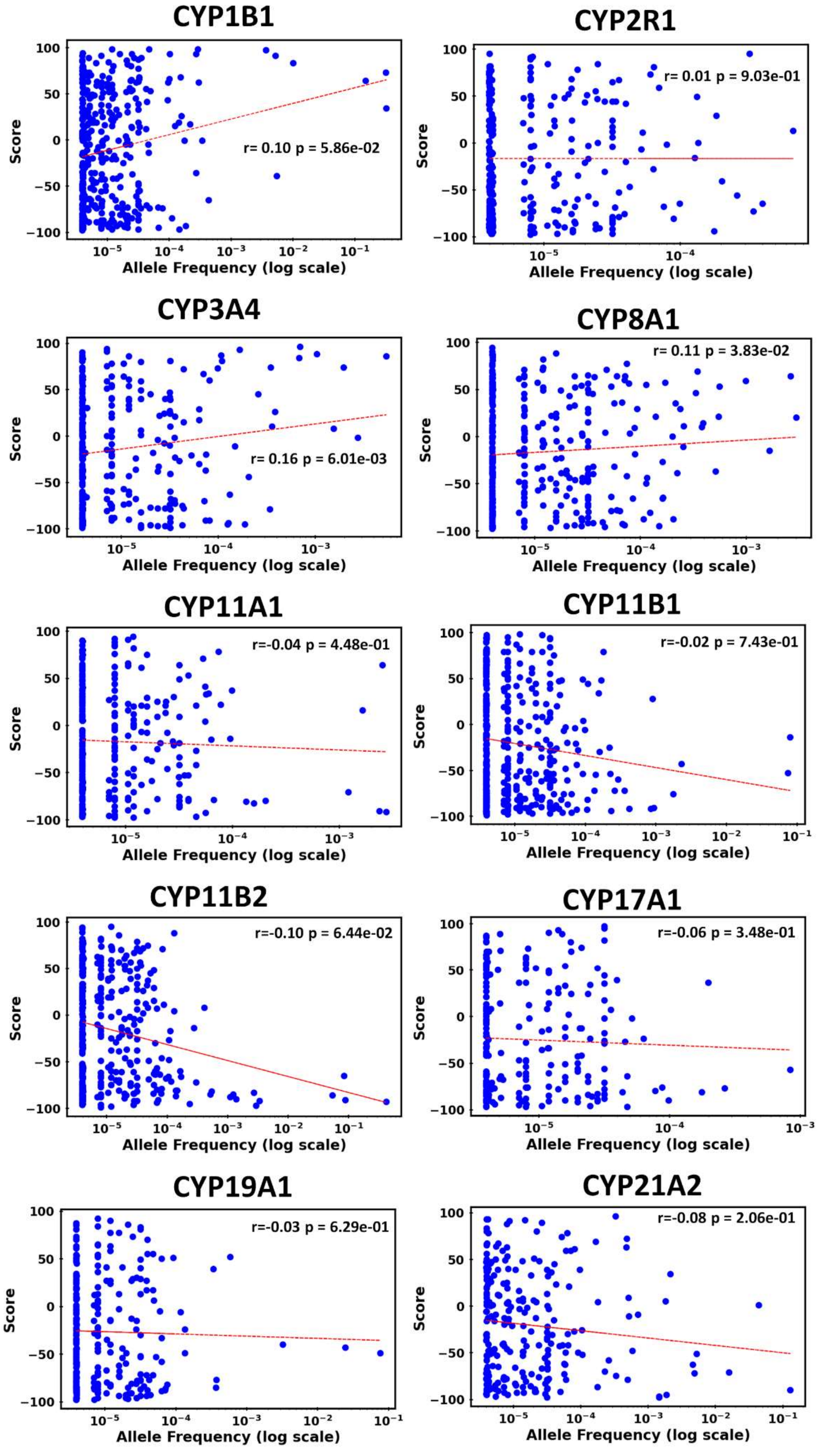


**Figure S14.** Correlation between allele frequency and SNAP2 score of 10 CYP450. Y-axis represents SNAP2 score and X-axis represents corresponding allele frequency.

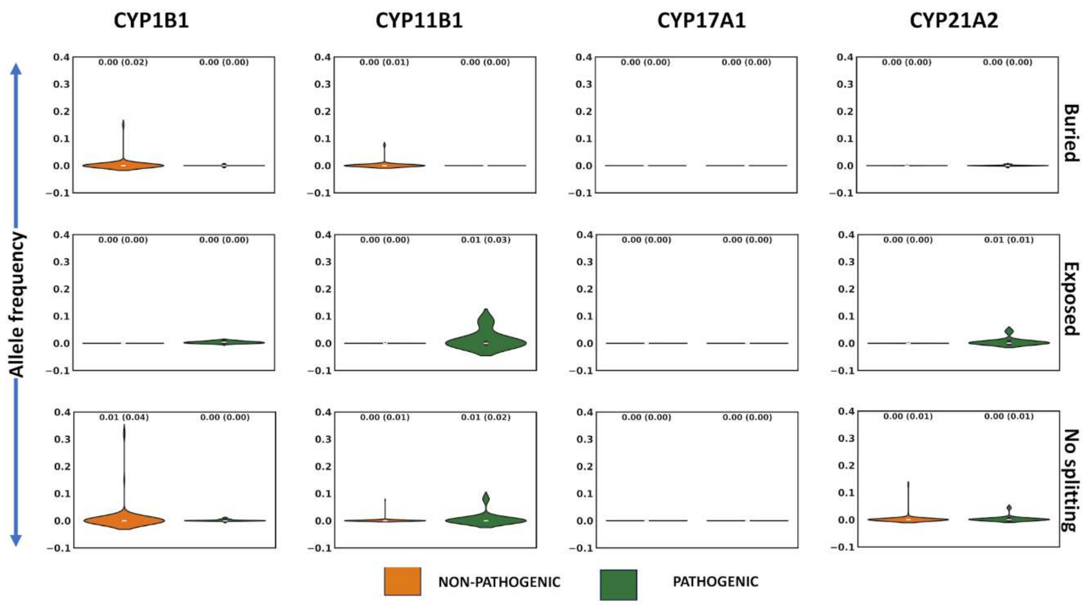


**Figure S15.** Violin plot using allele frequency as descriptor for the discrimination between pathogenic and non-pathogenic mutations.

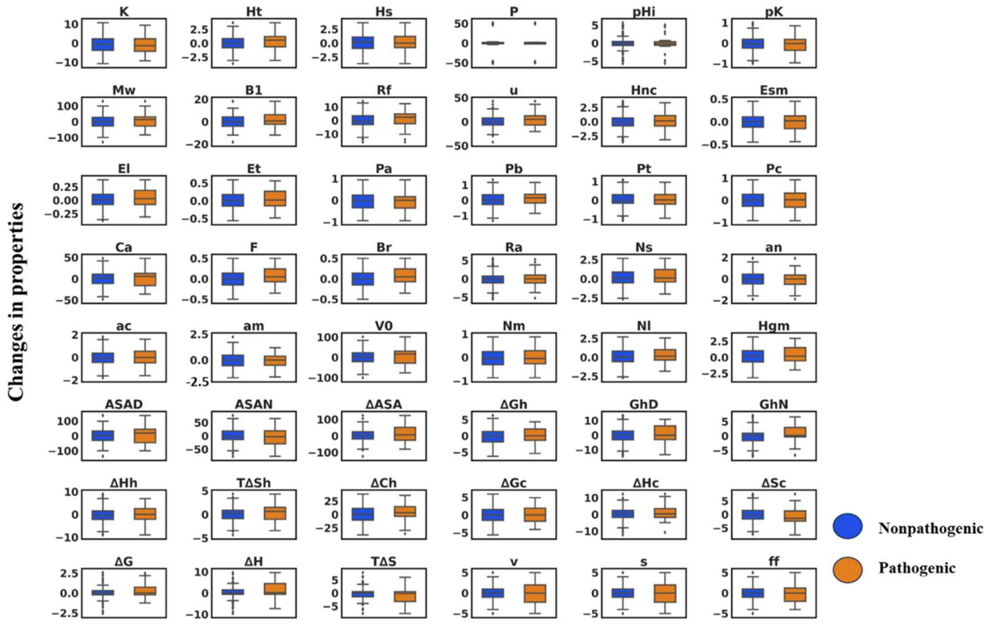


**Figure S16.** Box-whisker plots of 48 amino acid properties change for all possible mutation categories ('non-pathogenic mutation', and 'pathogenic mutation') CYP1B1.

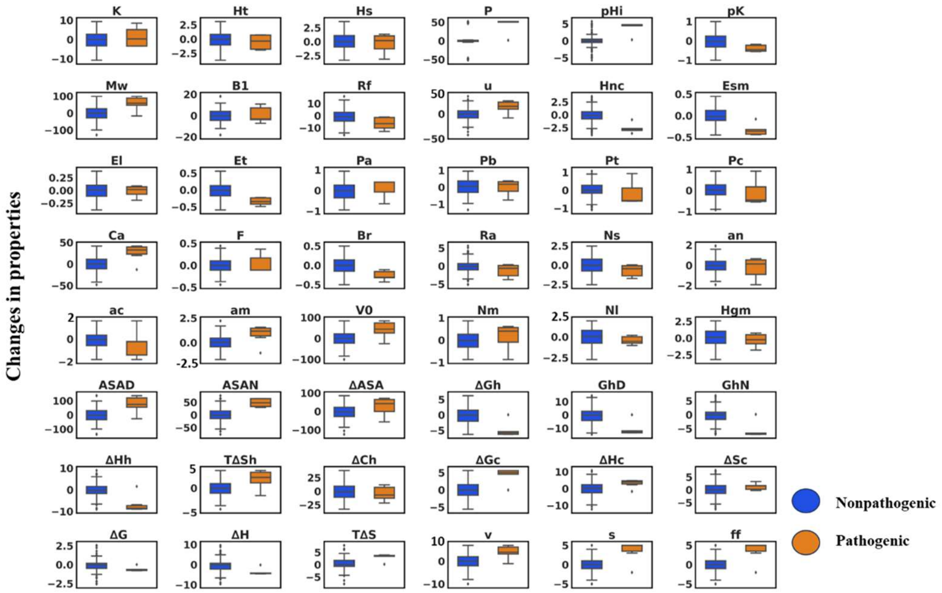


**Figure S17.** Box-whisker plots of 48 amino acid properties change for all possible mutation categories ( ‘non-pathogenic mutation’, and ‘pathogenic mutation’) CYP2R1.

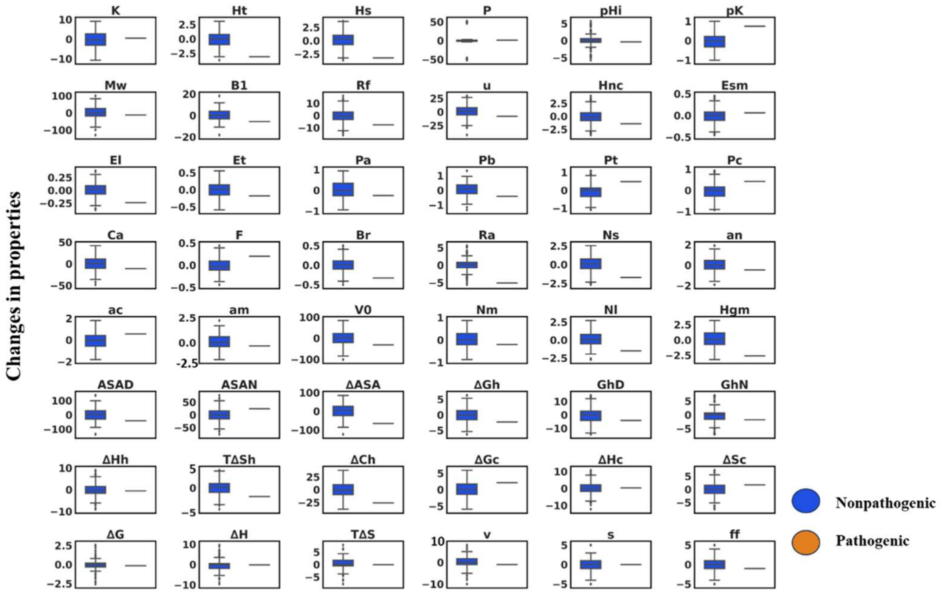


**Figure S18.** Box-whisker plots of 48 amino acid properties change for all possible mutation categories ( ‘non-pathogenic mutation’, and ‘pathogenic mutation’) CYP3A4.

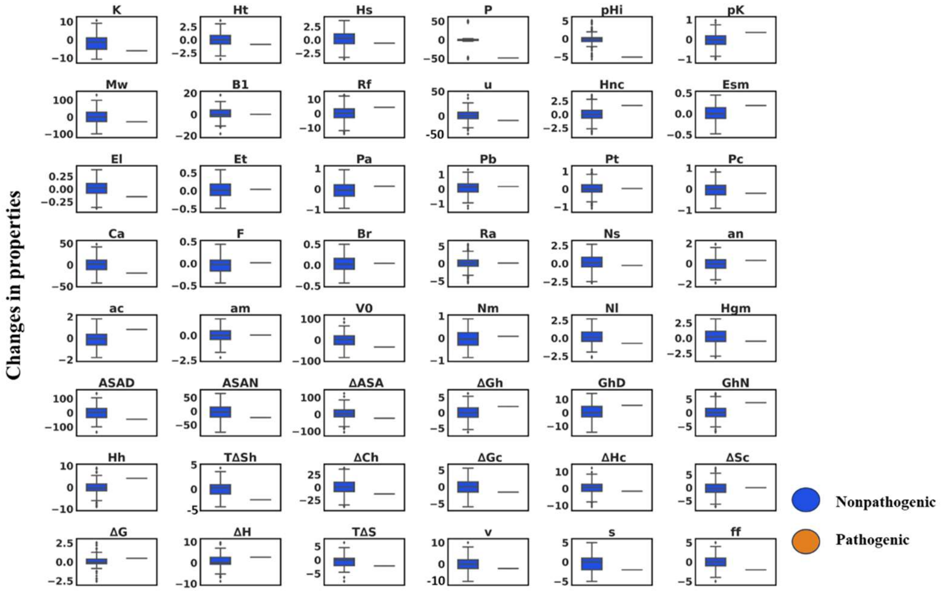


**Figure S19.** Box-whisker plots of 48 amino acid properties change for all possible mutation categories ('non-pathogenic mutation', and 'pathogenic mutation') CYP8A1.

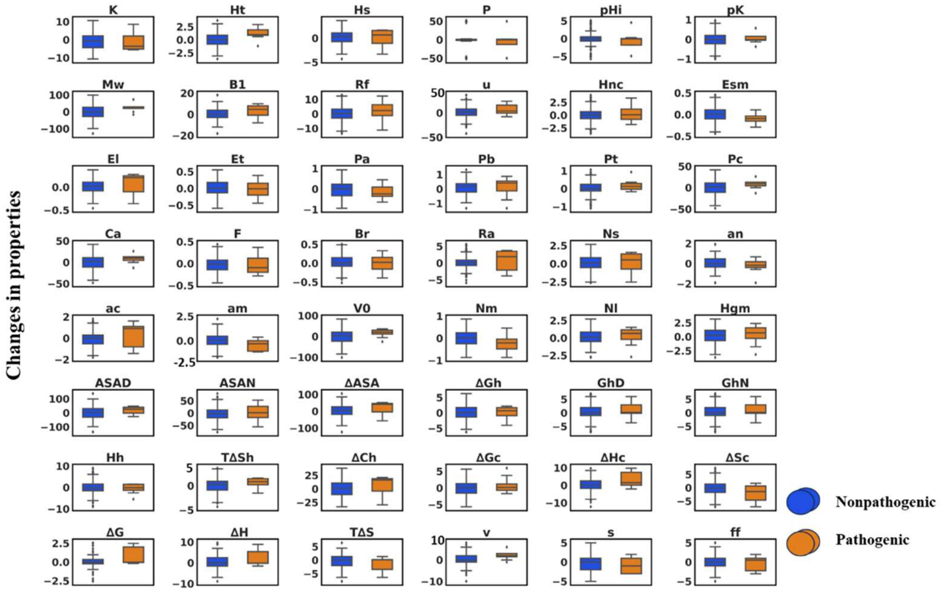


**Figure S20.** Box-whisker plots of 48 amino acid properties change for all possible mutation categories ('non-pathogenic mutation', and 'pathogenic mutation') CYP11A1.

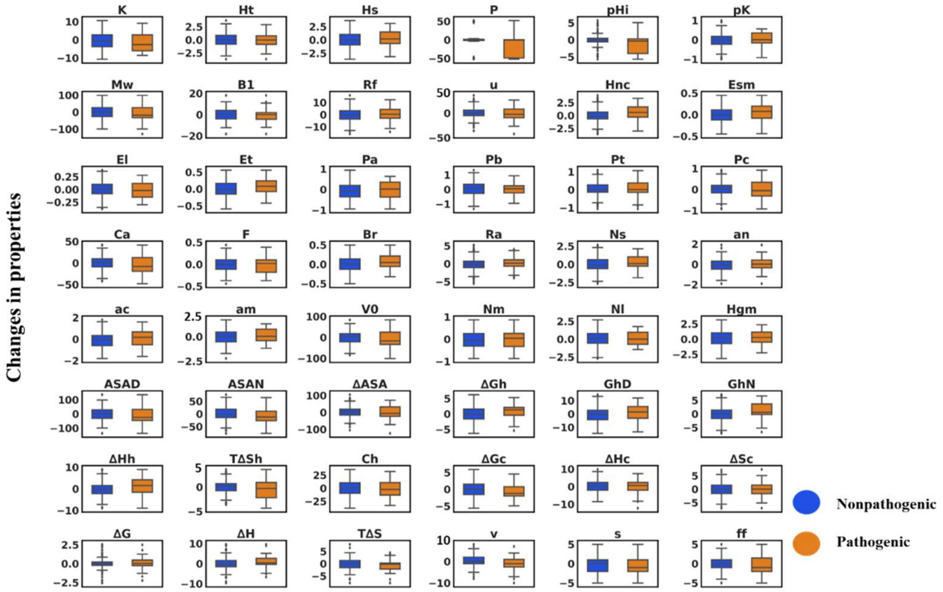


**Figure S21.** Box-whisker plots of 48 amino acid properties change for all possible mutation categories (‘non-pathogenic mutation’, and ‘pathogenic mutation’) CYP11B1.

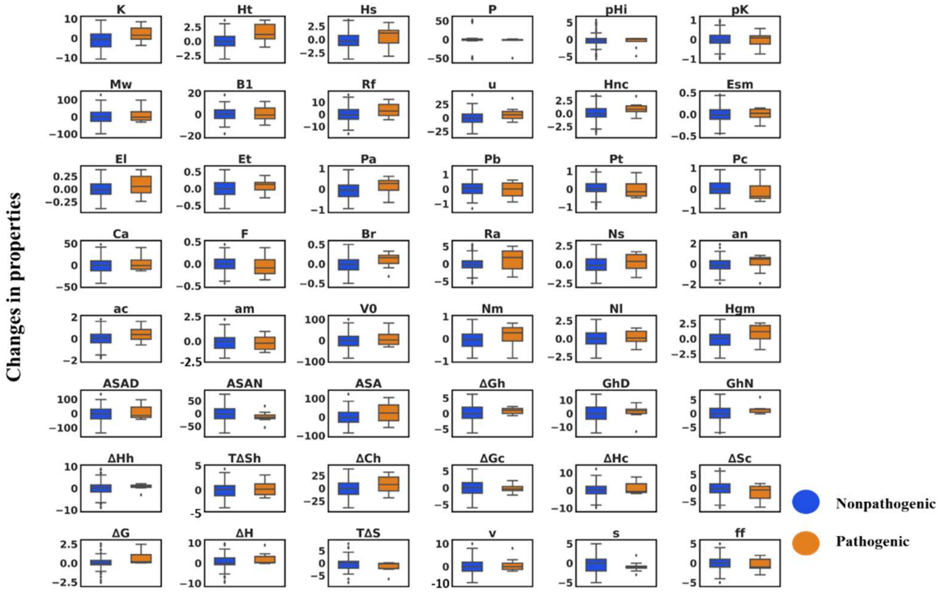


**Figure S22.** Box-whisker plots of 48 amino acid properties change for all possible mutation categories (‘non-pathogenic mutation’, and ‘pathogenic mutation’) CYP11B2.

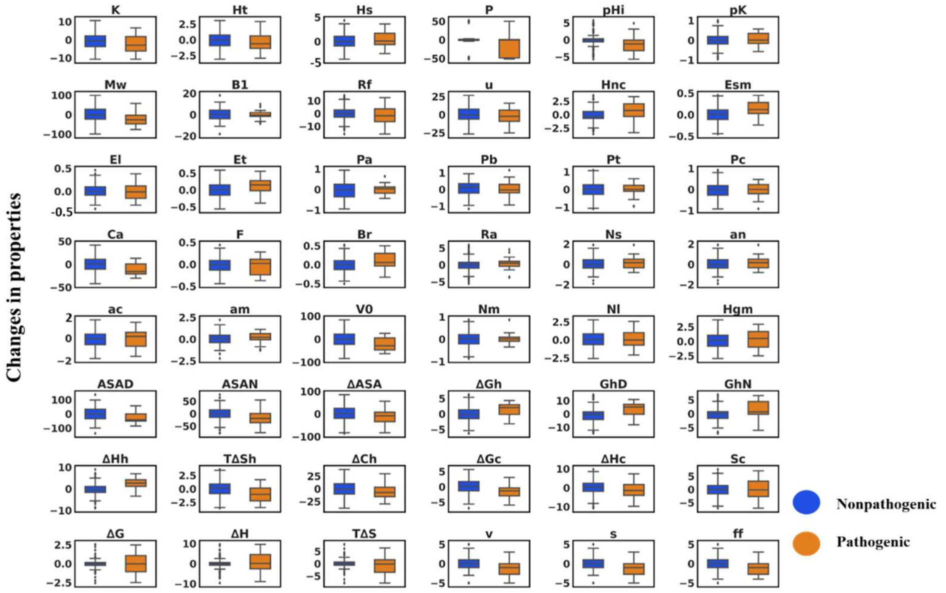


**Figure S23.** Box-whisker plots of 48 amino acid properties change for all possible mutation categories ('non-pathogenic mutation', and 'pathogenic mutation') CYP17A1.

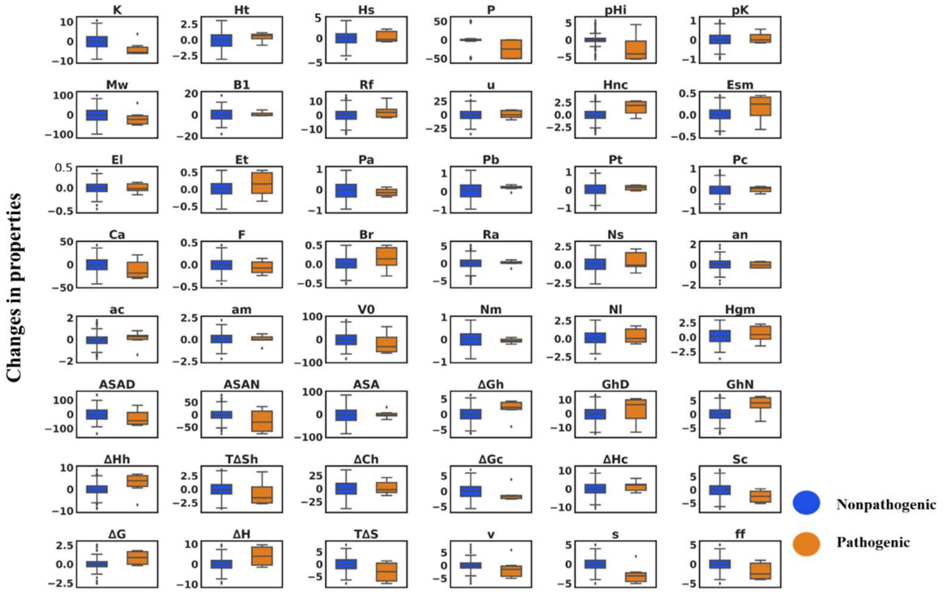


**Figure S24.** Box-whisker plots of 48 amino acid properties change for all possible mutation categories ('non-pathogenic mutation', and 'pathogenic mutation') CYP19A1.

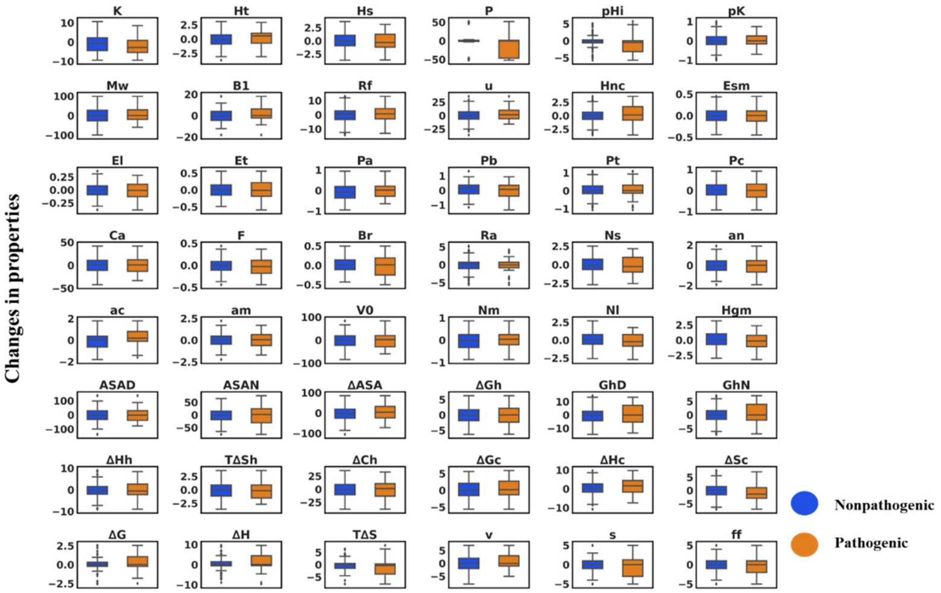

**Figure S25.** Box-whisker plots of 48 amino acid properties change for all possible mutation categories (‘non-pathogenic mutation’, and ‘pathogenic mutation’) CYP21A2.

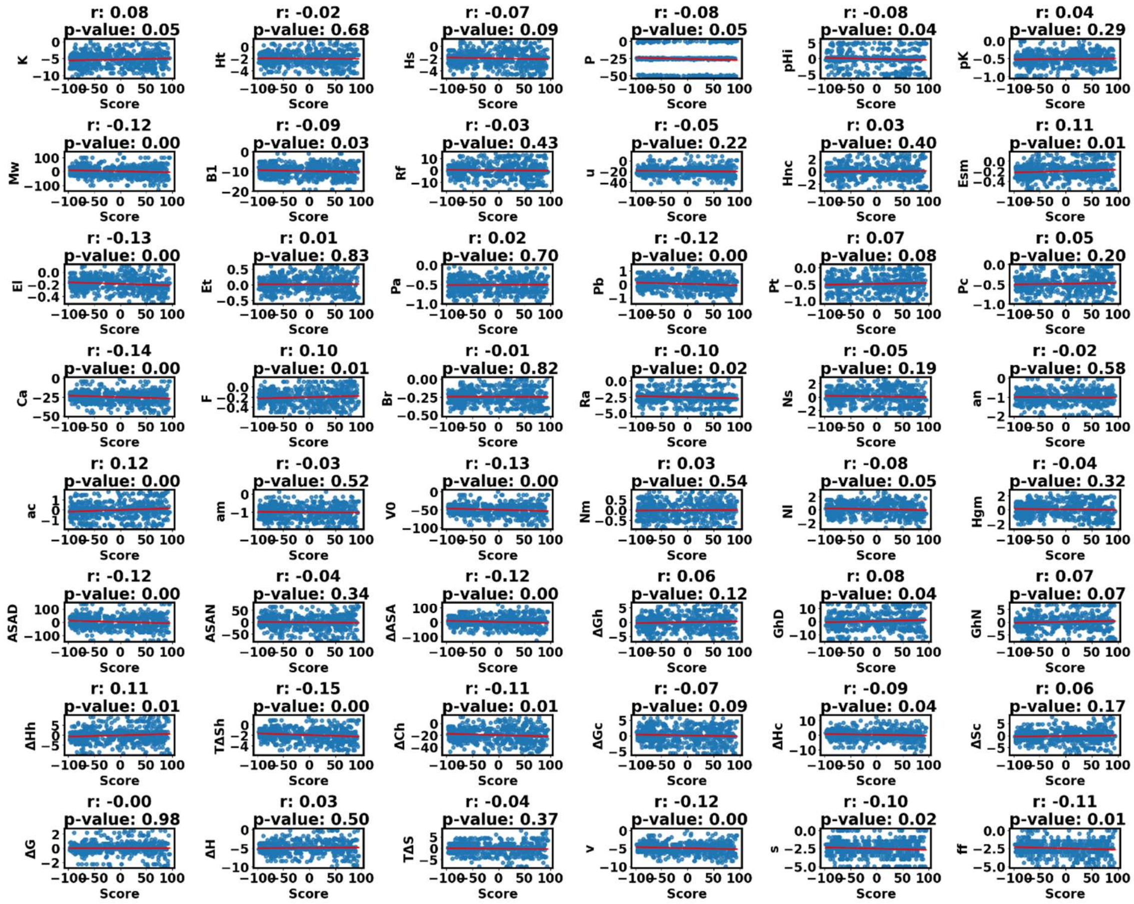

**Figure S26.** Correlation plots between change in 48 amino property and SNAP2 score of CYP1B1 (Non-Pathogenic). Correlation coefficient is represented by ‘r’,  closer to +1 represents positive correlation, more closer to -1 represents negative correlation, and 0 means no correlation.

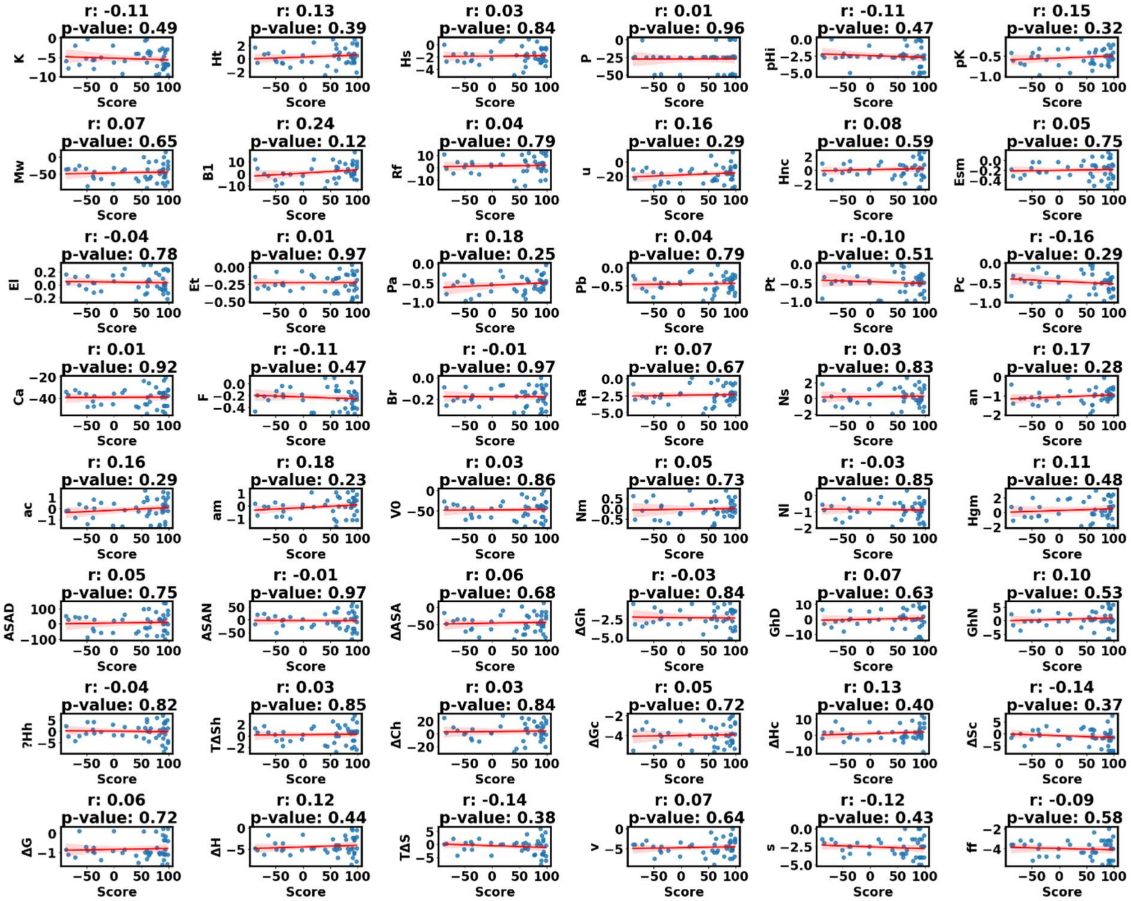

**Figure S27.** Correlation between change in 48 amino property and SNAP2 score of CYP1B1 (Pathogenic). Correlation coefficient is represented by 'r', closer to +1 represents positive correlation, more closer to -1 represents negative correlation, and 0 means no correlation.

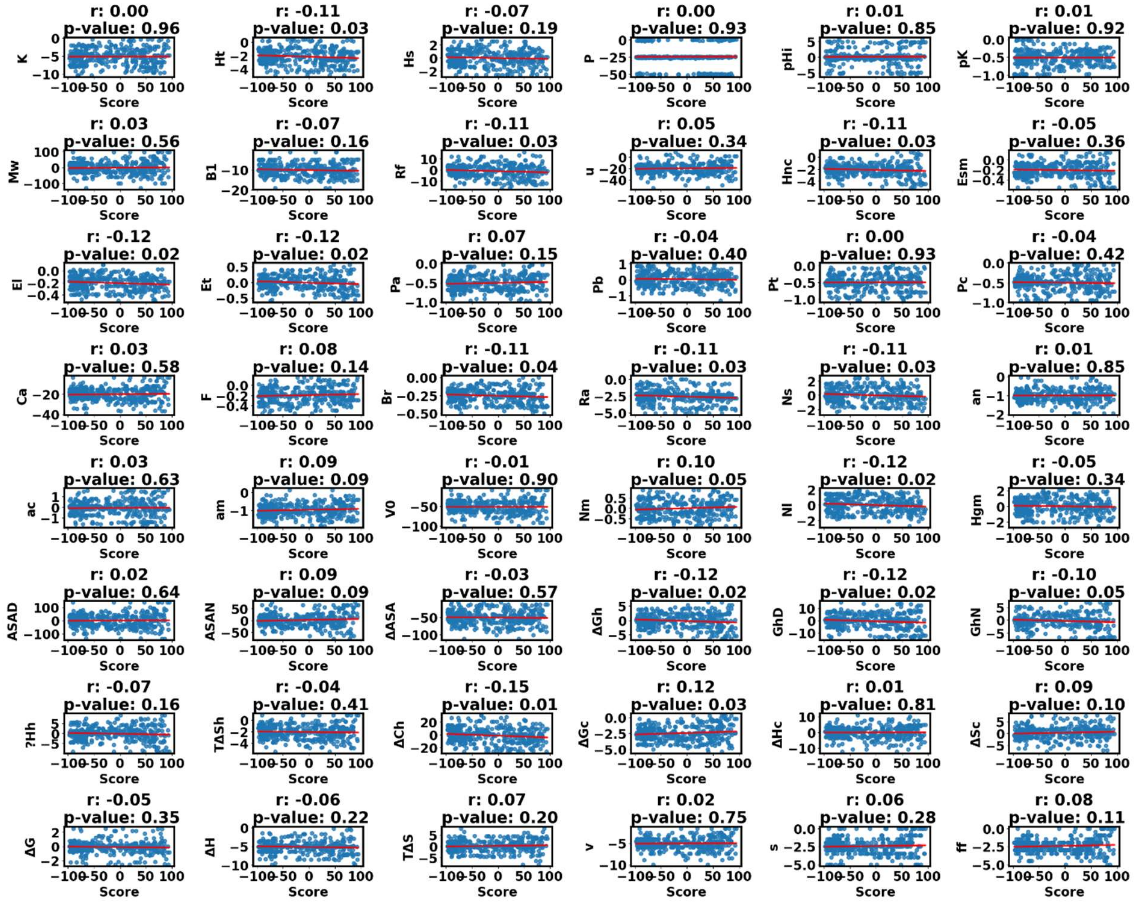

**Figure S28.** Correlation between change in 48 amino property and SNAP2 score of CYP2R1 (Non-Pathogenic). Correlation coefficient is represented by ‘r’, more closer to +1 represents positive correlation, more closer to -1 represents negative correlation, and 0 means no correlation.

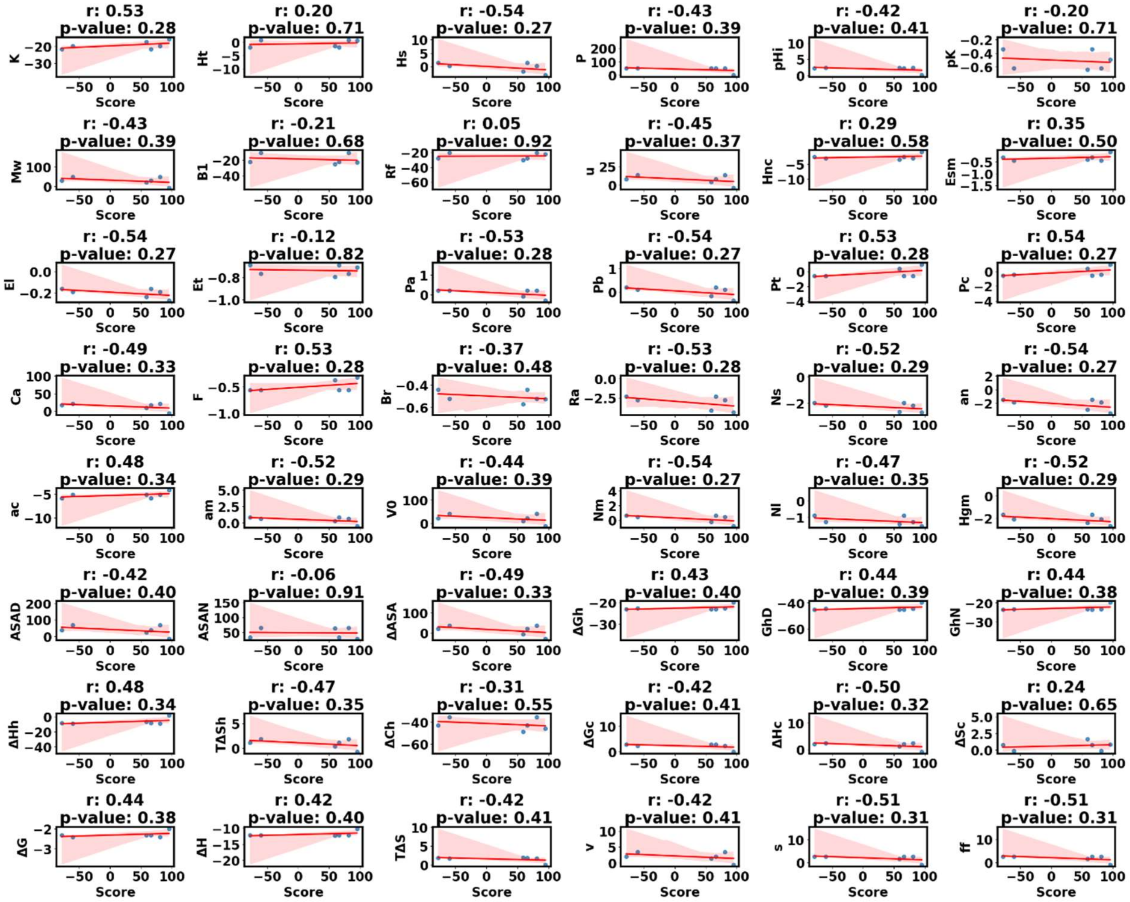


**Figure S29.** Correlation between change in 48 amino property and SNAP2 score of CYP2R1 (Pathogenic). Correlation coefficient is represented by 'r', more closer to +1 represents positive correlation, more closer to -1 represents negative correlation, and 0 means no correlation.

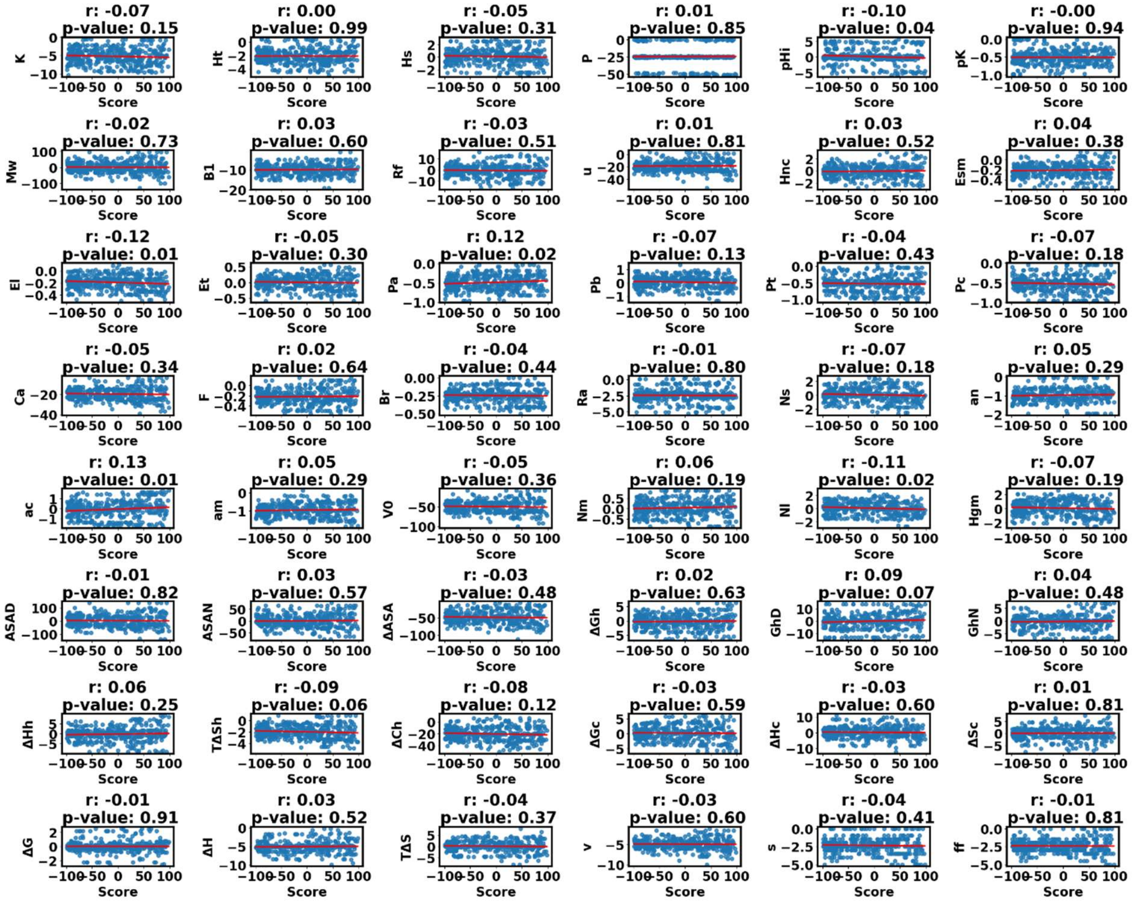


**Figure S30.** Correlation between change in 48 amino property and SNAP2 score of CYP3A4 (Non-Pathogenic). Correlation coefficient is represented by 'r', more closer to +1 represents positive correlation, more closer to -1 represents negative correlation, and 0 means no correlation.

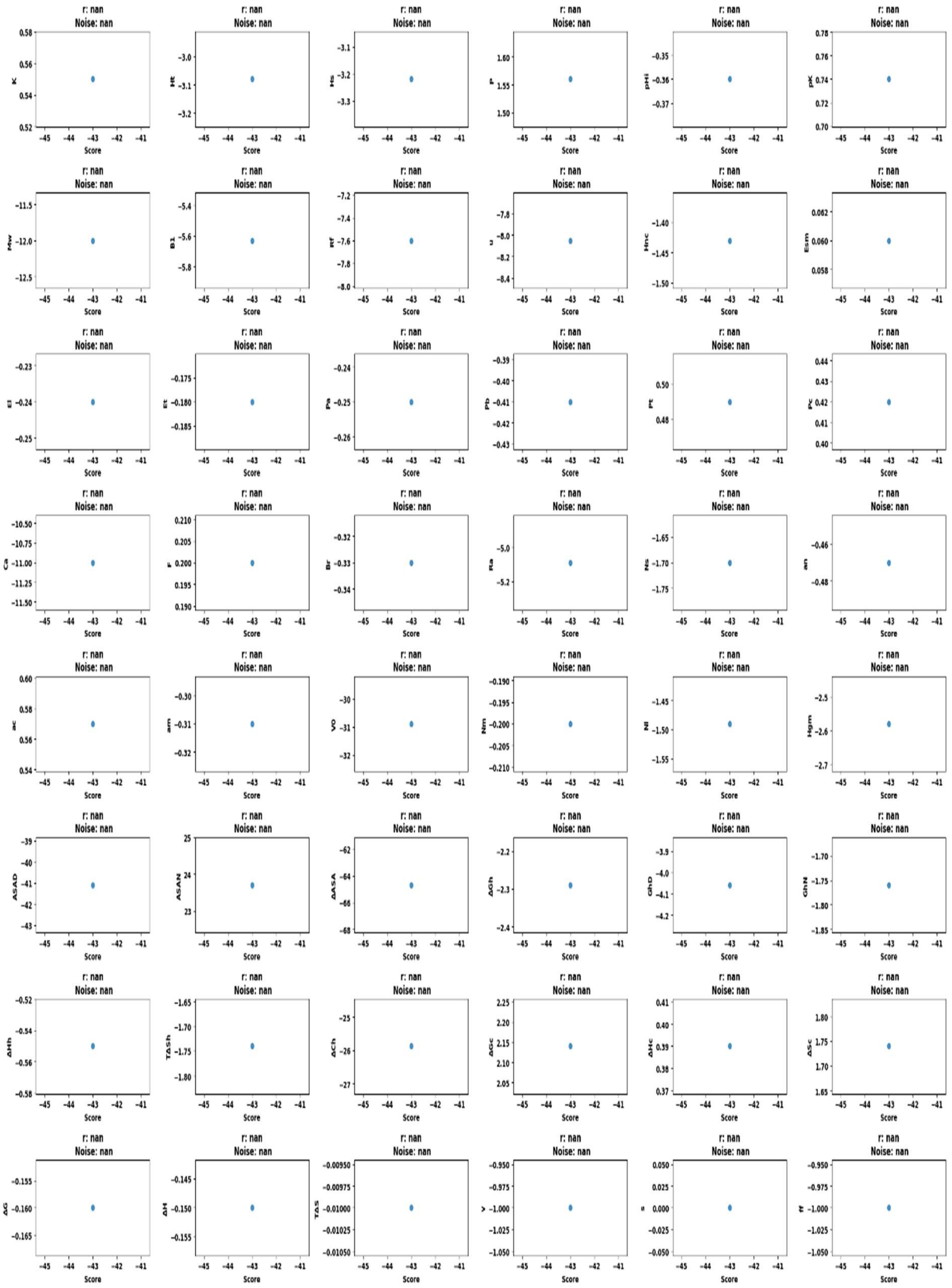


**Figure S31.** Correlation between change in 48 amino property and SNAP2 score of CYP3A4 (Pathogenic). Correlation coefficient is represented by 'r', more closer to +1 represents positive correlation, more closer to -1 represents negative correlation, and 0 means no correlation.

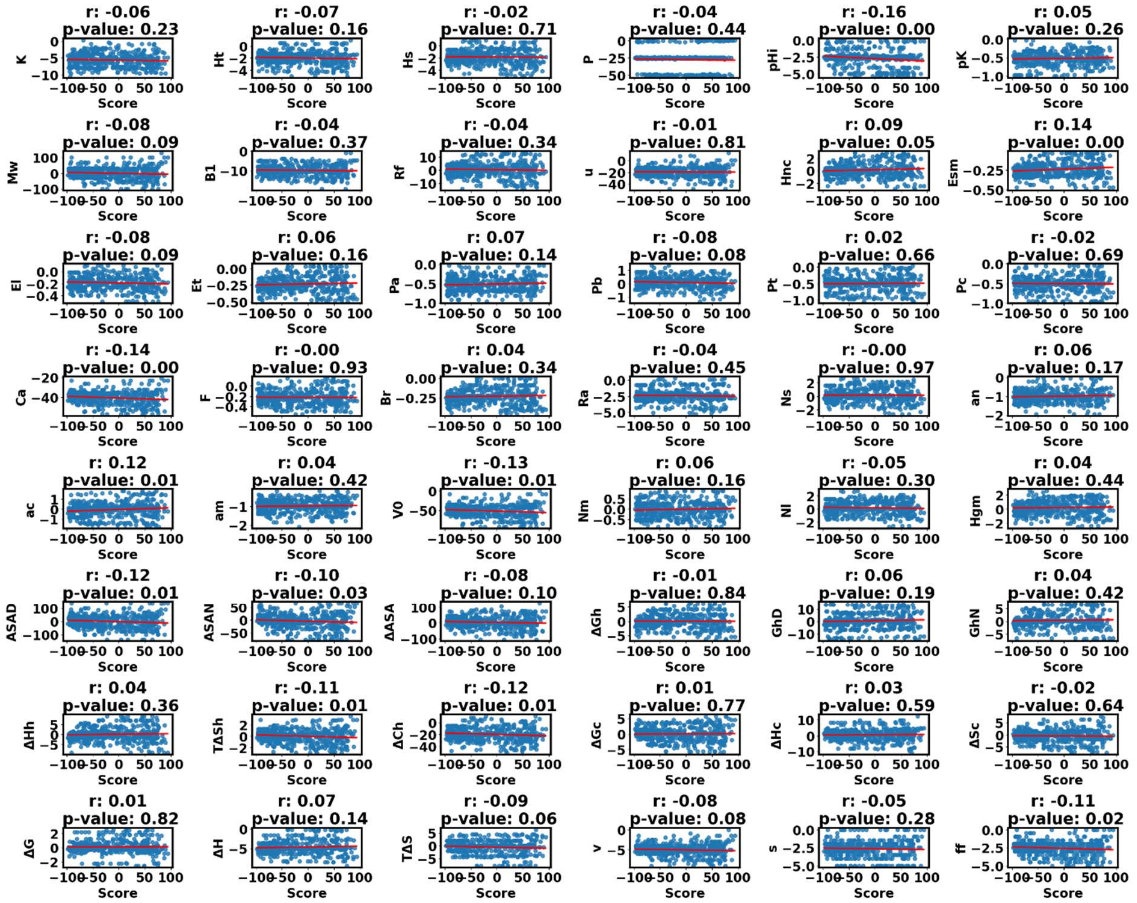


**Figure S32.** Correlation between change in 48 amino property and SNAP2 score of CYP8A1 (Non-Pathogenic). Correlation coefficient is represented by ‘r’, more closer to +1 represents positive correlation, more closer to -1 represents negative correlation, and 0 means no correlation.

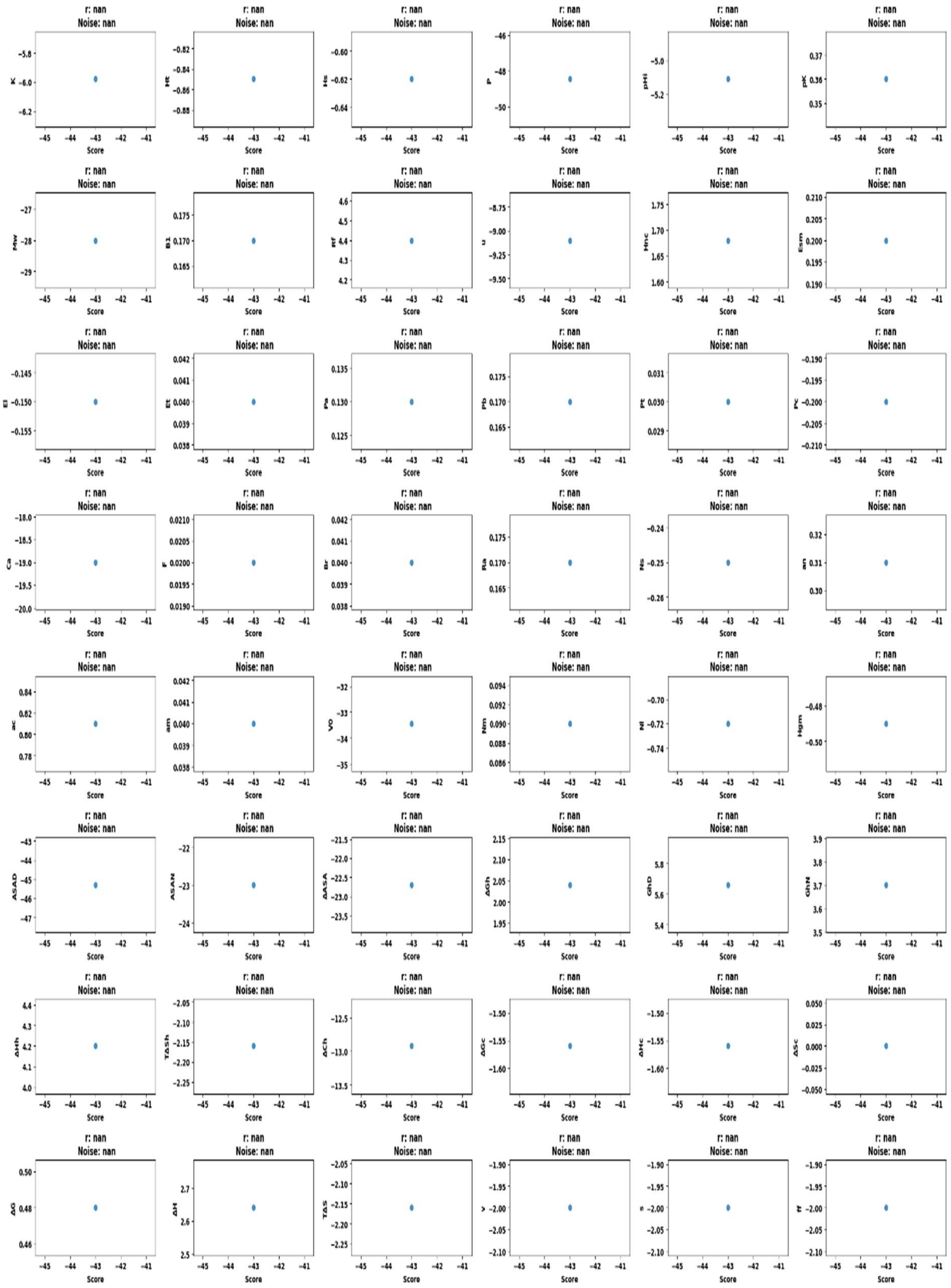


**Figure S33.** Correlation between change in 48 amino property and SNAP2 score of CYP8A1 (Pathogenic). Correlation coefficient is represented by 'r', more closer to +1 represents positive correlation, more closer to -1 represents negative correlation, and 0 means no correlation.

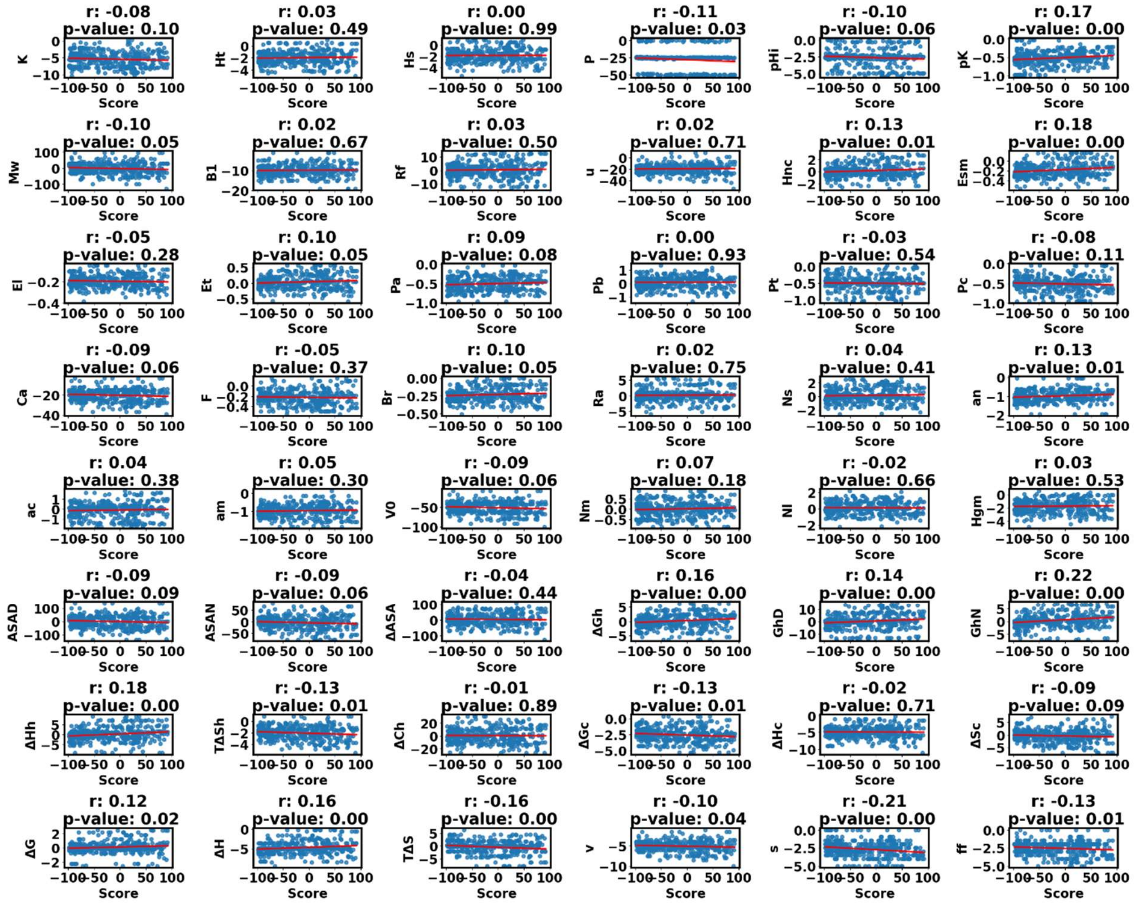

**Figure S34.** Correlation between change in 48 amino property and SNAP2 score of CYP11A1 (Non-Pathogenic). Correlation coefficient is represented by ‘r’, more closer to +1 represents positive correlation, more closer to -1 represents negative correlation, and 0 means no correlation.

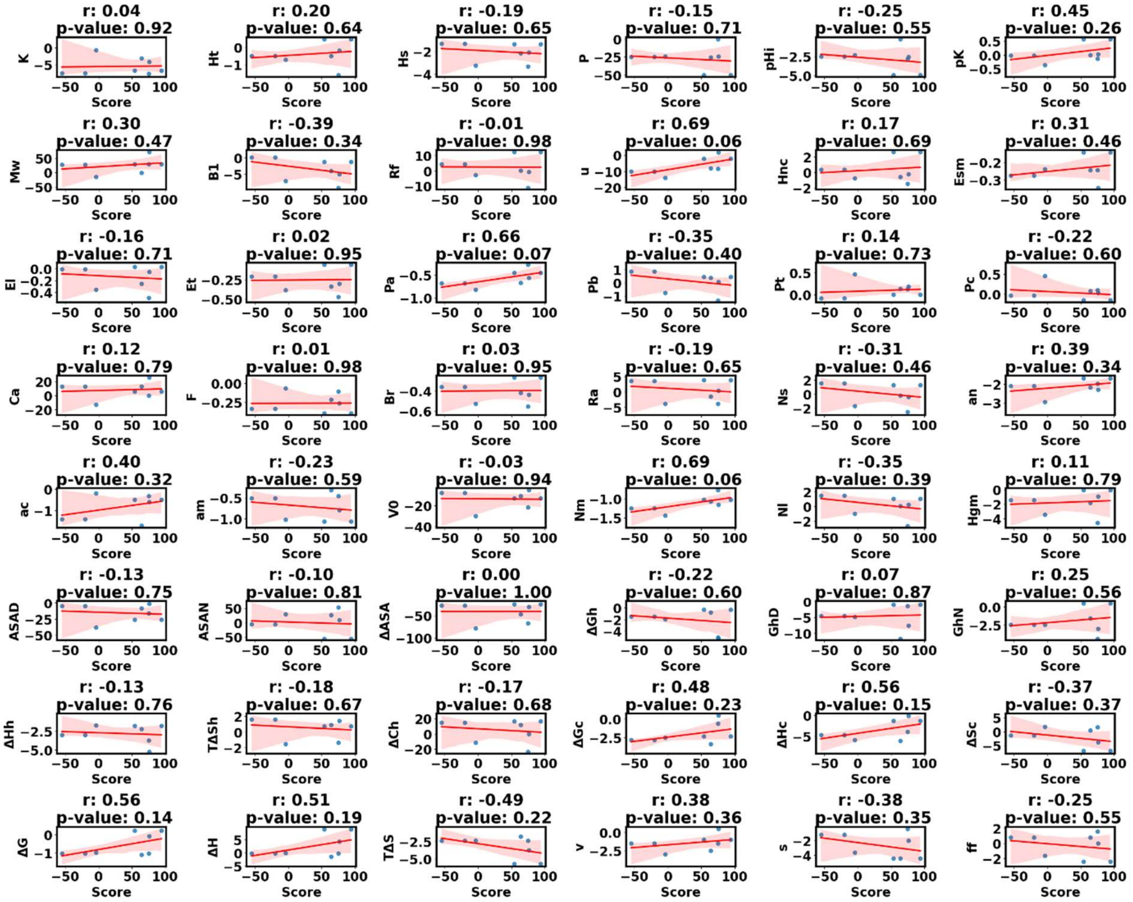


**Figure S35.** Correlation between change in 48 amino property and SNAP2 score of CYP11A1 (Pathogenic). Correlation coefficient is represented by ‘r’, more closer to +1 represents positive correlation, more closer to -1 represents negative correlation, and 0 means no correlation.

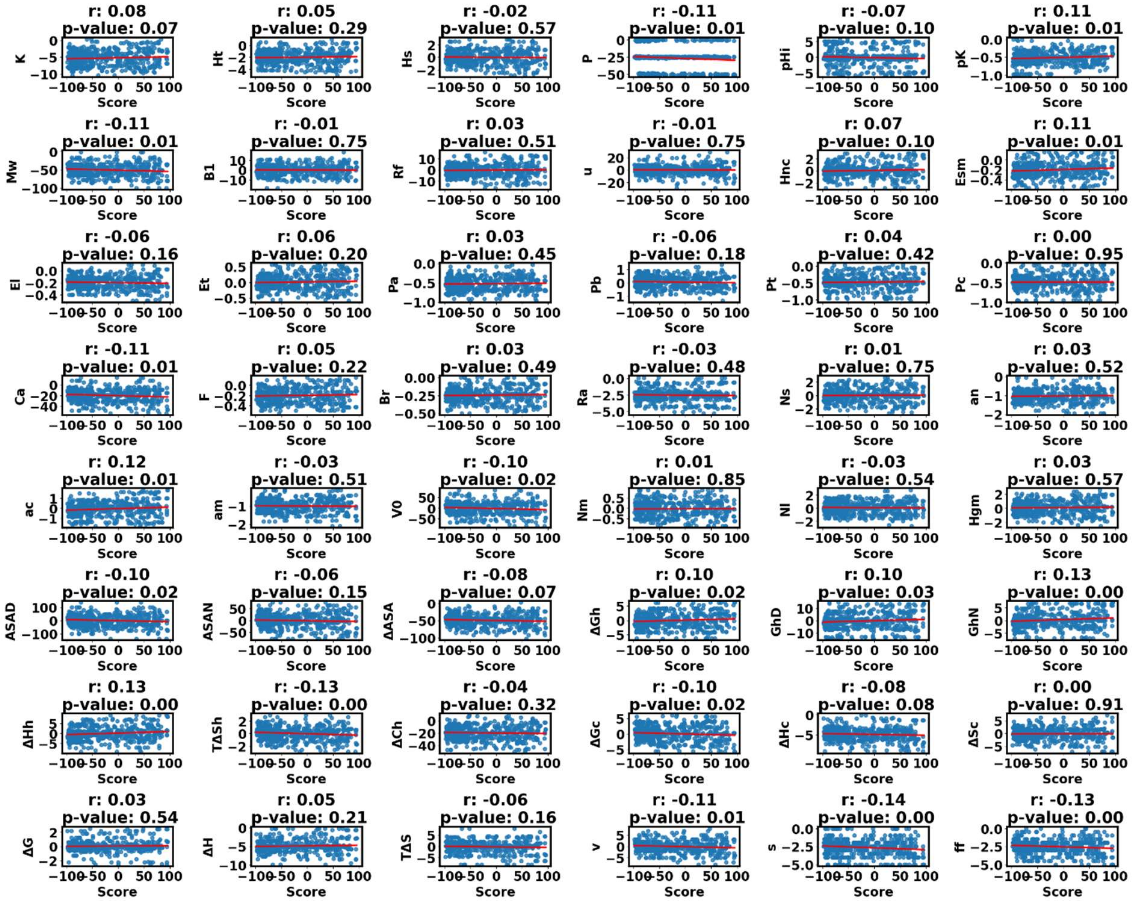


**Figure S36.** Correlation between change in 48 amino property and SNAP2 score of CYP11B1 (Non-Pathogenic). Correlation coefficient is represented by 'r', more closer to +1 represents positive correlation, more closer to -1 represents negative correlation, and 0 means no correlation.

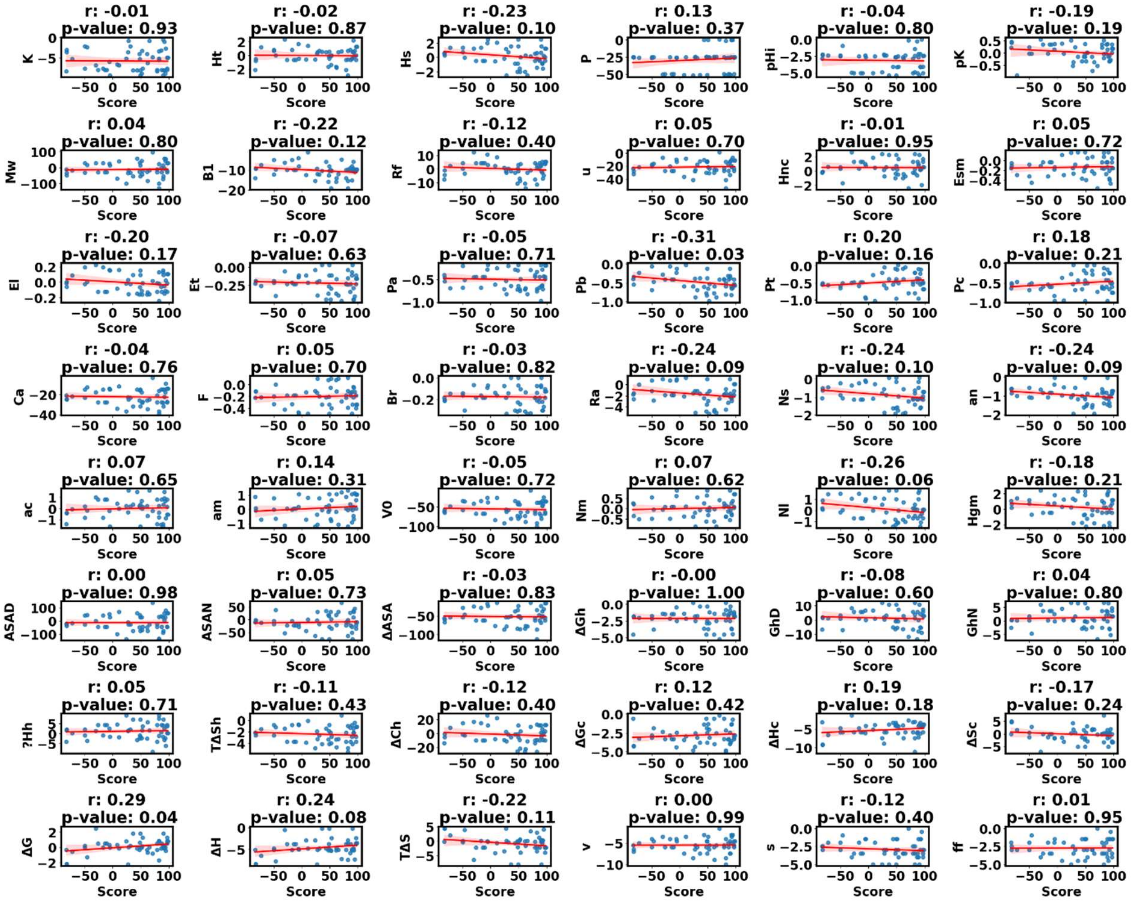


**Figure S37.** Correlation between change in 48 amino property and SNAP2 score of CYP11B1 (Pathogenic). Correlation coefficient is represented by 'r', more closer to +1 represents positive correlation, more closer to -1 represents negative correlation, and 0 means no correlation.

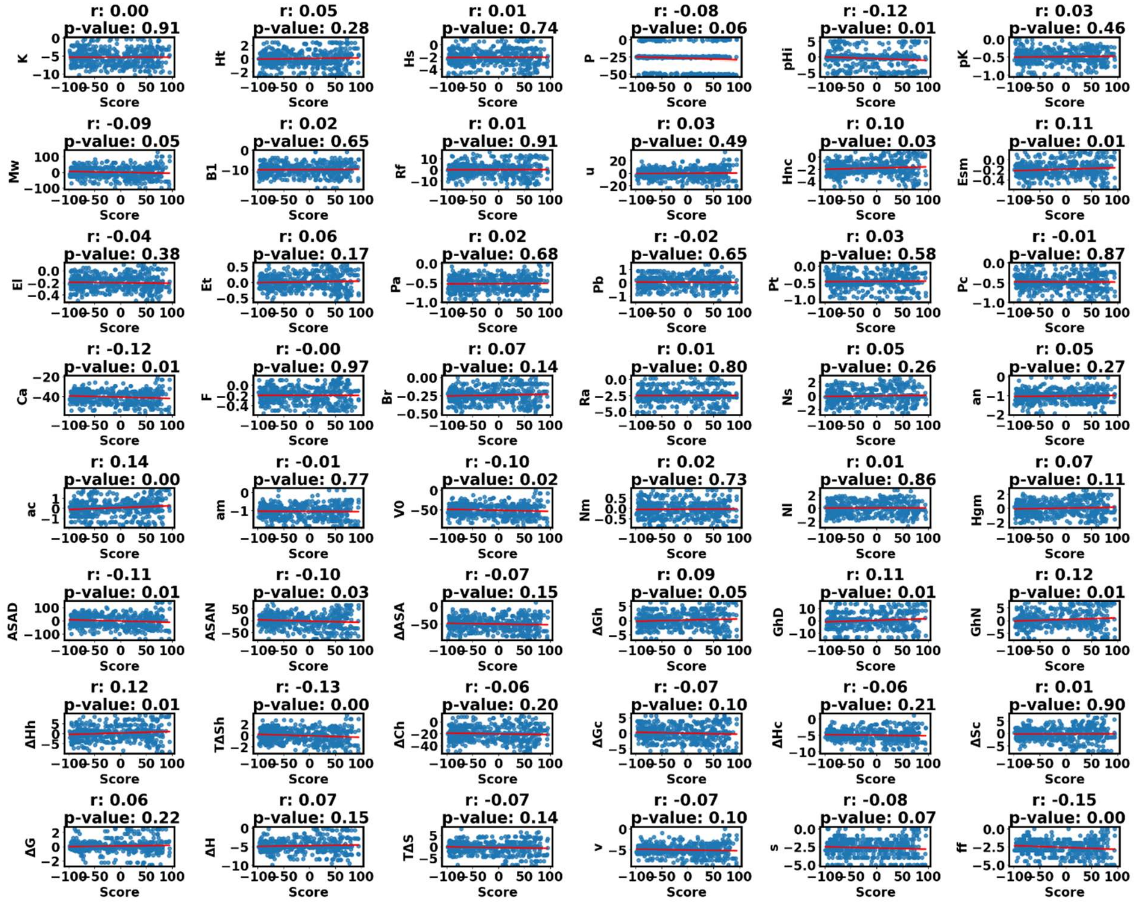


**Figure S38.** Correlation between change in 48 amino property and SNAP2 score of CYP11B2 (Non-Pathogenic). Correlation coefficient is represented by 'r', more closer to +1 represents positive correlation, more closer to -1 represents negative correlation, and 0 means no correlation.

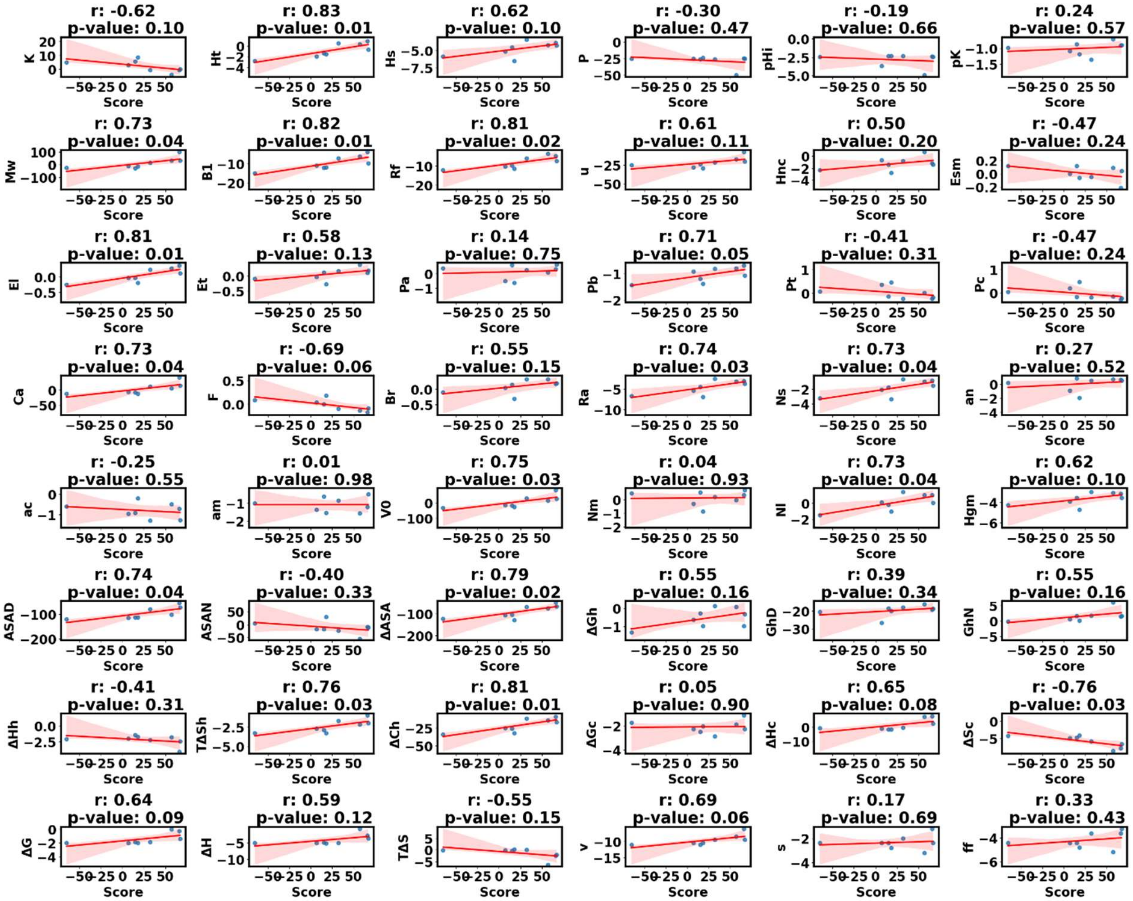


**Figure S39.** Correlation between change in 48 amino property and SNAP2 score of CYP11B2 (Pathogenic). Correlation coefficient is represented by 'r', more closer to +1 represents positive correlation, more closer to -1 represents negative correlation, and 0 means no correlation.

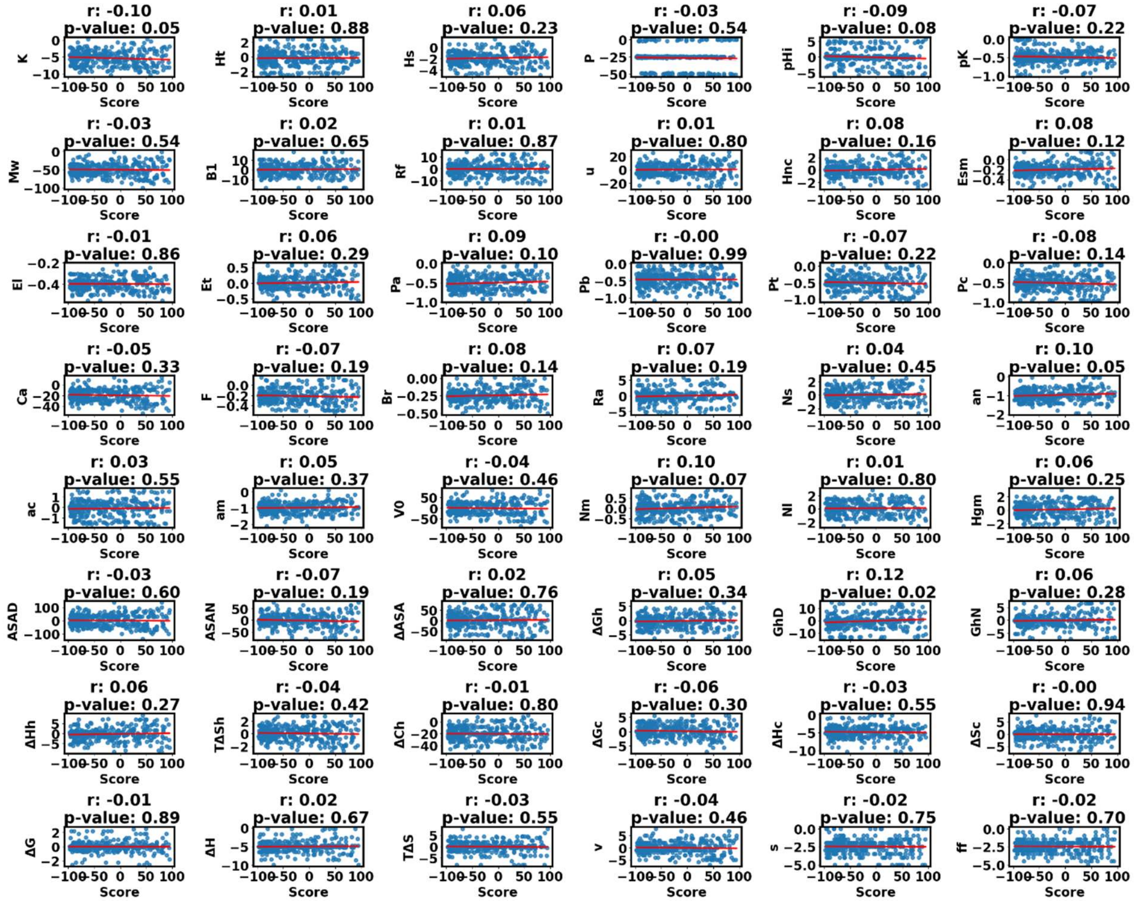


**Figure S40.** Correlation between change in 48 amino property and SNAP2 score of CYP17A1 (Non-Pathogenic). Correlation coefficient is represented by 'r', more closer to +1 represents positive correlation, more closer to -1 represents negative correlation, and 0 means no correlation.

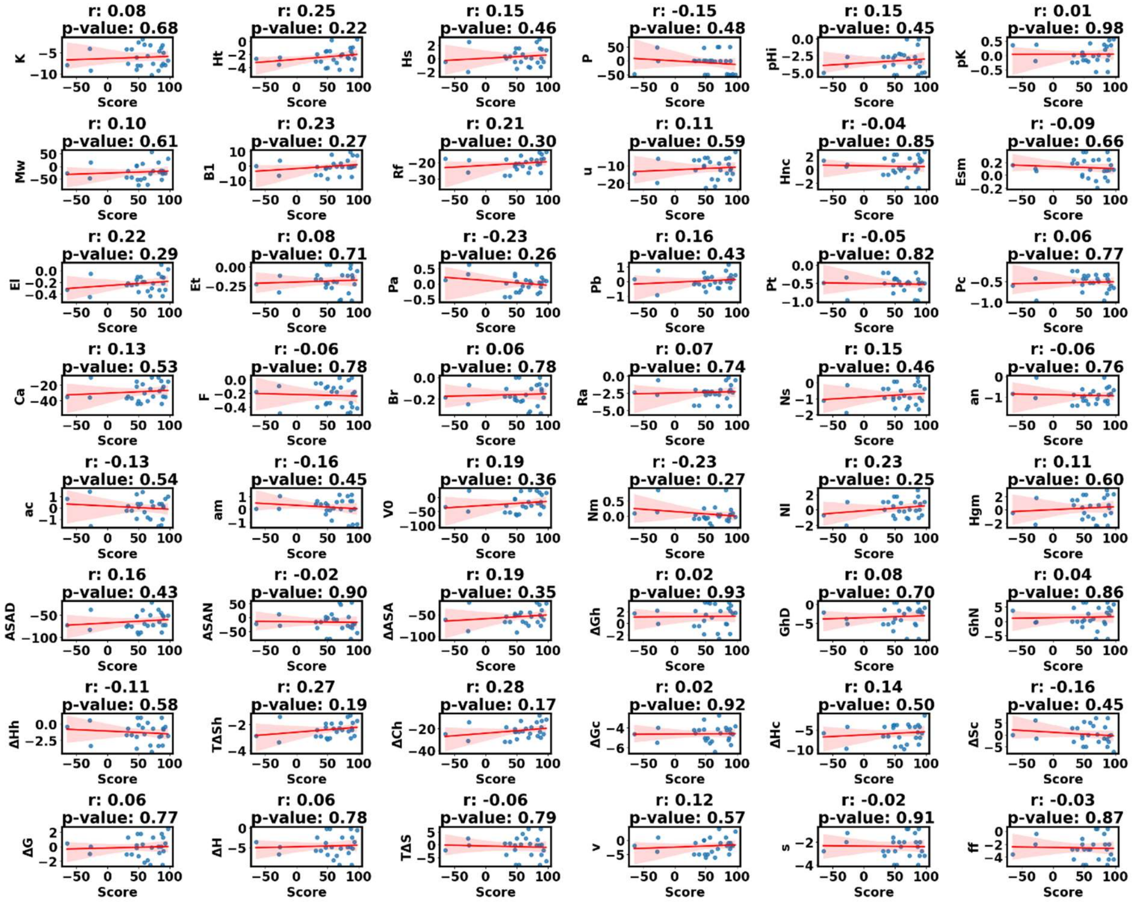

**Figure S41.** Correlation between change in 48 amino property and SNAP2 score of CYP17A1 (Pathogenic). Correlation coefficient is represented by ‘r’, more closer to +1 represents positive correlation, more closer to -1 represents negative correlation, and 0 means no correlation.

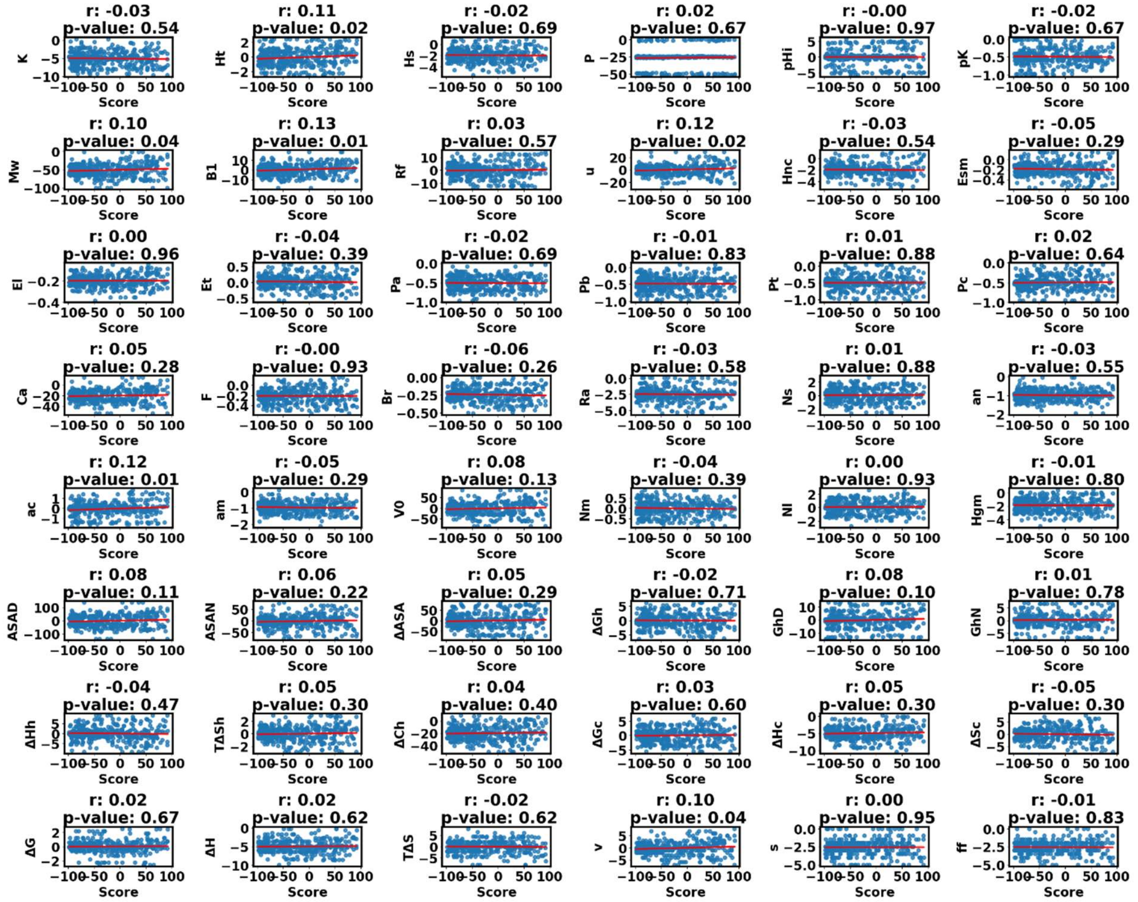

**Figure S42.** Correlation between change in 48 amino property and SNAP2 score of CYP19A1 (Non-Pathogenic). Correlation coefficient is represented by ‘r’, more closer to +1 represents positive correlation, more closer to -1 represents negative correlation, and 0 means no correlation.

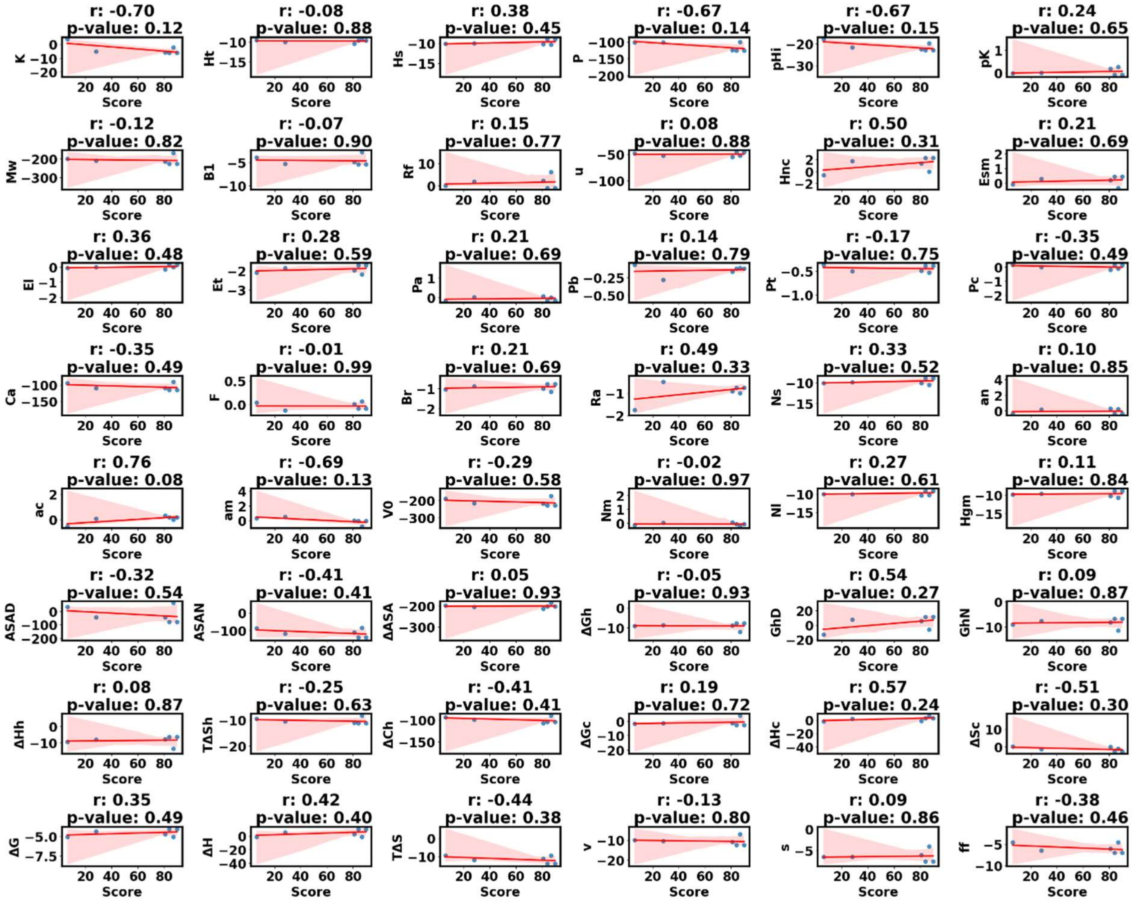

**Figure S43.** Correlation between change in 48 amino property and SNAP2 score of CYP19A1 (Pathogenic). Correlation coefficient is represented by ‘r’, more closer to +1 represents positive correlation, more closer to -1 represents negative correlation, and 0 means no correlation.

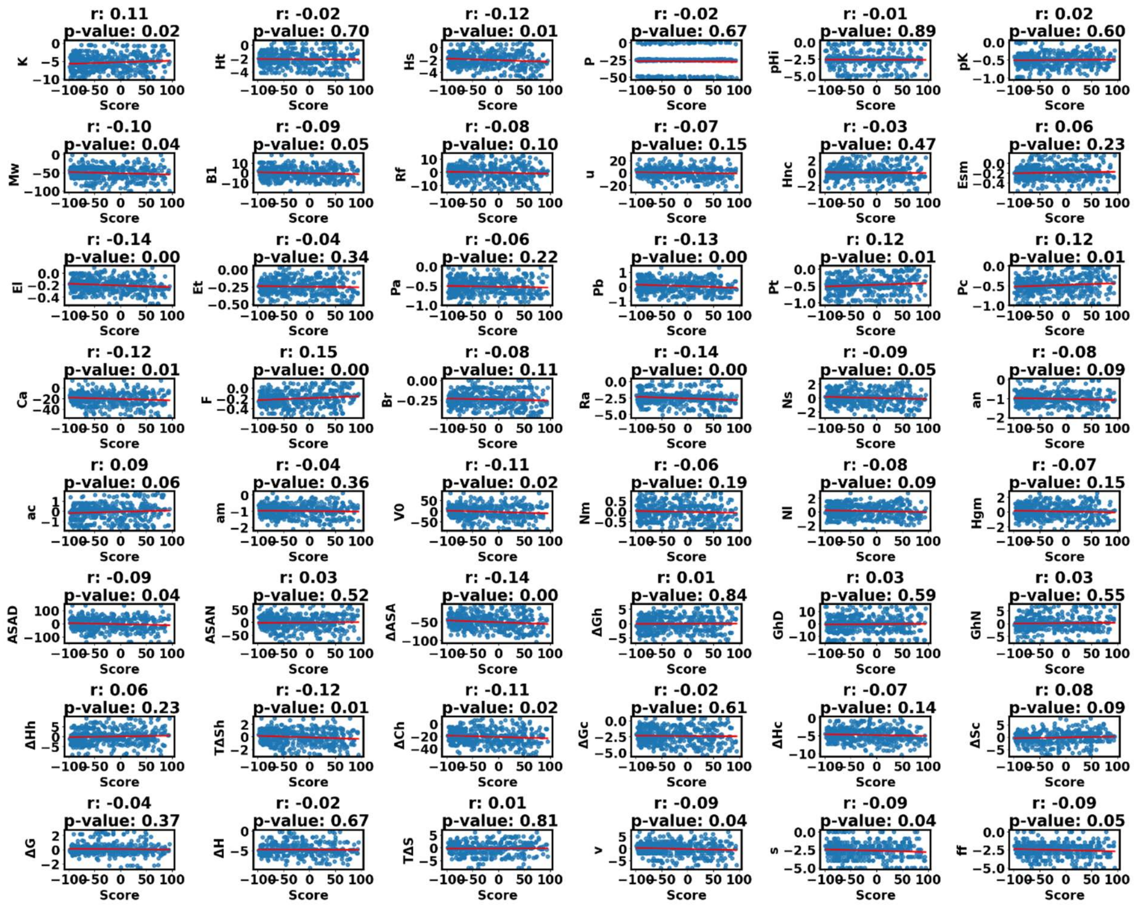

**Figure S44.** Correlation between change in 48 amino property and SNAP2 score of CYP21A2 (Non-Pathogenic). Correlation coefficient is represented by ‘r’, more closer to +1 represents positive correlation, more closer to -1 represents negative correlation, and 0 means no correlation.

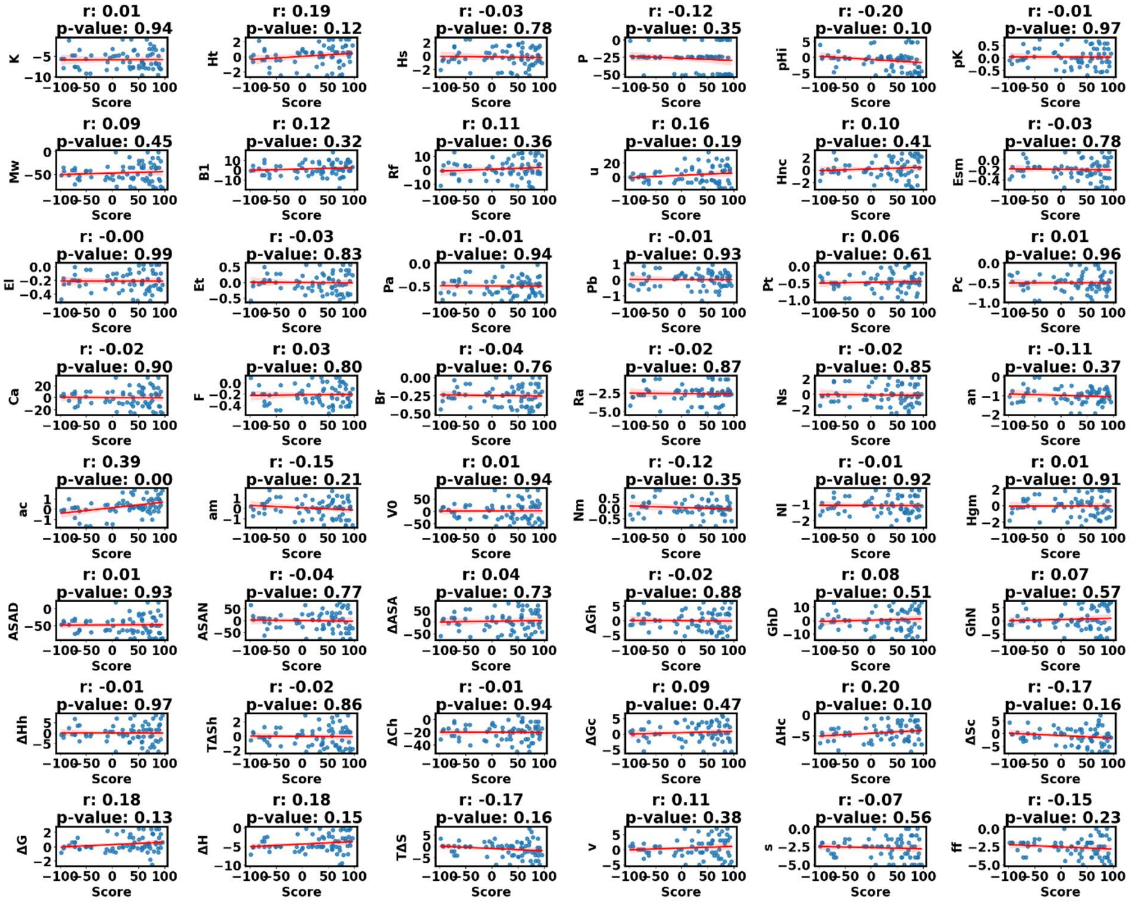

**Figure S45.** Correlation between change in 48 amino property and SNAP2 score of CYP21A2 (Pathogenic). Correlation coefficient is represented by 'r', more closer to +1 represents positive correlation, more closer to -1 represents negative correlation, and 0 means no correlation.

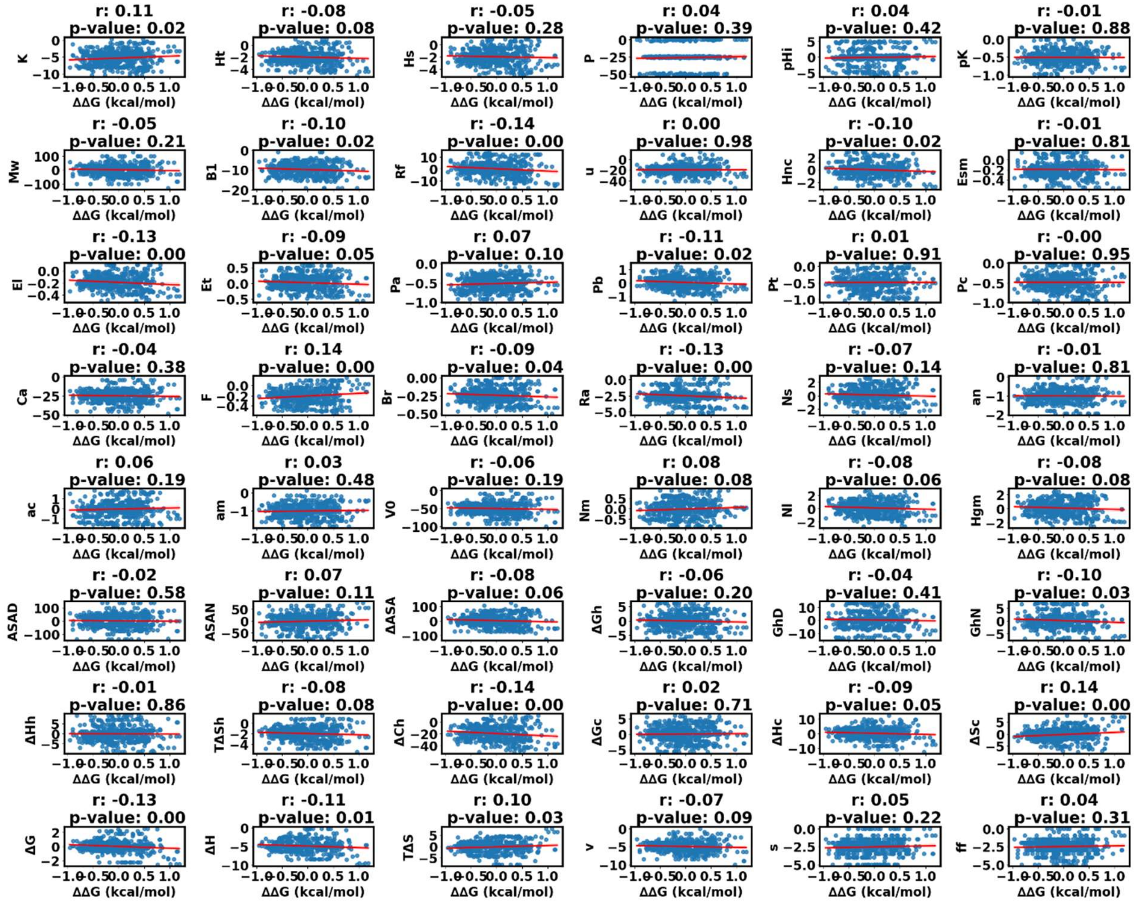

**Figure S46.** Correlation between change in 48 amino property and SNPMUSIC ΔΔG value of CYP1B1(Non-Pathogenic). Correlation coefficient is represented by ‘r’, more closer to +1 represents positive correlation, more closer to -1 represents negative correlation, and 0 means no correlation.

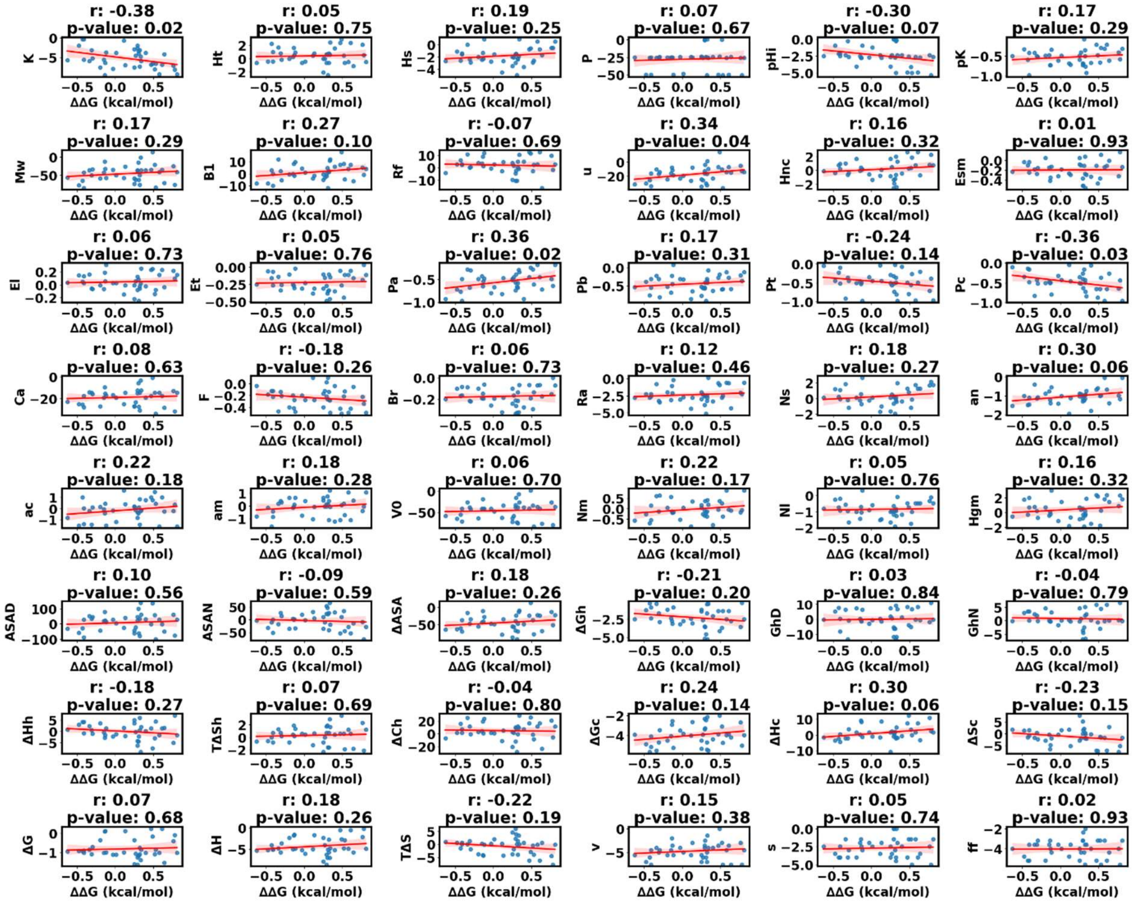


**Figure S47.** Correlation between change in 48 amino property and SNPMUSIC ΔΔG value of CYP1B1(Pathogenic). Correlation coefficient is represented by ‘r’, more closer to +1 represents positive correlation, more closer to -1 represents negative correlation, and 0 means no correlation.

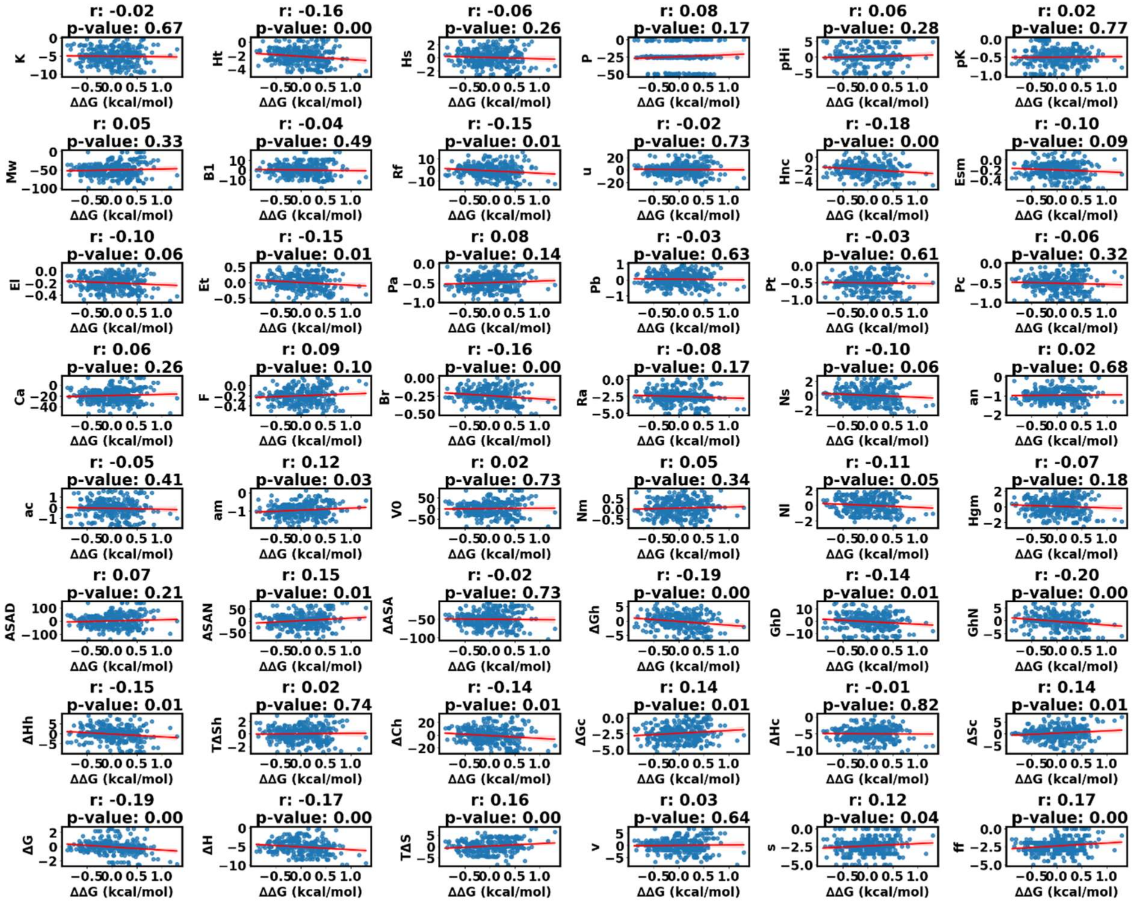

**Figure S48.** Correlation between change in 48 amino property and SNPMUSIC ΔΔG value of CYP2R1 (Non-Pathogenic). Correlation coefficient is represented by ‘r’, more closer to +1 represents positive correlation, more closer to -1 represents negative correlation, and 0 means no correlation.

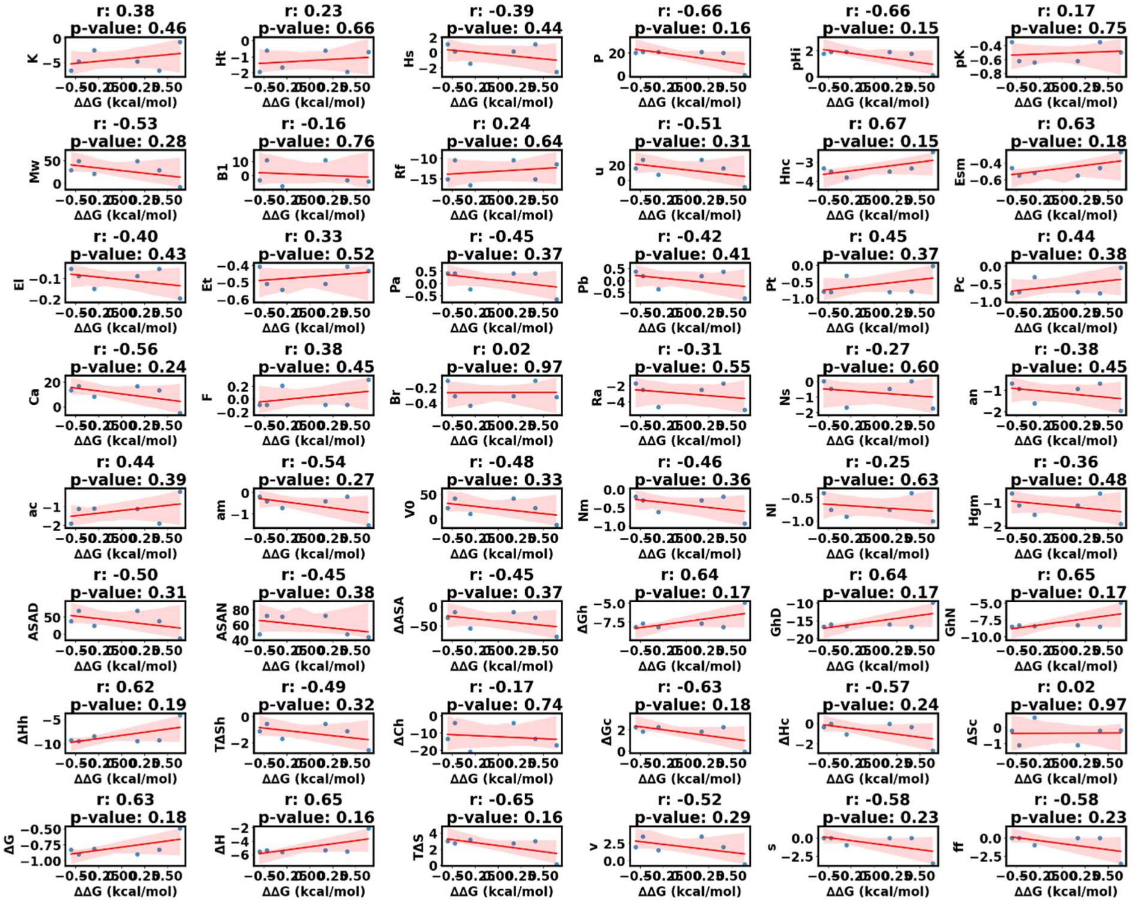

**Figure S49.** Correlation between change in 48 amino property and SNPMUSIC ΔΔG value of CYP2R1 (Pathogenic). Correlation coefficient is represented by 'r', more closer to +1 represents positive correlation, more closer to -1 represents negative correlation, and 0 means no correlation.

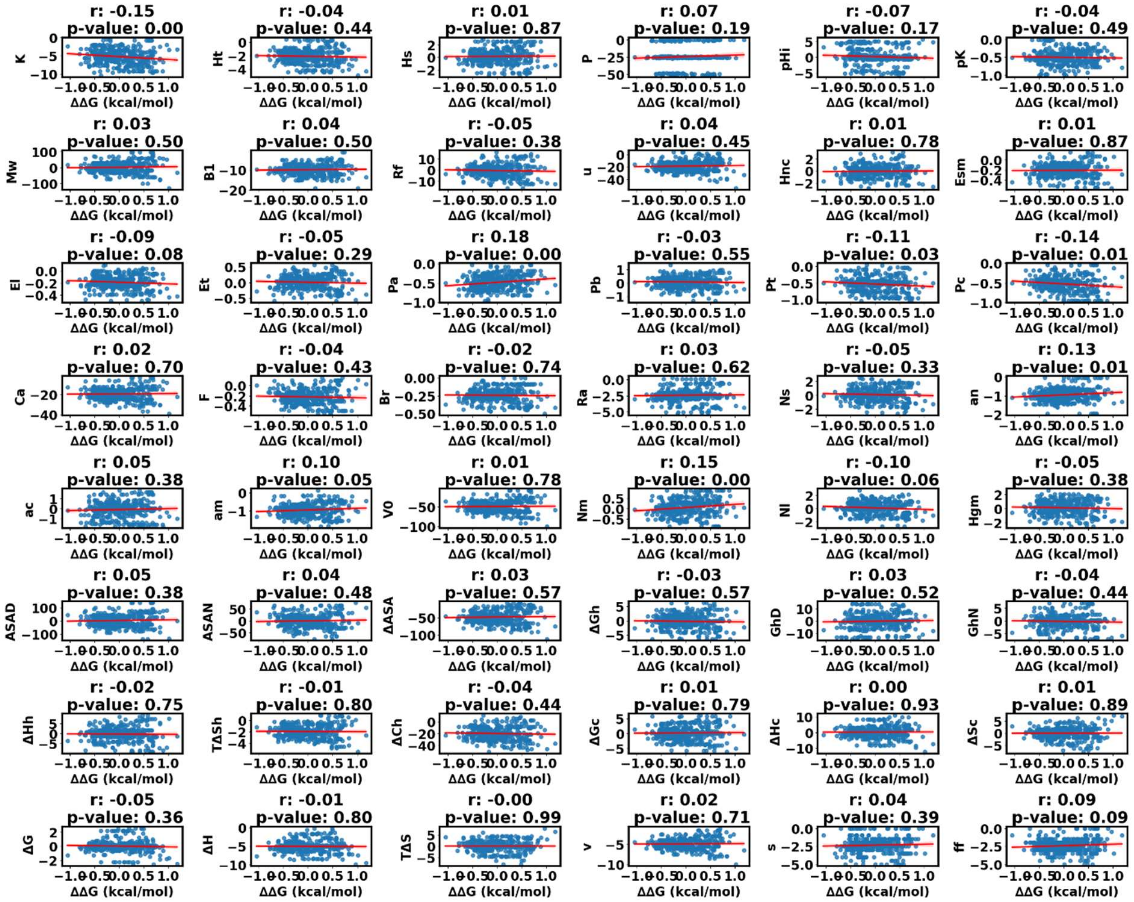


**Figure S50.** Correlation between change in 48 amino property and SNPMUSIC ΔΔG value of CY3A4 (Non-Pathogenic). Correlation coefficient is represented by ‘r’, more closer to +1 represents positive correlation, more closer to -1 represents negative correlation, and 0 means no correlation.

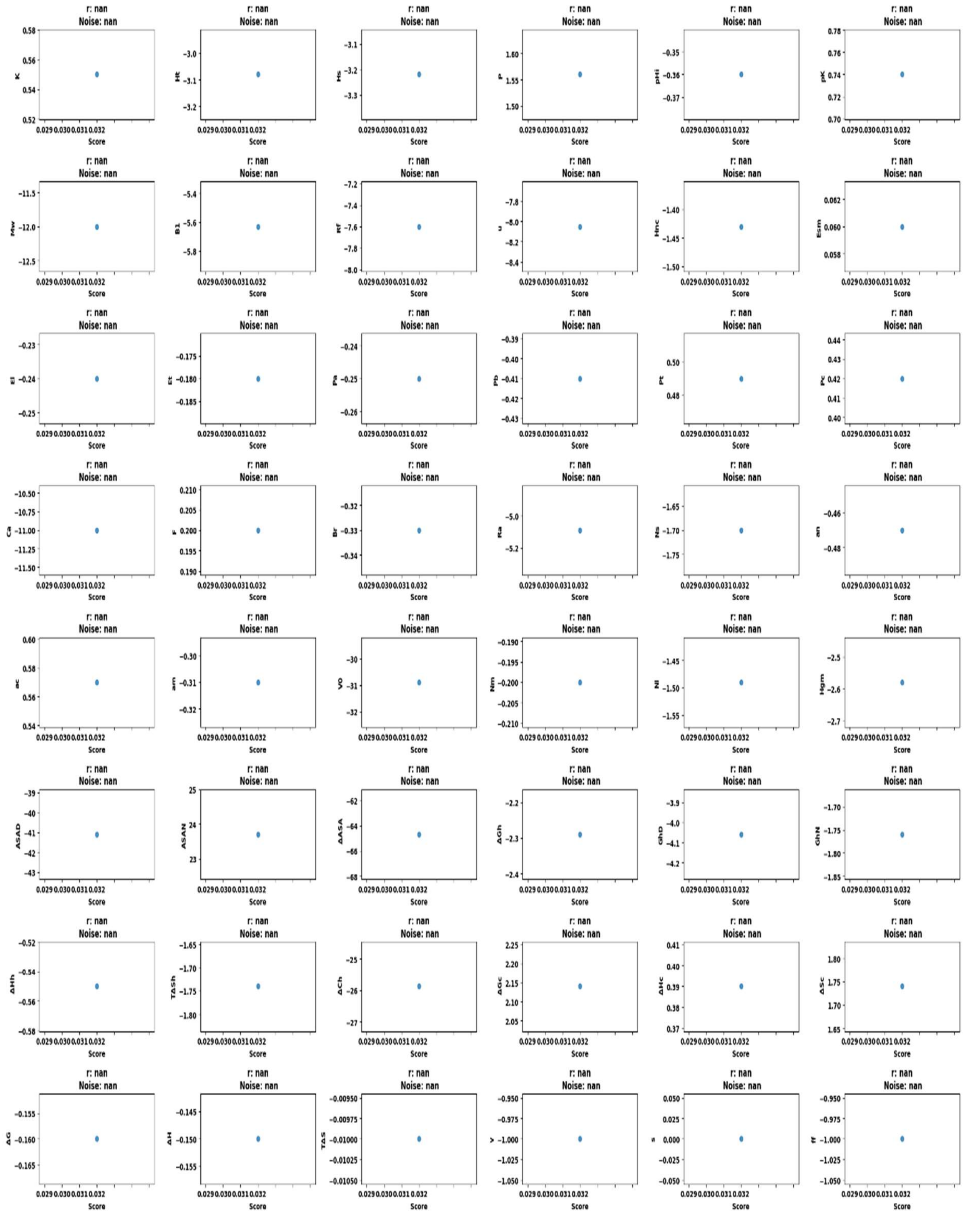

**Figure S51.** Correlation between change in 48 amino property and SNPMUSIC ΔΔG value of CYP3A4 (Pathogenic). Correlation coefficient is represented by ‘r’, more closer to +1 represents positive correlation, more closer to -1 represents negative correlation, and 0 means no correlation.

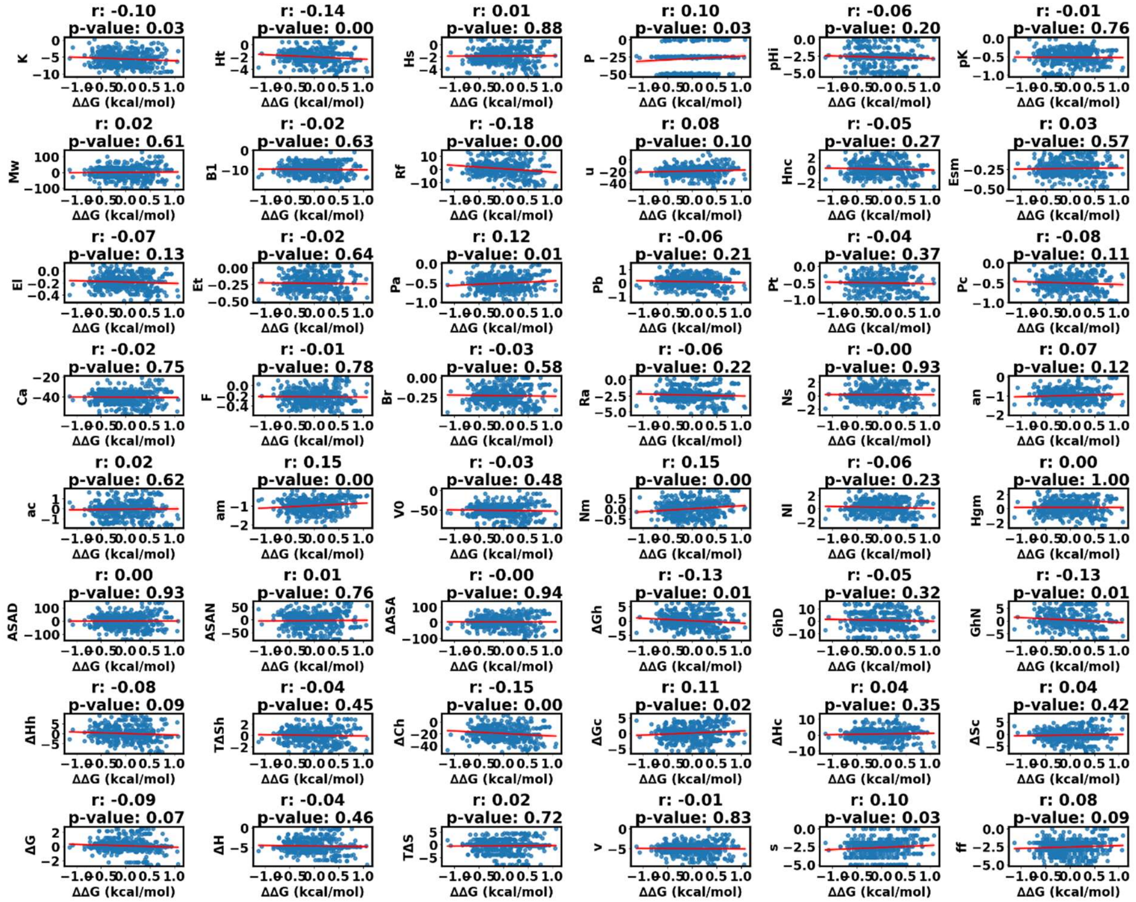

**Figure S52.** Correlation between change in 48 amino property and SNPMUSIC ΔΔG value of CYP8A1 (Non-Pathogenic). Correlation coefficient is represented by ‘r’, more closer to +1 represents positive correlation, more closer to -1 represents negative correlation, and 0 means no correlation.

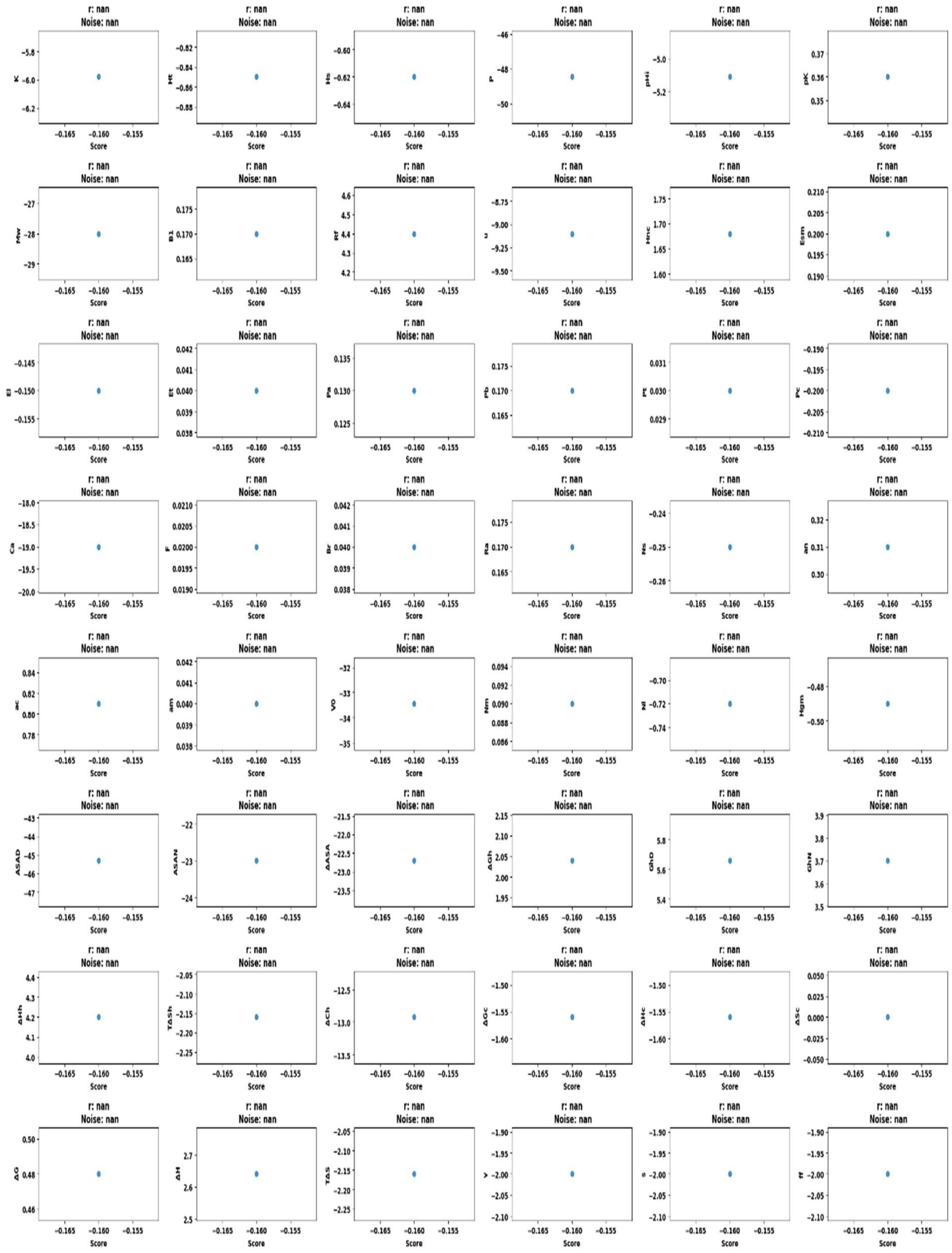


**Figure S53.** Correlation between change in 48 amino property and SNPMUSIC ΔΔG value of CYP8A1 (Pathogenic). Correlation coefficient is represented by 'r', more closer to +1 represents positive correlation, more closer to -1 represents negative correlation, and 0 means no correlation.

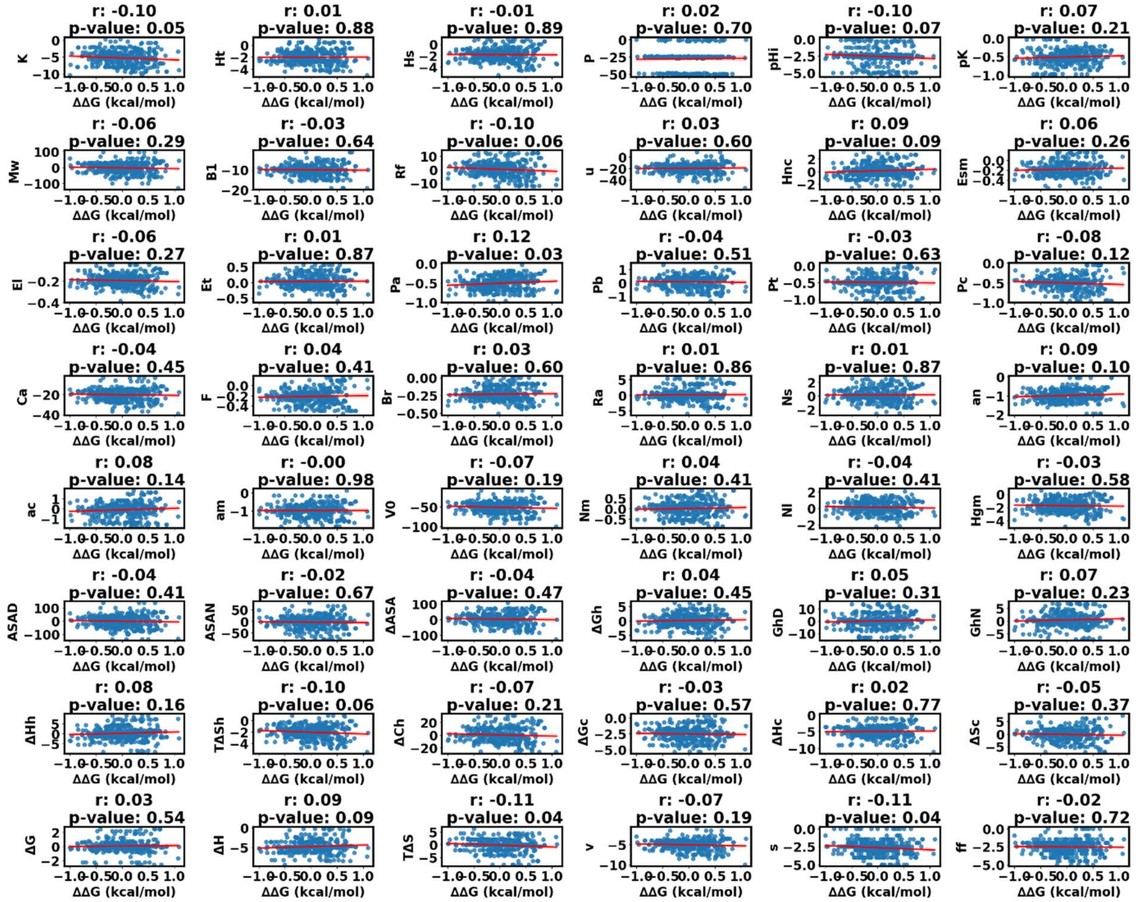

**Figure S54.** Correlation between change in 48 amino property and SNPMUSIC ΔΔG value of CYP11A1 (Non-Pathogenic). Correlation coefficient is represented by ‘r’, more closer to +1 represents positive correlation, more closer to -1 represents negative correlation, and 0 means no correlation.

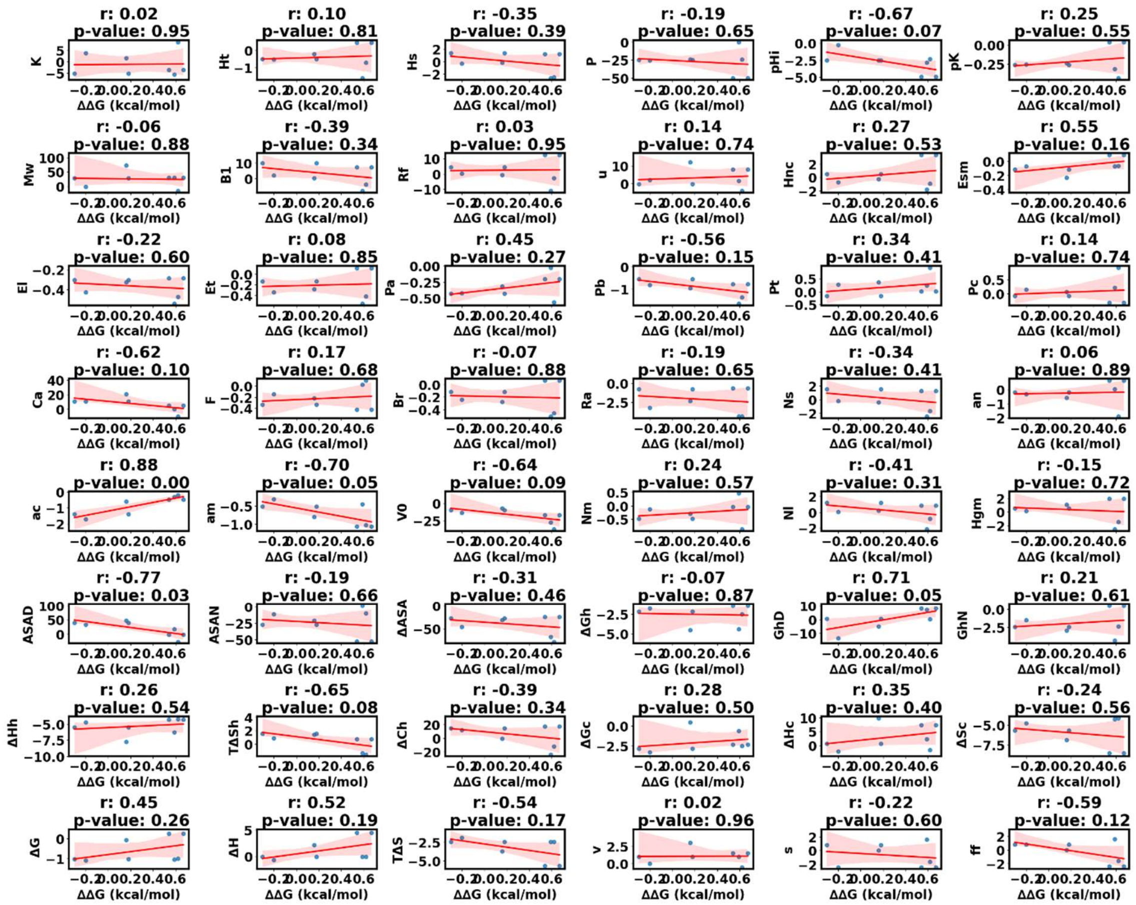


**Figure S55.** Correlation between change in 48 amino property and SNPMUSIC ΔΔG value of CYP11A1 (Pathogenic). Correlation coefficient is represented by 'r', more closer to +1 represents positive correlation, more closer to -1 represents negative correlation, and 0 means no correlation.

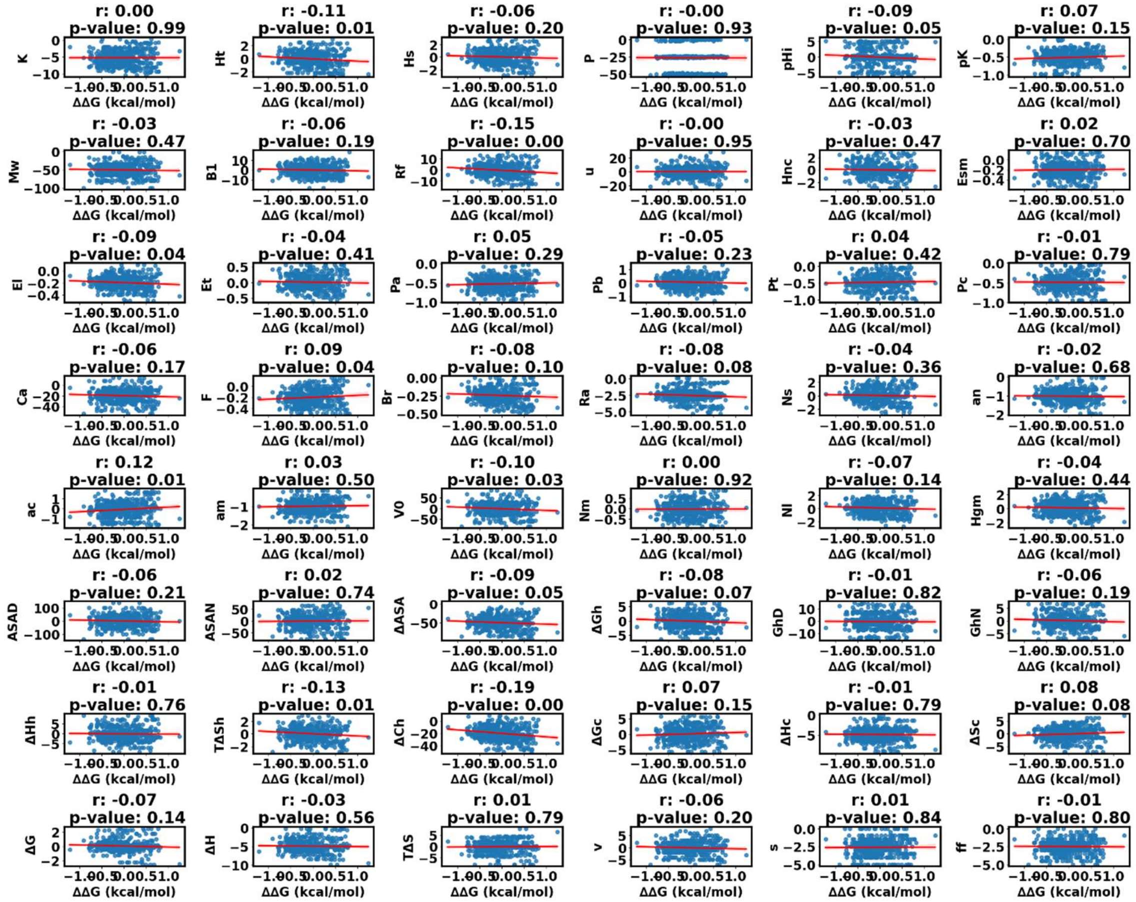

**Figure S56.** Correlation between change in 48 amino property and SNPMUSIC ΔΔG value of CYP11B1 (Non-Pathogenic). Correlation coefficient is represented by ‘r’, more closer to +1 represents positive correlation, more closer to -1 represents negative correlation, and 0 means no correlation.

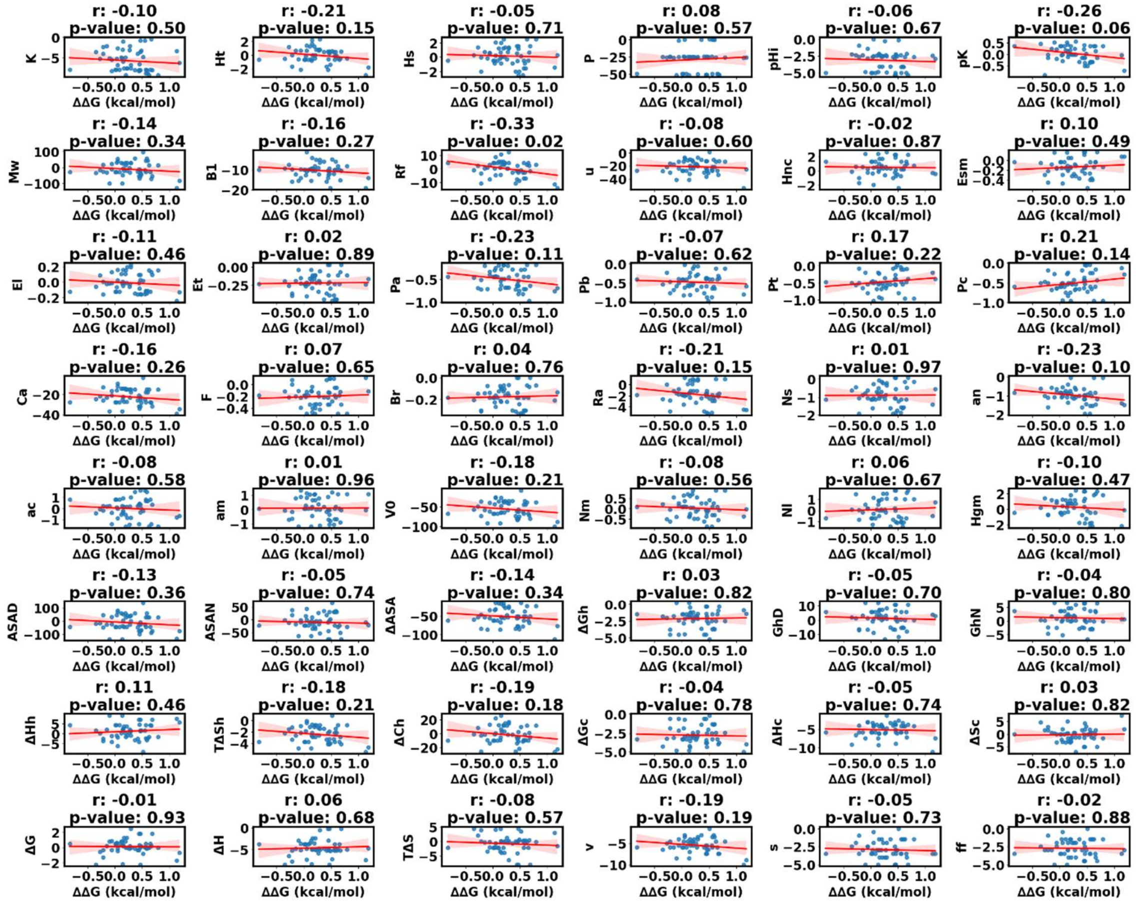

**Figure S57.** Correlation between change in 48 amino property and SNPMUSIC ΔΔG value of CYP11B1 (Pathogenic). Correlation coefficient is represented by ‘r’, more closer to +1 represents positive correlation, more closer to -1 represents negative correlation, and 0 means no correlation.

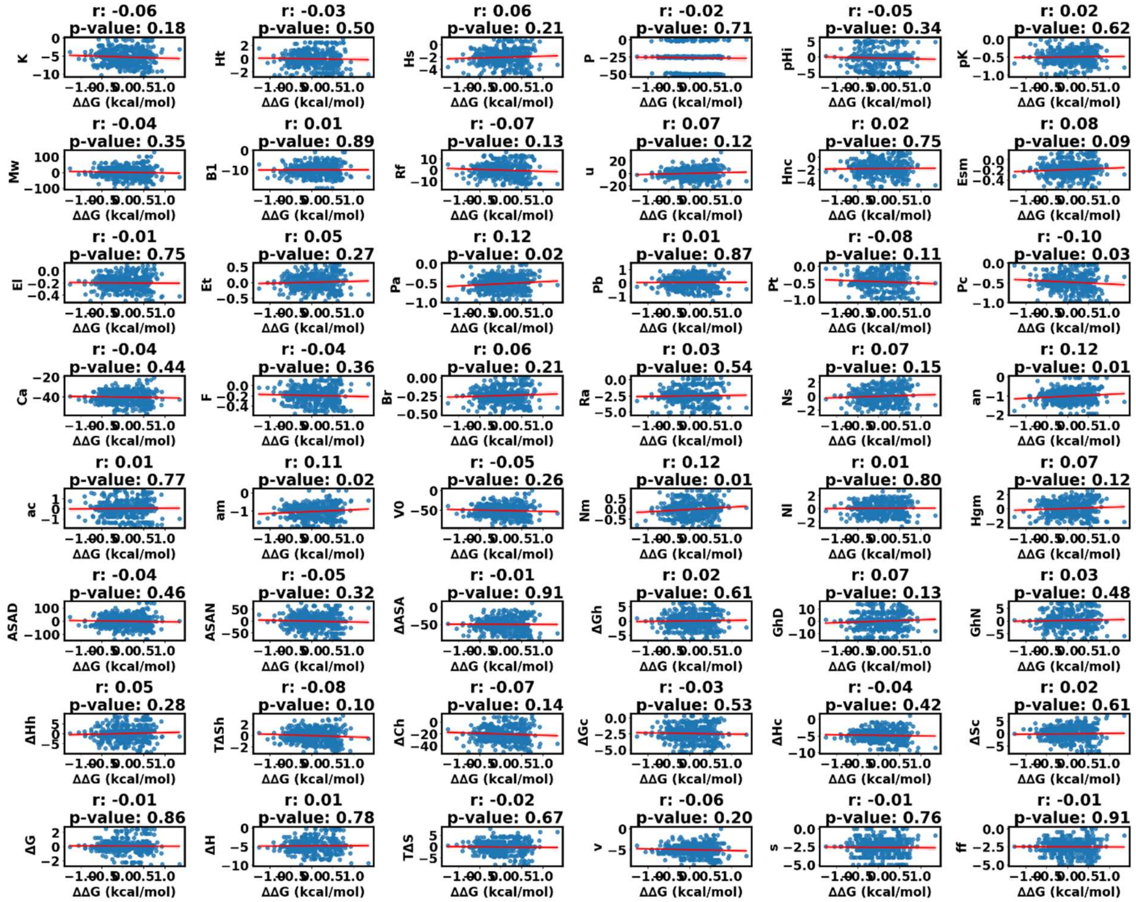

**Figure S58.** Correlation between change in 48 amino property and SNPMUSIC ΔΔG value of CYP11B2 (Non-Pathogenic). Correlation coefficient is represented by ‘r’, more closer to +1 represents positive correlation, more closer to -1 represents negative correlation, and 0 means no correlation.

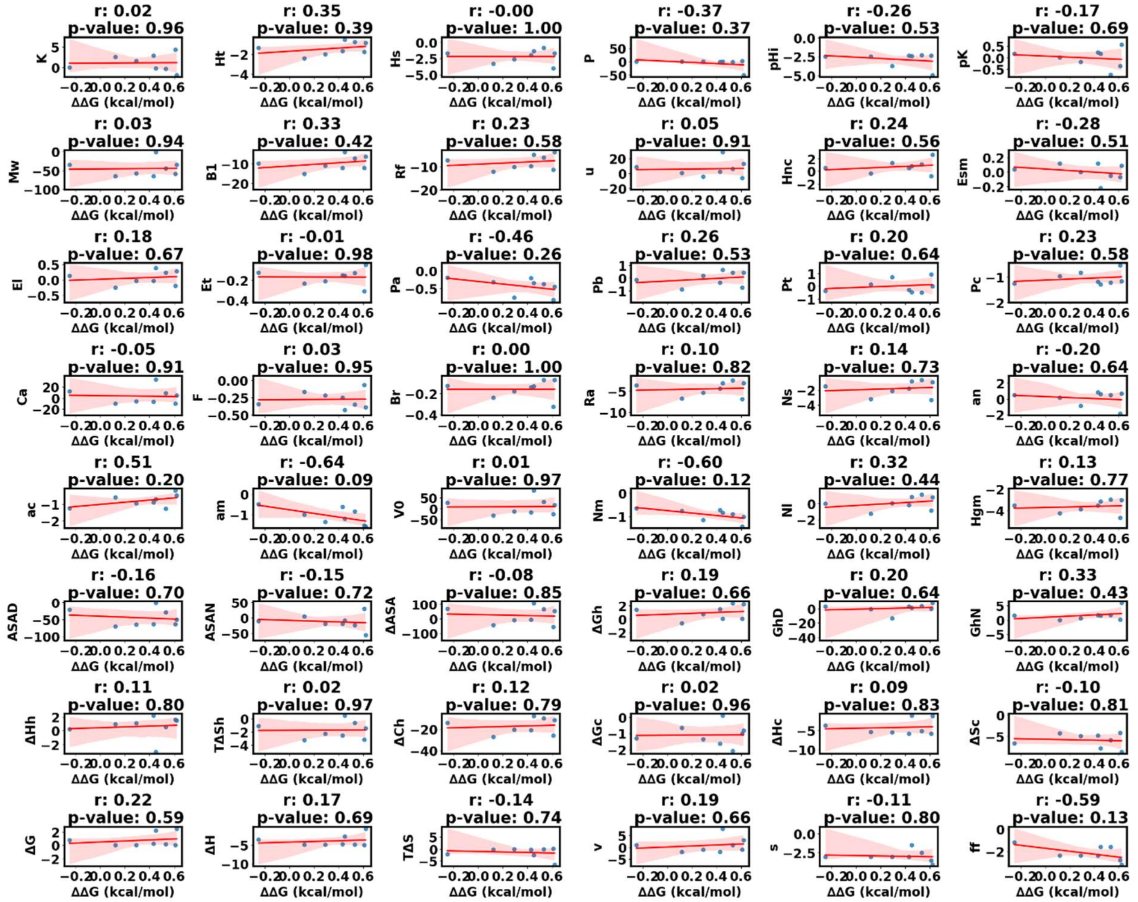

**Figure S59.** Correlation between change in 48 amino property and SNPMUSIC ΔΔG value of CYP11B2 (Pathogenic). Correlation coefficient is represented by 'r', more closer to +1 represents positive correlation, more closer to -1 represents negative correlation, and 0 means no correlation.

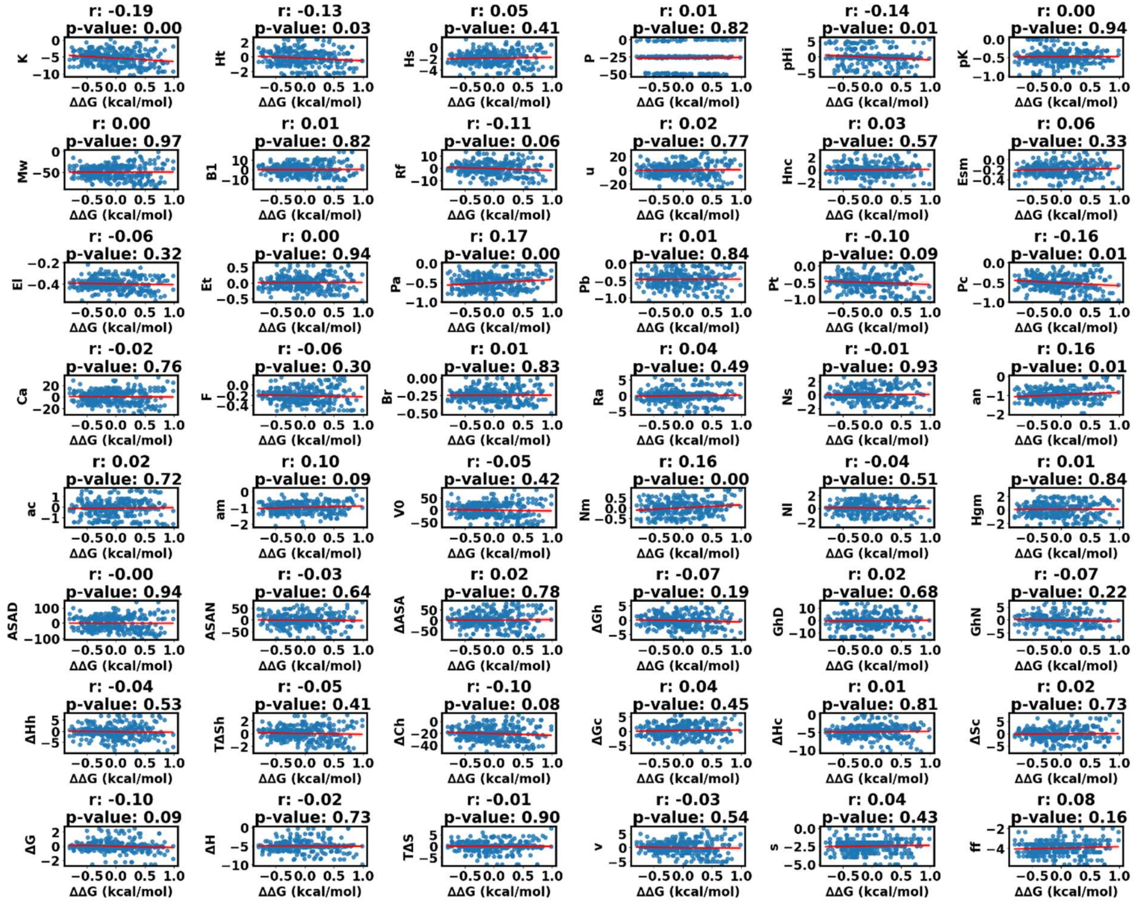

**Figure S60.** Correlation between change in 48 amino property and SNPMUSIC ΔΔG value of CYP17A1 (Non-Pathogenic). Correlation coefficient is represented by 'r', more closer to +1 represents positive correlation, more closer to -1 represents negative correlation, and 0 means no correlation.

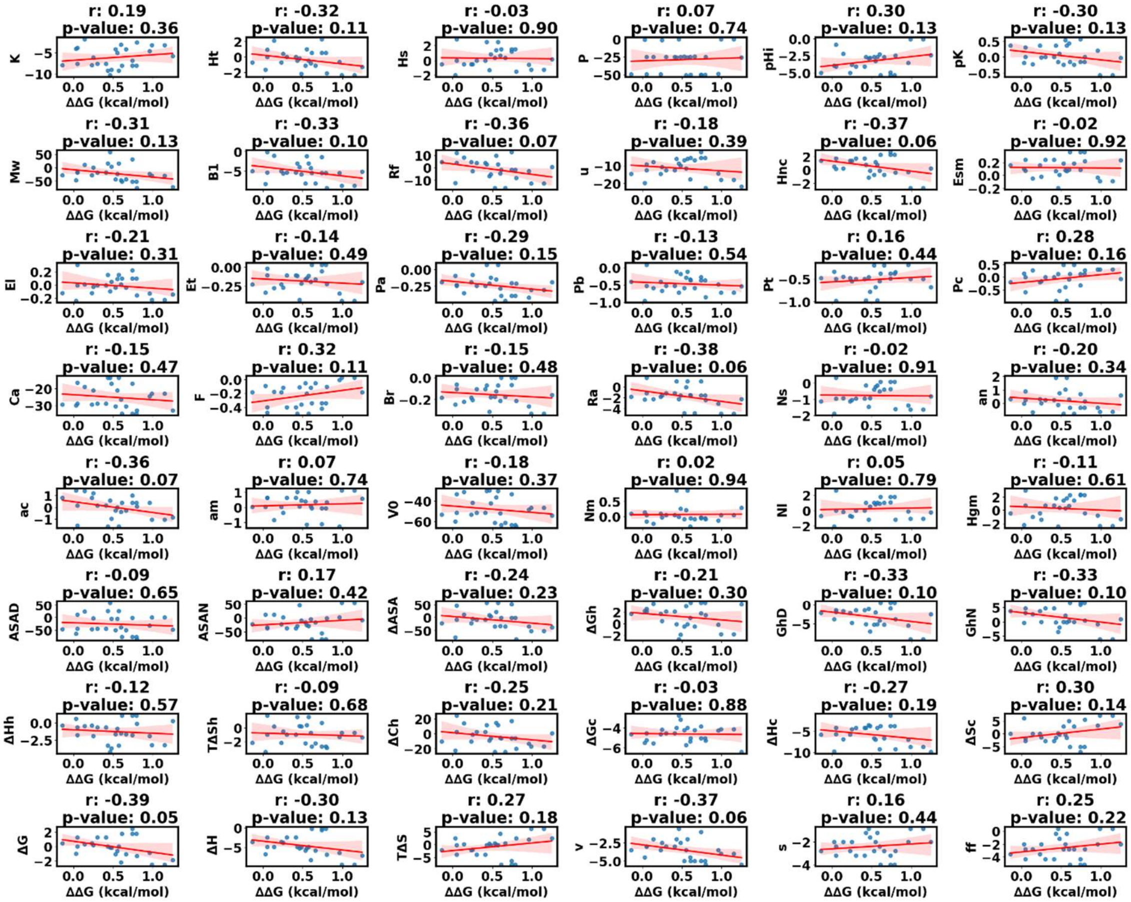


**Figure S61.** Correlation between change in 48 amino property and SNPMUSIC ΔΔG value of CYP17A1 (Pathogenic). Correlation coefficient is represented by ‘r’, more closer to +1 represents positive correlation, more closer to -1 represents negative correlation, and 0 means no correlation.

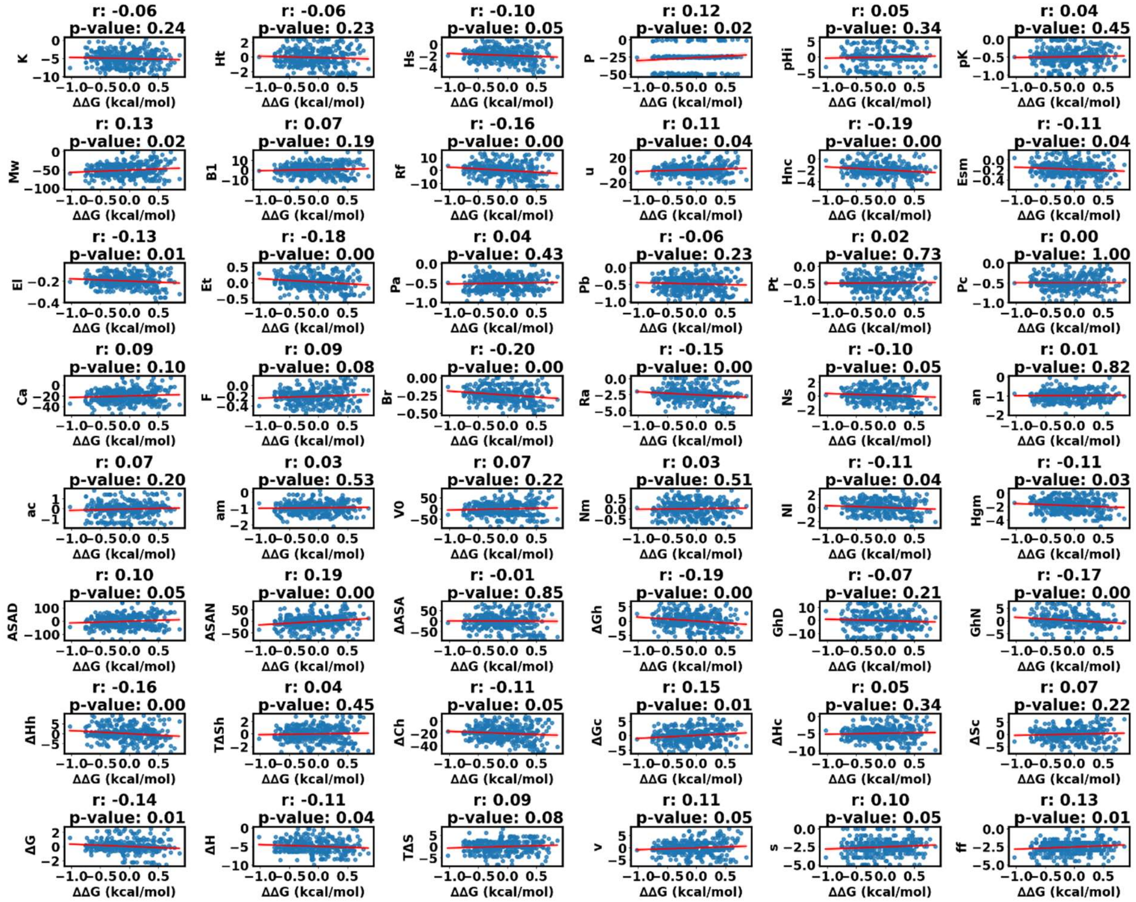


**Figure S62.** Correlation between change in 48 amino property and SNPMUSIC ΔΔG value of CYP19A1 (Non-Pathogenic). Correlation coefficient is represented by 'r', more closer to +1 represents positive correlation, more closer to -1 represents negative correlation, and 0 means no correlation.

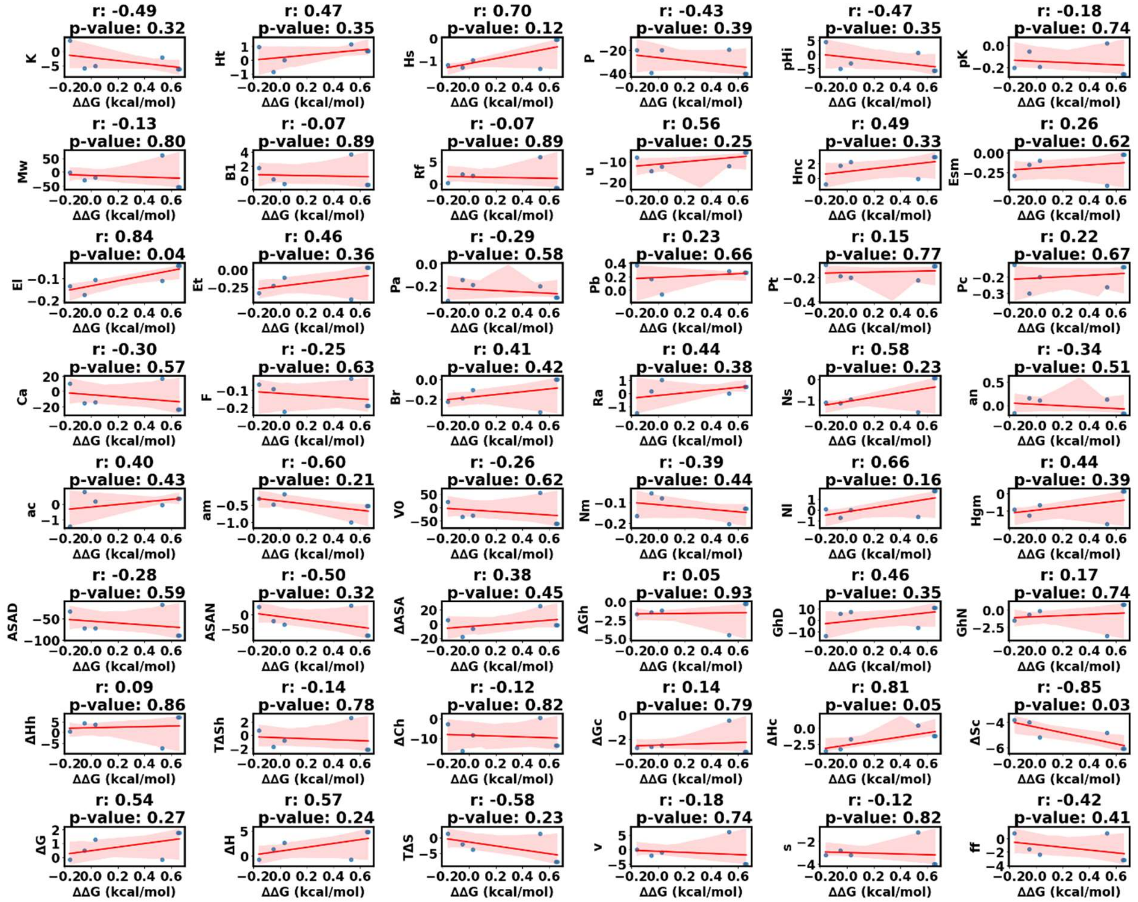

**Figure S63.** Correlation between change in 48 amino property and SNPMUSIC ΔΔG value of CYP19A1 (Pathogenic). Correlation coefficient is represented by ‘r’, more closer to +1 represents positive correlation, more closer to -1 represents negative correlation, and 0 means no correlation.

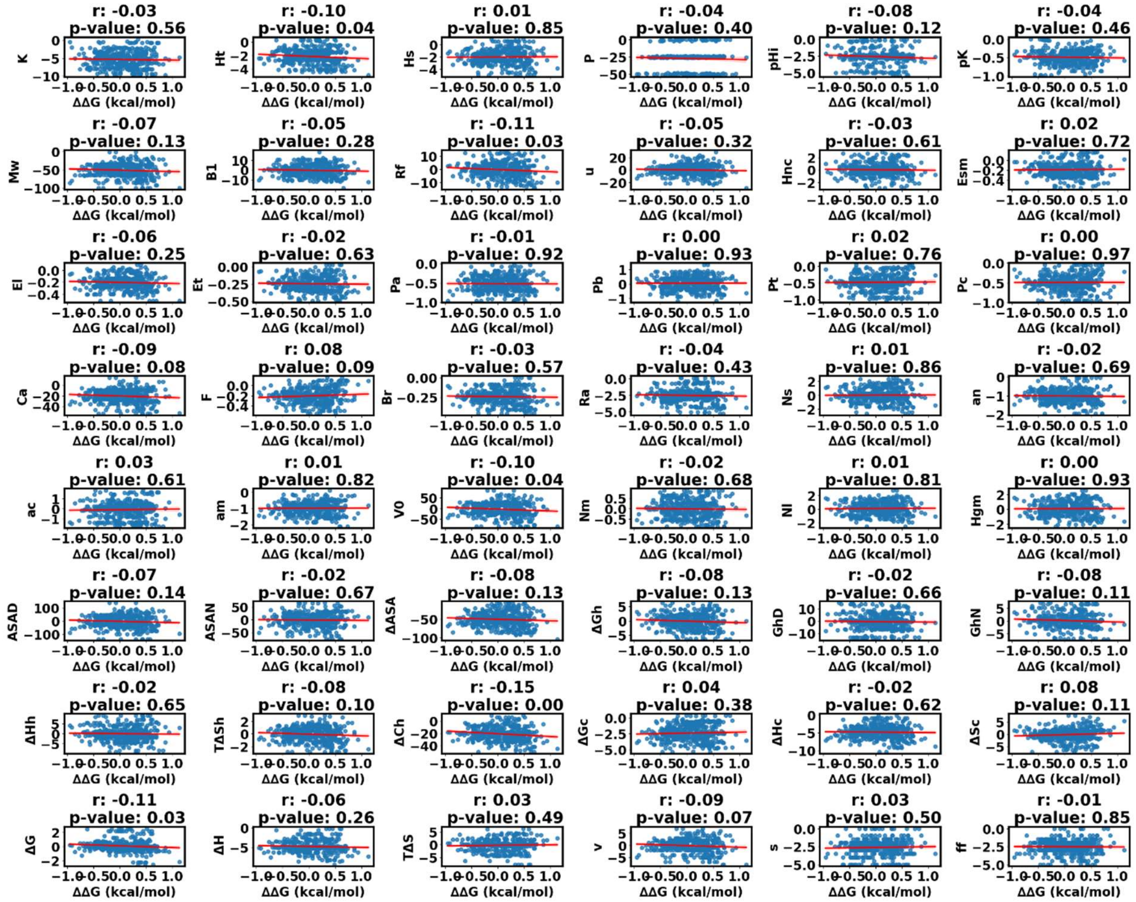


**Figure S64.** Correlation between change in 48 amino property and SNPMUSIC ΔΔG value of CYP21A2 (Non-Pathogenic). Correlation coefficient is represented by 'r', more closer to +1 represents positive correlation, more closer to -1 represents negative correlation, and 0 means no correlation.

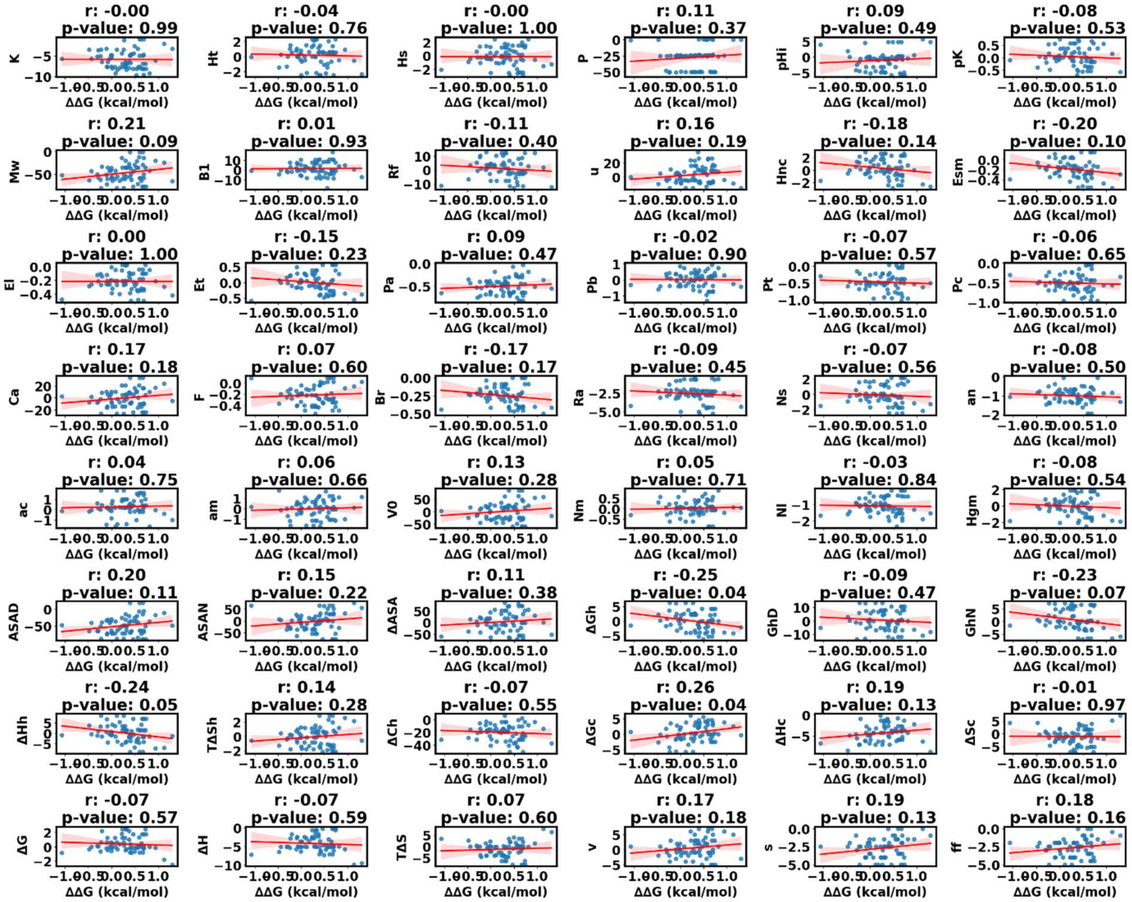


**Figure S65.** Correlation between change in 48 amino property and SNPMUSIC ΔΔG value of CYP21A2 (Pathogenic). Correlation coefficient is represented by 'r', more closer to +1 represents positive correlation, more closer to -1 represents negative correlation, and 0 means no correlation.

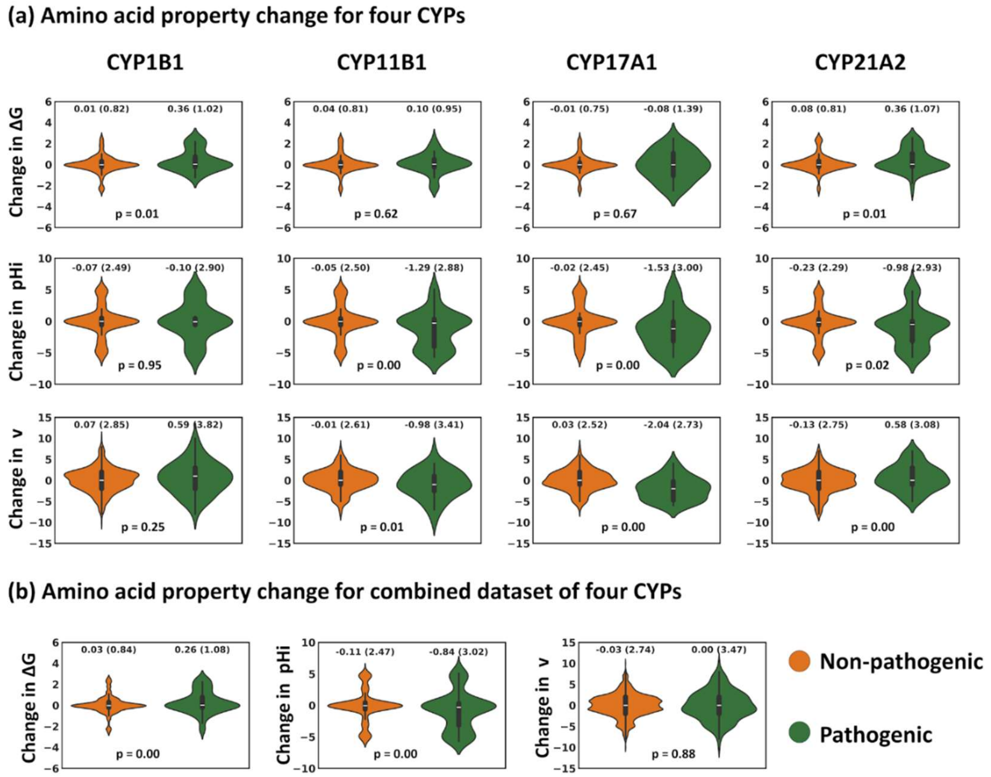


**Figure S66. Analysis of change in amino acid property between non-pathogenic and pathogenic mutations for change in ΔG, pHi, and v. (a)** Violin plots for each of the four CYPs. **(b)** Violin plots for the combined datasets of CYP1B1, CYP11B1, CYP17A1, and CYP21A2.
